# Thiamin-Diphosphate Enzymes Are an Ancient Family of Repeat Proteins

**DOI:** 10.1101/2021.03.21.436335

**Authors:** Matthew Merski, Maria Górna

## Abstract

A repeating sequence and structure pattern that is highly similar to the canonical cofactor binding motif has been identified in the thiamin-diphosphate dependent (ThDP) enzyme family. We have identified more than a thousand of these repeats in a non-redundant set (N = 58) of ThDP enzyme structures. The repeating element has a helix-turn-strand secondary structure which typically begins with an [G/A]{X(1,2)}[G/A] sequence motif with a typical length of 29 residues. The catalytically important diphosphate and aminopyrimidine interacting domains are comprised of a set of six of these repeats in a conserved architecture with a flavodoxin-like 213465 strand order. The canonical ThDP binding motif is the fourth repeat in the ThDP binding domain, while the conserved aminopyrimidine interacting glutamate is part of the second repeat in its domain. The third and fourth repeats form a contact between the functional domains, while the fifth repeat in the N-terminal domain forms an inter-chain contact. The conservation of these functional properties highlights the role of these repeats in the function and structure of this well-studied enzyme family and agrees with the principle of modular assembly in protein ancestry.

## INTRODUCTION

The origin of life on Earth at or before the beginning of the Archean period 4 billion years ago (BYA) required several events to occur: the organization of genetic information into a storage system, the formation of semi-permeable membranes, and the development of stable protein architectures^1^. It has been theorized, first by Dayhoff^2^, that small peptides formed spontaneously in the primordial Archean sea collected into oligomeric structures, possibly bound to nucleotides or similar compounds, which in turn then catalyzed additional synthesis of these oligomeric peptides, functioning as an early ribosome^3^. Fusion of these oligomeric peptide complexes into larger proteins could have given rise structures comprised of repeating sequences, which through genetic drift over a long enough time would eventually lead to complex, globular proteins^4–6^. These ancestral proteins may have had modular architectures, similar to modern repeat proteins in which a short (20-40 residue) sequence motif occurs several times within the protein chain^7^. Given that there are a number of globular proteins that are ubiquitous in all three kingdoms of life, the transition from the earliest repeat proteins to globular proteins likely preceded the last common universal ancestor (LUCA)^8^ that lived during the Archaean eon at least 2.5 BYA. The extreme length of time that has passed since then makes it likely that any repeating sequences have at best only weak similarity due to inevitable sequence drift. However, modern mammalian repeat proteins are quite well conserved despite 300 million years passing since the origin of that group suggesting that this divergence might occur quite slowly^9^. On the other hand, repeat proteins are more common in eukaryotic than bacterial genomes^10^ suggesting that many if not most date from the eukaryotic origin during the Proterozoic eon about 1-2 BYA^11^ although the repetitive nature of repeat proteins makes untangling the phylogenic relationships between these proteins difficult^12^. While as many as 25% of known, modern proteins can be classified as repeat proteins^13–14^, identification and analysis of the repeats themselves is complicated by several issues. Repeat sequences are essentially circular: there is no true start or end position^14^, insertions and deletions can occur within or between the repeats changing their apparent lengths, some repeats have a naturally variable length^7, 15^, and repeats can also occur a non-integer number of times within a protein^7^. Fortunately, a number of computational repeat detection methods exist, each focused around a different methodology and each with its own strengths and weaknesses. Unfortunately, a comparison of four distinct identification methods found little (0.2%) consensus agreement between them^16^. Despite these difficulties, a general consensus has emerged about the sequence properties of repeat proteins, namely that they are 1) self-homologous regions within a protein, 2) which have amino acid compositional biases, and 3) repeating sequence and structure patterns^17^. Awareness of these difficulties led to the development of a curated database of repeat protein structures (RepeatsDB) which uses a functional definition of a repeat protein as one that contains at least three copies of a repeating sequence with a clearly defined structure^18^. Recently, we developed a method to cluster related repeat protein sequences based on patterns of self-homology^19^ and we used this method to analyze the set of non-redundant known protein sequences (UniRef90)^20^. Upon examining our results we were surprised to find that a significant number of these related repeat protein clusters were comprised exclusively of thiamin-diphosphate (ThDP) dependent enzymes.

The ThDP enzymes are widely dispersed throughout the kingdoms of life and catalyze a wide range of reactions utilizing their nucleotide-like cofactor ThDP, the biologically active form of vitamin B_1_^21^. The reactions catalyzed by these enzymes have been the subject of 8 decades of investigation and the first crystal structures were reported nearly 30 years ago^22^ and since then several dozen genetically unique ThDP enzymes have been structurally characterized. A typical ThDP enzyme monomer is comprised of three similar domains each containing a core sheet of six β-strands surrounded by α-helices^23–24^ with the β-strands arranged in a parallel 213465 order that distinguishes it from other 3-layer alpha/beta/alpha folds such as the 321456 strand arrangement present in Rossmann domains^25–26^. One of these three domains (PP) interacts with the diphosphate moiety of the cofactor while a second domain (PYR) interacts with the cofactor’s aminopyrmidine ring through an almost universally conserved glutamate ^27–28^. All ThDP enzymes have at least one additional domain (here CFX, for cryptic function) which contain more sequence variation than the functional domains and have been previously sub-divided into several groups^29^. Some CFX domains are known to bind the nucleotide-like cofactor flavin adenine dinucleotide (FADH) or the nucleotide adenosine triphosphate (ATP)^30–31^. The ThDP family has been divided into two major groups based on the order of these domains in the protein^23^. In the first group (DH), which is comprised of the pyruvate dehydrogenases^32^ and the transketolases^33^, the domains are arranged PP, PYR, CFX from N to C. The second group (DC) which includes the acetohydroxyacid synthases^31^ and several decarboxylases^34–35^ is a circular permutation of the first group in which the domains are ordered PYR, CFX, followed by PP^23, 36^. While the ThDP family has low shared sequence identity between its members^37–38^ they all can be easily recognized by a canonical thiamin-binding motif, identified three decades ago which consists of a GDG sequence followed by about 30 variable residues and finally a conserved glutamine^39^. This canonical motif has been strictly maintained throughout the evolutionary history of this family, which is quite ancient and likely predates LUCA^8, 40–43^, allowing ready identification of ThDP enzymes from genomic sequences. Taking advantage of this recognizable motif, the TEED database of ThDP enzymes was assembled to aid bioengineering efforts and as of this writing it contains almost 120,000 protein sequences^29^.

Upon identifying the presence of protein repeats within the ThDP family we quickly noted that the repeating units themselves were homologous to the canonical ThDP binding motif. (Fig. 1). We were then able to analyze these protein repeats which underlie of all the members of the ThDP enzyme family revealing that the complex architecture of this diverse family of globular enzymes is easily built up from a set of simple, repeated oligopeptides.

**FIGURE 1:**
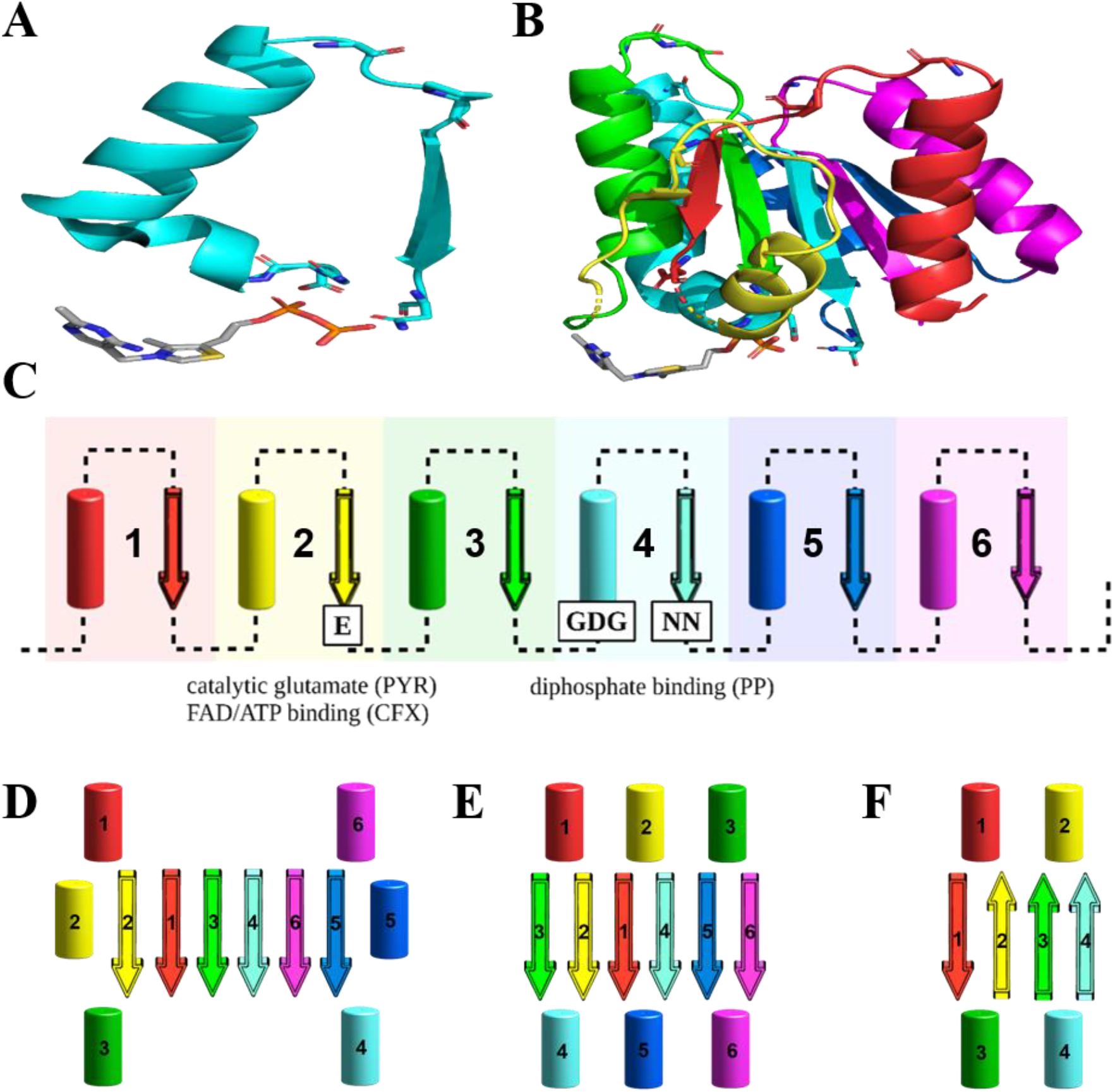
Illustration of the repeat architecture of ThDP enzymes. **A)** The canonical ThDP repeat from a pyruvate decarboxylase (PDB 2nxw) showing the helix-turn-strand structure along with the ThDP cofactor. The canonical N-terminal GDG and C-terminal “NN” residues are shown as sticks along with the conserved glycine and proline identified in the ThDP repeats. **B)** The repeats that make up the PP domain in a pyruvate dehydrogenase (PDB 1umb) colored in order from N to C red, yellow, green, cyan, blue and magenta. The ThDP cofactor is shown as sticks and colored as in panel A. Likewise the canonical ThDP binding residues (the GDG aspartate (D175) and the asparagine (N203)) are shown as sticks in the cyan color of their associated repeat. **C)** The arrangement of secondary structures in an idealized ThDP domain showing its tandem repeat structure. The repeats are colored in the same order as before and the position of the ligand interacting repeat in each of the domains is identified along with the positions of the canonical ThDP motif sequence in repeat four and the conserved catalytic glutamate in repeat three. Identically colored cartoon diagrams of the domain architectures for the **D)** PP & PYR domains, **E)** decarboxylase branch ThDP enzymes CFX domain and **F)** dehydrogenase branch ThDP enzymes CFX domain. A larger, more detailed version of this is given as SI Figure S6.

## RESULTS

### Identification of ThDP sequence repeats

After identifying the presence of a notable amount of self-homology in the ThDP enzymes present in UniRef90^19–20^, we engaged in a more thorough analysis of those proteins present in the PDB^44^. Sequence redundant proteins were removed and the remaining ones were organized phylogenetically (Fig. 2). Reverse calculation using DOTTER^45^ identified the canonical ThDP binding motif as one of the regions of self-homology and a (G/A){X(1,2)}(G/A) motif (where G/A indicated either a glycine or alanine residue and X(1,2) indicates one or two variable residues) at the beginning of a helix-turn-strand structural motif was quickly identified (Fig. 3). While the choice of the beginning of a repeat is largely arbitrary^14, 46^, we identified this motif as the start of the repeat for historical reasons^39^. Using these basic features, we were able to identify 1003 repeats from a set of 58 non-redundant ThDP proteins (Figure 3, SI Fig. S1, SI Table S1, S2, & S3) present in the Protein Data Bank in October 2019^47^ by manually examination (Materials and Methods). At least three repeat sequences were identified in each protein (*i.e.* one of the basic identifiers of protein repeats^16^). All the repeats had an average (mean) length of 27 residues (median = 26), although the repeats from the functional PP and PYR domains tended to be slightly longer than those from the CFX domain (mean = 28.7, 27.9, & 24.0 respectively) (Fig. 4, SI Table S1). 663 (66%) of all the identified repeats were 22-32 residues long with a mean length of 26.8 residues, suggesting a relatively standard repeat length (Figure 4) and these comprised the “best repeats” subset (SI Tables S1 & S2). While the strongest sequence conservation currently detectable was the (G/A){X(1,2)}(G/A) motif, some other sequence tendencies were also identified (SI Table S4). The helix itself is somewhat enriched in alanine residues, although this is typical of helical regions in the PDB in general. There is often a glycine or proline in the first or second position after the helix (37.4% of the repeats between 22-32 residues long and 36.6% of all the identified repeats) and a proline or a glycine in the position before the start of the strand (26.5% of the repeats between 22-32 residues long and 19.5% of the full set, see SI Tables S1-S4). Both of these proline/glycine residues were present in 7.8% of the repeats that were 22-32 residues long (5.4% among all the identified repeats). The strand itself is rich in valine, leucine, and isoleucine residues and the strand often ends on a hydrogen bond donating residue. These basic components were also noticed when the canonical ThDP binding motif was first identified^39^ and at least one later analysis of the canonical ThDP motif also noted a conserved valine residue and other sequence features preceding the canonical GDG motif, exactly as would be expected to occur due to the preference for hydrophobic residues in the strand at the C-terminus of the preceding repeat^48^. Several conserved functional positions were also associated with specific repeats (Figure 1C). The canonical ThDP binding sequence^39^ is always the fourth repeat in the PP domain. The conserved catalytic glutamate^27^ is present at the end of repeat two in the PYR domain. Repeat two in the CFX domain is associated with interaction with an additional nucleotide cofactor in those enzymes which also bind FADH or ATP^30–31^. With these repeats identified, we expanded our analysis to the entirety of the TEED database^29^. Sequences in the TEED database that had significant (>30%) pairwise identity to each of the 58 PDB structure sequences were used to generate matrices of consensus sequences. An increasing bias towards the amino acids alanine, glycine, and proline and a bias away from lysine and cysteine, which became more evident as the threshold for conservation (Simpson)^49–50^ increased and the sequence identity threshold decreased, was identified within the repeat regions (SI Figure S2). This bias became more and more pronounced as the overall conservation of the proteins decreased and the observed diversity at that specific position decreased. We also attempted to identify more specific sequence requirements within the repeats but this was stymied by the bias of the TEED database towards transketolases (TK), acetohydroxyacid synthases (AHAS) and pyruvate dehydrogenases (PDH) (SI Table S5). This largely restricted our analyses to these functional classes; we provide one analyzed example each from the DH class (the transketolases) (SI Figure S3) and one from the DC class (the acetohydroxyacid synthases, SI Figure S4). While there was no ubiquitous sequence pattern (other than the initial (G/A){X(1,2)}(G/A) motif), there was notable self-similarity between the copy of the repeat found in the PP and that found in the PYR domain for every repeat position in the representatives from both the DH and DC functional classes. The CFX domain repeats were much more diverged from the functional domain repeats and the domain topology is also different (see below) making it unclear if the repeats are present in the same order in the functional and CFX domains. While self-homologous residues were easier to detect near the edges of the repeats it was possible to identify a number of positions conserved in both classes of ThDP enzyme (*e.g.* “AATF” or “DKPTϕϕ” toward the C-terminus of the third and sixth DH repeats (SI Figure S3) or “PGPVLLDϕ” near the C-terminus of the sixth DC repeat (SI Figure S4) where ϕ is a hydrophobic residue). In several cases it was possible to detect the same conserved residue patterns from both the N and C termini allowing an additional estimate of the overall repeat length. Additionally, in both the DH and DC classes a conserved alanine was located at position 10 in repeat 3 and a conserved pair of residues was also identified at positions 5-6 in repeat 5 in both classes. These conserved residues were linked to conserved contacts in the ThDP enzymes (see below).

**FIGURE 2:**
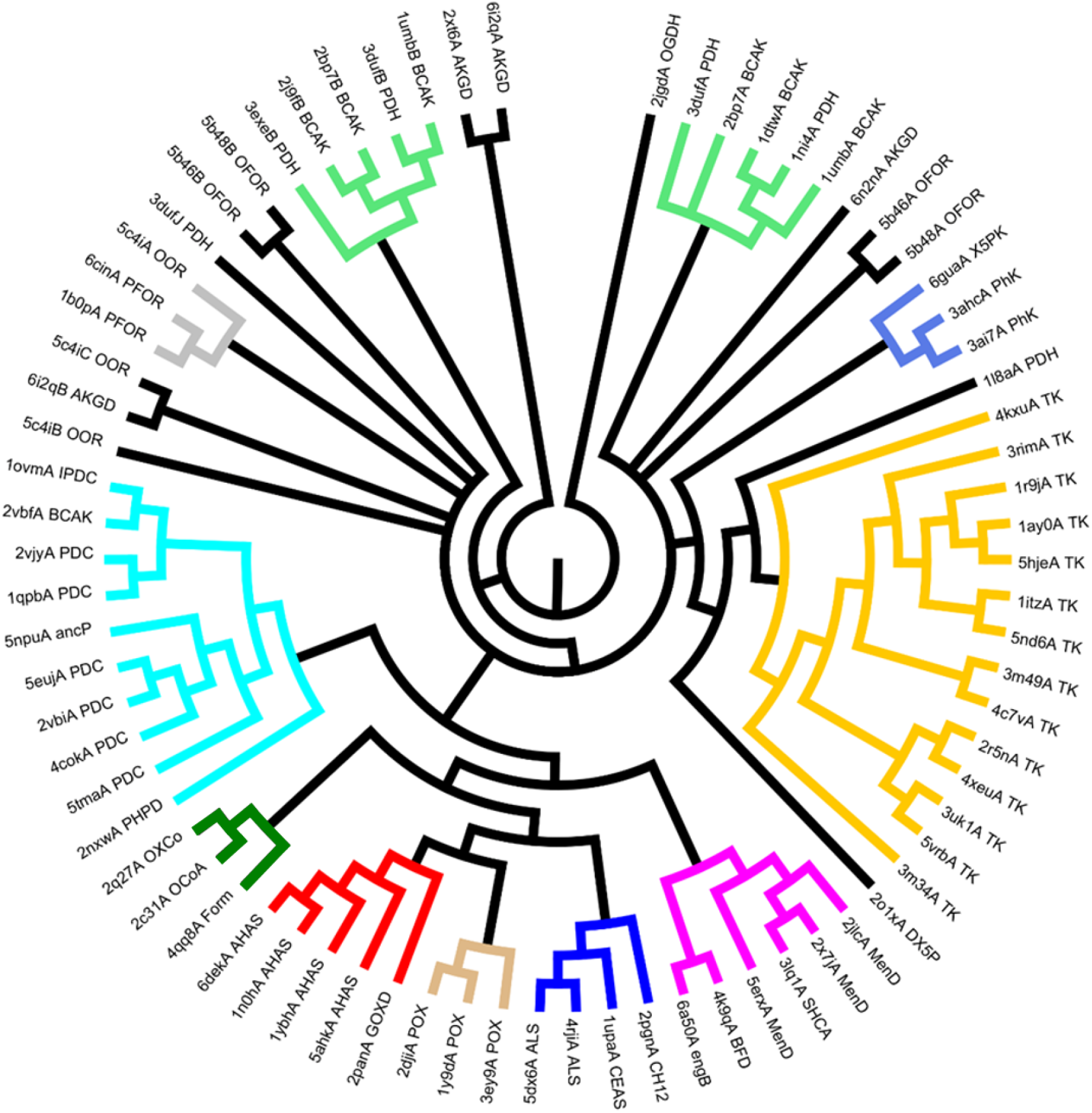
Phylogenic tree of the ThDP family enzymes. A phylogenic tree of the ThDP enzymes of known structure. Groups of more than 3 members were separated into different groupings indicated by color as follows: AHAS (red), ALS (dark blue), BFD (magenta), OxCDC (dark green), PDC (cyan), PDH (green), PhK (light blue), PFOR (grey), POX (beige), & TK (mustard). The PDH group of proteins consists of two separately expressed chains which do not co-cluster in this tree but are both indicated by the same color here.

**FIGURE 3:**
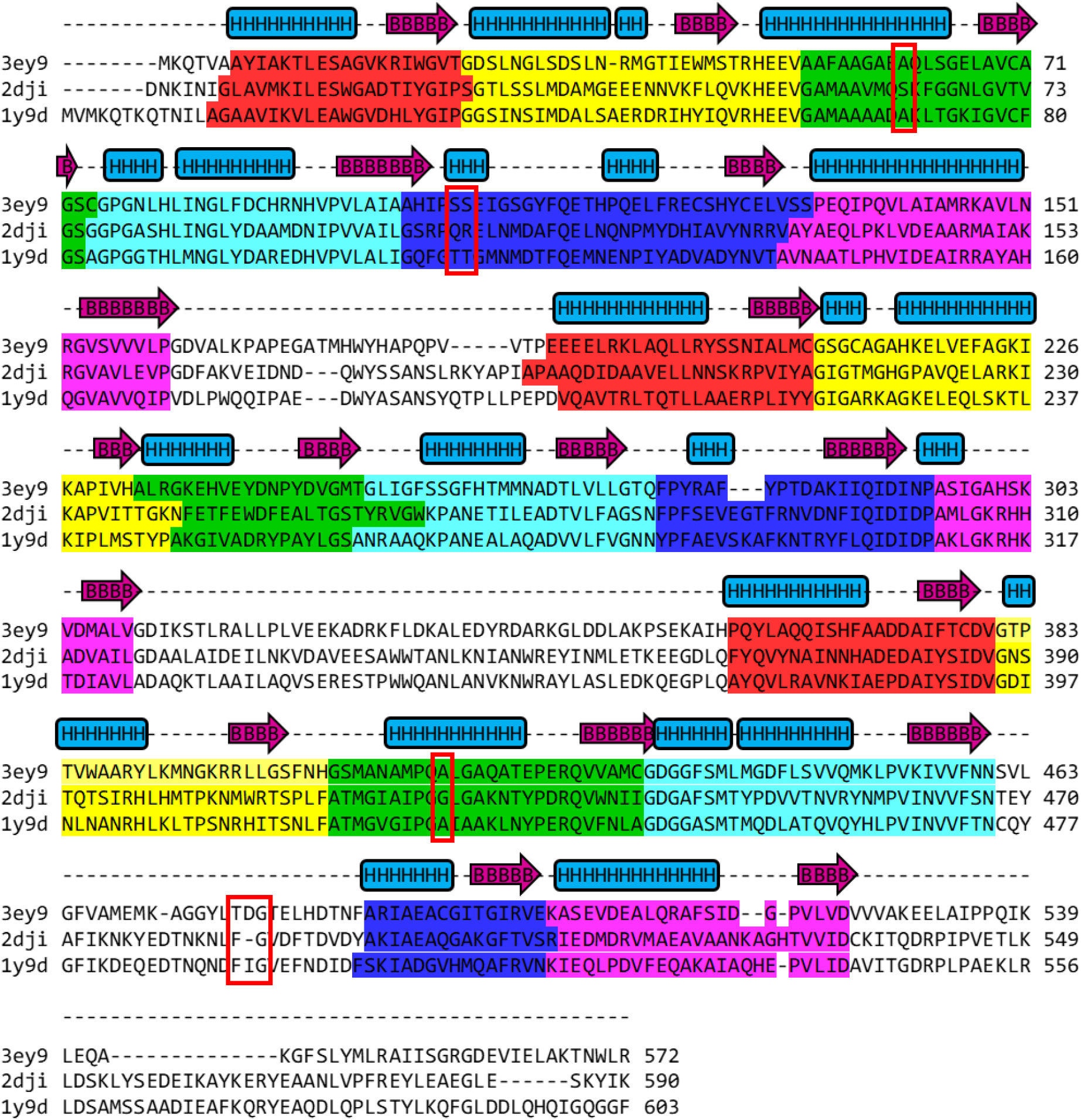
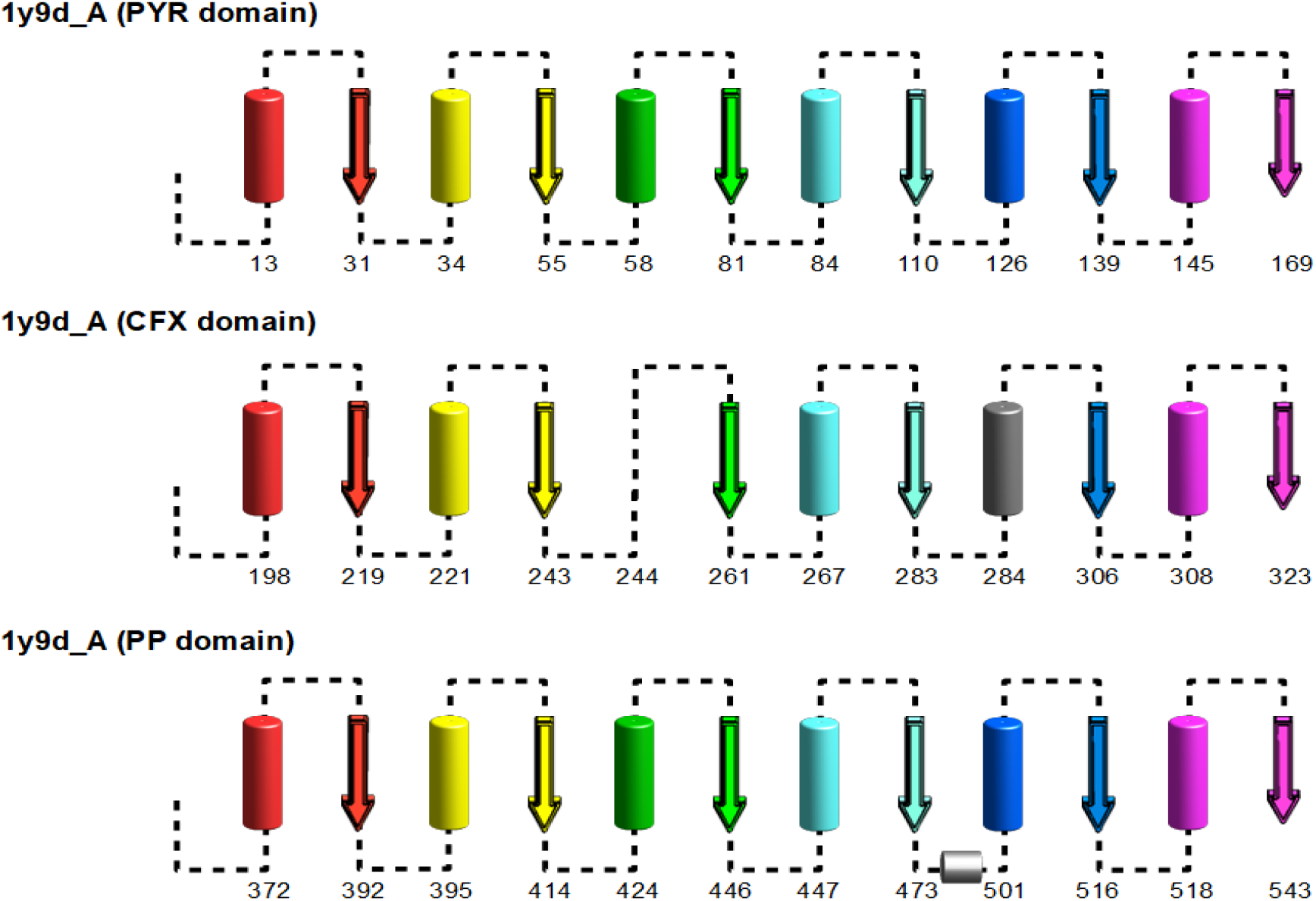
Illustration of the protein repeats present in the pyruvate oxidases cluster of ThDP enzymes. **A)** The repeats are highlighted in the order red, yellow, green, cyan, blue, and magenta for each of the three domains respectively. The three state secondary structure (α-helix (light blue), β-strand (plum), coil (dash)) for the repeat regions in PDB 3ey9 is shown above the multiple sequence alignment for reference. The conserved alanine in position 10 of repeat 3 and the conserved sequence in position 5–6 in repeat 5 is shown by a red box. In the DC enzymes, the conserved region in repeat 5 of the PP domain appears within the insertion between repeat 4 and 5. **B)** Cartoon illustration of the protein repeats present in the POX group of enzymes using the representative PDB 1y9d. The repeats are numbered and colored in the order red, yellow, green, cyan, blue, and magenta from N to C as in figure 1. Helical regions are indicated by a cylindrical tube and strand regions by an arrow. The number of the first helical residue in the helical region and the last beta residue in the strand region are also indicated. Insertions that contain secondary structure are indicated in grey while missing repeats are indicated by missing cartoon images. The three domains are shown in the order they occur in the protein. Secondary structure indication is derived from PyMol. Note that the helical region in the fifth repeat of the CFX domain is not resolved in the structure so it is colored grey here. Equivalent figures for a representative example from each of the other enzyme groups is given as SI Figure S5.

**FIGURE 4:**
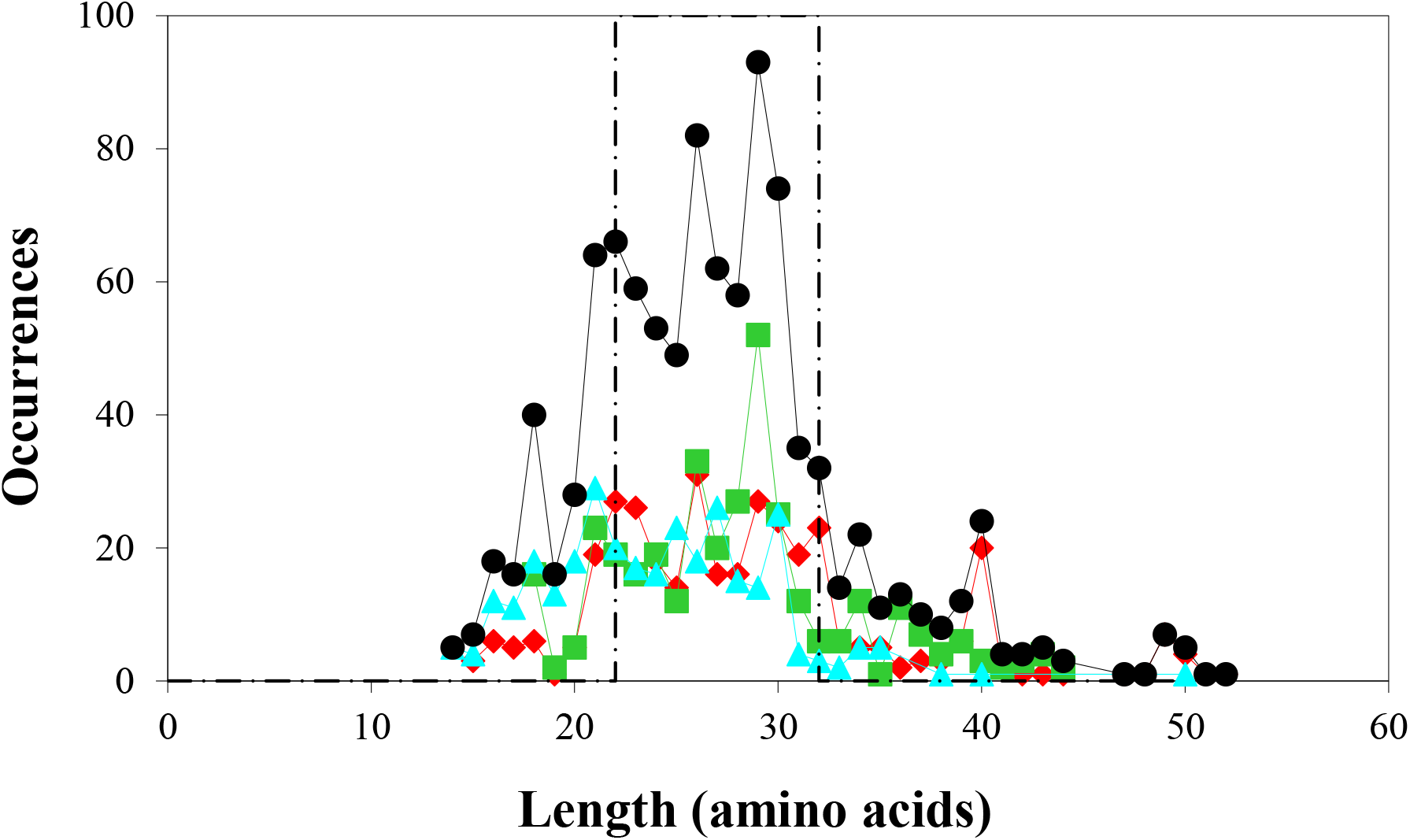
Size distribution of the ThDP repeats identified from the filtered PDB set. The number of repeats of each sequence length is shown for all the domains (black circles, N = 37), the diphosphate binding PP domain (red diamonds, N = 36), aminopyrimidine interacting PYR (green squares, N = 27), and extra third domain (cyan triangles, N = 25). The typical (66%) repeat length of 22–32 residues is indicated by a set of vertical, dashed black lines.

Our DOTTER-based self-homology method^19^ was not the only analysis method which could detect repeats in ThDP enzymes as a weak but noticeable positive signal was detected by other computational repeat identification methods (Supplemental Data S1). While not every repeat identification method could find the ThDP repeats (which is not unusual for repeat identification^16^) the fact that these repeats could be identified by multiple methodologies strengthens the assertion that these are protein repeats. That the divergence of this family occurred a remarkably long time ago suggests that ThDP-like repeats may exist in other protein families, either through gene transfer or due to their relative simplicity by having arisen multiple times during the existence of life on Earth. A search of the PDB using the ThDP repeat identified 881 non-ThDP enzyme structures with at least 3 copies of this repeat. Furthermore, these proteins tend to be enzymes that bind nucleotide-like molecules (Supplemental Data S2), as had been previously noted for a “transketolase motif” which also fits comfortably within our definition of the ThDP repeats^33^. Given the likely ancient origin of the ThDP enzymes^8, 41^ and the significant amount of time that has passed since divergence of the family, it is not surprising that these repeats are generally weakly conserved, although variation similar to what is seen in the ThDP enzymes is also not unusual for protein repeats^15^.

### Structural features of the repeats in the ThDP enzymes

The tandem repeat pattern can be easily identified in the structures of the individual domains in the ThDP enzymes and the arrangement of the repeats is highly conserved within the branches of ThDP family with a few notable exceptions (SI Table S6, SI Figure S5). The DH enzyme have a conserved insertion containing a secondary structure element after the first and fourth repeats in the PP domain (the insertion after the fourth repeat is missing in the phosphoketolase (PhK) enzymes) while the DC enzymes also have a conserved insertion after the fourth repeat in the PP domain. This insertion is also present in the PFOR enzymes, but is not associated with a secondary structure element. The spatial arrangement of the repeats is also highly conserved. The functional PP and PYR domains have a 213465 parallel strand arrangement with the 213 and 465 sets of helices partitioned above and below their respective strand sets in the DH, DC and PFOR enzymes (Figure 1 C, SI Figure S6A). The CFX domains of the DC class also contain 6 repeats but in a Rossmann-like 321456 parallel arrangement with the 321 and 456 sets of helices arranged exclusively above or below the plane of the strands (Figure 1D, SI Figure S6B). This domain is not a true Rossmann domain as it does not contain the conserved ligand binding geometry of the true Rossmann domains^26^ (SI Table S7). The CFX domains of the DH class are shorter, typically only containing 4 repeats with the first strand (Figure 1E, SI Table S6, SI Figure S6C). An additional two types of CFX domain with antiparallel strand arrangements are found in the PFOR enzymes (SI Figure S6D & S6E).

The repeat residues that are conserved in the both the DH and DC enzyme classes reflect the architecture of the ThDP enzymes. While it has long been known that the residues associated with repeat 4 in the PP domain bind the ThDP phosphate, the N-terminus of the helix in repeat 2 and the conserved sequences in repeat 5 (or in the equivalent insert region in the DC enzymes) also interact with the cofactor phosphate through the chelating magnesium. The conserved alanine at position 10 in repeat 3 is associated with the intra chain contact between the two functional domains although this alanine faces away from the contact surface (SI Figure S7). Likewise, the conserved residues in repeat 5 form an inter chain contact along with repeat 4 (SI Figure S8) and while there are no obvious hydrogen bonding interactions between these two helices, it should be noted that repeat 3 is in a central position in the functional domains and as this is an intra chain interaction and it is suspected that hydrophobic burial drives this interaction. Much like repeat 3, these residues are somewhat poorly oriented to directly participate in the interaction, they are difficult to identify from the sequences (SI Figure S1). While these residues can be readily identified from the structure, and there is a tendency for the sequence to be a hydrogen bonding residue followed by small one. Additionally, in the structure of a BFD enzyme (PDBID 6A50), there is a magnesium binding site formed by this inter chain contact through their backbone carbonyls in repeat 5 of the PYR domain.

## DISCUSSION

The first structure of a ThDP enzyme was solved in the early 1990s^51^, and dozens of additional structures have been reported since then, revealing a number of nearly perfectly conserved features of this family, including both the canonical ThDP binding motif^39^ and the conserved catalytic glutamate^27^. Yet numerous questions about this family of enzymes have remained unanswered, notably that the catalytic glutamate is not universally conserved^28^, there is significant sequence diversity among the members of the family^37–38^, and that even the domain arrangements within the ThDP family can be variable^36^, raising the question as to what is being fundamentally maintained by evolution within this family of enzymes, other than the canonical ThDP motif itself. But now the identification of the protein repeat architecture in the ThDP enzymes reveals an elegant organizing principle for this entire family of enzymes (Fig. 1).

Even though the ThDP enzymes diverged from their common ancestor in the distant past (even before LUCA), and the overall similarity between the repeats is quite weak, there are still substantial details to identify these repeats. Repeat identification is difficult, even under optimal conditions^13, 52^, and there is little consensus agreement between methods^16^ however a consensus has emerged for identification of repeats^17^, namely that repeats have 1) evident self-homology, 2) compositional biases, and 3) a repeating pattern that is identifiable in the sequence and structure of the repeats. All of these are present in the ThDP enzymes. First, self-homology is quite evident from self-alignment of the ThDP sequences^19, 45^. While there are not long stretches of regular patterns evident in the self-alignment plots, this is also not uncommon for many recognized repeat proteins and the previous analysis found that the amount of self-homology signal detected was well above the threshold amount expected as determined by comparison to other known repeat proteins^18–19^. Second, the ThDP repeats do have biased amino acid distributions (SI Figure S2). They show a preference for alanine, glycine, and proline and a bias against cysteine and lysine. While these preferences cannot be fully separated from the secondary structure propensities of these repeats (SI Table S8), the fact that these preferences were more evident as global sequence identity decreased suggests that they are an integral part of the architecture of these proteins as these preferences change more slowly than sequence does overall (SI Fig. S2). The fact that alanine occurs more often in helical sequences does not preclude it from being recognized as part of an identifying pattern in the ThDP repeats as there are conserved alanine residues identified in fully helical repeats^53^. Third, repeating patterns, in both sequence and structure, are readily identifiable in the ThDP enzymes. While the most obvious of these is the (G/A){X(1,2)}(G/A) motif, there are definite preferences at other positions within the repeats as well. This includes the enrichment of isoleucine, leucine, and valine residues at the end (strand portion) of the repeat (as previously recognized^48^) and the tendency of the turn portion of the repeat to be capped with proline and glycine residues (SI Table S4). It cannot be completely ruled out that these preferences are due to secondary structure constraints, however. And while no other universally conserved position was detected, when the individual repeats are compared between the two functional domains, patterns emerge both within functional classes (SI Figures S3 & S4) and throughout the ThDP family of repeated sequence patterns within individual proteins. By measuring the distances from both ends of the repeats (N and C) we were able to account for differences in repeat lengths and identify conserved features such as the conserved alanine at position 10 in repeat 3 that is associated with the intra-chain contact between the PP and PYR domains (SI Figure S7) and the conserved residues at position 5 & 6 in repeat 5 that is associated with the ThDP phosphate/magnesium and an inter-chain interface (SI Figure S8). To the best of our knowledge, none of these conserved sequences have been previously noted, demonstrating the utility of recognizing the repeat architecture in the ThDP enzymes, given that these residues have orientations which are not obviously involved in the interactions but these sequences are strongly conserved in both the DH and DC branches of the family. While the conserved section of repeat 5 is easily identified in the DH enzymes, it occurs within what was identified as the insertion region between repeat 4 & 5 in the DC enzymes highlights some of the limitations of our ability to identify the limits of the repeats; even now it is unclear if the insertion after repeat 4 in the DC PP domain should be included as part of repeat 5 or not as there is no clear initiation sequence in these insertions near the conserved sequence (SI Figure S1) even though they appear to interact with the cofactor diphosphate. The conserved residues in repeat 5 also appear to be possibly involved in a cryptic magnesium binding site^54^ (SI Figure 9) in the PYR domain, further suggesting the utility of identifying the repeat architecture of these enzymes, and perhaps the original, primitive function of these repeats. So, while the repeats themselves are quite diverged and sequence similarity is rather low, the identifying marks of self-homology, sequence bias, and repeating sequence and structure patterns are still notable suggesting that the modern ThDP enzymes still maintain a repeat architecture.

Identifying this repeat architecture also helps to simply structural analysis of these enzymes, which has been previously quite complex^23^. This complexity is partially due to breaks in the secondary structure, either due to natural structural drift or crystallographic assignments (Supplemental Figure S5), but recognition of the overarching repeat patterns alleviates this difficulty. The helix-turn-strand secondary structure pattern of the ThDP repeats (Supplemental Figure S6) also highlights many of the functions ThDP enzymes. Each of the three domains (PP, PYR, & CFX) has a unique function that is dependent on a specific repeat. The canonical repeat is always the fourth repeat in the PP domain (Fig. 1C) but it also cooperates with the conserved residues in repeat 5 and repeat 2 (at least in the DC branch and the transketolases). This repeat also typically overlaps with the repeat preceding it (*e.g*. the GDG aspartate also functions as the terminal carbonyl residue of the previous repeat). The conserved, catalytic glutamate that mediates the protein interaction with the aminopyrimidine nitrogen^35^ occurs just before the beginning of the third repeat. It bears noting that this repeat structure occurs even in the glyoxylate carboligases which are unique among the ThDP family in lacking the conserved glutamate that interacts with the aminopyrmidine moiety of the cofactor^28^. Residues from these two functional repeats have also recently been shown to mediate catalysis and communicate through a hydrogen bond between two the paired monomers in transketolase which emphasizes their functional significance^55^.

Both the PP and PYR domain have a universally conserved 213465 strand arrangement (Fig. 1, SI Fig. S6, SI Table S6) clearly indicating a common ancestral origin for these catalytic domains^56–57^. The CFX domains have a more diverse set of domain architectures with the members of the decarboxylase class having a Rossman-like parallel 321456 strand arrangement and the dehydrogenases an antiparallel 1(−2)(−3)(−4) arrangement (Fig. 1, SI Fig. S6)^23, 36, 38^. And while the functional ThDP domains have a similar strand arrangement to the Rossman-like flavodoxins, the (G/A){X(1,2)}(G/A) motif that starts the ThDP repeats is similar but not identical to the motifs that start either the Rossmann (GxGxxG) or the Walker A (GxxGxG) proteins suggesting convergent evolution rather than common descent^58–59^. Additionally, some ThDP enzymes are known to bind other nucleotide-like ligands such as FAD^30^ (which does not always have an obvious catalytic role^60^) or bind ADP in a functional role (which was suggested to be derived from the FAD binding site^31^). The diphosphate of these ligands is bound by the second repeat in the CFX domain. FAD and ADP binding domains confined to the decarboxylase branch of the family with its Rossmann-like CFX domain, however they do not display the conserved geometry of the true Rossmann domains further indicating convergent evolution rather than common descent (SI Table S7)^26^. More curious, however, are the two kinds of CFX domains which are exclusively present in the pyruvate-ferredoxin synthases/pyruvate synthases (PFOR) which have a PYR-CFX-PP domain arrangement similar to the decarboxylase branch, but domain architectures found in the dehydrogenase branch including a domain which differs from the dehydrogenase CFX domain by the addition/deletion of a single terminal repeat (SI Fig. S6, SI Table S6). These observations suggest that the PFORs may be the closest extant group to the root of the ThDP family, although we cannot rule out a polypheletic origin^61^.

It bears noting here that the ThDP repeats are not necessary to bind thiamin diphosphate^62–63^ and some proteins involved in ThDP biosynthesis are apparent true Rossmann proteins^56^ indicating that the use of the ThDP repeat architecture was likely a stochastic rediscovery of a suitable phosphate binding structure ^42–43^. That both the pyruvate dehydrogenases and the transketolases are widely dispersed among all three kingdoms of life, and both of these ThDP enzymes, as well as the possibly ancestral pyruvate synthases (PFOR) catalyze reactions believed to have been essential to life during the Archean eon^8, 40–41^ evidence the extreme age of this family. It is even more noteworthy in a family of enzymes that generally has low shared sequence homology^38^ in which even catalytically essential residues can be mutated without major losses in enzymatic activity^28, 64^, that we have been able to identify the universally conserved residues in repeats 3 and 5 by recognizing this architecture. That the ThDP repeats were conserved enough to be recognizable in addition to the observation of the conserved 213465 β-strand arrangement in the catalytic PP & PYR domains suggests that the repeat architecture is fundamental to the function and structure of these enzymes. All of these observations are consistent with the hypothesis that early proteins, such as the ancestor of the ThDP enzyme family, were repeat proteins derived from an assembly of short peptides^2, 4–6^.

## MATERIALS AND METHODS

Briefly, as described elsewhere^19^, protein chains in UniRef90^20^ were analyzed using DOTTER^45^. Protein chains with a threshold repeat signal from DOTTER were collated and compared pairwise using a Jaccard metric and then clustered using MCL^65^. Clusters containing between 5 and 200 members (N=8569) were analyzed manually and 55 (0.64 %) of these were comprised exclusively of ThDP enzymes.

ThDP enzyme sequences were collected from the PDB^44^ and redundancy was removed by CD-HIT^66^ at 90% sequence identity. A multiple sequence alignment was generated from this set (N_chains_ = 58) using MUSCLE^67^ and a phylogenic tree was generated using MrBayes^68^ (Fig. 2). Branches containing at least three leaves (N_branches_ = 10) were collected and then individual multiple sequence alignments were generated again using MUSCLE. The branches were named based on traditional functional groupings of ThDP enzymes. The members of these branches were analyzed by DOTTER and the DOTTER plots were decomposed to identify sections of sequence that generated self-similarity signal. The self-similarity score for any residue was defined as the maximum score in the in the row or column associated with that position. Plots of these DOTTER scores with sequence position and the multiple sequence alignments of the branches were used to identify the repeating sequence by looking for regions of conserved sequence identity. Putative repeats were then confirmed structurally using the protein structures as defined by PyMol^69^. This lead to the detection of the [G/A]{X(1,2)}[G/A] motif near the N terminus of a helix to start the repeat (a looser criterion than the “GDG” of the ThDP binding motif) and the set of non-redundant ThDP proteins was then manually analyzed. Because the repeats largely occurred in tandem, they were generally terminated by the appearance of the either the next [G/A]{X(1,2)}[G/A] motif or the start of another helix. When this was not possible, the borders were determined using the information available from the multiple sequence alignment and the protein structures. This identified a set of 1003 repeats which were then analyzed statistically to generate a “typical” repeat (Fig. S1, SI Table S1). The central 65% (from 20-85% of the distribution, Figure 4) was used to identify the repeat length. Sequence and structure propensities were calculated from the set of all the identified repeats. Domain architecture was identified manually using PyMol^69^. The ligand binding geometry for identification of true Rossmann domains was analyzed as described elsewhere^26^.

The non-redundant set of ThDP enzymes of known structure was compared to the TEED database^29^ by pairwise global sequence alignment using the BioConductor suite in R^70^. Sequences from the database were sorted based on their identity to each of the ThDP enzymes at cutoffs of 80, 70, 62, 50, 40, & 30 % identity. Within each set, for each repeat, the matching sequence was identified and recorded to produce a 21 member (all 20 amino acids and gap) matrix for each position in the repeat. At each position the degree of conservation was quantitated using the Simpson metric^49–50^. The amino acid distribution for each identity and Simpson threshold was also calculated. The distribution of amino acids for each position in the TEED set was calculated and used to formulate consensus repeats. The repeats from the functional PP and PYR domains were compared, with each position analyzed relative to its distance from either terminus to account for the variation in repeat lengths. Due to the bias in the TEED database towards transketolases, the repeats were generally analyzed within their functional subgroups.

The non-redundant PDB chains were also subject to other methods of *de novo* repeat identification with RADAR^71^, T-REKS^72^, HHREP^52^, XSTREAM^73^, and TRUST^74^ which gave weak results that were nevertheless consistent with our method (Supplemental Data S1). The ThDP repeat was used to search the set of all known protein structures using the sequence and secondary structure as defined by the PDB in Nov. 2019. The repeat was defined as [G/A][X][G/A] where X could be any one or two amino acids and [G/A] was either a glycine or alanine residue followed by a helical segment and then a beta strand segment where the last residue of the strand or the residue after the end of the strand was a glutamate, glutamine, aspartate, or asparagine. The helical region was allowed to include secondary structure “H”,”G” or “I” while the strand was “B” or “E”. In order to match the sequence preferences, helical regions were also required to have at least 9% alanine content and strand regions must have contained at least a 20% total of valine, leucine or isoleucine residues. The length of the repeat must also have been 14-36 residues long and at least three had to occur within one protein chain for the chain to be identified as repeat containing. This identified 953 PDB structures (72 ThDP structures and 881 non-ThDP structures) which represented 0.56% of the PDB at the time (Supplemental Data S2).

## ACKNOWLEDGEMENTS

The work was supported by the National Science Centre, Poland [grant agreement 2014/15/D/NZ1/00968] and the European Union’s Horizon 2020 research and innovation programme [Marie Skłodowska-Curie grant agreement No 655075]. The authors wish to thank Profs. Ben Luisi & Vikas Nanda for critical readings of the manuscript. Computer code for this work is available online at https://gorna.uw.edu.pl/en/research/software. The authors declare that they have no competing financial interests regarding this work.

## Notes

### Competing Interest Statement

The authors have declared no competing interest.

