## Supplemental Material for "Thiamin-Diphosphate Enzymes Are an Ancient Family of Repeat Proteins"

**Supplemental Material for**
**"Thiamin-Diphosphate Enzymes Are an Ancient Family of Repeat Proteins"**

Matthew Merski [1,*] & Maria Górna[1,*]

[1]Structural Biology Group, Biological and Chemical Research Centre, Department of Chemistry, University of Warsaw, Warsaw, Poland

**Supplemental Table S1:** Statistical analysis of the identified ThDP repeats.

| | **All repeats** | **Best repeats (22-32 residue length)** |
|---|---|---|
| **Number of repeats =** | 1003 | 663 |
| **Mean (median) repeat length =** | 27.0 (26) | 26.8 (27) |
| **Number of PP repeats =** | 349 (34.8 %) | 241 (36.3 %) |
| **Mean (median) PP domain length =** | 28.7 (28) | 26.9 (27) |
| **Max. PP domain length =** | 52 | 32 |
| **Min. PP domain length =** | 15 | 22 |
| **Number of PYR repeats =** | 348 (34.7 %) | 241 (36.3 %) |
| **Mean (median) PYR domain length =** | 27.9 (28) | 27.0 (28) |
| **Max. PYR domain length =** | 44 | 32 |
| **Min. PYR domain length =** | 18 | 22 |
| **Number of CFX repeats =** | 306 (30.5 %) | 181 (27.3 %) |
| **Mean (median) CFX domain length =** | 24.0 (24) | 26.3 (26) |
| **Max. CFX domain length =** | 50 | 32 |
| **Min. CFX domain length =** | 14 | 22 |

**Supplemental Table S2:** A list of the complete set of 663 “best” ThDP repeats found in the filtered PDB set listing the PDB ID, the sequence of the repeat, its length, and which of the three domains (PP, PYR, or CFX) it is located in. There are 241 (36.3%) PP domain repeats, 241 (36.3%) PYR domain repeats, and 181 (27.3%) CFX domain repeats in this set.

| | PDB ID | Class | Sequence | Length | Domain ID |
|---|---|---|---|---|---|
| 1 | 1b0p | PFO | AFFAALPASAKVITVLDRTKEP | 22 | CFX |
| 2 | 1b0p | PFO | GEKPIQSTYLVNRADYVACHNP | 22 | CFX |
| 3 | 1ni4 | PDH | GGCAKGKGGSMHMYAKNFYGGN | 22 | PP |
| 4 | 1ovm | PDC | QENFWRTLQTFIRPGDIILADQ | 22 | PP |
| 5 | 1qpb | PDC | QEWMWNQLGNFLQEGDVVIAET | 22 | PP |
| 6 | 1upa | ALS | AGVAADVLARITGRPQACWATL | 22 | PYR |
| 7 | 1upa | ALS | GPGMTNLSTGIATSVLDRSPVI | 22 | PYR |
| 8 | 1y9d | POX | AYQVLRAVNKIAEPDAIYSIDV | 22 | PP |
| 9 | 2bp7 | PDH | GIVGTAVGMGAYGLRPVVEIQF | 22 | PYR |
| 10 | 2bp7 | PDH | GTTVYVAQVAAEESGVDAEVID | 22 | CFX |
| 11 | 2c31 | OxC | AEDAIARAADLIKNAKRPVIML | 22 | CFX |
| 12 | 2c31 | OxC | AGYAASIAGYIEGKPGVCLTVS | 22 | PYR |
| 13 | 2dji | POX | FYQVYNAINNHADEDAIYSIDV | 22 | PP |
| 14 | 2j9f | PDH | GEEGTHVGSAAALDNTDLVFGQ | 22 | PP |
| 15 | 2jlc | BFD | SAEEGKKVALWAQTLGWPLIGD | 22 | CFX |
| 16 | 2pgn | ALS | GDIAGALQRAIDSGKPALIEIP | 22 | PP |
| 17 | 2q27 | OxC | AGYAAAASGFLTQKPGICLTVS | 22 | PYR |
| 18 | 2vbf | PDC | DSSTGAFTHHLDENKMISLNID | 22 | CFX |
| 19 | 2vbf | PDC | QDRLWQAVESLTQSNETIVAEQ | 22 | PP |
| 20 | 2vbf | PDC | YSKLPETFGATEDRVVSKIVRT | 22 | PP |
| 21 | 2vbi | PDC | DYSTVGWSAWPKGPNVILAEPD | 22 | CFX |
| 22 | 2vbi | PDC | NDEIVRHINALLTSNTTLVAET | 22 | PP |
| 23 | 2vbi | PDC | YAGLMEVFNAGEGHGLGLKATT | 22 | PP |
| 24 | 2vjy | PDC | GKGSIDEKHPRFGGVYVGTLSS | 22 | CFX |
| 25 | 2vjy | PDC | QEWVWTQVGEFLREGDVVITET | 22 | PP |
| 26 | 2x7j | BFD | EGNLYRILQHLVPENSSLFVGN | 22 | PP |
| 27 | 2x7j | BFD | SDADKENIIALSKALQYPILAD | 22 | CFX |
| 28 | 3ahc | PhK | ATDANMLLAISEKCFKSTNKIN | 22 | PYR |
| 29 | 3ai7 | PhK | ATDANMLLAIAEKCYKSTNKIN | 22 | PYR |
| 30 | 3exe | PDH | GGCAKGKGGSMHMYAKNFYGGN | 22 | PP |
| 31 | 3ey9 | POX | AAFAAGAEAQLSGELAVCAGSC | 22 | PYR |
| 32 | 3ey9 | POX | EEEELRKLAQLLRYSSNIALMC | 22 | CFX |
| 33 | 3ey9 | POX | KASEVDEALQRAFSIDGPVLVD | 22 | PP |
| 34 | 3ey9 | POX | PQYLAQQISHFAADDAIFTCDV | 22 | PP |
| 35 | 3lq1 | BFD | KKELEQPMVDLAKKLGWPILAD | 22 | CFX |
| 36 | 3lq1 | BFD | VSKPLKNWLEQLSDIRFYVVDP | 22 | CFX |
| 37 | 3m34 | TK | AMAAINNAFARYGIFLPFSATF | 22 | PYR |
| 38 | 3m49 | TK | AMGAAMNGIALHGGLKTYGGTF | 22 | PYR |
| 39 | 3rim | TK | AMGAILSGIVLHGPTRAYGGTF | 22 | PYR |
| 40 | 3rim | TK | GVPVLDGTDAEGVARGGYVLSD | 22 | CFX |
| 41 | 3uk1 | TK | AFNARTDAQLANVEKGGYVLRD | 22 | CFX |
| 42 | 3uk1 | TK | GMSAAINGLVLHGGYKPFGGTF | 22 | PYR |
| 43 | 4c7v | TK | GMACAMNGIMLHGGTRIFGSTF | 22 | PYR |
| 44 | 4cok | PDC | AAGAQAQAVALADALGCAVTTM | 22 | CFX |
| 45 | 4cok | PDC | NAEMARQIGALLTPRTTLTAET | 22 | PP |
| 46 | 4cok | PDC | YAGLMEVFNAGEGNGLGLRART | 22 | PP |
| 47 | 4kxu | TK | GGIGEAVSSAVVGEPGITVTHL | 22 | CFX |
| 48 | 4rji | ALS | AAFMAQAVGRLTGKPGVVLVTS | 22 | PYR |
| 49 | 4rji | ALS | GPGASNLATGLLTANTEGDPVV | 22 | PYR |

| | | | | | |
|---|---|---|---|---|---|
| 50 | 4rji | ALS | PLEIVKELRNAVDDHVTVTCDI | 22 | PP |
| 51 | 5ahk | AHA | PFGLFTQLNKLTERVALDYILD | 22 | PP |
| 52 | 5c4i | PFO | HAQLSVVYGASAAGARVFTGSS | 22 | PYR |
| 53 | 5c4i | PFO | SIDTILEFLGDTGNLAQIVTVD | 22 | CFX |
| 54 | 5erx | BFD | AGYLAIGLAIGAGAPVCVAMTS | 22 | PYR |
| 55 | 5erx | BFD | GLHVAAAVSHALRPGDQLVLGA | 22 | PP |
| 56 | 5euj | PDC | GAQELVENADAILCLAPVFNDY | 22 | CFX |
| 57 | 5euj | PDC | GFSAEGYARARGAAAAIVTFSV | 22 | PYR |
| 58 | 5euj | PDC | NDEMTRQIQSLITSDTTLTAET | 22 | PP |
| 59 | 5euj | PDC | YAGLIDVFNDEDGHGLGLKAST | 22 | PP |
| 60 | 5npu | PDC | AAGAEEAVVELADALGCAVATM | 22 | CFX |
| 61 | 5npu | PDC | RAELCRQIQGLLNPNTTLIAET | 22 | PP |
| 62 | 5npu | PDC | YAGLMEVFNAEDGKGLGLKATT | 22 | PP |
| 63 | 5tma | PDC | AAAAEEAAVKFADALGGAVATM | 22 | CFX |
| 64 | 5tma | PDC | NAEIARQIEDLLTPNTTVIAET | 22 | PP |
| 65 | 5vrb | TK | GMGAIMNGLVLHGGVKPFGATF | 22 | PYR |
| 66 | 6dek | AHA | QSNPDFMKLAESMNVKGIRITN | 22 | PP |
| 67 | 1b0p | PFO | GCGETPYVRVITQLFGERMFIAN | 23 | PP |
| 68 | 1dtw | PDH | GKGRQMPVHYGCKERHFVTISSP | 23 | PP |
| 69 | 1itz | TK | AGSTLGWQKYVGAQGKAIGIDKF | 23 | CFX |
| 70 | 1n0h | AHA | GAGHMAEGYARASGKPGVVLVTS | 23 | PYR |
| 71 | 1n0h | AHA | PQTVIKKLSKVANDTGRHVIVTT | 23 | PP |
| 72 | 1upa | ALS | GYGIPAAIGAQMARPDQPTFLIA | 23 | PP |
| 73 | 1upa | ALS | REELLAALRKGAELGRPFLIEVP | 23 | PP |
| 74 | 1y9d | POX | YPFAEVSKAFKNTRYFLQIDIDP | 23 | CFX |
| 75 | 1ybh | AHA | GGVFAAEGYARSSGKPGICIATS | 23 | PYR |
| 76 | 2c31 | OxC | GANALDNTRMIVDMLKPRKRLDS | 23 | PP |
| 77 | 2c31 | OxC | YSNSLGVVRDFMLANPDISLVNE | 23 | PP |
| 78 | 2dji | POX | FPFSEVEGTFRNVDNFIQIDIDP | 23 | CFX |
| 79 | 2jlc | BFD | GEAQLAHRICDYLPEQGQLFVGN | 23 | PP |
| 80 | 2jlc | BFD | GLGHLALGLAKVSKQPVAVIVTS | 23 | PYR |
| 81 | 2jlc | BFD | SLVVRLIDALSQLPAGYPVYSNR | 23 | PP |
| 82 | 2jlc | BFD | WQELETAFADAWRTPTTTVIEMV | 23 | PP |
| 83 | 2nxw | PDC | AVGFAADAAARYSSTLGVAAVTY | 23 | PYR |
| 84 | 2pgn | ALS | GCGFPMALGAQLAEPNSRVFLGT | 23 | PP |
| 85 | 2pgn | ALS | GGAWMVNGYNYVKDRSAAVGAWH | 23 | PYR |
| 86 | 2q27 | OxC | GANTLDNARNIIDMYKPRRRLDC | 23 | PP |
| 87 | 2q27 | OxC | YFNALSAVRDVLRENQDIYLVNE | 23 | PP |
| 88 | 2r5n | TK | AGIADYWYKYVGLNGAIVGMTTF | 23 | CFX |
| 89 | 2vbf | PDC | FGKSAVDESLPSFLGIYNGKLSE | 23 | CFX |
| 90 | 2x7j | BFD | SAGFFALGLAKAKQRPVLLICTS | 23 | PYR |
| 91 | 3ahc | PhK | YAQDVRGLIYDRPNHDNFHVVGY | 23 | CFX |
| 92 | 3ai7 | PhK | YAHDVRGLIYDRPNHDNFNVHGY | 23 | CFX |
| 93 | 3exe | PDH | GQEACCVGLEAGINPTDHLITAY | 23 | PP |
| 94 | 3lq1 | BFD | SAGFFALGLAKASKRPVVLLCTS | 23 | PYR |
| 95 | 3lq1 | BFD | VDELEEAIDKASYHKGLDIIEVK | 23 | PP |
| 96 | 3m49 | TK | GAKDDTYEKVAKGAYVVSASKKE | 23 | CFX |
| 97 | 3m49 | TK | MGATFGWHRYVGLEGDVLGIDTF | 23 | CFX |
| 98 | 3rim | TK | AGVAQCWHQLVGDTGEIVSIEHY | 23 | CFX |
| 99 | 3uk1 | TK | AGVTDFWRKYVGLEGGVVGIDTF | 23 | CFX |
| 100 | 4c7v | TK | AGTTYGWAKYAGDHGVMIGIDEF | 23 | CFX |
| 101 | 4cok | PDC | GFSAEGYARANGAAAAIVTFSVG | 23 | PYR |
| 102 | 4k9q | BFD | CPSIVPLMQDVFRINQPDTFYTF | 23 | PP |
| 103 | 4k9q | BFD | GSTEETFLKDFPSDFNYVLALQE | 23 | PYR |

| | | | | | |
|---|---|---|---|---|---|
| 104 | 4k9q | BFD | LDELKTAYLEALSFKGTSVIVVP | 23 | PP |
| 105 | 4kxu | TK | QNMVSIAVGCATRNRTVPFCSTF | 23 | PYR |
| 106 | 4qq8 | OxC | AAAGHAAEGYARAGAKLGVALVT | 23 | PYR |
| 107 | 4rji | ALS | GVALPWAIGASLVKPGEKVVSVS | 23 | PP |
| 108 | 4rji | ALS | PDQLADVLRQGMNAEGPVIIDVP | 23 | PP |
| 109 | 4xeu | TK | AAHADYWYKYVGLDGRIIGMTSF | 23 | CFX |
| 110 | 4xeu | TK | LPHQARDVAQVADIARGGYVLKD | 23 | CFX |
| 111 | 5c4i | PFO | AGCGPALTYRLVAKAAGPNTIFI | 23 | PP |
| 112 | 5dx6 | ALS | AEALEPTLRAAMDVDGPAVVAIP | 23 | PP |
| 113 | 5dx6 | ALS | GVALPWAIGAWLVNPERKVVSVS | 23 | PP |
| 114 | 5erx | BFD | SNPVRDVALAGLDTRGIRVRSNR | 23 | PP |
| 115 | 5nd6 | TK | AATSFGWAKYIGLKGKHVGIDTF | 23 | CFX |
| 116 | 5nd6 | TK | AMGAICNGIALHKSGLIPYCATF | 23 | PYR |
| 117 | 5npu | PDC | GFSAEGYARANGVGAAVVTFSVG | 23 | PYR |
| 118 | 5tma | PDC | GFSAEGYARAKGAAAAVVTYSVG | 23 | PYR |
| 119 | 5vrb | TK | AGHTNGWYKYVGLNGAVVGINRF | 23 | CFX |
| 120 | 6a50 | BFD | GSNELPFLKDFPEDFRYILALQE | 23 | PYR |
| 121 | 6a50 | BFD | LEQLKGSLQEALSAKGPVLIEVS | 23 | PP |
| 122 | 6a50 | BFD | STSTTAQMWQRLNMRNPGSYYFC | 23 | PP |
| 123 | 6cin | PFO | PENCIQCNQCSLVCPHAAIRPYL | 23 | CFX |
| 124 | 6dek | AHA | GAGHMAEGYARASGKPGVVLVTS | 23 | PYR |
| 125 | 6dek | AHA | PQTLIKEISDQAQTYNKEVIVTT | 23 | PP |
| 126 | 1ay0 | TK | AVATRKLSETVLEDVYNQLPELIG | 24 | PYR |
| 127 | 1b0p | PFO | GMGKSQDVMNTAVKSGYWPLFRYD | 24 | PP |
| 128 | 1dtw | PDH | GEEGTHVGSAAALDNTDLVFGQYR | 24 | PP |
| 129 | 1n0h | AHA | GLTGGQIFNEMMSRQNVDTVFGYP | 24 | PYR |
| 130 | 1n0h | AHA | KQEELDAKLKEFVSTKGPVLLEVE | 24 | PP |
| 131 | 1ni4 | PDH | WNSEDAKGLIKSAIRDNNPVVVLE | 24 | PYR |
| 132 | 1ovm | PDC | ACYEIDRVLTTMLRERRPGYLMLP | 24 | PYR |
| 133 | 1ovm | PDC | GTSAFGAIDLRLPADVNFIVQPLW | 24 | PP |
| 134 | 1qpb | PDC | AESEKEVIDTILVLIKDAKNPVIL | 24 | CFX |
| 135 | 1qpb | PDC | GTSAFGINQTTFPNNTYGISQVLW | 24 | PP |
| 136 | 1r9j | TK | AIATRKASENCLAVLFPAIPALMG | 24 | PYR |
| 137 | 1r9j | TK | RQNTEPQSGSSIEGVRHGAYSVVD | 24 | CFX |
| 138 | 1y9d | POX | KIEQLPDVFEQAKAIAQHEPVLID | 24 | PP |
| 139 | 2dji | POX | APAAQDIDAAVELLNNSKRPVIYA | 24 | CFX |
| 140 | 2dji | POX | IEDMDRVMAEAVAANKAGHTVVID | 24 | PP |
| 141 | 2nxw | PDC | ALAACADEVLAAMRSATSPVLMVC | 24 | CFX |
| 142 | 2nxw | PDC | APAEIARVLGAARAQSRPVYLEIP | 24 | PYR |
| 143 | 2pan | AHA | GASHMAEGYTRATAGNIGVCLGTS | 24 | PYR |
| 144 | 2q27 | OxC | GIALARAIRVSVSGRPGGVYLDLP | 24 | PYR |
| 145 | 2r5n | TK | ARNAVRMAALMKQRQVMVYTHDSI | 24 | PYR |
| 146 | 2vbf | PDC | GTSFFGASTIFLKSNSRFIGQPLW | 24 | PP |
| 147 | 2vbf | PDC | TTEQVILSKIEESLKNAQKPVVIA | 24 | CFX |
| 148 | 2vbi | PDC | AAAKGFFPEDHAGFRGLYWGEVSN | 24 | CFX |
| 149 | 2vbi | PDC | CGFSAEGYARSNGAAAAVVTFSVG | 24 | PYR |
| 150 | 2vbi | PDC | GDSWFNAMRMTLPRGARVELEMQW | 24 | PP |
| 151 | 2vjy | PDC | GTSAFGINQTHFPNNTYGISQVLW | 24 | PP |
| 152 | 2vjy | PDC | PEAEEEVIENVLQLIKEAKNPVIL | 24 | CFX |
| 153 | 2x7j | BFD | SMPIRDVDTFFEKQDRPFRIYSNR | 24 | PP |
| 154 | 2x7j | BFD | YDSFLKDDELKRKLRPDVVIRFGP | 24 | CFX |
| 155 | 3ahc | PhK | AIFAGKQPAPTWVTLDEARAELEA | 24 | CFX |
| 156 | 3ai7 | PhK | AIIAGKQPAATWLTLDEARAELEK | 24 | CFX |
| 157 | 3duf | PDH | GQEASQIASHFALEKEDFILPGYR | 24 | PP |

| | | | | | |
|---|---|---|---|---|---|
| 158 | 3ey9 | POX | GLIGFSSGFHTMMNADTLVLLGTQ | 24 | CFX |
| 159 | 3ey9 | POX | GSGCAGAHKELVEFAGKIKAPIVH | 24 | CFX |
| 160 | 3lq1 | BFD | DSSIQKMVTECTGKKGVFVVGPID | 24 | CFX |
| 161 | 3lq1 | BFD | SMPIRDVDTYFSQIDKKIKMLANR | 24 | PP |
| 162 | 3m34 | TK | FELFEKQDKAYQERLLKGEVIGVE | 24 | CFX |
| 163 | 4c7v | TK | ATRAASQKAINALAKEVSSLWGGA | 24 | PYR |
| 164 | 4cok | PDC | GDSWFNAVRMKLPHGARVELEMQW | 24 | PP |
| 165 | 4k9q | BFD | ASVVAIADGLSQSLRKPVIVNIHT | 24 | PYR |
| 166 | 4xeu | TK | ARNAVRMSALMKQRVLYVFTHDSI | 24 | PYR |
| 167 | 5dx6 | ALS | ANAAFMAAAVGRITGKAGVALVTS | 24 | PYR |
| 168 | 5dx6 | ALS | GPGCSNLITGMATANSEGDPVVAL | 24 | PYR |
| 169 | 5dx6 | ALS | GYSPVEYEPAMWNSGNATLVHIDV | 24 | CFX |
| 170 | 5erx | BFD | GTGANQTMEQLGYFGTQVRASISL | 24 | PYR |
| 171 | 5euj | PDC | GDSWFNASRMPIPGGARVELEMQW | 24 | PP |
| 172 | 5hje | TK | AAGAVRLSALSEFPITWVATHDSI | 24 | PYR |
| 173 | 5npu | PDC | GDSWFNAMRMKLPHGARVELEMQW | 24 | PP |
| 174 | 5tma | PDC | GDSWFNAQRMKLPNGARVEYEMQW | 24 | PP |
| 175 | 6a50 | BFD | ACVVGIADGYAQASRKPAFINLHS | 24 | PYR |
| 176 | 6cin | PFO | AGCGETPYVKLVTQLFGDRMIIAN | 24 | PP |
| 177 | 6dek | AHA | GLTGGEIFHEMMLRHKVDTVFGYA | 24 | PYR |
| 178 | 6gua | PhK | HIVASKQPRQQWFTKEEAEKLATD | 24 | CFX |
| 179 | 1b0p | PFO | GIVAEYMQKVASLTGRSYKLFDYVG | 25 | CFX |
| 180 | 1dtw | PDH | WDVDTICKSVIKTGRLLISHEAPLT | 25 | CFX |
| 181 | 1itz | TK | RQKLPHLPGTSIEGVEKGGYTISDN | 25 | CFX |
| 182 | 1ni4 | PDH | SRPVGHCLEAAAVLSKEGVECEVIN | 25 | CFX |
| 183 | 1ovm | PDC | GTRFTDTLTAGFTHQLTPAQTIEVQ | 25 | CFX |
| 184 | 1umb | PDH | GTVMPEVLQAAAELAKAGVSAEVLD | 25 | CFX |
| 185 | 1umb | PDH | WDYEAVMNSVAKTGRVVLVSDAPRH | 25 | CFX |
| 186 | 1y9d | POX | ANRAAQKPANEALAQADVVLFVGNN | 25 | CFX |
| 187 | 1y9d | POX | GDINLNANRHLKLTPSNRHITSNLF | 25 | PP |
| 188 | 1ybh | AHA | GEAIPPQYAIKVLDELTDGKAIIST | 25 | PP |
| 189 | 2dji | POX | GNSTQTSIRHLHMTPKNMWRTSPLF | 25 | PP |
| 190 | 2j9f | PDH | GKGRQMPVHYGCKERHFVTISSPLA | 25 | PP |
| 191 | 2j9f | PDH | WDVDTICKSVIKTGRLLISHEAPLT | 25 | CFX |
| 192 | 2jlc | BFD | ARWLVSTIDHALGTLHAGGVHINCP | 25 | PYR |
| 193 | 2jlc | BFD | SSLTGKRLLQWQASCEPEEYWIVDD | 25 | CFX |
| 194 | 2nxw | PDC | PMDIARAVNDRVRAGQEPLLIAADM | 25 | PP |
| 195 | 2pan | AHA | APAFEQAKALMAQYRVPVVVEVILE | 25 | PP |
| 196 | 2vbi | PDC | AISAMNALGGAYAENLPVILISGAP | 25 | PYR |
| 197 | 2vbi | PDC | KLRAANALAATETLADKLQCAVTIM | 25 | CFX |
| 198 | 2x7j | BFD | ASWDEFKTAYAPQADKPGLHLIEIK | 25 | PP |
| 199 | 2x7j | BFD | MPVSKPVFLWLKDDPTIQQIVIDED | 25 | CFX |
| 200 | 2x7j | BFD | PITHYIGSFIDEFALSGITDAVVCP | 25 | PYR |
| 201 | 2x7j | BFD | RESLSDVAEMLAEAEKGMIVCGELH | 25 | CFX |
| 202 | 3duf | PDH | GAMVHESLKAAAELEKEGISAEVVD | 25 | CFX |
| 203 | 3duf | PDH | LDIETIIGSVEKTGRAIVVQEAQRQ | 25 | CFX |
| 204 | 3exe | PDH | SRPVGHCLEAAAVLSKEGVECEVIN | 25 | CFX |
| 205 | 3ey9 | POX | GPGNLHLINGLFDCHRNHVPVLAIA | 25 | PYR |
| 206 | 3ey9 | POX | GTPTVWAARYLKMNGKRRLLGSFNH | 25 | PP |
| 207 | 3m34 | TK | KLKALNEPVFGDVKNGAYLLKESKE | 25 | CFX |
| 208 | 4cok | PDC | ALSAFNALGGAYAENLPVILISGAP | 25 | PYR |
| 209 | 4kxu | TK | TKNSTFSEIFKKEHPDRFIECYIAE | 25 | PYR |
| 210 | 4qq8 | OxC | GVGFGTALGAQVADLEAGRRTILVT | 25 | PP |
| 211 | 4rji | ALS | GPAADDAISAAIAKIQTAKLPVVLV | 25 | CFX |

| | | | | | |
|---|---|---|---|---|---|
| 212 | 4rji | ALS | GYDPIEYDPKFWNINGDRTIIHLDE | 25 | CFX |
| 213 | 5ahk | AHA | GAAFAASAVSRVTHHKTLGLALATS | 25 | PYR |
| 214 | 5ahk | AHA | GSMGFAIPAAIGACYAGKKPIIVIT | 25 | PP |
| 215 | 5c4i | PFO | GIMSELARMVADGELDAEFVHGEGE | 25 | PYR |
| 216 | 5dx6 | ALS | ALHPLRIVRAMQDIVNSDVTLTVDM | 25 | PP |
| 217 | 5dx6 | ALS | GAAPDDAIDQVAKLIAQAKNPIFLL | 25 | CFX |
| 218 | 5erx | BFD | PSTTQARVVVDELIRGGVRDVVLCP | 25 | PYR |
| 219 | 5euj | PDC | AAAKGFFPEDHPNFRGLYWGEVSSE | 25 | CFX |
| 220 | 5euj | PDC | QTSVTAAVDAAVEWLQDRQNVVMLV | 25 | CFX |
| 221 | 5hje | TK | AIAAAIDEAKKVTNKPTLVRLTTTI | 25 | PP |
| 222 | 5nd6 | TK | GLRAAIAQAKAVKDKPTLIKVSTLI | 25 | PP |
| 223 | 5npu | PDC | AAAKSFFPEDHPGYVGTYWGEVSSP | 25 | CFX |
| 224 | 5npu | PDC | ALSAFNAIGGAYAENLPVILISGAP | 25 | PYR |
| 225 | 5tma | PDC | ALSAFDAIGGAYAENLPVILISGAP | 25 | PYR |
| 226 | 6a50 | BFD | GVEAGETNVDAANLPRPLVKWSYEP | 25 | PYR |
| 227 | 6dek | AHA | QQELKSGVKEFLDATEPVLLEVIVE | 25 | PP |
| 228 | 1ay0 | TK | AGIAKAIAQAKLSKDKPTLIKMTTTI | 26 | PP |
| 229 | 1ay0 | TK | FFTFDKQPLEYRLSVLPDNVPIMSVE | 26 | CFX |
| 230 | 1b0p | PFO | AYVGIYDILEGIKDGGTFVLNSPWSS | 26 | CFX |
| 231 | 1dtw | PDH | GTQVHVIREVASMAKEKLGVSCEVID | 26 | CFX |
| 232 | 1dtw | PDH | RSPFQAKGLLLSCIEDKNPCIFFEPK | 26 | PYR |
| 233 | 1itz | TK | ADGNETAGAYKVAVLNRKRPSILALS | 26 | PYR |
| 234 | 1itz | TK | GDATRNLSQQCLNALANVVPGLIGGS | 26 | PYR |
| 235 | 1itz | TK | GMGAICNGIALHSPGFVPYCATFFVF | 26 | PYR |
| 236 | 1n0h | AHA | GGAILPVYDAIHNSDKFNFVLPKHEQ | 26 | PYR |
| 237 | 1n0h | AHA | GTMGYGLPAAIGAQVAKPESLVIDID | 26 | PP |
| 238 | 1ni4 | PDH | VRDAINQGMDEELERDEKVFLLGEEV | 26 | PYR |
| 239 | 1ovm | PDC | GSIGYTLAAAFGAQTACPNRRVIVLT | 26 | PP |
| 240 | 1ovm | PDC | SACLKAFRDAAENKLAMSKRTALLAD | 26 | CFX |
| 241 | 1qpb | PDC | ATAPAEIDRCIRTTYVTQRPVYLGLP | 26 | PYR |
| 242 | 1r9j | TK | AMCAILNGLDAHDGIIPFGGTFLNFI | 26 | PYR |
| 243 | 1r9j | TK | GYALGAVRLAAISHHRVIYVATHDSI | 26 | PYR |
| 244 | 1r9j | TK | QELFDAQPDTYRQAVLPAGVPVVSVE | 26 | CFX |
| 245 | 1r9j | TK | SDQTETSGAWAVALSSIHTPTVLCLS | 26 | PYR |
| 246 | 1umb | PDH | STPYDAKGLLKAAIRDEDPVVFLEPK | 26 | PYR |
| 247 | 1upa | ALS | GYDYAEDLRPSMWQKGIEKKTVRISP | 26 | CFX |
| 248 | 1y9d | POX | AGPGGTHLMNGLYDAREDHVPVLALI | 26 | PYR |
| 249 | 1y9d | POX | ATMGVGIPGAIAAKLNYPERQVFNLA | 26 | PP |
| 250 | 1ybh | AHA | GAMGFGLPAAIGASVANPDAIVVDID | 26 | PP |
| 251 | 1ybh | AHA | GDPAQEDEIFPNMLLFAAACGIPAAR | 26 | PP |
| 252 | 1ybh | AHA | GGASMEIHQALTRSSSIRNVLPRHEQ | 26 | PYR |
| 253 | 2bp7 | PDH | LDLDTIVESVKKTGRCVVVHEATRTC | 26 | CFX |
| 254 | 2c31 | OxC | TPAELKAALEEAVASGKPCLINAMID | 26 | PP |
| 255 | 2dji | POX | ATMGIAIPGGLGAKNTYPDRQVWNII | 26 | PP |
| 256 | 2dji | POX | GGPGASHLINGLYDAAMDNIPVVAIL | 26 | PYR |
| 257 | 2j9f | PDH | GTQVHVIREVASMAKEKLGVSCEVID | 26 | CFX |
| 258 | 2j9f | PDH | TQIPQAVGAAYAAKRANANRVVICYF | 26 | PP |
| 259 | 2jlc | BFD | GSRSTPLTLAAAENSAFIHHTHFDER | 26 | PYR |
| 260 | 2nxw | PDC | AGMGFGVPAGIGAQCVSGGKRILTVV | 26 | PP |
| 261 | 2nxw | PDC | RAELKAALDKAFATRGRFQLIEAMIP | 26 | PP |
| 262 | 2pan | AHA | GAAINPFYSAMRKHGGIRHILARHVE | 26 | PYR |
| 263 | 2pan | AHA | GPLGWTIPAALGVCAADPKRNVVAIS | 26 | PP |
| 264 | 2pan | AHA | GYGVDHVKVAEGLGCKAIRVFKPEDI | 26 | PP |
| 265 | 2q27 | OxC | TTDELRHALTTGIQSRKPTIINVVID | 26 | PP |

| | | | | | |
|---|---|---|---|---|---|
| 266 | 2vbf | PDC | ENEFVSVMKEAQADVNRMYWIELVLE | 26 | PP |
| 267 | 2vbf | PDC | GSIGYTFPAALGSQIADKESRHLLFI | 26 | PP |
| 268 | 2vbi | PDC | GHIGWSVPSAFGNAMGSQDRQHVVMV | 26 | PP |
| 269 | 2vbi | PDC | PKELTEAIARAKANTRGPTLIECQID | 26 | PP |
| 270 | 2x7j | BFD | APQAINQHFLFGNFVKFFTDSALPEE | 26 | PYR |
| 271 | 2x7j | BFD | GSRSTPLAVLCAAHPDISVHVQIDER | 26 | PYR |
| 272 | 3ahc | PhK | GELGYALSHAYGAVMNNPSLFVPCII | 26 | PP |
| 273 | 3ai7 | PhK | GELGYALSHAYGAIMDNPSLFVPAIV | 26 | PP |
| 274 | 3duf | PDH | STPYDAKGLLISAIRDNDPVIFLEHL | 26 | PYR |
| 275 | 3exe | PDH | VRDAINQGMDEELERDEKVFLLGEEV | 26 | PYR |
| 276 | 3exe | PDH | WNSEDAKGLIKSAIRDNNPVVVLENE | 26 | PYR |
| 277 | 3ey9 | POX | GSMANAMPQALGAQATEPERQVVAMC | 26 | PP |
| 278 | 3lq1 | BFD | APQAMDQLHLYGSHVKDFTDMALPEN | 26 | PYR |
| 279 | 3lq1 | BFD | GANGIDGVVSSALGASVVFQPMFLLI | 26 | PP |
| 280 | 3lq1 | BFD | GSRSTPLALMMAEHPILKIYVDVDER | 26 | PYR |
| 281 | 3lq1 | BFD | YDAFLKEAEIIDKLTPEVVIRFGSMP | 26 | CFX |
| 282 | 3m34 | TK | YEEINKALEQAKKSTKPCLIIAKTTI | 26 | PP |
| 283 | 3rim | TK | ALATRAASGAVLSALGPKLPELWGGS | 26 | PYR |
| 284 | 4c7v | TK | ERFEAQSEEYKNTVIPPELKKRMTIE | 26 | CFX |
| 285 | 4cok | PDC | AAAKSFFPEDHPGYRGHYWGEVSSPG | 26 | CFX |
| 286 | 4cok | PDC | APAKIDHVIRTALREKKPAYLEIACN | 26 | PYR |
| 287 | 4cok | PDC | GHIGWSVPAAFGNALAAPERQHVLMV | 26 | PP |
| 288 | 4k9q | BFD | PDKVKEFAQRITASKNPLLIYGSDIA | 26 | CFX |
| 289 | 4kxu | TK | ATRKAYGQALAKLGHASDRIIALDGD | 26 | PYR |
| 290 | 4qq8 | OxC | AGGGFTNAVTPIANARTDRTPVLFLT | 26 | PYR |
| 291 | 5ahk | AHA | GGMITHLVDSINLLGKTKLVSMHHEQ | 26 | PYR |
| 292 | 5ahk | AHA | VGNNQMWAAHTLRLNAQQAMHHSGGL | 26 | PP |
| 293 | 5c4i | PFO | EESLIKGVPILKGIRPGSTLVVNTKR | 26 | CFX |
| 294 | 5c4i | PFO | GCVAVAHGVRLADVDVICSYPIRPYT | 26 | PYR |
| 295 | 5euj | PDC | GAISAMNAIGGAYAENLPVILISGSP | 26 | PYR |
| 296 | 5euj | PDC | GHIGWSVPSAFGNAVGSPERRHIMMV | 26 | PP |
| 297 | 5nd6 | TK | GLATRQHSQTMINALAPALPGLIGGS | 26 | PYR |
| 298 | 5npu | PDC | GHIGWSVPATFGYAVAEPERRNVLMV | 26 | PP |
| 299 | 5tma | PDC | APAKIDHVIKTALREKKPVYLEIACN | 26 | PYR |
| 300 | 5tma | PDC | GHIGWSVPAAFGYAVGAPERRNILMV | 26 | PP |
| 301 | 5tma | PDC | YAALMEVFNGNGGYDSGAGKGLKAKT | 26 | PP |
| 302 | 5vrb | TK | TSVFDRQDAAYQAAVLPEGLPRIAVE | 26 | CFX |
| 303 | 6a50 | BFD | DQDLDILVKALNSASNPAIVLGPDVD | 26 | CFX |
| 304 | 6cin | PFO | FSAEHFLKVLPASVKRIAVLDRTKEP | 26 | CFX |
| 305 | 6cin | PFO | PSYVGRYNLLEGIKPGGIFLLNSTWS | 26 | CFX |
| 306 | 6dek | AHA | GGAILPVFDAIYNSDKFKFVLPRHEQ | 26 | PYR |
| 307 | 6dek | AHA | GTMGYGLPAAIGAQVAKPDAIVIDID | 26 | PP |
| 308 | 6gua | PhK | GELGYVLSHATGAILDQPEQIAFAVV | 26 | PP |
| 309 | 6gua | PhK | GFHGYEDLIESIFYQRGHDGLIVHGY | 26 | CFX |
| 310 | 1b0p | PFO | AAAGKRTGKKDLARMVMTYGYVYVATV | 27 | PP |
| 311 | 1b0p | PFO | AGQKDGLLGQIAAMSDLYTKKSVWIFG | 27 | PP |
| 312 | 1b0p | PFO | GNTATAHVAYAMSEVAAIYPITPSSTM | 27 | PYR |
| 313 | 1b0p | PFO | LEDMDKHLPSGIKRTIANKKLKFYNID | 27 | CFX |
| 314 | 1dtw | PDH | NLFQSVTSALDNSLAKDPTAVIFGEDV | 27 | PYR |
| 315 | 1itz | TK | TDYMRGAMRISALSEAGVIYVMTHDSI | 27 | PYR |
| 316 | 1itz | TK | WELFDEQSDEYKESVLPAAVTARISIE | 27 | CFX |
| 317 | 1n0h | AHA | AVGARFDDRVTGNISKFAPEARRAAAE | 27 | CFX |
| 318 | 1n0h | AHA | GVGQHQMWAAQHWTWRNPHTFITSGGL | 27 | PP |
| 319 | 1umb | PDH | GHEAAQVAIAHAIRPGFDWVFPYYRDH | 27 | PP |

| | | | | | |
|---|---|---|---|---|---|
| 320 | 1umb | PDH | TMVQALNRALDEEMAKDPRVVVLGEDV | 27 | PYR |
| 321 | 1upa | ALS | VHQVIDSMNTVMEEAAEPGEGTIVSDI | 27 | PP |
| 322 | 1y9d | POX | GIGARKAGKELEQLSKTLKIPLMSTYP | 27 | CFX |
| 323 | 1ybh | AHA | GGGCLNSSDELGRFVELTGIPVASTLM | 27 | CFX |
| 324 | 1ybh | AHA | GVGQHQMWAAQFYNYKKPRQWLSSGGL | 27 | PP |
| 325 | 1ybh | AHA | VTKKADLREAIQTMLDTPGPYLLDVIC | 27 | PP |
| 326 | 2c31 | OxC | GKGAAYAQCDDEIRALVEETGIPFLPM | 27 | CFX |
| 327 | 2dji | POX | GTLSSLMDAMGEEENNVKFLQVKHEEV | 27 | PYR |
| 328 | 2j9f | PDH | NLFQSVTSALDNSLAKDPTAVIFGEDV | 27 | PYR |
| 329 | 2jlc | BFD | GASGIDGLLSTAAGVQRASGKPTLAIV | 27 | PP |
| 330 | 2nxw | PDC | GVAGDAEITRLVEESDGLFLLGAILSD | 27 | CFX |
| 331 | 2pan | AHA | GPAGTDMITALYSASADSIPILCITGQ | 27 | PYR |
| 332 | 2pan | AHA | TIGLSQIAAAQMLHVFKDRHWINCGQA | 27 | PP |
| 333 | 2pgn | ALS | GSRLSDWGIAQGYITKMPKFVHVDTDP | 27 | CFX |
| 334 | 2q27 | OxC | GARLNWLLAHGKKGWAADTQFIQLDIE | 27 | CFX |
| 335 | 2q27 | OxC | GKGAAYSQADEQLREFIESAQIPFLPM | 27 | CFX |
| 336 | 2r5n | TK | TDAFDKQDAAYRESVLPKAVTARVAVE | 27 | CFX |
| 337 | 2vbf | PDC | AENATYEIDRVLSQLLKERKPVYINLP | 27 | PYR |
| 338 | 2vbi | PDC | AHSAPAKIDHVIRTALRERKPAYLDIA | 27 | PYR |
| 339 | 2vbi | PDC | HTSLKAAVDATVALLEKSASPVMLLGS | 27 | CFX |
| 340 | 2vjy | PDC | GALLSDFNTGSFSYSYKTKNIVEFHSD | 27 | CFX |
| 341 | 2x7j | BFD | GANGIDGVVSSAMGVCEGTKAPVTLVI | 27 | PP |
| 342 | 3duf | PDH | GAQYIQAAGVALGLKMRGKKAVAITYT | 27 | PP |
| 343 | 3duf | PDH | TMVQAITDALRIELKNDPNVLIFGEDV | 27 | PYR |
| 344 | 3ey9 | POX | GDSLNGLSDSLNRMGTIEWMSTRHEEV | 27 | PYR |
| 345 | 3ey9 | POX | PEQIPQVLAIAMRKAVLNRGVSVVVLP | 27 | PYR |
| 346 | 3m34 | TK | ADGVENVKAWQIALNADIPSAFVLSRQ | 27 | PYR |
| 347 | 3m49 | TK | IEAIAKAIEEAKADEKRPTLIEVRTTI | 27 | PP |
| 348 | 3m49 | TK | MDRFEAQTAEYKESVLPKAVTKRFAIE | 27 | CFX |
| 349 | 3rim | TK | LEWFEAQPYEYRDAVLPPTVSARVAVE | 27 | CFX |
| 350 | 3uk1 | TK | SSDVFDRQDAEYRERVLPHGVRRVAIE | 27 | CFX |
| 351 | 4kxu | TK | GAGVTLHEALAAAELLKKEKINIRVLD | 27 | CFX |
| 352 | 4kxu | TK | GHSVEELCKAFGQAKHQPTAIIAKTFK | 27 | PP |
| 353 | 4kxu | TK | SDGVATEKAVELAANTKGICFIRTSRP | 27 | PYR |
| 354 | 4qq8 | OxC | GLHGIHIDTIFQACLDHDVPIIDTRHE | 27 | PYR |
| 355 | 4qq8 | OxC | GSEASRTARKTALSAFVAATGVPVFAD | 27 | CFX |
| 356 | 4xeu | TK | TSVYEQQDESYKQSVLPVEVGARIAIE | 27 | CFX |
| 357 | 5ahk | AHA | SMQAFSSALESFLESPRPLLLEVSMSD | 27 | PP |
| 358 | 5dx6 | ALS | GIPGAKIDKVFDSLLDSSIRIIPVRHE | 27 | PYR |
| 359 | 5euj | PDC | GSKLRAAAAEKQAVALADRLGCAVTIM | 27 | CFX |
| 360 | 5hje | TK | DQLTFDKQSEEYKLSVLPDGVPILSVE | 27 | CFX |
| 361 | 5nd6 | TK | WELFEEQSAEYKESVLPSDVTARVSVE | 27 | CFX |
| 362 | 5npu | PDC | AEDAPALIDHAIRTALREKKPAYIEIA | 27 | PYR |
| 363 | 5tma | PDC | GEVSYPGVEKTMKEADAVIALAPVFND | 27 | CFX |
| 364 | 5vrb | TK | AKAETVATRKASQNSIEILAKELPELV | 27 | PYR |
| 365 | 6cin | PFO | AAGGKFTKKKDLGLMAMSYGYVYVASV | 27 | PP |
| 366 | 6cin | PFO | AEEMDSRLPADMKRTIATKKLKFYNID | 27 | CFX |
| 367 | 6cin | PFO | GNTAAAHVAYAMSEVATIYPITPSSPM | 27 | PYR |
| 368 | 6cin | PFO | VQEVMDLALVAHLATLKARVPFVHFFD | 27 | PYR |
| 369 | 6dek | AHA | ALGARFDDRVTGNISKFAPEAKLAASE | 27 | CFX |
| 370 | 6dek | AHA | AVSEFTSEAIKRAANILNKAKKPIIYA | 27 | CFX |
| 371 | 6dek | AHA | GVGQHQMWAAQHFTWTQPRTMITSGGL | 27 | PP |
| 372 | 1b0p | PFO | AFEADGRFPLGTSQFEKRGVAINVPQWV | 28 | CFX |
| 373 | 1b0p | PFO | AIAAHALSIFGDHQDIYAARQTGFAMLA | 28 | PYR |

| | | | | | |
|---|---|---|---|---|---|
| 374 | 1b0p | PFO | GEEADDWAAQGRKNIFGQTLTIREMQSE | 28 | PYR |
| 375 | 1dtw | PDH | LATQIPQAVGAAYAAKRANANRVVICYF | 28 | PP |
| 376 | 1n0h | AHA | AIGTDAFQEADVVGISRSCTKWNVMVKS | 28 | PYR |
| 377 | 1n0h | AHA | GRGGIIHFEVSPKNINKVVQTQIAVEGD | 28 | CFX |
| 378 | 1n0h | AHA | VEELPLRINEAFEIATSGRPGPVLVDLP | 28 | PYR |
| 379 | 1ni4 | PDH | GMGTSVERAAASTDYYKRGDFIPGLRVD | 28 | PP |
| 380 | 1ovm | PDC | AEQLADVLEKVAHHERLSLIEVMLPKAD | 28 | PP |
| 381 | 1ovm | PDC | FLVLRHGLKHALQKWVKEVPMAHATMLM | 28 | CFX |
| 382 | 1ovm | PDC | GVGELSAMNGIAGSYAEHVPVLHIVGAP | 28 | PYR |
| 383 | 1qpb | PDC | GVGELSALNGIAGSYAEHVGVLHVVGVP | 28 | PYR |
| 384 | 1umb | PDH | IASHVPPAAGAAISMKLLRTGQVAVCTF | 28 | PP |
| 385 | 1upa | ALS | GFFRHYGVLFARADQPFGFLTSAGCSSF | 28 | PP |
| 386 | 1upa | ALS | GVVADGWQKAADQAAALLAEAKHPVLVV | 28 | CFX |
| 387 | 1upa | ALS | GVVGREAASILFDEVEGIDFVLTRHEFT | 28 | PYR |
| 388 | 1y9d | POX | GGSINSIMDALSAERDRIHYIQVRHEEV | 28 | PYR |
| 389 | 2bp7 | PDH | GFGAELVSLVQEHCFHHLEAPIERVTGW | 28 | CFX |
| 390 | 2bp7 | PDH | MTMIQALRSAMDVMLERDDNVVVYGQDV | 28 | PYR |
| 391 | 2c31 | OxC | GTWGVMGIGMGYCVAAAAVTGKPVIAVE | 28 | PP |
| 392 | 2c31 | OxC | IKDIPIGIARAVRTAVSGRPGGVYVDLP | 28 | PYR |
| 393 | 2dji | POX | GIGTMGHGPAVQELARKIKAPVITTGKN | 28 | CFX |
| 394 | 2jlc | BFD | VSAFNRRWAAVILEALTRHGVRHICIAP | 28 | PYR |
| 395 | 2nxw | PDC | GAGAFNMVNAVAGAYAEKSPVVVISGAP | 28 | PYR |
| 396 | 2nxw | PDC | VEVRRYGLEAKVAELAQRLGVPVVTTFM | 28 | CFX |
| 397 | 2pan | AHA | AALVPRVLQQAFHLMRSGRPGPVLVDLP | 28 | PYR |
| 398 | 2pgn | ALS | GNHTLPMFGGAILQRPRRLVTSMAEGIL | 28 | PP |
| 399 | 2q27 | OxC | GTWGVMGIGMGYAIGASVTSGSPVVAIE | 28 | PP |
| 400 | 2r5n | TK | GMTAIANGISLHGGFLPYTSTFLMFVEY | 28 | PYR |
| 401 | 2vbf | PDC | GHEVISFGLEKTVTQFVSETKLPITTLN | 28 | CFX |
| 402 | 2vbf | PDC | GVGELSAINGLAGSYAENLPVVEIVGSP | 28 | PYR |
| 403 | 2vjy | PDC | GVGELSALNGIAGSYAEHVGVLHVVGVP | 28 | PYR |
| 404 | 3exe | PDH | GMGTSVERAAASTDYYKRGDFIPGLRVD | 28 | PP |
| 405 | 3lq1 | BFD | HEQVLTDYLAAFIEELVQAGVKEAIISP | 28 | PYR |
| 406 | 3m34 | TK | GKDLATRDSNGEILNVLAKNLEGFLGGS | 28 | PYR |
| 407 | 3m49 | TK | GSKAATRNSSGAVINAIAESVPSFFGGS | 28 | PYR |
| 408 | 4k9q | BFD | GAPVFRYYPWIAGQFIPEGSTLLQVSDD | 28 | CFX |
| 409 | 4k9q | BFD | PEDVPGAFMRAYATAMQQPQGPVFLSLP | 28 | PYR |
| 410 | 4kxu | TK | AAFFTRAFDQIRMAAISESNINLCGSHC | 28 | PYR |
| 411 | 4qq8 | OxC | GARFGLNTGHGSGQLIPHSAQVIQVDPD | 28 | CFX |
| 412 | 4qq8 | OxC | SVESFSAALAQALAHNRPACINVAVALD | 28 | PP |
| 413 | 4rji | ALS | GSHAIWMSRYFRSYEPLTLMISNGMQTL | 28 | PP |
| 414 | 4xeu | TK | ANKGETIASRKASQNALNAFGPLLPELL | 28 | PYR |
| 415 | 4xeu | TK | GMSAIMNGVALHGGFIPYGATFLIFMEY | 28 | PYR |
| 416 | 5ahk | AHA | VLGSRMDVRQTGAQPEDFARNAEIIQID | 28 | CFX |
| 417 | 5c4i | PFO | ASIGWPVDLMNKVRKGLNQEGPAYIHIH | 28 | PP |
| 418 | 5c4i | PFO | MNKDARDIVVALTEAAAKEGKYVQAWEN | 28 | CFX |
| 419 | 5dx6 | ALS | GSFHIWIARYLYSFRARQVMISNGQQTM | 28 | PP |
| 420 | 5tma | PDC | AAELEEAIKVALDNTDGPTLIECFIARE | 28 | PP |
| 421 | 6a50 | BFD | AAGGLGFALPAAIGVQLAEPERQVIAVI | 28 | PP |
| 422 | 6a50 | BFD | GAPVFRYVFYDPGQYLKPGTRLISVTCD | 28 | CFX |
| 423 | 6cin | PFO | AEIADEWAAHGRKNIFGKTLQVAEMQSE | 28 | PYR |
| 424 | 6cin | PFO | ASGEVKDLLLDIDRQKDYLTKKSIWIIG | 28 | PP |
| 425 | 6cin | PFO | ATPGIVAQVMEQVAGLTGRHYHLFDYAG | 28 | CFX |
| 426 | 6dek | AHA | AIGTDAFQEADIVGISRSCTKWNVMVKN | 28 | PYR |
| 427 | 6dek | AHA | GPGATNVITPMADALMDGVPLVVFSGQV | 28 | PYR |

| | | | | | |
|---|---|---|---|---|---|
| 428 | 6dek | AHA | GRGGILHFEISPKNINKVVEATEAIEGD | 28 | CFX |
| 429 | 6dek | AHA | VAELPRRINEAFEIATTGRPGPVLVDLP | 28 | PYR |
| 430 | 1ay0 | TK | ADGNEVSAAYKNSLESKHTPSIIALSRQN | 29 | PYR |
| 431 | 1ay0 | TK | GVGEDGPTHQPIETLAHFRSLPNIQVWRP | 29 | PYR |
| 432 | 1b0p | PFO | ASQGLLLMIPNMYKISGELLPGVFHVTAR | 29 | PYR |
| 433 | 1b0p | PFO | GDGWAYDIGYGGLDHVLASGEDVNVFVMD | 29 | PP |
| 434 | 1dtw | PDH | AFGGVFRCTVGLRDKYGKDRVFNTPLCEQ | 29 | PYR |
| 435 | 1dtw | PDH | GCVGHGALYHSQSPEAFFAHCPGIKVVIP | 29 | PYR |
| 436 | 1itz | TK | GLGEDGPTHQPIEHLVSFRAMPNILMLRP | 29 | PYR |
| 437 | 1n0h | AHA | GDASFNMTLTELSSAVQAGTPVKILILNN | 29 | PP |
| 438 | 1ni4 | PDH | GASAGVAAQHSQCFAAWYGHCPGLKVVSP | 29 | PYR |
| 439 | 1r9j | TK | ASGSEVSLAVDAAKALSGELRVRVVSMPC | 29 | CFX |
| 440 | 1r9j | TK | GVGEDGPTHQPVELVAALRAMPNLQVIRP | 29 | PYR |
| 441 | 1umb | PDH | ATKADPNKGRQMPEHPGSKALNFFTVASP | 29 | PP |
| 442 | 1umb | PDH | GGGVRGGHHHSQSPEAHFVHTAGLKVVAV | 29 | PYR |
| 443 | 1upa | ALS | GAAAIRSGAVPAIRALAERLNIPVITTYI | 29 | CFX |
| 444 | 1upa | ALS | RPHEITDLVDSAVNAAMTEPVGPSFISLP | 29 | PYR |
| 445 | 1y9d | POX | GDGGASMTMQDLATQVQYHLPVINVVFTN | 29 | PP |
| 446 | 1ybh | AHA | AFGVRFDDRVTGKLEAFASRAKIVHIDID | 29 | CFX |
| 447 | 1ybh | AHA | DVEDIPRIIEEAFFLATSGRPGPVLVDVP | 29 | PYR |
| 448 | 1ybh | AHA | GDGSFIMNVQELATIRVENLPVKVLLLNN | 29 | PP |
| 449 | 1ybh | AHA | GPGATNLVSGLADALLDSVPLVAITGQVP | 29 | PYR |
| 450 | 1ybh | AHA | RRMIGTDAFQETPIVEVTRSITKHNYLVM | 29 | PYR |
| 451 | 2bp7 | PDH | GGGIYGGQTHSQSPEAMFTQVCGLRTVMP | 29 | PYR |
| 452 | 2c31 | OxC | APGFLNGVTSLAHATTNCFPMILLSGSSE | 29 | PYR |
| 453 | 2c31 | OxC | GARLNWLMQHGKGKTWGDELKKYVQIDIQ | 29 | CFX |
| 454 | 2c31 | OxC | GDSAFGFSGMELETICRYNLPVTVIIMNN | 29 | PP |
| 455 | 2c31 | OxC | GVVGIPITNLARMWQDDGQRFYSFRHEQH | 29 | PYR |
| 456 | 2dji | POX | AYAEQLPKLVDEAARMAIAKRGVAVLEVP | 29 | PYR |
| 457 | 2dji | POX | GDGAFSMTYPDVVTNVRYNMPVINVVFSN | 29 | PP |
| 458 | 2j9f | PDH | AFGGVFRCTVGLRDKYGKDRVFNTPLCEQ | 29 | PYR |
| 459 | 2j9f | PDH | GCVGHGALYHSQSPEAFFAHCPGIKVVIP | 29 | PYR |
| 460 | 2j9f | PDH | RSPFQAKGLLLSCIEDKNPCIFFEPKILY | 29 | PYR |
| 461 | 2jlc | BFD | ANQAIRQPGMFASHPTHSISLPRPTQDIP | 29 | PYR |
| 462 | 2nxw | PDC | GDGAFQMTGWELGNCRRLGIDPIVILFNN | 29 | PP |
| 463 | 2nxw | PDC | GIPGDFALPFFKVAEETQILPLHTLSHEP | 29 | PYR |
| 464 | 2pan | AHA | GDFDFQFLIEELAVGAQFNIPYIHVLVNN | 29 | PP |
| 465 | 2pan | AHA | GGGVINADAAALLQQFAELTSVPVIPTLM | 29 | CFX |
| 466 | 2pan | AHA | GIGNRFANRHTGSVEKYTEGRKIVHIDIE | 29 | CFX |
| 467 | 2pgn | ALS | CVGNLLLHAAMQEARTGRIPAVHIGLNSD | 29 | PYR |
| 468 | 2pgn | ALS | GDGMPASMFRAMAEVRKVQRPEDIIVTDI | 29 | PP |
| 469 | 2pgn | ALS | GGGVARSGGSEALLKLAEMVGVPVVTTST | 29 | CFX |
| 470 | 2pgn | ALS | GRLAGRSEAAQQVPWQSFTPIARSTQRVE | 29 | PYR |
| 471 | 2pgn | ALS | RLDKVGEAIHEAFRVAEGHPAGPAYVDIP | 29 | PYR |
| 472 | 2q27 | OxC | APGFLNGLTALANATVNGFPMIMISGSSD | 29 | PYR |
| 473 | 2q27 | OxC | GDSAFGFSGMEIETICRYNLPVTIVIFNN | 29 | PP |
| 474 | 2q27 | OxC | GVVGIPVTDMARHAQAEGIRYIGFRHEQS | 29 | PYR |
| 475 | 2r5n | TK | GLGEDGPTHQPVEQVASLRVTPNMSTWRP | 29 | PYR |
| 476 | 2vbi | PDC | GDGSFQLTAQEVAQMVRYELPVIIFLINN | 29 | PP |
| 477 | 2vjy | PDC | INTAPAEIDRCIRTTYVSQRPVYLGLPAN | 29 | PYR |
| 478 | 3duf | PDH | AGIAANVVAEINERAILSLEAPVLRVAAP | 29 | CFX |
| 479 | 3duf | PDH | GGGVHTPELHSDSLEGLVAQQPGLKVVIP | 29 | PYR |
| 480 | 3exe | PDH | GASAGVAAQHSQCFAAWYGHCPGLKVVSP | 29 | PYR |
| 481 | 3ey9 | POX | GDGGFSMLMGDFLSVVQMKLPVKIVVFNN | 29 | PP |

| | | | | | |
|---|---|---|---|---|---|
| 482 | 3m34 | TK | GVGEDGPTHQPIEQLSTFRAMPNFLTFRP | 29 | PYR |
| 483 | 3m49 | TK | AVGEDGPTHEPIEQLAALRAMPNVSVIRP | 29 | PYR |
| 484 | 3rim | TK | GLGEDGPTHQPIEHLSALRAIPRLSVVRP | 29 | PYR |
| 485 | 3uk1 | TK | ADTVETAVAWTYAVAHQHPSCLIFSRQNL | 29 | PYR |
| 486 | 3uk1 | TK | ATGSEVELAMKAVEPLAQQGIAARVVSMP | 29 | CFX |
| 487 | 3uk1 | TK | GLGEDGPTHQSVEHVASLRLIPNLDVWRP | 29 | PYR |
| 488 | 4c7v | TK | AVGKDGPTHEPIEQLASLRTIPNVQVFRP | 29 | PYR |
| 489 | 4cok | PDC | GDGSFQLTAQEVAQMIRHDLPVIIFLINN | 29 | PP |
| 490 | 4cok | PDC | GGELAAAIEQARANRNGPTLIECTLDRDD | 29 | PP |
| 491 | 4k9q | BFD | MLLNEPLLTNIEAINMPKPWVKWSYEPAR | 29 | PYR |
| 492 | 4qq8 | OxC | ATEHIPRLVMQAIRAALSAPRGPVLLDLP | 29 | PYR |
| 493 | 4rji | ALS | DVKNIPEAVTNAFRIASAGQAGAAFVSFP | 29 | PYR |
| 494 | 4rji | ALS | GDGGFLFSAMELETAVRLKAPIVHIVWND | 29 | PP |
| 495 | 4rji | ALS | GIPGAKIDAVFDALQDKGPEIIVARHEQN | 29 | PYR |
| 496 | 4rji | ALS | GMKGGRPEAIKAVRKLLKKVQLPFVETYQ | 29 | CFX |
| 497 | 4xeu | TK | ADAVESAVAWKHAIERADGPSALIFSRQN | 29 | PYR |
| 498 | 4xeu | TK | GHDADEIKTAIDTARKSDQPTLICCKTVI | 29 | PP |
| 499 | 4xeu | TK | GLGEDGPTHQPIEQLASLRLTPNLDTWRP | 29 | PYR |
| 500 | 5ahk | AHA | GDGGAQLNIQELDIIARDKLPILTIVMNN | 29 | PP |
| 501 | 5ahk | AHA | GPGATNLITGIADCWLDSHPCIFLTGQVN | 29 | PYR |
| 502 | 5ahk | AHA | SADELVPCLRKAIQIAKEGRPGPVLLDIP | 29 | PYR |
| 503 | 5c4i | PFO | GPTGCMYVANTSYGCGPWRVPWIHAQITN | 29 | PP |
| 504 | 5dx6 | ALS | APDALAEVVSNAFRAAEQGRPGSAFVSLP | 29 | PYR |
| 505 | 5dx6 | ALS | GLMASQPENSKALRRLLETSHIPVTSTYQ | 29 | CFX |
| 506 | 5euj | PDC | AEEAPAKIDHVIRTALRERKPAYLEIACN | 29 | PYR |
| 507 | 5euj | PDC | GAELEGAIKKALDNRRGPTLIECNIAQDD | 29 | PP |
| 508 | 5euj | PDC | GDGSFQLTAQEVAQMIRYEIPVIIFLINN | 29 | PP |
| 509 | 5hje | TK | AMGAIMNGIAAFGANYKNYGGTFLNFVSY | 29 | PYR |
| 510 | 5hje | TK | ATGSEVSLAVDALKVLEGQGIKAGVVSLP | 29 | CFX |
| 511 | 5hje | TK | GLGEDGPTHQPIETLAHFRATPNISVWRP | 29 | PYR |
| 512 | 5nd6 | TK | GLGEDGPTHQPIEHLASFRAMPDMLMIRP | 29 | PYR |
| 513 | 5npu | PDC | GDGSFQLTAQEVAQMVRRKLPIIIFLINN | 29 | PP |
| 514 | 5npu | PDC | GGELAEAIKKALAHREGPTLIECVIDRDD | 29 | PP |
| 515 | 5tma | PDC | ASLNAAVEETLKFIENRDKVAVLVGSKLR | 29 | CFX |
| 516 | 5tma | PDC | GDGSFQLTAQEVAQMVRLKLPVIIFLINN | 29 | PP |
| 517 | 5vrb | TK | AIEAARAETGKPSIICCKTLIGKGSANKE | 29 | PP |
| 518 | 5vrb | TK | GLGEDGPTHQPIEQTATLRLIPNMDVWRP | 29 | PYR |
| 519 | 6a50 | BFD | GDGSANYSISALWTAAQYNIPTIFVIMNN | 29 | PP |
| 520 | 6cin | PFO | GSLGEPLYEDVQTVLAEHGKNILVVGGRY | 29 | CFX |
| 521 | 6dek | AHA | GDASFNMTLTELSSAVQAGAPIKVCVLNN | 29 | PP |
| 522 | 6gua | PhK | DHNGYTHQDPGMLTHLAEKKSDFIRQYLP | 29 | PYR |
| 523 | 1ay0 | TK | ATGSEVSLSVEAAKTLAAKNIKARVVSLPD | 30 | CFX |
| 524 | 1b0p | PFO | GAPGDPLYLDVCSAFVERGEAMPKILAGRY | 30 | CFX |
| 525 | 1b0p | PFO | GLGSKEFSPAMVKSVYDNMSGAKKNHFTVG | 30 | CFX |
| 526 | 1b0p | PFO | PENCIQCNQCAFVCPHSAILPVLAKEEELV | 30 | CFX |
| 527 | 1b0p | PFO | SSSVQEAHDMALVAHLAAIESNVPFMHFFD | 30 | PYR |
| 528 | 1dtw | PDH | GGFASEISSTVQEECFLNLEAPISRVCGYD | 30 | CFX |
| 529 | 1itz | TK | GTGSELEIAAKAADELRKEGKTVRVVSFVS | 30 | CFX |
| 530 | 1n0h | AHA | GAGILNHADGPRLLKELSDRAQIPVTTTLQ | 30 | CFX |
| 531 | 1n0h | AHA | TSRAQDEFVMQSINKAADLINLAKKPVLYV | 30 | CFX |
| 532 | 1ni4 | PDH | AQYDGAYKVSRGLWKKYGDKRIIDTPISEM | 30 | PYR |
| 533 | 1ni4 | PDH | GIVGAQVPLGAGIALACKYNGKDEVCLTLY | 30 | PP |
| 534 | 1ni4 | PDH | GMDILCVREATRFAAAYCRSGKGPILMELQ | 30 | PP |
| 535 | 1ovm | PDC | GDGAAQLTIQELGSMLRDKQHPIILVLNNE | 30 | PP |

| | | | | | |
|---|---|---|---|---|---|
| 536 | 1ovm | PDC | GVPGDYNLQFLDHVIDSPDICWVGCANELN | 30 | PYR |
| 537 | 1qpb | PDC | ADACCSRHDVKAETKKLIDLTQFPAFVTPM | 30 | CFX |
| 538 | 1qpb | PDC | GDGSLQLTVQEISTMIRWGLKPYLFVLNND | 30 | PP |
| 539 | 1qpb | PDC | GLPGDFNLSLLDKIYEVEGMRWAGNANELN | 30 | PYR |
| 540 | 1qpb | PDC | GSIGFTTGATLGAAFAAEEIDPKKRVILFI | 30 | PP |
| 541 | 1qpb | PDC | TGEWDKLTQDKSFNDNSKIRMIEVMLPVFD | 30 | PP |
| 542 | 1umb | PDH | ASFVSEVAATIAEDLLDMLLAPPIRVTGFD | 30 | CFX |
| 543 | 1umb | PDH | GKRGGVFLVTEGLLQKYGPDRVMDTPLSEA | 30 | PYR |
| 544 | 1y9d | POX | AVNAATLPHVIDEAIRRAYAHQGVAVVQIP | 30 | PYR |
| 545 | 2bp7 | PDH | GNLATQFVQAVGWAMASAIKGDTKIASAWI | 30 | PP |
| 546 | 2bp7 | PDH | GYFGGVFRCTEGLQTKYGKSRVFDAPISES | 30 | PYR |
| 547 | 2j9f | PDH | DVFAVYNATKEARRRAVAENQPFLIEAMTY | 30 | PP |
| 548 | 2j9f | PDH | GGFASEISSTVQEECFLNLEAPISRVCGYD | 30 | CFX |
| 549 | 2jlc | BFD | GDLSALYDLNALALLRQVSAPLVLIVVNNN | 30 | PP |
| 550 | 2pgn | ALS | GDGALYYHFNEFRVAVEHKLPVITMVFTNE | 30 | PP |
| 551 | 2pgn | ALS | GFIGHTSHFVADAFSKSHLGKRVINPATEL | 30 | PYR |
| 552 | 2r5n | TK | ANPAKIASRKASQNAIEAFGPLLPEFLGGS | 30 | PYR |
| 553 | 2r5n | TK | ATGSEVELAVAAYEKLTAEGVKARVVSMPS | 30 | CFX |
| 554 | 2r5n | TK | GHDAASIKRAVEEARAVTDKPSLLMCKTII | 30 | PP |
| 555 | 2vbf | PDC | GDGSLQLTVQELGLSIREKLNPICFIINND | 30 | PP |
| 556 | 2vbf | PDC | GVPGDYNLQFLDQIISREDMKWIGNANELN | 30 | PYR |
| 557 | 2vbi | PDC | AVAGDYNLVLLDQLLLNKDMKQIYCCNELN | 30 | PYR |
| 558 | 2vjy | PDC | ADACCSRHDAKAETKKLIDLTQFPAFVTPM | 30 | CFX |
| 559 | 2vjy | PDC | GDGSLQLTVQEISTMIRWGLKPYLFVLNND | 30 | PP |
| 560 | 2vjy | PDC | GLPGDFNLSLLDNIYEVPGMRWAGNANELN | 30 | PYR |
| 561 | 2vjy | PDC | GSIGFTTGATLGAAFAAEEIDPKKRVILFI | 30 | PP |
| 562 | 2vjy | PDC | TGEWNKLTTDEKFQDNTRIRLIEVMLPTMD | 30 | PP |
| 563 | 2x7j | BFD | GDLSFYHDLNGLLAAKKLGIPLTVILVNND | 30 | PP |
| 564 | 3duf | PDH | GVNGGVFRATEGLQAEFGEDRVFDTPLAES | 30 | PYR |
| 565 | 3exe | PDH | AQYDGAYKVSRGLWKKYGDKRIIDTPISEM | 30 | PYR |
| 566 | 3exe | PDH | GIVGAQVPLGAGIALACKYNGKDEVCLTLY | 30 | PP |
| 567 | 3exe | PDH | GMDILCVREATRFAAAYCRSGKGPILMELQ | 30 | PP |
| 568 | 3lq1 | BFD | GDLSFYHDMNGLLMAKKYKMNLTIVIVNND | 30 | PP |
| 569 | 3m34 | TK | ADLGPSNKTELHSMGDFVEGKNIHFGIREH | 30 | PYR |
| 570 | 3m34 | TK | ASGSEVWLCLESANELEKQGFACNVVSMPC | 30 | CFX |
| 571 | 3m34 | TK | FIFSEYLKPAARIAALMKIKHFFIFTHDSI | 30 | PYR |
| 572 | 3m49 | TK | ATGSEVSLAVEAQKALAVDGVDASVVSMPS | 30 | CFX |
| 573 | 3m49 | TK | FVFSDYLRPAIRLAALMQLPVTYVFTHDSI | 30 | PYR |
| 574 | 3rim | TK | ADANETAYAWRTILARRNGSGPVGLILTRQ | 30 | PYR |
| 575 | 3rim | TK | ATGSEVQLAVAAQTLLADNDILARVVSMPC | 30 | CFX |
| 576 | 3rim | TK | ISIEDDTNIALCEDTAARYRAYGWHVQEVE | 30 | PP |
| 577 | 3rim | TK | LQFSDYMRPAVRLAALMDIDTIYVWTHDSI | 30 | PYR |
| 578 | 3uk1 | TK | LTFSDYSRNALRVAALMKVPSIFVFTHDSI | 30 | PYR |
| 579 | 4c7v | TK | FVFSDYLKAAIRLSAIQKLPVIYVLTHDSV | 30 | PYR |
| 580 | 4c7v | TK | GFNLEEIDKAIVQAKAESDKPTIIEIKTTI | 30 | PP |
| 581 | 4cok | PDC | EASLKAAVDAALAFIEQRGSVTMLVGSRIR | 30 | CFX |
| 582 | 4qq8 | OxC | GDGSVGYSIGEFDTLVRKQLPLIVIIMNNQ | 30 | PP |
| 583 | 4xeu | TK | ATGSEVGLAVQAYDKLSEQGRKVRVVSMPC | 30 | CFX |
| 584 | 5dx6 | ALS | GDGGFLQSSMELETAVRLKANVLHLIWVDN | 30 | PP |
| 585 | 5erx | BFD | APRSGDNPLHPLALPLLRPQQVIMLGRPTL | 30 | CFX |
| 586 | 5erx | BFD | GSRNAPLAFALQDADRSGRIRLHVRIDERT | 30 | PYR |
| 587 | 5nd6 | TK | GTGSELELATAAAGILEKEGKNVRVVSFPC | 30 | CFX |
| 588 | 5nd6 | TK | YIFTDYMRNAMRMSALSEAGVVYVMTHDSI | 30 | PYR |
| 589 | 5npu | PDC | EETLKAAVEAALDFIEKREKPVLLVGGKLR | 30 | CFX |

| | | | | | |
|---|---|---|---|---|---|
| 590 | 5vrb | TK | ATGSEVGLAVEAQKVLAGQGIAVRVVSMPS | 30 | CFX |
| 591 | 5vrb | TK | LMFSEYERNALRMAALMKINPVFVFTHDSI | 30 | PYR |
| 592 | 6a50 | BFD | ASAAEVPHAMSRAIHMASMAPQGPVYLSVP | 30 | PYR |
| 593 | 6cin | PFO | GLGSKEFNPSMVKAVFDNLAATTPKNKFTV | 30 | CFX |
| 594 | 6cin | PFO | SSCEVIEETVNYLVEKGEKVGLIKVRLFRP | 30 | CFX |
| 595 | 6dek | AHA | GAGILNNEQGPKLLKELADKANIPVTTTLQ | 30 | CFX |
| 596 | 6gua | PhK | GHWGTVSGQTFLYAHANRLINKYDQKMFYM | 30 | PP |
| 597 | 1b0p | PFO | GADGTVGANKQAIKIIGDNTDLFAQGYFSYD | 31 | CFX |
| 598 | 1b0p | PFO | GAPANFTALEAKGKELKGYKFRIQINTLDCM | 31 | CFX |
| 599 | 1b0p | PFO | SMGYSKQQFLKVLKEAESFPGPSLVIAYATC | 31 | PP |
| 600 | 1dtw | PDH | GEGAASEGDAHAGFNFAATLECPIIFFCRNN | 31 | PP |
| 601 | 1dtw | PDH | GNDVFAVYNATKEARRRAVAENQPFLIEAMT | 31 | PP |
| 602 | 1itz | TK | GNTGYDDIRAAIKEAKAVTDKPTLIKVTTTI | 31 | PP |
| 603 | 1n0h | AHA | GPGATNVVTPMADAFADGIPMVVFTGQVPTS | 31 | PYR |
| 604 | 1r9j | TK | GDTDYEGLRKALAEAKATKGKPKMIVQTTTI | 31 | PP |
| 605 | 1umb | PDH | GDGATSEGDWYAGINFAAVQGAPAVFIAENN | 31 | PP |
| 606 | 1y9d | POX | GQFGTTGMNMDTFQEMNENPIYADVADYNVT | 31 | PYR |
| 607 | 2bp7 | PDH | DFVAVYAASRWAAERARRGLGPSLIEWVTYR | 31 | PP |
| 608 | 2bp7 | PDH | SNPYDAKGLLIASIECDDPVIFLEPKRLYNG | 31 | PYR |
| 609 | 2j9f | PDH | GEGAASEGDAHAGFNFAATLECPIIFFCRNN | 31 | PP |
| 610 | 3ahc | PhK | GHWGTTPGLNFLLAHINRLIADHQQNTVFIM | 31 | PP |
| 611 | 3ai7 | PhK | GHWGTTPGLNFLIGHINRFIADHGQNTVIIM | 31 | PP |
| 612 | 3duf | PDH | GDGGTSQGDFYEGINFAGAFKAPAIFVVQNN | 31 | PP |
| 613 | 3duf | PDH | GMDPLAVYAAVKAARERAINGEGPTLIETLC | 31 | PP |
| 614 | 3rim | TK | GGENVVGIEEAIANAQAVTDRPSFIALRTVI | 31 | PP |
| 615 | 3uk1 | TK | AGANERGETVATRKASQQTIEGLAAVLPELL | 31 | PYR |
| 616 | 4c7v | TK | ATGSEVGLALKAKEELQKKGKDVIVVSLPSW | 31 | CFX |
| 617 | 4cok | PDC | AVAGDYNLVLLDQLLLNTDMQQIYCSNELNC | 31 | PYR |
| 618 | 4k9q | BFD | GAGLGNAMGCLLTAYQNKTPLIITAGQQTRE | 31 | PYR |
| 619 | 4k9q | BFD | GDGSFQYSVQGIYTGVQQKTHVIYVVFQNEE | 31 | PP |
| 620 | 4kxu | TK | GSLGQGLGAACGMAYTGKYFDKASYRVYCLL | 31 | PP |
| 621 | 4kxu | TK | GVSIGEDGPSQMALEDLAMFRSVPTSTVFYP | 31 | PYR |
| 622 | 4kxu | TK | LGQSDPAPLQHQMDIYQKRCEAFGWHAIIVD | 31 | PP |
| 623 | 4qq8 | OxC | ADGGLTYLWLSEVMSRVKPGGFLCHGYLNSM | 31 | PP |
| 624 | 5c4i | PFO | ATGIVDVENLAAVVKNPAAMRRGYAEAQVRQ | 31 | CFX |
| 625 | 5c4i | PFO | GDGGAVDIGLQALSAMLYRGHDVLFICYDNE | 31 | PP |
| 626 | 5euj | PDC | AVAGDYNLVLLDQLLLNKDMEQVYCCNELNC | 31 | PYR |
| 627 | 5hje | TK | ADGNETSAAYKSAIESTHTPHILALTRQNLP | 31 | PYR |
| 628 | 5nd6 | TK | AGGNETAGAYKVAIANRKRPTTIALSRQNMP | 31 | PYR |
| 629 | 5npu | PDC | AVAGDYNLVLLDQLLKNKDLEQVYCCNELNC | 31 | PYR |
| 630 | 5tma | PDC | AVAGDYNLVLLDNLLDNKNMEQVYCCNELNC | 31 | PYR |
| 631 | 6cin | PFO | AMGASHSQLMKALIEAEKYDGPSLIIAYAPC | 31 | PP |
| 632 | 1ay0 | TK | GDGCLQEGISSEASSLAGHLKLGNLIAIYDDN | 32 | PP |
| 633 | 1dtw | PDH | GYAISTPTSEQYRGDGIAARGPGYGIMSIRVD | 32 | PP |
| 634 | 1itz | TK | GDGCQMEGIANEACSLAGHWGLGKLIAFYDDN | 32 | PP |
| 635 | 1itz | TK | HISIDGDTEIAFTEDVSTRFEALGWHTIWVKN | 32 | PP |
| 636 | 1r9j | TK | GDGCLMEGVCQEALSLAGHLALEKLIVIYDSN | 32 | PP |
| 637 | 1r9j | TK | YISIDGSTSLSFTEQCHQKYVAMGFHVIEVKN | 32 | PP |
| 638 | 1umb | PDH | FYAISVDYRHQTHSPTIADKAHAFGIPGYLVD | 32 | PP |
| 639 | 1upa | ALS | GDGGFHSNSSDLETIARLNLPIVTVVVNNDTN | 32 | PP |
| 640 | 2bp7 | PDH | GDGATAESDFHTALTFAHVYRAPVILNVVNNQ | 32 | PP |
| 641 | 2c31 | OxC | REIVDLQQGDYEEMDQMNVARPHCKASFRINS | 32 | PYR |
| 642 | 2dji | POX | GSRPQRELNMDAFQELNQNPMYDHIAVYNRRV | 32 | PYR |
| 643 | 2pan | AHA | APRARLHKEDFQAVDIEAIAKPVSKMAVTVRE | 32 | PYR |

| | | | | | |
|---|---|---|---|---|---|
| 644 | 2r5n | TK | GDGCMMEGISHEVCSLAGTLKLGKLIAFYDDN | 32 | PP |
| 645 | 2r5n | TK | GISIDGHVEGWFTDDTAMRFEAYGWHVIRDID | 32 | PP |
| 646 | 2x7j | BFD | SPQMLRYIRTLASRAAGEAQKRPMGPVHVNVP | 32 | PYR |
| 647 | 3ahc | PhK | ASAGDVPTQELMAASDALNKMGIKFKVVNVVD | 32 | CFX |
| 648 | 3ai7 | PhK | AAAGDVPTQEIMAASDKLKELGVKFKVVNVAD | 32 | CFX |
| 649 | 3duf | PDH | RFAISTPVEKQTVAKTLAQKAVAAGIPGIQVD | 32 | PP |
| 650 | 3lq1 | BFD | SEEMLRYAKWHGSRAVDIAMKTPRGPVHLNFP | 32 | PYR |
| 651 | 3uk1 | TK | GDGCLMEGISHEACSLAGTLKLNKLIALYDDN | 32 | PP |
| 652 | 4c7v | TK | GDGDLMEGVASEAASLAGHLKLGKLIALYDSN | 32 | PP |
| 653 | 4c7v | TK | GISLDGKTSASFTENVGARFEAYGWQYILVED | 32 | PP |
| 654 | 4k9q | BFD | ASGGLGWDLPAAVGLALGEEVSGRNRPVVTLM | 32 | PP |
| 655 | 4kxu | TK | PFTIKPLDRKLILDSARATKGRILTVEDHYYE | 32 | CFX |
| 656 | 4xeu | TK | GDGCMMEGISHEVASLAGTLRLNKLIAFYDDN | 32 | PP |
| 657 | 4xeu | TK | GISIDGEVHGWFTDDTPKRFEAYGWQVIRNVD | 32 | PP |
| 658 | 5c4i | PFO | GGAVASGIEAAYKAMIRKKKTDAEFPNIIVMA | 32 | PP |
| 659 | 5erx | BFD | GDLTFVHDSSGLLIGPTEPIPRSLTIVVSNDN | 32 | PP |
| 660 | 5hje | TK | ADAAVATRKLSEIVLSKIIPEVPEIIGGSADL | 32 | PYR |
| 661 | 5hje | TK | GDGCLMEGVSSEASSLAGHLQLGNLIAFWDDN | 32 | PP |
| 662 | 5nd6 | TK | GDGCMMEGISNEACSLAGHWGLGKLIALYDDN | 32 | PP |
| 663 | 5vrb | TK | GDGCLMEGVSHEACSLAGTLGLGKLIVLYDDN | 32 | PP |

**Supplemental Table S3:** Distribution of ThDP repeats by domain and enzymatic group

| | AHAS | ALS | BFD | OxCDC | PDC | PDH | PFOR | PhK | POX | TK | All |
|---|---|---|---|---|---|---|---|---|---|---|---|
| **# of standard size repeats (%)** | 66 (74.2) | 48 (66.7) | 68 (65.4) | 34 (63.0) | 124 (73.4) | 79 (70.5) | 49 (66.2) | 17 (33.3) | 36 (66.7) | 142 (63.4) | 663 (100) |
| **% standard size repeats from PP domain** | 40.9 | 41.7 | 35.7 | 41.2 | 44.4 | 41.8 | 30.6 | 35.3 | 41.7 | 21.8 | 37.5 |
| **% standard size repeats from PYR domain** | 39.4 | 35.4 | 37.1 | 38.2 | 29.0 | 36.7 | 22.4 | 17.6 | 33.3 | 46.5 | 33.6 |
| **% standard size repeats from CFX domain** | 19.7 | 22.9 | 21.4 | 20.6 | 26.6 | 21.5 | 46.9 | 47.1 | 25.0 | 31.7 | 28.3 |
| **# of standard size PP repeats** | 27 | 20 | 25 | 14 | 55 | 33 | 10 | 6 | 15 | 31 | 236 |
| **# of standard size PYR repeats** | 26 | 17 | 28 | 13 | 36 | 29 | 11 | 3 | 12 | 66 | 241 |
| **# of standard size CFX repeats** | 13 | 11 | 15 | 7 | 33 | 17 | 28 | 8 | 9 | 45 | 186 |

**Supplemental Table S4:** Statistical analysis of amino acid sequences in the ThDP repeats organized by repeat length. The average (mean) length of the repeats in that size range (LENGTH), the fraction which start with the [G/A]{X(1,2)}[G/A] motif (GXG), the average start position of the helical region as defined by the PDB structure (HLX START), the average end position of the helical region (HLX END), the average start position of the strand (STRN STRT) and end position of the strand (STRN END), the fraction of the repeats which have a proline or glycine residue in the one or two residues after the end of the helix (HELIX PRO), the fraction of repeats which have a proline or glycine residue at the position immediately preceding the start of the strand (STRAND PRO), and finally the fraction of repeats which have both of the aforementioned proline/glycine residues. Statistics are calculated for repeats from all the domains and from just the functional PP & PYR domains further subdivided by the length range of the repeats. The statistics off all the repeats are highlighted in grey while the statistics calculated from just those repeats 22-32 residues in length (see Fig. 4) are highlighted in green.

| | LENGTH | GXG | HLX START | HLX END | STRN STRT | STRN END | HELIX PRO | STRAND PRO | BOTH PRO |
|---|---|---|---|---|---|---|---|---|---|
| **All Domains** | | | | | | | | | |
| **TOTAL =** | 26.98 | 40 | 3.44 | 14.95 | 20.27 | 24.6 | 36.6 | 19.5 | 5.4 |
| **20-32 aa** | 26.05 | 53.2 | 3.44 | 14.72 | 19.94 | 24.02 | 37.5 | 23.8 | 6.9 |
| **22-32 aa** | 26.79 | 60.8 | 3.5 | 15.24 | 20.51 | 24.63 | 37.4 | 26.5 | 7.8 |
| **20-30 aa** | 25.52 | 58.7 | 3.32 | 14.29 | 19.52 | 23.49 | 37.5 | 23 | 6.4 |
| **22-30 aa** | 26.26 | 67.8 | 3.37 | 14.81 | 20.09 | 24.09 | 37.4 | 25.8 | 7.4 |
| **PP & PYR domains** | | | | | | | | | |
| **TOTAL =** | 27.13 | 39.1 | 3.26 | 14.82 | 20.23 | 24.66 | 34.5 | 20.2 | 5.2 |
| **20-32 aa** | 25.97 | 56.2 | 3.18 | 14.46 | 19.79 | 23.93 | 34.8 | 26.1 | 7.1 |
| **22-32 aa** | 26.98 | 67.1 | 3.22 | 15.11 | 20.53 | 24.73 | 34.2 | 30.2 | 8.5 |
| **20-30 aa** | 25.52 | 62.3 | 3.18 | 14.17 | 19.48 | 23.58 | 35.4 | 26.2 | 7.9 |
| **22-30 aa** | 26.55 | 75.6 | 3.22 | 14.85 | 20.25 | 24.41 | 34.9 | 30.9 | 9.5 |

**Supplemental Table S5:** Distribution of sequence matches (% of total sequences found) in the TEED database to the sequences in the PDB. Data is arranged by enzyme class, sequences detected within the listed pairwise identity cutoff and by repeat number and domain (domains are ordered N to C regardless of enzyme class).

| Repeat | # : | **1A** | **1B** | **1C** | **2A** | **2B** | **2C** | **3A** | **3B** | **3C** | **4A** | **4B** | **4C** | **5A** | **5B** | **5C** | **6A** | **6B** | **6C** |
|---|---|---|---|---|---|---|---|---|---|---|---|---|---|---|---|---|---|---|---|
| 80 % ID | AHA | 9.0 | 9.1 | 9.1 | 9.1 | 9.0 | 9.1 | 9.1 | 9.0 | 9.1 | 9.1 | 9.0 | 8.8 | 9.1 | 9.3 | 18.1 | 9.1 | 10.8 | 19.3 |
| | ALS | 5.0 | 4.8 | 4.8 | 5.0 | 4.7 | 4.8 | 5.0 | 4.7 | 4.8 | 5.0 | 4.7 | 4.8 | 5.0 | 4.9 | 10.0 | 5.0 | 5.7 | 10.7 |
| | BFD | 13.6 | 12.9 | 12.9 | 13.5 | 12.9 | 12.9 | 13.5 | 12.8 | 12.9 | 13.5 | 12.8 | 12.9 | 13.5 | 11.9 | 26.7 | 13.5 | 2.7 | 28.6 |
| | OxCDC | 3.0 | 2.8 | 2.8 | 3.0 | 2.8 | 2.8 | 2.9 | 2.8 | 2.8 | 2.9 | 2.8 | 2.8 | 2.9 | 2.8 | 5.8 | 2.9 | 3.3 | 6.2 |
| | PDC | 6.0 | 5.8 | 5.8 | 6.0 | 5.8 | 5.8 | 6.1 | 5.8 | 5.8 | 6.1 | 5.8 | 5.9 | 6.1 | 4.2 | 11.8 | 6.1 | 0.0 | 12.7 |
| | PDH | 12.5 | 15.9 | 16.1 | 12.5 | 16.0 | 16.1 | 12.4 | 16.1 | 16.1 | 12.4 | 16.1 | 16.1 | 12.4 | 16.6 | 0.0 | 12.4 | 19.3 | 0.0 |
| | PHK | 3.2 | 3.0 | 3.1 | 3.2 | 3.0 | 3.1 | 3.2 | 3.0 | 3.1 | 3.2 | 3.0 | 3.1 | 3.2 | 3.1 | 6.4 | 3.2 | 3.7 | 0.0 |
| | POX | 10.5 | 10.0 | 10.0 | 10.5 | 10.0 | 10.0 | 10.5 | 10.0 | 10.0 | 10.5 | 10.0 | 10.1 | 10.5 | 10.3 | 21.0 | 10.5 | 12.0 | 22.4 |
| | TK | 37.2 | 35.7 | 35.4 | 37.3 | 35.6 | 35.4 | 37.2 | 35.6 | 35.3 | 37.2 | 35.6 | 35.5 | 37.2 | 36.7 | 0.0 | 37.2 | 42.5 | 0.0 |
| | PFOR | 0.1 | 0.1 | 0.1 | 0.1 | 0.1 | 0.1 | 0.1 | 0.1 | 0.1 | 0.1 | 0.1 | 0.1 | 0.1 | 0.1 | 0.2 | 0.1 | 0.0 | 0.0 |
| 70 % ID | AHA | 12.7 | 12.4 | 12.5 | 12.8 | 12.4 | 12.5 | 12.7 | 12.4 | 12.5 | 12.7 | 12.4 | 12.4 | 12.8 | 12.7 | 34.0 | 12.8 | 13.8 | 36.0 |
| | ALS | 3.0 | 2.8 | 2.8 | 3.0 | 2.8 | 2.8 | 3.0 | 2.8 | 2.8 | 3.0 | 2.8 | 2.9 | 3.0 | 2.9 | 7.9 | 3.0 | 3.2 | 8.4 |
| | BFD | 7.7 | 7.4 | 7.4 | 7.7 | 7.3 | 7.4 | 7.7 | 7.3 | 7.4 | 7.7 | 7.4 | 7.4 | 7.7 | 6.4 | 20.3 | 7.7 | 1.5 | 21.5 |
| | OxCDC | 1.5 | 1.4 | 1.4 | 1.5 | 1.4 | 1.4 | 1.5 | 1.4 | 1.4 | 1.5 | 1.4 | 1.4 | 1.5 | 1.4 | 4.0 | 1.5 | 1.6 | 4.2 |
| | PDC | 4.2 | 4.0 | 4.0 | 4.1 | 4.0 | 4.0 | 4.2 | 4.0 | 4.0 | 4.2 | 4.0 | 4.1 | 4.2 | 3.1 | 10.9 | 4.2 | 0.0 | 11.5 |
| | PDH | 9.0 | 12.5 | 12.7 | 9.0 | 12.6 | 12.8 | 9.0 | 12.6 | 12.8 | 8.9 | 12.7 | 12.8 | 8.9 | 13.0 | 0.0 | 8.9 | 14.2 | 0.0 |
| | PHK | 2.1 | 2.0 | 2.0 | 2.1 | 2.0 | 2.0 | 2.1 | 2.0 | 2.0 | 2.1 | 2.0 | 2.0 | 2.1 | 2.1 | 5.6 | 2.1 | 2.3 | 0.0 |
| | POX | 6.5 | 6.2 | 6.3 | 6.5 | 6.2 | 6.3 | 6.5 | 6.2 | 6.3 | 6.5 | 6.2 | 6.3 | 6.5 | 6.3 | 17.3 | 6.5 | 6.9 | 18.4 |
| | TK | 53.2 | 51.2 | 50.7 | 53.2 | 51.2 | 50.7 | 53.3 | 51.1 | 50.6 | 53.3 | 51.1 | 50.7 | 53.3 | 52.1 | 0.0 | 53.3 | 56.5 | 0.0 |
| | PFOR | 0.1 | 0.1 | 0.1 | 0.1 | 0.1 | 0.1 | 0.1 | 0.1 | 0.1 | 0.1 | 0.1 | 0.1 | 0.1 | 0.1 | 0.2 | 0.1 | 0.0 | 0.0 |
| 62 % ID | AHA | 9.3 | 8.9 | 9.0 | 9.3 | 8.9 | 9.0 | 9.3 | 8.9 | 9.0 | 9.3 | 8.9 | 8.9 | 9.3 | 9.1 | 30.8 | 9.3 | 10.0 | 32.5 |
| | ALS | 2.4 | 2.3 | 2.3 | 2.4 | 2.3 | 2.3 | 2.4 | 2.3 | 2.3 | 2.4 | 2.3 | 2.3 | 2.4 | 2.3 | 7.9 | 2.4 | 2.5 | 8.3 |
| | BFD | 5.9 | 5.6 | 5.7 | 5.9 | 5.6 | 5.7 | 5.9 | 5.6 | 5.7 | 5.9 | 5.6 | 5.7 | 5.9 | 4.8 | 19.5 | 5.9 | 0.9 | 20.6 |
| | OxCDC | 1.0 | 1.0 | 1.0 | 1.0 | 1.0 | 1.0 | 1.0 | 1.0 | 1.0 | 1.0 | 1.0 | 1.0 | 1.0 | 1.0 | 3.4 | 1.0 | 1.1 | 3.5 |
| | PDC | 4.2 | 4.0 | 4.0 | 4.2 | 4.0 | 4.0 | 4.2 | 4.0 | 4.0 | 4.2 | 4.0 | 4.0 | 4.2 | 3.4 | 13.8 | 4.2 | 0.0 | 14.5 |
| | PDH | 5.5 | 9.7 | 9.8 | 5.4 | 9.7 | 9.8 | 5.4 | 9.7 | 9.8 | 5.4 | 9.7 | 9.8 | 5.4 | 9.9 | 0.0 | 5.4 | 10.9 | 0.0 |
| | PHK | 1.3 | 1.2 | 1.2 | 1.3 | 1.2 | 1.2 | 1.3 | 1.2 | 1.2 | 1.3 | 1.2 | 1.2 | 1.3 | 1.2 | 4.2 | 1.3 | 1.4 | 0.0 |
| | POX | 5.9 | 5.6 | 5.6 | 5.9 | 5.6 | 5.6 | 5.9 | 5.6 | 5.6 | 5.9 | 5.6 | 5.6 | 5.9 | 5.7 | 19.5 | 5.9 | 6.3 | 20.6 |
| | TK | 62.9 | 60.1 | 59.8 | 62.9 | 60.1 | 59.8 | 62.9 | 60.1 | 59.7 | 62.9 | 60.0 | 59.8 | 62.9 | 61.0 | 0.0 | 62.9 | 66.9 | 0.0 |
| | PFOR | 1.7 | 1.6 | 1.6 | 1.7 | 1.6 | 1.6 | 1.7 | 1.6 | 1.6 | 1.7 | 1.6 | 1.6 | 1.7 | 1.6 | 1.0 | 1.7 | 0.0 | 0.0 |
| 50 % ID | AHA | 5.0 | 4.7 | 4.7 | 5.0 | 4.7 | 4.7 | 5.0 | 4.7 | 4.7 | 5.0 | 4.7 | 4.7 | 5.0 | 4.7 | 20.5 | 5.0 | 5.4 | 26.8 |
| | ALS | 2.5 | 2.3 | 2.3 | 2.5 | 2.3 | 2.3 | 2.4 | 2.3 | 2.3 | 2.4 | 2.3 | 2.3 | 2.4 | 2.3 | 10.0 | 2.4 | 2.6 | 13.1 |
| | BFD | 3.1 | 2.8 | 2.8 | 3.1 | 2.8 | 2.8 | 3.1 | 2.8 | 2.8 | 3.1 | 2.8 | 2.8 | 3.1 | 2.4 | 12.4 | 3.1 | 0.8 | 16.2 |
| | OxCDC | 1.9 | 1.8 | 1.8 | 1.9 | 1.8 | 1.8 | 1.9 | 1.8 | 1.8 | 1.9 | 1.8 | 1.8 | 1.9 | 1.8 | 7.8 | 1.9 | 2.1 | 10.1 |
| | PDC | 2.8 | 2.6 | 2.6 | 2.8 | 2.6 | 2.6 | 2.8 | 2.6 | 2.6 | 2.8 | 2.6 | 2.6 | 2.8 | 2.3 | 11.3 | 2.8 | 0.0 | 14.7 |
| | PDH | 3.7 | 10.8 | 10.9 | 3.7 | 10.8 | 10.9 | 3.7 | 10.8 | 10.9 | 3.7 | 10.8 | 10.9 | 3.7 | 10.9 | 0.0 | 3.7 | 12.7 | 0.0 |
| | PHK | 2.1 | 1.9 | 1.9 | 2.1 | 1.9 | 1.9 | 2.1 | 1.9 | 1.9 | 2.1 | 1.9 | 1.9 | 2.1 | 1.9 | 8.4 | 2.1 | 2.2 | 0.0 |
| | POX | 3.6 | 3.3 | 3.3 | 3.6 | 3.3 | 3.3 | 3.6 | 3.3 | 3.3 | 3.6 | 3.3 | 3.3 | 3.6 | 3.4 | 14.7 | 3.6 | 3.9 | 19.1 |
| | TK | 65.1 | 60.4 | 60.2 | 65.2 | 60.4 | 60.2 | 65.2 | 60.4 | 60.2 | 65.2 | 60.4 | 60.2 | 65.2 | 60.8 | 0.0 | 65.2 | 70.2 | 0.0 |
| | PFOR | 10.2 | 9.4 | 9.5 | 10.2 | 9.4 | 9.5 | 10.2 | 9.4 | 9.5 | 10.2 | 9.4 | 9.5 | 10.2 | 9.5 | 14.9 | 10.2 | 0.0 | 0.0 |
| 40 % ID | AHA | 6.9 | 6.1 | 6.1 | 7.0 | 6.1 | 6.1 | 6.9 | 6.1 | 6.1 | 7.0 | 6.1 | 6.1 | 7.0 | 6.1 | 32.4 | 7.0 | 6.5 | 45.9 |
| | ALS | 1.6 | 1.4 | 1.4 | 1.6 | 1.4 | 1.4 | 1.6 | 1.4 | 1.4 | 1.6 | 1.4 | 1.4 | 1.6 | 1.4 | 7.6 | 1.6 | 1.5 | 10.8 |
| | BFD | 2.1 | 1.9 | 1.9 | 2.1 | 1.9 | 1.9 | 2.1 | 1.9 | 1.9 | 2.1 | 1.9 | 1.9 | 2.1 | 1.7 | 9.9 | 2.1 | 0.9 | 14.0 |
| | OxCDC | 0.7 | 0.6 | 0.6 | 0.7 | 0.6 | 0.6 | 0.7 | 0.6 | 0.6 | 0.7 | 0.6 | 0.6 | 0.7 | 0.6 | 3.1 | 0.7 | 0.6 | 4.3 |
| | PDC | 2.0 | 1.7 | 1.7 | 2.0 | 1.7 | 1.7 | 2.0 | 1.7 | 1.7 | 2.0 | 1.7 | 1.7 | 2.0 | 1.5 | 9.1 | 2.0 | 0.0 | 12.9 |
| | PDH | 2.6 | 15.0 | 15.1 | 2.6 | 15.0 | 15.1 | 2.5 | 15.0 | 15.1 | 2.5 | 15.1 | 15.1 | 2.5 | 15.1 | 0.0 | 2.5 | 16.2 | 0.0 |
| | PHK | 4.1 | 3.6 | 3.6 | 4.1 | 3.6 | 3.6 | 4.1 | 3.6 | 3.6 | 4.1 | 3.6 | 3.6 | 4.1 | 3.6 | 19.2 | 4.1 | 3.8 | 0.0 |
| | POX | 1.8 | 1.6 | 1.6 | 1.8 | 1.6 | 1.6 | 1.8 | 1.6 | 1.6 | 1.8 | 1.6 | 1.6 | 1.8 | 1.6 | 8.4 | 1.8 | 1.7 | 12.0 |
| | TK | 73.8 | 64.4 | 64.2 | 73.8 | 64.4 | 64.2 | 73.8 | 64.4 | 64.2 | 73.8 | 64.3 | 64.2 | 73.8 | 64.6 | 0.0 | 73.8 | 68.8 | 0.0 |
| | PFOR | 4.4 | 3.8 | 3.8 | 4.4 | 3.8 | 3.8 | 4.4 | 3.8 | 3.8 | 4.4 | 3.8 | 3.8 | 4.4 | 3.8 | 10.2 | 4.4 | 0.0 | 0.0 |
| 30 % ID | AHA | 18.4 | 15.7 | 15.8 | 18.4 | 15.7 | 15.8 | 18.4 | 15.7 | 15.8 | 18.3 | 15.7 | 15.8 | 18.3 | 15.9 | 48.8 | 18.3 | 15.6 | 55.4 |
| | ALS | 2.0 | 1.7 | 1.7 | 2.0 | 1.7 | 1.7 | 2.0 | 1.7 | 1.7 | 2.0 | 1.7 | 1.7 | 2.0 | 1.7 | 5.3 | 2.0 | 1.9 | 6.0 |
| | BFD | 3.0 | 2.6 | 2.6 | 3.0 | 2.6 | 2.6 | 3.0 | 2.6 | 2.6 | 3.0 | 2.6 | 2.6 | 3.0 | 2.3 | 7.9 | 3.0 | 1.7 | 9.0 |
| | OxCDC | 0.8 | 0.7 | 0.7 | 0.8 | 0.7 | 0.7 | 0.8 | 0.7 | 0.7 | 0.8 | 0.7 | 0.7 | 0.8 | 0.7 | 2.2 | 0.8 | 0.8 | 2.5 |
| | PDC | 6.6 | 5.6 | 5.7 | 6.6 | 5.6 | 5.7 | 6.6 | 5.6 | 5.7 | 6.6 | 5.6 | 5.7 | 6.6 | 4.9 | 17.5 | 6.6 | 0.0 | 19.8 |
| | PDH | 6.9 | 20.1 | 20.2 | 6.9 | 20.1 | 20.2 | 6.9 | 20.2 | 20.2 | 6.9 | 20.2 | 20.2 | 6.9 | 20.4 | 0.0 | 6.9 | 22.7 | 0.0 |
| | PHK | 3.1 | 2.6 | 2.6 | 3.1 | 2.6 | 2.6 | 3.1 | 2.6 | 2.6 | 3.1 | 2.6 | 2.6 | 3.1 | 2.7 | 8.1 | 3.1 | 3.0 | 0.0 |
| | POX | 2.4 | 2.1 | 2.1 | 2.4 | 2.1 | 2.1 | 2.4 | 2.1 | 2.1 | 2.4 | 2.1 | 2.1 | 2.4 | 2.1 | 6.4 | 2.4 | 2.3 | 7.3 |
| | TK | 53.9 | 46.4 | 46.2 | 54.0 | 46.3 | 46.2 | 54.0 | 46.3 | 46.1 | 54.0 | 46.3 | 46.2 | 54.0 | 46.8 | 0.0 | 54.0 | 51.9 | 0.0 |
| | PFOR | 2.9 | 2.5 | 2.5 | 2.9 | 2.5 | 2.5 | 2.9 | 2.5 | 2.5 | 2.9 | 2.5 | 2.5 | 2.9 | 2.5 | 3.9 | 2.9 | 0.0 | 0.0 |

**Supplemental Table S6:** Table listing the domains of the non-redundant set ThDP enzyme structures based on the three dimensional arrangement of their repeat types (A-E, see SI Fig. S3). Repeat deletions within a domain (due to actual deletion or lack of experimental electron density in the model) are indicated by (Δ). Insertions of extra repeats are indicated by (+1).

| DC | PDB | 1 | 2 | 3 | DH | PDB | 1 | 2 | 3 | PFOR | PDB | 1 | 2 | 3 | 4 | 5 |
|---|---|---|---|---|---|---|---|---|---|---|---|---|---|---|---|---|
| **AHAS** | 1noh | A | B | A | **PDH** | 1dtw | A | A | C | **PFOR** | 1b0p | A | C(+1) | A(Δ) | D | A |
| | 1ybh | A | B | A | | 1ni4 | A | A | C | | 5c4i | A | C(+1) | E | A | |
| | 2pan | A | B | A | | 1umb | A | A | C | | 6cin | A | C(+1) | B(Δ) | D | A |
| | 5ahk | A | B | A | | 2bp7 | A | A | C | | | | | | | |
| | 6dek | A | B | A | | 2j9f | A | A | C | | | | | | | |
| **ALS** | 1upa | A | B | A | | 3duf | A | A | C | | | | | | | |
| | 2pgn | A | B | A | | 3exe | A | A | C | | | | | | | |
| | 4rji | A | B | A | **PhK** | 3ahc | A | A | C (+1) | | | | | | | |
| | 5dx6 | A | B | A | | 3ai7 | A(Δ) | A(Δ) | C (+1) | | | | | | | |
| **BFD** | 2jlc | A | B | A | | 6gua | A | A | C (+1) | | | | | | | |
| | 2x7j | A | B | A | **TK** | 1ay0 | A | A | C | | | | | | | |
| | 3lq1 | A | B | A | | 1itx | A | A | C | | | | | | | |
| | 4k9q | A | B | A | | 1r9j | A | A | C | | | | | | | |
| | 5erx | A | B | A | | 2r5n | A | A | C | | | | | | | |
| | 6a50 | A | B | A | | 3m34 | A | A | C | | | | | | | |
| **OxCDC** | 2c31 | A | B | A | | 3m49 | A | A | C | | | | | | | |
| | 2q27 | A | B | A | | 3rim | A | A | C | | | | | | | |
| | 4qq8 | A | B | A | | 3uk1 | A | A | C | | | | | | | |
| **PDC** | 1ovm | A | B(Δ) | A | | 4x7v | A | A | C | | | | | | | |
| | 1qpb | A | B(Δ) | A | | 4kxu | A | A | C | | | | | | | |
| | 2nxw | A | B(Δ) | A | | 4xeu | A | A | C | | | | | | | |
| | 2vbf | A | B(Δ) | A | | 5hje | A | A | C | | | | | | | |
| | 2vbi | A | B(Δ) | A | | 5nd6 | A | A | C | | | | | | | |
| | 2vjy | A | B(Δ) | A | | 5vrb | A | A | C | | | | | | | |
| | 4cok | A | B(Δ) | A | | | | | | | | | | | | |
| | 5euj | A | B(Δ) | A | | | | | | | | | | | | |
| | 5npu | A | B(Δ) | A | | | | | | | | | | | | |
| | 5tma | A | B(Δ) | A | | | | | | | | | | | | |
| **POX** | 1y9d | A | B | A | | | | | | | | | | | | |
| | 2dji | A | B | A | | | | | | | | | | | | |
| | 3ey9 | A | B | A | | | | | | | | | | | | |

**Supplemental Table S7:** A list of the canonical Rossmann ligand/carboxylate interaction angles in ThDP enzymes indicating the PDB ID and chain, the identity of the ligand, either flavin adenine dinucleotide (FAD) or adenosine diphosphate (ADP), the identity of the interacting carboxylate residue, its secondary structure, the hydrogen bonding distances between the carboxylate and ribose oxygens (in Å), the manually determined ring configuration, and the value of the angle α. It is clear that the values of are widely distributed suggesting that the Rossmann-like geometry (ring configuration = E, $90^\circ < \alpha < 140^\circ$ ) of the B type domains are the result of convergent evolution rather than common ancestry. Calculation of the angle α was as described in Laurino *et al. PLoS Biol*. **14**(3): e1002396.

| PDB | chain | Ligand | residue | 2˚ struct | D1 (Å) | D2 (Å) | ring configuration | α |
|---|---|---|---|---|---|---|---|---|
| 2c31 | A | ADP | D306 | B | 2.7 | 2.5 | E | 134.0 |
| | B | ADP | D306 | B | 2.9 | 2.5 | E | 137.5 |
| 1n0h | A | FAD | E407 | B | 2.6 | 2.5 | E | 32.6 |
| | B | FAD | E407 | B | 2.8 | 2.6 | E | 32.3 |
| 2pgn | A | FAD | D302 | B | 2.6 | 2.7 | T | 55.8 |
| | B | FAD | D302 | B | 2.6 | 2.7 | T | 53.6 |
| 2pan | A | FAD | D302 | B | 2.5 | 2.7 | T | 56.6 |
| | B | FAD | D302 | B | 2.6 | 2.6 | T | 64.0 |
| | C | FAD | D302 | B | 2.6 | 2.7 | T | 63.3 |
| | D | FAD | D302 | B | 2.5 | 2.8 | E | 63.8 |
| | E | FAD | D303 | B | 2.6 | 2.7 | T | 61.9 |
| | F | FAD | D304 | B | 2.7 | 2.7 | T | 61.2 |
| 5ahk | A | FAD | D308 | B | 2.7 | 2.7 | T | 66.3 |
| | B | FAD | D308 | B | 2.7 | 2.6 | T | 64.3 |
| 1ybh | A | FAD | D395 | B | 3.1 | 2.7 | E | 61.5 |
| 6dek | A | FAD | E403 | B | 3 | 2.2 | E | 55.7 |
| 2q28 | A | ADP | D302 | B | 2.7 | 2.6 | E | 132.5 |
| | B | ADP | D302 | B | 2.7 | 2.6 | E | 131.8 |

**Supplemental Table S8:** Sequence propensity for each amino acid as calculated from the PDB (Nov. 2018) according to the overall abundance of each amino acid and their presence within specific secondary structure elements based on a three state (helix, strand, coil) secondary structure definition.

| Amino Acid | General | Helix | Strand | Coil |
|---|---|---|---|---|
| **A** | 7.996 | 11.027 | 6.168 | 6.536 |
| **C** | 1.358 | 1.159 | 1.987 | 1.224 |
| **D** | 5.624 | 4.908 | 3.105 | 7.307 |
| **E** | 6.608 | 8.780 | 4.402 | 5.969 |
| **F** | 3.914 | 4.074 | 5.616 | 3.021 |
| **G** | 7.369 | 3.619 | 4.910 | 11.315 |
| **H** | 2.662 | 2.124 | 2.255 | 3.252 |
| **I** | 5.617 | 6.140 | 9.577 | 3.427 |
| **K** | 5.893 | 6.486 | 4.690 | 5.992 |
| **L** | 9.058 | 11.986 | 10.256 | 6.304 |
| **M** | 2.343 | 2.796 | 2.227 | 2.054 |
| **N** | 4.200 | 3.273 | 2.664 | 5.595 |
| **P** | 4.645 | 2.383 | 2.032 | 7.537 |
| **Q** | 3.798 | 4.629 | 3.018 | 3.524 |
| **R** | 5.177 | 6.013 | 4.504 | 4.851 |
| **S** | 6.362 | 4.921 | 5.453 | 7.861 |
| **T** | 5.593 | 4.302 | 6.872 | 5.989 |
| **V** | 7.014 | 6.403 | 13.342 | 4.608 |
| **W** | 1.339 | 1.535 | 1.819 | 0.974 |
| **Y** | 3.430 | 3.442 | 5.103 | 2.663 |

**Supplemental Figure S1A**: Multiple sequence alignment with example secondary structure for the acetohydroxyacid synthase (AHAS) group of ThDP enzymes. The PDB ID used in this figure is given. The three state secondary structure assignment (H=helix, B=strand) for the repeat regions of PDB 2pan is shown on top. The identified repeats for each domain are colored red, yellow, green, cyan, blue, and magenta respectively. The conserved residues in the functional domains (repeat 3 position 10 alanine, repeat 5 positions 5, 6) are indicated with bold underlined yellow text where they could be identified from sequence (and red where they could only be identified in the structures). Residue numbers are shown at the end of each line.

```
     --------------------------------------------------------------------------------
2pan MGSSHHHHHHSSGLVPRGSHMA---------------------------------------------------------- 22
5ahk MGSSHHHHHHSSGLVPRGSHMK---------------------------------------------------------- 22
1ybh ---TFISRFAPDQ--PRK-------------------------------------------------------------- 13
6dek ---MHHHHHHSSGLVPRGSGMKETAAAKFERQHMDSPDLGTDDDDKAMAFNTADTSTQPIIN-DPTLNKHQSSAISRK-K 75
1n0h ---MHHHHHHSSGLVPRGSGMKETAAAKFERQHMDSPDLGTDDDDKAMG-------SAPSFNVDPLEQPAEPSKLAKKLR 70

     -------------HHHHHHHHHHH----BBBB---HHHHHHHHHHHHH---BBBB---HHHHHHHHHHHHHH------BB
2pan -------SMAKMRAVDAAMYVLEKEGITTAFGVPGAAINPFYSAMRKHGGIRHILARHVEGASHMAEGYTRATAGN-IGV 94
5ahk -------------ASDAVAKILADNNVLYGFELIGGMITHLVDSINLLGKTKLVSMHHEQGAAFAASAVSRVTHHKTLGL 89
1ybh -------------GADILVEALERQGVETVFAYPGGASMEIHQALTRSSSIRNVLPRHEQGGVFAAEGYARSSGKP--GI 78
6dek KEQLMDDSFIGLTGGEIFHEMMLRHKVDTVFGYAGGAILPVFDAIYNSDKFKFVLPRHEQGAGHMAEGYARASGKP--GV 153
1n0h AEPDMDTSFVGLTGGQIFNEMMSRQNVDTVFGYPGGAILPVYDAIHNSDKFNFVLPKHEQGAGHMAEGYARASGKP--GV 148

     BBB---HHHH--HHHHHHHHH----BBBBBBB--H------HH----------HHHHHHHH--BBBB---HHHHHHHHHH
2pan CLGTSGPAGTDMITALYSASADSIPILCITGQAPR------ARLHKEDFQAVDIEAIAKPVSKMAVTVREAALVPRVLQQ 168
5ahk ALATSGPGATNLITGIADCWLDSHPCIFLTGQVNTHELKGKRDIRQQGFQELDSVALVTSITKYAYQIKSADELVPCLRK 169
1ybh CIATSGPGATNLVSGLADALLDSVPLVAITGQVPR------RMIGTDAFQETPIVEVTRSITKHNYLVMDVEDIPRIIEE 152
6dek VLVTSGPGATNVITPMADALMDGVPLVVFSGQVPT------TAIGTDAFQEADIVGISRSCTKWNVMVKNVAELPRRINE 227
1n0h VLVTSGPGATNVVTPMADAFADGIPMVVFTGQVPT------SAIGTDAFQEADVVGISRSCTKWNVMVKSVEELPLRINE 222

     HHHHHH------BBBBBB--------------------------------HH---HHHHHHHHHH----BBBBB-HHHHH
2pan AFHLMRSGRPGPVLVDLPFDVQVAEIEFDPDMYEPLPVY------KPAASRM---QIEKAVEMLIQAERPVIVAGGGVIN 239
5ahk AIQIAKEGRPGPVLLDIPMDIQRA------DIDEALLNNPMTPEPKVQRPSIAMSDLDFIINKLQNAKKPLLLIGGGAVN 243
1ybh AFFLATSGRPGPVLVDVPKDIQQQLAIPNWEQAMRLPGY-MSRMPKPPEDS----HLEQIVRLISESKKPVLYVGGGCLN 227
6dek AFEIATTGRPGPVLVDLPKDVTASILRESIPINTTLPSNALSQITKKAVSEFTSEAIKRAANILNKAKKPIIYAGAGILN 307
1n0h AFEIATSGRPGPVLVDLPKDVTAAILRNPIPTKTTLPSNALNQLTSRAQDEFVMQSINKAADLINLAKKPVLYVGAGILN 302

     ----HHHHHHHHHHH---BBB--------------B----------HHHHHHHHH--BBBBB-----HHHH---HHHHH-
2pan -ADAAALLQQFAELTSVPVIPTLMGWGCIPDDHELMAGMVGLQTAHRYGNATLLASDMVFGIGNRFANRHTG-SVEKYTE 317
5ahk SSGFQKWLEQI-ELRGIPYVASLKGAEKIKASD-LYLGMLGAY-GTRAANHAVQNCDLLLVLGSRMDVRQTGAQPEDFAR 320
1ybh SSDE---LGRFVELTGIPVASTLMGLGSYPCDDELSLHMLGMH-GTVYANYAVEHSDLLLAFGVRFDDRVTG-KLEAFAS 302
6dek NEQGPKLLKELADKANIPVTTTLQGLGAFDQRDPKSLDMLGMH-GSAAANTAIQNADCIIALGARFDDRVTG-NISKFAP 385
1n0h HADGPRLLKELSDRAQIPVTTTLQGLGSFDQEDPKSLDMLGMH-GCATANLAVQNADLIIAVGARFDDRVTG-NISKFAP 380

     --B---------BBBBB--HHH----------BB----------------------------------------------
2pan GRK---------IVHIDIEPTQIGRVLCPDLGIVSDAKAALTLLVEVAQEMQKAGRLPCRKEWVADCQQRKRTLLRKTHF 388
5ahk NAE---------IIQIDLQEGQLNNRVIADFSYQIELSEYFSRFSPLQIPVNN--------DWSVWTALLKEK-FRVTFI 382
1ybh RAK---------IVHIDIDSAEIGKNKTPHVSVCGDVKLALQGMNKVLENRAEELKLD-FGVWRNELNVQKQK-FPLSF- 370
6dek EAKLAASEGRGGILHFEISPKNINKVVEATEAIEGDVTANLQSFIPLVDSIEN------RPEWFNKINEWKKK-YPYSYQ 458
1n0h EARRAAAEGRGGIIHFEVSPKNINKVVQTQIAVEGDATTNLGKMMSKIFPVKE------RSEWFAQINKWKKE-YPYAYM 453

     ---------HHHHHHHHHH---H----BBBBB---HHHHHHHHH--------BBB---------HHHHHHHHHHH----B
2pan DNVP---VKPQRVYEEMNK---AFGRDVCYVTTIGLSQIAAAQMLHVFKDRHWINCGQAGPLGWTIPAALGVCAADPKRN 462
5ahk DEYTTWNLSPFGLFTQLNKLTERVALDYIL--DVGNNQMWAAHTLRLNAQQAMHHSGGLGSMGFAIPAAIGACYAGKKPI 460
1ybh -KTFGEAIPPQYAIKVLDELTDG---KAIISTGVGQHQMWAAQFYNYKKPRQWLSSGGLGAMGFGLPAAIGASVANPDAI 446
6dek LETPGSLIKPQTLIKEISDQAQTYNKEVIVTTGVGQHQMWAAQHFTWTQPRTMITSGGLGTMGYGLPAAIGAQVAKPDAI 538
1n0h EETPGSKIKPQTVIKKLSKVANDTGRHVIVTTGVGQHQMWAAQHWTWRNPHTFITSGGLGTMGYGLPAAIGAQVAKPESL 533
```

```
     BBBBBBHHHHHH-HHHHHHHHHH-----BBBBBB---------------------------------------HHHHHHH-
2pan VVAISGDFDFQFLIEELAVGAQFNIPYIHVLVNNAYLGLIRQSQRAFDMDYCVQLAFENI-NSSEVNGYGVDHVKVAEGL 541
5ahk IV-ITGDGGAQLNIQELDIIARDKLPILTIVMNNHSLGMVRGFQEMY---FEGRNSSTY------WNGYTSQFKKIGEAY 530
1ybh VVDIDGDGSFIMNVQELATIRVENLPVKVLLLNNQHLGMVMQWEDRF---YKANRAHTFLGDPAQEDEIFPNMLLFAAAC 523
6dek VIDIDGDASFNMTLTELSSAVQAGAPIKVCVLNNEEQGMVTQWQSLF---YEHRYSHTH--------QSNPDFMKLAESM 607
1n0h VIDIDGDASFNMTLTELSSAVQAGTPVKILILNNEEQGMVTQWQSLF---YEHRYSHTH--------QLNPDFIKLAEAM 602
     --BBBBB--HHHHHHHHHHHHHHHHHH---BBBBBBB--------------------------------------
2pan GCKAIRVFKPEDIAPAFEQAKALMAQYRVPVVVEVILERVTNISMGSELDNVMEFEDIADNAADAPTETCFMHYE 616
5ahk RVESKTIISMQAFSSALESFLESPRPLLLEVSMSDARECRPRLEYGRAID------------QQSPRHDG----- 588
1ybh GIPAARVTKKADLREAIQTMLDTPGPYLLDVICPHQEHVLPMIPSGGTFNDVI--------TEGDGRLEHHHHHH 590
6dek NVKGIRITNQQELKSGVKEFLDATEPVLLEVIVEKKVPVLPMVPAGKALDDFILWDAEVEKQQNDLRKERTGGKY 682
1n0h GLKGLRVKKQEELDAKLKEFVSTKGPVLLEVEVDKKVPVLPMVAGGSGLDEFINFDPEVERQQTELRHKRTGGKH 677
```

**Supplemental Figure S1B:** Multiple sequence alignment with example secondary structure for the acetolactatate synthase (ALS) group of ThDP enzymes. The PDB ID used in this figure is given. The three state secondary structure assignment (H=helix, B=strand) for the repeat regions of PDB 2pgn is shown on top. The identified repeats for each domain are colored red, yellow, green, cyan, blue, and magenta respectively. The conserved residues in the functional domains (repeat 3 position 10 alanine, repeat 5 positions 5, 6) are indicated with bold underlined yellow text where they could be identified from sequence (and red where they could only be identified in the structures). Residue numbers are shown at the end of each line.

```
     --------------------------------HHHHHHHHHHH----BBBBB--HHHHHH-HHHHH-------BB----H
2pgn ----------------MAIK-----------RGADLIVEALEEYGTEQVVGFIGHTSHFV-ADAFSKSHLGKRVINPATE 52
1upa --------------MSRVSTAPSGKP-----TAAHALLSRLRDHGVGKVFGVVGREAASILFDEV--E--GIDFVLTRHE 57
5dx6 MGSSHHHHHHSSGLVPRGSHMDKQYPVRQWAHGADLVVSQLEAQGVRQVFGIPGAKIDKV-FDSLLDS--SIRIIPVRHE 77
4rji ----------------MLTKATKEQKSLVKNRGAELVVDCLVEQGVTHVFGIPGAKIDAV-FDALQDK--GPEIIVARHE 61

     HHHHHHHHHHHHHH----BBBBBBHHHHHH-HHHHHHHHH----BBBBBBB--HHH-----------HHH-H-----BBB
2pgn LGGAWMVNGYNYVKDRSAAVGAWHCVGNLLLHAAMQEARTGRIPAVHIGLNSDGRLAGRSEAAQQVPWQS-FTPIARSTQ 131
1upa FTAGVAADVLARITGRPQACWATLGPGMTNLSTGIATSVLDRSPVIALAAQSESHDIFPNDTHQCLDSVAIVAPMSKYAV 137
5dx6 ANAAFMAAAVGRITGKAGVALVTSGPGCSNLITGMATANSEGDPVVALGGAVKRADKAK-QVHQSMDTVAMFSPVTKYAI 156
4rji QNAAFMAQAVGRLTGKPGVVLVTSGPGASNLATGLLTANTEGDPVVALAGNVIRADRLK-RTHQSLDNAALFQPITKYSV 140

     B---HHHHHHHHHHHHHHH------BBBBBBB------------------------------H---HHHHHHHHHHHH--
2pgn RVERLDKVGEAIHEAFRVAEGHPAGPAYVDIPFDLT--ADQIDDKALVPRGATRAKSVLHAPN---EDVREAAAQLVAAK 206
1upa ELQRPHEITDLVDSAVNAAMTEPVGPSFISLPVDLLGSSEGIDTTVPNPPANTPAKPVGVVADGWQKAADQAAALLAEAK 217
5dx6 EVTAPDALAEVVSNAFRAAEQGRPGSAFVSLPQDVV---DGPVSGKVLPASGAP--QMGAAPD---DAIDQVAKLIAQAK 228
4rji EVQDVKNIPEAVTNAFRIASAGQAGAAFVSFPQDVV---NEVTNTKNVRAVAAP--KLGPAAD---DAISAAIAKIQTAK 212

     -BBBBB-HHHHHH--HHHHHHHHHHH---BBB--------------BB-BB-----------HHHHHHHHH--BBBBB--
2pgn NPVILAGGGVARSGGSEALLKLAEMVGVPVVTTSTGAGVFPETHALAM-GSA-----GFCGWKSANDMMAAADFVLVLGS 280
1upa HPVLVVGAAAIRSGAVPAIRALAERLNIPVITTYIAKGVLPVGHELNY-GAVTGYMDGILNFPALQTMFAPVDLVLTVGY 296
5dx6 NPIFLLGLMASQPENSKALRRLLETSHIPVTSTYQAAGAVNQDNFSRFAGRV-----GLFNNQAGDRLLQLADLVICIGY 303
4rji LPVVLVGMKGGRPEAIKAVRKLLKKVQLPFVETYQAAGTLSRDLEDQYFGRI-----GLFRNQPGDLLEQADVVLTIGY 287

     ------------------BBBBB--HHH--------BBBB----------------------------------------
2pgn R-LSDWGIAQGYITKMPKFVHVDTDPAVLGTFYFPLLSVVADAKTFMEQLIEVLPGTSGFKAVRYQERENFRQATEFRAA 359
1upa DYAEDLRPSMWQKGIEKKTVRISPTVNPIPRVYRPDVDVVTDVLAFVEHFE---TATASFGA---KQRHDIEPLRARIAE 370
5dx6 S-PVEYEPAMWN-SGNATLVHIDVLPAYEERNYTPDVELVGDIAGTLNKLAQNIDHRLVLSP---QAAEILRDRQHQREL 378
4rji D-PIEYDPKFWNINGDRTIIHLDEIIADIDHAYQPDLELIGDIPSTINHIEHD-AVKVEFAE---REQKILSDLKQYMHE 362

     ---------------B-HHHHHHHH--HH------BBBB-----HHHHHHH--------BB----------HHHHHHHHH
2pgn WDGWVREQESGDGMPASMFRAMAEV--RKVQRPEDIIVTDIGNHTLPMFGGAILQRPRRLVTSMAEGILGCGFPMALGAQ 437
1upa FLADPETYEDG-MRVHQVIDSMNTVMEEAAEPGEGTIVSDIGFFRHYGVLFARADQPFGFLTSAGCSSFGYGIPAAIGAQ 449
5dx6 LDRRGAQLNQFALHPLRIVRAMQDIVNSDV-----TLTVDMGSFHIWIARYLYSFRARQVMISNGQQTMGVALPWAIGAW 453
4rji GEQVPADWKSDRAHPLEIVKELRNAVDDHV-----TVTCDIGSHAIWMSRYFRSYEPLTLMISNGMQTLGVALPWAIGAS 437

     HH-----BBBBBHHHHHHHHHHHHHHHHH-----BBBBBB-----------------------------HHHHHHHH--B
2pgn LAEPNSRVFLGTGDGALYYHFNEFRVAVEHKLPVITMVFTNESYGANWTLMNHQFGQN--NWTEFMNPDWVGIAKAFGAY 515
1upa MARPDQPTFLIAGDGGFHSNSSDLETIARLNLPIVTVVVNNDTNGLIELYQNIGHHRSHDPAVKFGGVDFVALAEANGVD 529
5dx6 LVNPERKVVSVSGDGGFLQSSMELETAVRLKANVLHLIWVDNGYNMVAIQEEKKYQRL--SGVEFGPMDFKAYAESFGAK 531
4rji LVKPGEKVVSVSGDGGFLFSAMELETAVRLKAPIVHIVWNDSTYDMVAFQQLKKYNRT--SAVDFGNIDIVKYAESFGAT 515

     BBB------HHHHHHHHHHH---BBBBBB------------------------------------------------
2pgn GESVRETGDIAGALQRAIDSGKPALIEIPVSKTQGL---ASDPVGGVGPNLLLKGREIPVDTGGSMYPGENLLHLKS 589
1upa ATRATNREELLAALRKGAELGRPFLIEVPVNYDFQ--------PGGFGALSI------------------------- 573
5dx6 GFAVESAEALEPTLRAAMDVDGPAVVAIPVDYRDN--------PLLMGQLHLSQ-----IL---------------- 579
4rji GLRVESPDQLADVLRQGMNAEGPVIIDVPVDYSDNINLASDKLPKEFGELMKTK-----ALLSGRPVLGSSHHHHHH 587
```

**Supplemental Figure S1C:** Multiple sequence alignment with example secondary structure for the benzoylformate decarboxylase (BFD) group of ThDP enzymes. The PDB ID used in this figure is given. The three state secondary structure assignment (H=helix, B=strand) for the repeat regions of PDB 6a50 is shown on top. The identified repeats for each domain are colored red, yellow, green, cyan, blue, and magenta respectively. The conserved residues in the functional domains (repeat 3 position 10 alanine, repeat 5 positions 5, 6) are indicated with bold underlined yellow text where they could be identified from sequence (and red where they could only be identified in the structures). Residue numbers are shown at the end of each line.

```
     --------------------------------HHHHHHHHHHH----BBBB---HHHHHHH------------BBBB---
6a50 MGSSHHHHHHSSG--LVPRGSHM------MASVHGTTYELLRRQGIDTVFGNPGSNELPFLKDFPE-----DFRYILALQ 67
4k9q -----------------------------MRTVKEITFDLLRKLQVTTVVGNPGSTEETFLKDFPS-----DFNYVLALQ 46
5erx MGSSHHHHHHSSG--LVPRGSHMNP----STTQARVVVDELIRGGVRDVVLCPGSRNAPLAFALQDADRSGRIRLHVRID 74
2jlc MGSSHHHHHHSSGENLYFQGHMSVSAF--NRRWAAVILEALTRHGVRHICIAPGSRSTPLTLAAAE---NSAFIHHTHFD 75
3lq1 MSLTNHEQV--------------------LTDYLAAFIEELVQAGVKEAIISPGSRSTPLALMMAE---HPILKIYVDVD 57
2x7j MGSSHHHHHHSSGENLYFQGHMLEMTVNPITHYIGSFIDEFALSGITDAVVCPGSRSTPLAVLCAA---HPDISVHVQID 77

     HHHHHHHHHHHHHHH---BBBBBBHHHHHHH-HHHHHHHHH----BBBBBBB--HHHH----------HHH-------BB
6a50 EACVVGIADGYAQASRKPAFINLHSAAGTGNAMGALSNARTSHSPLIVTAGQQTRAMIGVEAGETNVDAANLPRPLVKWS 147
4k9q EASVVAIADGLSQSLRKPVIVNIHTGAGLGNAMGCLLTAYQNKTPLIITAGQQTREMLLNEPLLTNIEAINMPKPWVKWS 126
5erx ERTAGYLAIGLAIGAGAPVCVAMTSGTAVANLGPAVVEANYARVPLIVLSANRPYELLGTGANQT-MEQLGYFGTQVRAS 153
2jlc ERGLGHLALGLAKVSKQPVAVIVTSGTAVANLYPALIEAGLTGEKLILLTADRPPELIDCGANQA-IRQPGMFASHPTHS 154
3lq1 ERSAGFFALGLAKASKRPVVLLCTSGTAAANYFPAVAEANLSQIPLIVLTADRPHELRNVGAPQA-MDQLHLYGSHVKDF 136
2x7j ERSAGFFALGLAKAKQRPVLLICTSGTAAANFYPAVVEAHYSRVPIIVLTADRPHELREVGAPQA-INQHFLFGNFVKFF 156

     B----H-HHHHHHH---------HH-----HHHHHH------BBBBBB--------------------------------
6a50 YEPASA-AEVPHAM---------SR-----AIHMASMAPQGPVYLSVPYDDWDK---DADPQSHHLFDRHVS-------- 201
4k9q YEPARP-EDVPGAF---------MR-----AYATAMQQPQGPVFLSLPLDDWEKLIPEVD------VARTVS-------- 177
5erx ISLGLA-EDAPERTSALNATWRSATCRVLAAATGARTANAGPVHFDIPLR--EPLVP--DPEPLGAVTPPGR--PAGKPW 226
2jlc ISLPRPTQDIP-------ARWLVST-----IDHALGTLHAGGVHINCPFA--EPLYGEMDDTGLSWQQRLGDWWQDDKPW 220
3lq1 TDMALP-ENSEEMLR--YAKWHGSR-----AVDIAMKTPRGPVHLNFPLR--EPLVPILEPSPFTATGKKHH--HVHIYY 204
2x7j TDSALP-EESPQMLR--YIRTLASR-----AAGEAQKRPMGPVHVNVPLR--EPLMPDLSDEPFGRM-RTGR--HVSVKT 223

     ------HHHHHHHHHHHHH----BBBB-HHHHH---HHHHHHHHHHH---BBB-----B-----------BBBB----HH
6a50 SSVRLNDQDLDILVKALNSASNPAIVLGPDVDAANANADCVMLAERLKAPVWVAPSAPRCPFPTRH--PCFRGLMPAGIA 279
4k9q TRQGPDPDKVKEFAQRITASKNPLLIYGSDIARSQAWSDGIAFAERLNAPVWAAPFAERTPFPEDH--PLFQGALTSGIG 255
5erx TYTPPVTFDQPLDIDL---SVDTVVISG---HGAGVHPNLAAL------PTVAEPTAPRSG---DN--PLHPLALP---- 285
2jlc LREAPRLESEKQRDWFFWRQKRGVVVAGR--MSAEEGKKVALWAQTLGWPLIGDVLSQTGQ-------PLPCADLWLGNA 291
3lq1 THEVLDDSSIQKMVTE-CTGKKGVFVVGP-IDKKELEQPMVDLAKKLGWPILADPLSGLRSYGALDEVVIDQYDAFLKEA 282
2x7j GTQSVDRESLSDVAEMLAEAEKGMIVCGE-LHSDADKENIIALSKALQYPILADPLSNLRNGVHDKSTVIDAYDSFLKDD 302

     HHHHH-----BBBBB----------------------------------BBBBBB--HHHHHH----BBBB---------
6a50 AISQLLEGHDVVLVIGA-PV-------------FRYVFYDPGQYLKPGTRLISVTCDPLEAARAPMGDAIVADIGAMASA 345
4k9q SLEKQIQGHDLIVVIGA-PV-------------FRYYPWIAGQFIPEGSTLLQVSDDPNMTSKAVVGDSLVSDSKLFLIE 321
5erx -----LLRPQQVIMLGR-PTLHRPVSVLLADAEVPVFALTTGPRWPDVSGNSQATGTRAVTTGAP--------------R 345
2jlc KATSELQQAQIVVQLGSSLTGKRLLQWQASCEPEEYWIVDDIEGRLD---PAHHRGRRLIANIADWLE-----LHPAEKR 363
3lq1 EIIDKL-TPEVVIRFGSMPVSKPLKNWLEQLSDIRFYVVDPGAAWKD---PIKAVTDMIHCDERFLLDIMQQNMPDDAKD 358
2x7j ELKRKL-RPDVVIRFGPMPVSKPVFLWLKDDPTIQQIVIDEDGGWRD---PTQASAHMIHCNASVFAEEIMAGLTAATRS 378

     ------------------------------HHHHHHHHHHH-----BBBBB-HHHHHHH--HHH--------BBB-----
6a50 LANLVEESSRQLPTAAPEPAKVDQDAGRLHPETVFDTLNDMAPENAIYLNESTSTTAQM--WQRLNMRNPGSYYFCAAGG 423
4k9q ALKLIDQREKNNTPQRSPMTKEDRTAMPLRPHAVLEVLKENSPKEIVLVEECPSIVPLMQDVFRIN--QPDTFYTFASGG 399
5erx PAWLDRCAAMNRHAIAAVREQLAAHPLTTGLHVAAAVSHALRPGDQLVLGASNPVRDVA--LAGLDTRGIRVRSNRGVAG 423
2jlc QPW---CVEIPRLAEQAMQAVIARRDAFGEAQLAHRICDYLPEQGQLFVGNSLVVRLIDA-LSQLP-AGYPVYSNRGASG 438
3lq1 AAWLNGWTSYNKVAREIVLAEMANTTILEEGKIVAELRRLLPDKAGLFIGNSMPIRDVDTYFSQID-KKIKMLANRGANG 437
2x7j SEWLEKWQFVNGRFREHLQT-ISSEDVSFEGNLYRILQHLVPENSSLFVGNSMPIRDVDTFFEKQD-RPFRIYSNRGANG 456
```

```
     ---HHHHHHHHHHH----------------------BBBBBBHH-----HHHHHHHHH-------BBBBBB---------
6a50 LGFALPAAIGVQLAEPE-----RQ------------VIAVIGDGSANYSISALWTAAQYNIP--TIFVIMNNGTYGMLRW 484
4k9q LGWDLPAAVGLALGE-EVSGRNRP------------VVTLMGDGSFQYSVQGIYTGVQQKTH--VIYVVFQNEEYGILKQ 464
5erx IDGTVSTAIGAALAY-E-----GAHERTGSPDSPPRTIALIGDLTFVHDSSGLLIGPTEPIPRSLTIVVSNDNGGGIFEL 497
2jlc IDGLLSTAAGVQRAS-G-----KP------------TLAIVGDLSALYDLNALALLRQVSAP--LVLIVVNNNGGQIFSL 498
3lq1 IDGVVSSALGASVVF-------QP------------MFLLIGDLSFYHDMNGLLMAKKYKMN--LTIVIVNNDGGGIFSF 496
2x7j IDGVVSSAMGVCEGT-K-----AP------------VTLVIGDLSFYHDLNGLLAAKKLGIP--LTVILVNNDGGGIFSF 516
     --------------------------HHHHH---BBBBB--HHHHHHHHHHHHH----BBBBBB----------------
6a50 FAG--VLEAE---NVPGLDVPGIDFRALAKGYGVQALKADNLEQLKGSLQEALSAKGPVLIEVSTVS-----------PV 548
4k9q FAE--LEQTP---NVPGLDLPGLDIVAQGKAYGAKSLKVETLDELKTAYLEALSFKGTSVIVVPITK-----------EL 528
5erx LEQGDPRFSDVSSRIFGTP-HDVDVGALCRAYHVESRQIE-VDELGPTLDQP--GAGMRVLEVKADRSSLRQLHAAIKAA 573
2jlc LPT--PQSER--ERFYLMP-QNVHFEHAAAMFELKYHRPQNWQELETAFADAWRTPTTTVIEMVVNDTDGAQTLQQLLAQ 573
3lq1 LPQ--ANEPKYFESLFGTS-TELDFRFAAAFYDADYHEAKSVDELEEAIDKASYHKGLDIIEVKTNR------HENKANH 567
2x7j LPQ--ASEKTHFEDLFGTP-TGLDFKHAAALYGGTYSCPASWDEFKTAYAPQADKPGLHLIEIKTDRQSRVQLHRDMLNE 593

     -----------
6a50 K---------- 549
4k9q KPLFGHHHHHH 539
5erx L---------- 574
2jlc VS-------HL 577
3lq1 QALEGHHHHHH 578
2x7j AVREVKKQWEL 604
```

**Supplemental Figure S1D:** Multiple sequence alignment with example secondary structure for the oxalyl CoA decarboxylase (OxCDC) group of ThDP enzymes. The PDB ID used in this figure is given. The three state secondary structure assignment (H=helix, B=strand) for the repeat regions of PDB 4qq8 is shown on top. The identified repeats for each domain are colored red, yellow, green, cyan, blue, and magenta respectively. The conserved residues in the functional domains (repeat 3 position 10 alanine, repeat 5 positions 5, 6) are indicated with bold underlined yellow text where they could be identified from sequence (and red where they could only be identified in the structures). Residue numbers are shown at the end of each line.

```
     -----------HHHHHHHHHHH-----BBBB--HHHHHHHHHHHHHH---BBB---HHHHHHHHHHHHHHHH---BBBBB--
4qq8 ------MAMITGGELVVRTLIKAGVEHLFGLHGIHIDTIFQACLDHDVPIIDTRHEAAAGHAAEGYARAGAKLGVALVTA 74
2q27 --MSDQLQMTDGMHIIVEALKQNNIDTIYGVVGIPVTDMARHAQAEGIRYIGFRHEQSAGYAAAASGFLTQKPGICLTVS 78
2c31 MSNDDNVELTDGFHVLIDALKMNDIDTMYGVVGIPITNLARMWQDDGQRFYSFRHEQHAGYAASIAGYIEGKPGVCLTVS 80

     -HHHHH-HHHHHHHHHH---BBBBBBB--HHH--------------HHHHHHHH--BBBB---HHHHHHHHHHHHHHHH
4qq8 GGGFTNAVTPIANARTDRTPVLFLTGSGALRDDETNTLQAG----IDQVAMAAPITKWAHRVMATEHIPRLVMQAIRAAL 150
2q27 APGFLNGLTALANATVNGFPMIMISGSS---DRAIVDLQQGDYEELDQMNAAKPYAKAAFRVNQPQDLGIALARAIRVSV 155
2c31 APGFLNGVTSLAHATTNCFPMILLSGSS---EREIVDLQQGDYEEMDQMNVARPHCKASFRINSIKDIPIGIARAVRTAV 157

     ------BBBBBB----------------------------HHHHHHHHHHHHH---BBBBB-HHHHHH--HHHHHHHH
4qq8 SAPRGPVLLDLPWDILMNQI--DEDSVIIPDLVLSAHGAHPDPADLDQALALLRKAERPVIVLGSEASRTARKTALSAFV 228
2q27 SGRPGGVYLDLPANVLAATMEKDEALTTIVKVENPSPALLPCPKSVTSAISLLAKAERPLIILGKGAAYSQADEQLREFI 235
2c31 SGRPGGVYVDLPAKLFGQTISVEEANKLLFKPIDPAPAQIPAEDAIARAADLIKNAKRPVIMLGKGAAYAQCDDEIRALV 237

     HHH---BBB----HHH-----HHHBBBBHHHHHHH--------BBBBB-----HHH------------BBBBBB--HHH
4qq8 AATGVPVFADYEGLSMLSGLPDAMRGGLVQNLYSFAKADAAPDLVLMLGARFGLNTGHGSGQLIPHS-AQVIQVDPDACE 307
2q27 ESAQIPFLP----MSMAKGILEDTHPLSAAAARSFALANA--DVVMLVGARLNWLLAHGK-KGWAAD-TQFIQLDIEPQE 307
2c31 EETGIPFLP----MGMAKGLLPDNHPQSAAATRAFALAQC--DVCVLIGARLNWLMQHGKGKTWGDELKKYVQIDIQANE 311

     --------BBBB-------------------------------------------------------HHHHHHHHHHHH-
4qq8 LGRLQGIALGIVADVGGTIEALAQATAQDAAWPDRGDWCAKVTDLAQERYASIAAKSSSEHA---LHPFHASQVIAKHV- 383
2q27 IDSNRPIAVPVVGDIASSMQGML-AELKQNTFTTPLVWRDILNIHKQQNAQKMHEKLSTDTQP--LNYFNALSAVRDVLR 384
2c31 MDSNQPIAAPVVGDIKSAV-SLLRKALKGAPKAD-AEWTGALKAKVDGNKAKLAGKMTAETPSGMMNYSNSLGVVRDFML 389

     ----BBBBB---HHHHHHHHHH------BBB----------HHHHHHHHHHHHHH---BBBBBBBHHHHHHH--HHHHHH
4qq8 -DAGVTVVADGGLTYLWLSEVMSRVKPGGFLCHGYLNSMGVGFGTALGAQVADLEAGRRTILVTGDGSVGYSIGEFDTLV 462
2q27 ENQDIYLVNEGANTLDNARNIIDMYKPRRRLDCGTWGVMGIGMGYAIGASVT---SGSPVVAIEGDSAFGFSGMEIETIC 461
2c31 ANPDISLVNEGANALDNTRMIVDMLKPRKRLDSGTWGVMGIGMGYCVAAAAV---TGKPVIAVEGDSAFGFSGMELETIC 466

     H-----BBBBBB-----------------------------HHHHHHH---BBBBB--HHHHHHHHHHHHH----BBB
4qq8 RKQLPLIVIIMNNQSWGWTLHFQQLAV-GPNRVTGTR-LENGSYHGVAAAFGADGYHVDSVESFSAALAQALAHNRPACI 540
2q27 RYNLPVTIVIFNN---GGIYRGDGVDLSGAGAPSPTDLLHHARYDKLMDAFRGVGYNVTTTDELRHALTTGIQSRKPTII 538
2c31 RYNLPVTVIIMNN---GGIYKGNEADP-QPGVISCTR-LTRGRYDMMMEAFGGKGYVANTPAELKAALEEAVASGKPCLI 541

     BBB-------------------------------------
4qq8 NVAVALDPIPPEELILIGMDPFAGSTENLYFQSGALEHHHHHH 583
2q27 NVVI---------------DPAAG-TESGHITKLNPKQVAGN- 564
2c31 NAMI---------------DPDAG-VESGRIKSLNVVSKVGKK 568
```

**Supplemental Figure S1E:** Multiple sequence alignment with example secondary structure for the pyruvate decarboxylase (PDC) group of ThDP enzymes. The PDB ID used in this figure is given. The three state secondary structure assignment (H=helix, B=strand) for the repeat regions of PDB 2nxw is shown on top. The identified repeats for each domain are colored red, yellow, green, cyan, blue, and magenta respectively. The conserved residues in the functional domains (repeat 3 position 10 alanine, repeat 5 positions 5, 6) are indicated with bold underlined yellow text where they could be identified from sequence (and red where they could only be identified in the structures). Residue numbers are shown at the end of each line.

```
     --------------------------HHHHHHHHHHH-----BBB---HHHHHHHHHHHHHH----BBB---HHHHHHHHH
2nxw MGSSHHHHHHSSGLVPRG----SHMKLAEALLRALKDRGAQAMFGIPGDFALPFFKVAEETQILPLHTLSHEPAVGFAAD 76
5tma ----------------------MDYTVGTYLAERLVQIGLKHHFAVAGDYNLVLLDNLLDNKNMEQVYCCNELNCGFSAE 58
5npu ----------------------MTYTVGHYLATRLAQIGLKHHFAVAGDYNLVLLDQLLKNKDLEQVYCCNELNCGFSAE 58
4cok ----------------------MTYTVGRYLADRLAQIGLKHHFAVAGDYNLVLLDQLLLNTDMQQIYCSNELNCGFSAE 58
5euj -----------------------MYTVGMYLAERLAQIGLKHHFAVAGDYNLVLLDQLLLNKDMEQVYCCNELNCGFSAE 57
2vbi ----------------------MTYTVGMYLAERLVQIGLKHHFAVAGDYNLVLLDQLLLNKDMKQIYCCNELNCGFSAE 58
2vjy ---------------------MSEITLGRYLFERLKQVEVQTIFGLPGDFNLSLLDNIYEVPGMRWAGNANELNAAYAAD 59
1qpb ---------------------MSEITLGKYLFERLKQVNVNTVFGLPGDFNLSLLDKIYEVEGMRWAGNANELNAAYAAD 59
2vbf MGSSHHHHHHSSGLVPRGSHMASMYTVGDYLLDRLHELGIEEIFGVPGDYNLQFLDQIISREDMKWIGNANELNASYMAD 80
1ovm --------------------MRTPYCVADYLLDRLTDCGADHLFGVPGDYNLQFLDHVIDSPDICWVGCANELNASYAAD 60

     HHHHHH---BBBBB---HHHH--HHHHHHHHH----BBBBBBB----------------------HHHHHHH-----BBB
2nxw AAARYSSTLGVAAVTYGAGAFNMVNAVAGAYAEKSPVVVISGAPGTTEGNAGLLLHHQ--GRTLDTQFQVFKEITVAQAR 154
5tma GYAR-AKGAAAAVVTYSVGALSAFDAIGGAYAENLPVILISGAPNNNDHAAAHVLHHALGKTDYHYQLEMAKNITAAAEA 137
5npu GYAR-ANGVGAAVVTFSVGALSAFNAIGGAYAENLPVILISGAPNTNDHGSGHILHHTIGTTDYGYQLEMAKQITCAAVS 137
4cok GYAR-ANGAAAAIVTFSVGALSAFNALGGAYAENLPVILISGAPNANDHGTGHILHHTLGTTDYGYQLEMARHITCAAES 137
5euj GYAR-ARGAAAAIVTFSVGAISAMNAIGGAYAENLPVILISGSPNTNDYGTGHILHHTIGTTDYNYQLEMVKHVTCAAES 136
2vbi GYAR-SNGAAAAVVTFSVGAISAMNALGGAYAENLPVILISGAPNSNDQGTGHILHHTIGKTDYSYQLEMARQVTCAAES 137
2vjy GYAR-LKGMSCIITTFGVGELSALNGIAGSYAEHVGVLHVVGVPSVSSQAKQLLLHHTLGNGDFTVFHRMSSNISETTAM 138
1qpb GYAR-IKGMSCIITTFGVGELSALNGIAGSYAEHVGVLHVVGVPSISAQAKQLLLHHTLGNGDFTVFHRMSANISETTAM 138
2vbf GYAR-TKKAAAFLTTFGVGELSAINGLAGSYAENLPVVEIVGSPTSKVQNDGKFVHHTLADGDFKHFMKMHEPVTAARTL 159
1ovm GYAR-CKGFAALLTTFGVGELSAMNGIAGSYAEHVPVLHIVGAPGTAAQQRGELLHHTLGDGEFRHFYHMSEPITVAQAV 139

     ------HHHHHHHHHHHHHHH---BBBBBB---------------------------HHHHHHHHHHHHHHH---BBBBB
2nxw LDDPAKAPAEIARVLGAARAQSRPVYLEIPRNMVNAEVEPVGDDPAW-----PVDRDALAACADEVLAAMRSATSPVLMV 229
5tma IYTPEEAPAKIDHVIKTALREKKPVYLEIACNIASMPCAAPGPASAL-FNDEASDEASLNAAVEETLKFIENRDKVAVLV 216
5npu ITHAEDAPALIDHAIRTALREKKPAYIEIACNVAAQPCARPGPVSAL-LNEPTSDEETLKAAVEAALDFIEKREKPVLLV 216
4cok IVAAEDAPAKIDHVIRTALREKKPAYLEIACNVAGAPCVRPGGIDAL-LSPPAPDEASLKAAVDAALAFIEQRGSVTMLV 216
5euj IVSAEEAPAKIDHVIRTALRERKPAYLEIACNVAGAECVRPGPINSL-LRELEVDQTSVTAAVDAAVEWLQDRQNVVMLV 215
2vbi ITDAHSAPAKIDHVIRTALRERKPAYLDIACNIASEPCVRPGPVSSL-LSEPEIDHTSLKAAVDATVALLEKSASPVMLL 216
2vjy ITDINTAPAEIDRCIRTTYVSQRPVYLGLPANLVDLTVPASLLDTPIDLSLKPNDPEAEEEVIENVLQLIKEAKNPVILA 218
1qpb ITDIATAPAEIDRCIRTTYVTQRPVYLGLPANLVDLNVPAKLLQTPIDMSLKPNDAESEKEVIDTILVLIKDAKNPVILA 218
2vbf LTA-ENATYEIDRVLSQLLKERKPVYINLPVDVAAAKAEKPAL--SL-EKESSTTNTTEQVILSKIEESLKNAQKPVVIA 235
1ovm LTE-QNACYEIDRVLTTMLRERRPGYLMLPADVAKKAATPP--VNALTHKQAHADSACLKAFRDAAENKLAMSKRTALLA 216

     -HHHHH---HHHHHHHHHHH---BBB-HHH-------------B--HHH--HHHHHHHH---BBBBB---B------B-H
2nxw CVEVRRYGLEAKVAELAQRLGVPVVTTFMGRGLLADAPTPPLGTYIGVAGDAEITRLVEESDGLFLLGAILSDTNFAVSQ 309
5tma GSKLRAAAAEEAAVKFADALGGAVATMAAAKSFFPEENPHYIGTSWGEVSYPGVEKTMKEADAVIALAPVFNDYSTTGWT 296
5npu GGKLRAAGAEEAVVELADALGCAVATMAAAKSFFPEDHPGYVGTYWGEVSSPGVEEIVDWADGIICLGPVFNDYSTVGWT 296
4cok GSRIRAAGAQAQAVALADALGCAVTTMAAAKSFFPEDHPGYRGHYWGEVSSPGAQQAVEGADGVICLAPVFNDYATVGWS 296
5euj GSKLRAAAAEKQAVALADRLGCAVTIMAAAKGFFPEDHPNFRGLYWGEVSSEGAQELVENADAILCLAPVFNDYATVGWN 295
2vbi GSKLRAANALAATETLADKLQCAVTIMAAAKGFFPEDHAGFRGLYWGEVSNPGVQELVETSDALLCIAPVFNDYSTVGWS 296
2vjy DACCSRHDAKAETKKLIDLTQFPAFVTPMGKGSIDEKHPRFGGVYVGTLSSPAVKEAVESADLVLSVGALLSDFNTGSFS 298
1qpb DACCSRHDVKAETKKLIDLTQFPAFVTPMGKGSIDEQHPRYGGVYVGTLSKPEVKEAVESADLILSVGALLSDFNTGSFS 298
2vbf GHEVISFGLEKTVTQFVSETKLPITTLNFGKSAVDESLPSFLGIYNGKLSEISLKNFVESADFILMLGVKLTDSSTGAFT 315
1ovm DFLVLRHGLKHALQKWVKEVPMAHATMLMGKGIFDERQAGFYGTYSGSASTGAVKEAIEGADTVLCVGTRFTDTLTAGFT 296
```

```
     HH--HHHBBBBB-----------------------------------------------------------HHHHHHHHH
2nxw RKIDLRKTIHAFDRAVTLGYHTYADIPLAGLVDALLERLPPSDRT-----TRGKEPHAYPTGLQADGEPIAPMDIARAVN 384
5tma DIPDPKKLVLAEPRSVVVNGIRFPSVHLKDYLTRLAEKVSKKTGALDFFKSLNAGELKKADPA-DPSAPLVNAEIARQIE 375
5npu AWPKGENVVLVDPHHITVGGEEFTGIHLKDFLTALTERVPKKDATLDQFKARVGKPAAEKVPAADPNAPLTRAELCRQIQ 376
4cok AWPKGDNVMLVERHAVTVGGVAYAGIDMRDFLTRLAAHTVRRDAT-----ARGGAYVTPQTPAAAPTAPLNNAEMARQIG 371
5euj SWPKGDNVMVMDTDRVTFAGQSFEGLSLSTFAAALAEKAPSRPAT-----TQGTQAPVLGIEAAEPNAPLTNDEMTRQIQ 370
2vbi AWPKGPNVILAEPDRVTVDGRAYDGFTLRAFLQALAEKAPARPAS-----AQKSSVPTCSLTATSDEAGLTNDEIVRHIN 371
2vjy YSYKTKNIVEFHSDYTKIRSATFPGVQMKFALQKLLTKVADAAKG---YKPVPVPSEPEHNEAVADSTPLKQEWVWTQVG 375
1qpb YSYKTKNIVEFHSDHMKIRNATFPGVQMKFVLQKLLTAIADAAKG---YKPVAVPARTPANAAVPASTPLKQEWMWNQLG 375
2vbf HHLDENKMISLNIDEGIIFNKVVEDFDFRAVVSSLSELKGIE--------YEGQYIDKQYEEFIPSSAPLSQDRLWQAVE 387
1ovm HQLTPAQTIEVQPHAARVGDVWFTGIPMNQAIETLVELCKQHVH------AGLMSSSSGAIPFPQPDGSLTQENFWRTLQ 370

     HHHH------BBBB---HHHHHH----------BB----------HHHHHHHHHHH---------BBBBBHHHHHHHHHH
2nxw DRVRAGQEPLLIAADMGDCLFTAMDMI---DAGLMAPGYYAGMGFGVPAGIGAQCV----SGGKRILTVVGDGAFQMTGW 457
5tma DLLTPNT---TVIAETGDSWFNAQRMKLPNGARVEYEMQWGHIGWSVPAAFGYAVG----APERRNILMVGDGSFQLTAQ 448
5npu GLLNPNT---TLIAETGDSWFNAMRMKLPHGARVELEMQWGHIGWSVPATFGYAVA----EPERRNVLMVGDGSFQLTAQ 449
4cok ALLTPRT---TLTAETGDSWFNAVRMKLPHGARVELEMQWGHIGWSVPAAFGNALA----APERQHVLMVGDGSFQLTAQ 444
5euj SLITSDT---TLTAETGDSWFNASRMPIPGGARVELEMQWGHIGWSVPSAFGNAVG----SPERRHIMMVGDGSFQLTAQ 443
2vbi ALLTSNT---TLVAETGDSWFNAMRMTLPRGARVELEMQWGHIGWSVPSAFGNAMG----SQDRQHVVMVGDGSFQLTAQ 444
2vjy EFLREGD---VVITETGTSAFGINQTHFPNNTYGISQVLWGSIGFTTGATLGAAFAAEEIDPKKRVILFIGDGSLQLTVQ 452
1qpb NFLQEGD---VVIAETGTSAFGINQTTFPNNTYGISQVLWGSIGFTTGATLGAAFAAEEIDPKKRVILFIGDGSLQLTVQ 452
2vbf SLTQSNE---TIVAEQGTSFFGASTIFLKSNSRFIGQPLWGSIGYTFPAALGSQIA----DKESRHLLFIGDGSLQLTVQ 460
1ovm TFIRPGD---IILADQGTSAFGAIDLRLPADVNFIVQPLWGSIGYTLAAAFGAQTA----CPNRRVIVLTGDGAAQLTIQ 443

     HHHHHHHH----BBBBBB-------------------------HHHHHHHH-----------BBBBB-BHHHHHH-HHHH
2nxw ELGNCRRLGIDPIVILFNNASWEMLRTFQ--PESAFNDLDDWRFADMAAGMGG---------DGVRVRTRAELKA-ALDK 525
5tma EVAQMVRLKLPVIIFLINNYGYTIEVMIHDGP---YNNIKNWDYAALMEVFNGNGGYDSGAGKGLKAKTAAELEE-AIKV 524
5npu EVAQMVRRKLPIIIFLINNRGYTIEVKIHDGP---YNNIKNWDYAGLMEVFNA----EDGKGLGLKATTGGELAE-AIKK 521
4cok EVAQMIRHDLPVIIFLINNHGYTIEVMIHDGP---YNNVKNWDYAGLMEVFNA----GEGNGLGLRARTGGELAA-AIEQ 516
5euj EVAQMIRYEIPVIIFLINNRGYVIEIAIHDGP---YNYIKNWNYAGLIDVFND----EDGHGLGLKASTGAELEG-AIKK 515
2vbi EVAQMVRYELPVIIFLINNRGYVIEIAIHDGP---YNYIKNWDYAGLMEVFNA----GEGHGLGLKATTPKELTE-AIAR 516
2vjy EISTMIRWGLKPYLFVLNNDGYTIERLIH-GETAQYNCIQNWQHLELLPTFGA------KDYEAVRVSTTGEWNKLTTDE 525
1qpb EISTMIRWGLKPYLFVLNNDGYTIEKLIH-GPKAQYNEIQGWDHLSLLPTFGA------KDYETHRVATTGEWDKLTQDK 525
2vbf ELGLSIREKLNPICFIINNDGYTVEREIH-GPTQSYNDIPMWNYSKLPETFGA----TEDRVVSKIVRTENEFVS-VMKE 534
1ovm ELGSMLRDKQHPIILVLNNEGYTVERAIH-GAEQRYNDIALWNWTHIPQALSL-----DPQSECWRVSEAEQLAD-VLEK 516

     HHH-----BBBBBB--------------------------------------------
2nxw AFATRGRFQLIEAMIPRGVLSDTLARFVQGQKRLHAAPRE------------------ 565
5tma ALDNTDGPTLIECFIAREDCTEELVKWGERVAAANSRKPVNKLL------LEHHHHHH 576
5npu ALAHREGPTLIECVIDRDDCTPELVTWGKKVATANARPPQAI---------------- 563
4cok ARANRNGPTLIECTLDRDDCTQELVTWGKRVAAANARPPRAG---------------- 558
5euj ALDNRRGPTLIECNIAQDDCTETLIAWGKRVAATNSRKPQALVPRGSGGGLEHHHHHH 573
2vbi AKANTRGPTLIECQIDRTDCTDMLVQWGRKVASTNARKTTLA--------LEHHHHHH 566
2vjy KFQDNTRIRLIEVMLPTMDAPSNLVKQAQLTAATNAKN-------------------- 563
1qpb SFNDNSKIRMIEVMLPVFDAPQNLVEQAKLTAATNAKQ-------------------- 563
2vbf AQADVNRMYWIELVLEKEDAPKLLKKMGKLFAEQNK---------------------- 570
1ovm -VAHHERLSLIEVMLPKADIPPLLGALTKALEACNNA--------------------- 552
```

**Supplemental Figure S1F:** Multiple sequence alignment with example secondary structure for the pyruvate dehydrogenase (PDH) group of ThDP enzymes. The PDB ID used in this figure is given, for this protein class the A and B chains were stitched together to give a three domain set. The three state secondary structure assignment (H=helix, B=strand) for the repeat regions of PDB 1ni4 is shown on top. The identified repeats for each domain are colored red, yellow, green, cyan, blue, and magenta respectively. The conserved residues in the functional domains (repeat 3 position 10 alanine, repeat 5 positions 5, 6) are indicated with bold underlined yellow text where they could be identified from sequence (and red where they could only be identified in the structures). Note that this group of proteins is made up of two separate chains which are split into two groups in the phylogenic tree (Fig. 2).

```
     --------------------------------------------------------------------------------
1ni4 -------------------MRGSF--------------------------ANDATFEI-KKCDLHRLEEGPPVTTVLTRE 34
3exe --MGSSHHHHHHSSGLVPRGSHMF--------------------------ANDATFEI-KKCDLHRLEEGPPVTTVLTRE 51
1dtw --------SSLDDKPQFPGASAEF-----------------IDKLEFIQPNVISGIPIYRVMDRQGQIINPSEDPHLPKE 55
2j9f --------SSLDDKPQFPGASAEF-----------------IDKLEFIQPNVISGIPIYRVMDRQGQIINPSEDPHLPKE 55
2bp7 MNEYAPLRLHVPEPTGRPGCQTDFSYLRLNDAGQARKPPVDVDAAD----TADLSYSLVRVLDEQGDAQGPWAE-DIDPQ 75
3duf --------------MGVKTFQFPF-----------------AEQLE----KVAEQFPTFQILNEEGEVVNEEAMPELSDE 45
1umb ----------------MVKETHRF--------------------------ETFTEEPI-RLIGEEGEWLGDFPL-DLEGE 36

     --------------------------------------HHHHHHHHHH------BBB-----------------------
1ni4 DGLKYYRMMQTVRRMELKADQLYKQKIIRGFCHLCDGQEACCVGLEAGINPT-DHLITAYRAHGFTFTRGLSVREILAEL 113
3exe DGLKYYRMMQTVRRMELKADQLYKQKIIRGFCHLCDGQEACCVGLEAGINPT-DHLITAYRAHGFTFTRGLSVREILAEL 130
1dtw KVLKLYKSMTLLNTMDRILYESQRQGRI-SFYMTNYGEEGTHVGSAAALDNT-DLVFGQYREAGVLMYRDYPLELFMAQC 133
2j9f KVLKLYKSMTLLNTMDRILYESQRQGRI-SFYMTNYGEEGTHVGSAAALDNT-DLVFGQYREAGVLMYRDYPLELFMAQC 133
2bp7 ILRQGMRAMLKTRIFDSRMVVAQRQKKM-SFYMQSLGEEAIGSGQALALNRT-DMCFPTYRQQSILMARDVSLVEMICQL 153
3duf QLKELMRRMVYTRILDQRSISLNRQGRL-GFYAPTAGQEASQIASHFALEKE-DFILPGYRDVPQIIWHGLPL----YQA 119
1umb KLRRLYRDMLAARMLDERYTILIRTGKT-SFIAPAAGHEAAQVAIAHAIRPGFDWVFPYYRDHGLALALGIPLKELLGQM 115

     --------------------B--B---------HHHHHHHHHHHHHHH-----BBBBBB--HHH-HHHHHHHHHHHH---
1ni4 TGRKGGCAKGKGGSMH--MYAKNFYGGNGIVGAQVPLGAGIALACKYNGKDEVCLTLYGDGAANQGQIFEAYNMAALWKL 191
3exe TGRKGGCAKGKGGSMH--MYAKNFYGGNGIVGAQVPLGAGIALACKYNGKDEVCLTLYGDGAANQGQIFEAYNMAALWKL 208
1dtw YGNISDLGKGRQMPVHYGCKERHFVTISSPLATQIPQAVGAAYAAKRANANRVVICYFGEGAASEGDAHAGFNFAATLEC 213
2j9f YGNISDLGKGRQMPVHYGCKERHFVTISSPLATQIPQAVGAAYAAKRANANRVVICYFGEGAASEGDAHAGFNFAATLEC 213
2bp7 LSNERDPLKGRQLPIMYSVREAGFFTISGNLATQFVQAVGWAMASAIKGDTKIASAWIGDGATAESDFHTALTFAHVYRA 233
3duf FLFSRGHFHGNQIPE-----GVNVLPPQIIIGAQYIQAAGVALGLKMRGKKAVAITYTGDGGTSQGDFYEGINFAGAFKA 194
1umb LATKADPNKGRQMPEHPGSKALNFFTVASPIASHVPPAAGAAISMKLLRTGQVAVCTFGDGATSEGDWYAGINFAAVQGA 195

     -BBBBBBB--BB--B-BHHHH-----HHH--------BBBBB---HHHHHHHHHHHHHHHH-----BBBBBB--------
1ni4 PCIFICENNRYGMGT-SVERAAASTDYYKRGDF--IPGLRVDGMDILCVREATRFAAAYCRSGKGPILMELQTYRYHGHS 268
3exe PCIFICENNRYGMGT-SVERAAASTDYYKRGDF--IPGLRVDGMDILCVREATRFAAAYCRSGKGPILMELQTYRYHGHS 285
1dtw PIIFFCRNNGYAIST-PTSEQYRGDGIAARGPGYGIMSIRVDGNDVFAVYNATKEARRRAVAENQPFLIEAMTYRIGHHS 292
2j9f PIIFFCRNNGYAIST-PTSEQYRGDGIAARGPGYGIMSIRVDGNDVFAVYNATKEARRRAVAENQPFLIEAMTYRIGHHS 292
2bp7 PVILNVVNNQWAISTFQAIAGGESTTFAGRGVGCGIASLRVDGNDFVAVYAASRWAAERARRGLGPSLIEWVTYRAGPHS 313
3duf PAIFVVQNNRFAIST-PVEKQTVAKTLAQKAVAAGIPGIQVDGMDPLAVYAAVKAARERAINGEGPTLIETLCFRYGPHT 273
1umb PAVFIAENNFYAISV-DYRHQTHSPTIADKAHAFGIPGYLVDGMDVLASYYVVKEAVERARRGEGPSLVELRVYRYGPHS 274

     --------------------------------------------------------------------------------
1ni4 MS-DPGVSYRTREEIQEVRSKSDPIMLLKDRMVNSNLASVEELKEIDVEVRKEIEDAAQFATADPE------PPLEELGY 341
3exe MS-DPGVSYRTREEIQEVRSKSDPIMLLKDRMVNSNLASVEELKEIDVEVRKEIEDAAQFATADPE------PPLEELGY 358
1dtw TS-DDSSAYRSVDEVNYWDKQDHPISRLRHYLLSQGWWDEEQEKAWRKQSRRKVMEAFEQAERKPK------PNPNLLFS 365
2j9f TS-DDSSAYRPVDEVNYWDKQDHPISRLRHYLLSQGWWDEEQEKAWRKQSRRKVMEAFEQAERKPK------PNPNLLFS 365
2bp7 TS-DDPSKYRPADDWSHFPLG-DPIARLKQHLIKIGHWSEEEHQATTAEFEAAVIAAQKEAEQYGTLANGHIPSAASMFE 391
3duf MSGDDPTRYRSKELENEWAKK-DPLVRFRKFLEAKGLWSEEEENNVIEQAKEEIKEAIKKADETPK------QKVTDLIS 346
1umb SA-DDDSRYRPKEEVAFWRKK-DPIPRFRRFLEARGLWNEEWEEDVREEIRAELERGLKEAEEAGP------VPPEWMFE 346
```

```
     ----------------------------------------------------------HHHHHHHHHHHHHHH---BBBB
1ni4 HIYSSDPPFEVRGANQWIKFKSVSMRGSHHHHHHGSL------------------QVTVRDAINQGMDEELERDEKVFLL 403
3exe HIYSSDPPFEVRGANQWIKFKSVSL------------------------------QVTVRDAINQGMDEELERDEKVFLL 408
1dtw DVYQEM-PAQLRKQQESLARHLQTYGEHYPLDHFDKVAHFTFQPDPEPREYGQTQKMNLFQSVTSALDNSLAKDPTAVIF 444
2j9f DVYQEM-PAQLRKQQESLARHLQTYGEHYPLDHFDKVAHFTFQPDPEPREYGQTQKMNLFQSVTSALDNSLAKDPTAVIF 444
2bp7 DVYKEM-PDHLRRQRQELGVMATTT-------------------------------MTMIQALRSAMDVMLERDDNVVVY 439
3duf IMFEEL-PFNLKEQYEIYKEKESKMA-----------------------------QMTMVQAITDALRIELKNDPNVLIF 396
1umb DVFAEK-PWHLLRQEALLKEELMAL-------------------------------MTMVQALNRALDEEMAKDPRVVVL 394
     B---------------HHHHH----BBB----HHHHHHHHHHHHH---BBBBB---HHHHHHHHHHHHH----HHHH---
1ni4 GEEVAQYDGAYKVSRGLWKKYGDKRIIDTPISEMGFAGIAVGAAMAGLRPICEFMTFNFSMQAIDQVINSAAKTYYMSGG 483
3exe GEEVAQYDGAYKVSRGLWKKYGDKRIIDTPISEMGFAGIAVGAAMAGLRPICEFMTFNFSMQAIDQVINSAAKTYYMSGG 488
1dtw GEDVA-FGGVFRCTVGLRDKYGKDRVFNTPLCEQGIVGFGIGIAVTGATAIAEIQFADYIFPAFDQIVNEAAKYRYRSGD 523
2j9f GEDVA-FGGVFRCTVGLRDKYGKDRVFNTPLCEQGIVGFGIGIAVTGATAIAEIQFADYIFPAFDQIVNEAAKYRYRSGD 523
2bp7 GQDVGYFGGVFRCTEGLQTKYGKSRVFDAPISESGIVGTAVGMGAYGLRPVVEIQFADYFYPASDQIVSEMARLRYRSAG 519
3duf GEDVGVNGGVFRATEGLQAEFGEDRVFDTPLAESGIGGLAIGLALQGFRPVPEIQFFGFVYEVMDSICGQMARIRYRTGG 476
1umb GEDVGKRGGVFLVTEGLLQKYGPDRVMDTPLSEAAIVGAALGMAAHGLRPVAEIQFADYIFPGFDQLVSQVAKLRYRSGG 474

     -------BBBBB--------HHH----HHHHH-----BBB----HHHHHHHHHHHHH----BBBBB--------------
1ni4 LQPV-PIVFRGPNGASAGVAAQHSQCFAAWYGHCPGLKVVSPWNSEDAKGLIKSAIRDNNPVVVLENELMYGVP------ 556
3exe LQPV-PIVFRGPNGASAGVAAQHSQCFAAWYGHCPGLKVVSPWNSEDAKGLIKSAIRDNNPVVVLENELMYGVP------ 561
1dtw LFNCGSLTIRSPWGCVGHGALYHSQSPEAFFAHCPGIKVVIPRSPFQAKGLLLSCIEDKNPCIFFEPKILY--------- 594
2j9f LFNCGSLTIRSPWGCVGHGALYHSQSPEAFFAHCPGIKVVIPRSPFQAKGLLLSCIEDKNPCIFFEPKILY--------- 594
2bp7 EFIA-PLTLRMPCGGGIYGGQTHSQSPEAMFTQVCGLRTVMPSNPYDAKGLLIASIECDDPVIFLEPKRLYNGPFDGHHD 598
3duf RYHM-PITIRSPFGGGVHTPELHSDSLEGLVAQQPGLKVVIPSTPYDAKGLLISAIRDNDPVIFLEHLKLY--------- 546
1umb QFTA-PLVVRMPSGGGVRGGHHHSQSPEAHFVHTAGLKVVAVSTPYDAKGLLKAAIRDEDPVVFLEPKRLY--------- 544

     -----------HHHH-----------BBBB------------HHHHHHHHHH-HHHH----BBBBB--------HHHHHH
1ni4 -------FEFPPEAQSKDFLIPIGKAKIERQGTHITVVSHSRPVGHCLEAAA-VLSKEGVECEVINMRTIRPMDMETIEA 628
3exe -------FEFPPEAQSKDFLIPIGKAKIERQGTHITVVSHSRPVGHCLEAAA-VLSKEGVECEVINMRTIRPMDMETIEA 633
1dtw -------RAAAEEVPIEPYNIPLSQAEVIQEGSDVTLVAWGTQVHVIREVASMAKEKLGVSCEVIDLRTIIPWDVDTICK 667
2j9f -------RAAAEEVPIEPYNIPLSQAEVIQEGSDVTLVAWGTQVHVIREVASMAKEKLGVSCEVIDLRTIIPWDVDTICK 667
2bp7 RPVTPWSKHPHSAVPDGYYTVPLDKAAITRPGNDVSVLTYGTTVYVAQVAAE----ESGVDAEVIDLRSLWPLDLDTIVE 674
3duf -------RSFRQEVPEGEYTIPIGKADIKREGKDITIIAYGAMVHESLKAAA-ELEKEGISAEVVDLRTVQPLDIETIIG 618
1umb -------RSVKEEVPEEDYTLPIGKAALRREGKDLTLICYGTVMPEVLQAAA-ELAKAGVSAEVLDLRTLMPWDYEAVMN 616

     HHHHH--BBBB--------HHHHHHHHHHH-HHHHH-----BBB------------------------------------
1ni4 SVMKTNHLVTVEGGWPQFGVGAEICARIMEGPAFNFLDAPAVRVTGADVPMPYAKILEDNSIPQVKDIIFAIKKTLNI-- 706
3exe SVMKTNHLVTVEGGWPQFGVGAEICARIMEGPAFNFLDAPAVRVTGADVPMPYAKILEDNSIPQVKDIIFAIKKTLNIFA 713
1dtw SVIKTGRLLISHEAPLTGGFASEISSTVQE-ECFLNLEAPISRVCGYDTPFPHIF--EPFYIPDKWKCYDALRKMINY-- 742
2j9f SVIKTGRLLISHEAPLTGGFASEISSTVQE-ECFLNLEAPISRVCGYDTPFPHIF--EPFYIPDKWKCYDALRKMINYGG 744
2bp7 SVKKTGRCVVVHEATRTCGFGAELVSLVQE-HCFHHLEAPIERVTGWDTPYPHAQ--EWAYFPGPSRVGAALKKVMEV-- 749
3duf SVEKTGRAIVVQEAQRQAGIAANVVAEINE-RAILSLEAPVLRVAAPDTVYPFAQA-ESVWLPNFKDVIETAKKVMNF-- 694
1umb SVAKTGRVVLVSDAPRHASFVSEVAATIAE-DLLDMLLAPPIRVTGFDTPYPYAQ--DKLYLPTVTRILNAAKRALDY-- 691

     -------
1ni4 ------- 706
3exe IKKTLNI 720
1dtw ------- 742
2j9f HHHHHH- 750
2bp7 ------- 749
3duf ------- 694
1umb ------- 691
```

**Supplemental Figure S1G:** Multiple sequence alignment with example secondary structure for the phosphoketolase (PhK) group of ThDP enzymes. The PDB ID used in this figure is given. For this protein class, the A & B chains were stitched together to give a three domain structure. The three state secondary structure assignment (H=helix, B=strand) for the repeat regions of PDB 3ahc is shown on top. The identified repeats for each domain are colored red, yellow, green, cyan, blue, and magenta respectively. The conserved residues in the functional domains (repeat 3 position 10 alanine, repeat 5 positions 5, 6) are indicated with bold underlined yellow text where they could be identified from sequence (and red where they could only be identified in the structures). Residue numbers are shown at the end of each line.

```
     --------------------------------------------------------------------------------
6gua -----------------------------MTEYN---SEAYLKKLDKWWRAATYLGAGMIFLKENPLFSVTGTPIKAENL 48
3ahc MGSSHHHHHHSSGLVPRGSHMTNPVIGTPWQKLDRPVSEEAIEGMDKYWRVTNYMSIGQIYLRSNPLMK---EPFTRDDV 77
3ai7 --------------------MTSPVIGTPWKKLNAPVSEEALEGVDKYWRVANYLSIGQIYLRSNPLMK---EPFTREDV 57

     ---------HHHHHHHHHHHHHHHHHHH-B-B--BB--------------------------------------------
6gua KANPIGHWGTVSGQTFLYAHANRLINKYDQKMFYMGGPGHGGQAMVVPSYLDGSYTEAYPEITQDLEGMSRLFKRFSFPG 128
3ahc KHRLVGHWGTTPGLNFLLAHINRLIADHQQNTVFIMGPGHGGPAGTSQSYVDGTYTEYYPNITKDEAGLQKFFRQFSYPG 157
3ai7 KHRLVGHWGTTPGLNFLIGHINRFIADHGQNTVIIMGPGHGGPAGTSQSYLDGTYTETFPKITKDEAGLQKFFRQFSYPG 137

     -B--------------------HHHHHHHHH------BBBBBBB--HHH-HHHHHHHHHHH--------BBBBBBBB--B
6gua GIGSHMTAQTPGSLHEGGELGYVLSHATGAILDQPEQIAFAVVGDGEAETGPLMTSWHSIKFINPKNDGAILPILDLNGF 208
3ahc GIPSHFAPETPGSIHEGGELGYALSHAYGAVMNNPSLFVPCIIGDGEAETGPLATGWQSNKLVNPRTDGIVLPILHLNGY 237
3ai7 GIPSHFAPETPGSIHEGGELGYALSHAYGAIMDNPSLFVPAIVGDGEAETGPLATGWQSNKLVNPRTDGIVLPILHLNGY 217

     ----B-HHHH--HHHHHHHHHH--BBBBBBB-------HHHHHHHHHHHHHHHHHHHHHHHHHHH------------BBB
6gua KISNPTLFARTSDVDIRKFFEGLGYSP-RYIENDDIHDYMAYHKLAAEVFDKAIEDIHQIQKDAREDNRYQNGEIPAWPI 287
3ahc KIANPTILARISDEELHDFFRGMGYHPYEFVAGFDNEDHMSIHRRFAELFETIFDEICDIKAAAQTDDMTR----PFYPM 313
3ai7 KIANPTILSRISDEELHEFFHGMGYEPYEFVAGFDDEDHMSIHRRFAELWETIWDEICDIKATAQTDNVHR----PFYPM 293

     BBBB----------------------------------------------------------------------------
6gua VIARLPKGWGGPRYNDWSGPKFDGKGMPIEHSFRAHQVPLPLSSKNMGTLPEFVKWMTSYQPETLFNADGSLKEELRDFA 367
3ahc LIFRTPKGWTCPKFID---------GKKTEGSWRAHQVPLASARDTEEHFEVLKGWMESYKPEELFNADGSIKDDVTAFM 384
3ai7 LIFRTPKGWTCPKYID---------GKKTEGSWRSHQVPLASARDTEAHFEVLKNWLESYKPEELFDANGAVKDDVLAFM 364

     -----------------------------------------BBBB--H-----HHHHHHHHHHHHH----BBBBB-----
6gua PKGEMRMASNPVTNGGVDSSNLVLPDWQEF-ANPISENNRGKLLPDTNDNMDMNVLSKYFAEIVKLNPTRFRLFGPDETM 446
3ahc PKGELRIGANPNANGGVIREDLKLPELDQYEVTGVKEYGHGWGQVEAP-----RALGAYCRDIIKNNPDSFRIFGPDETA 459
3ai7 PKGELRIGANPNANGGVIRNDLKLPNLEDYEVKEVAEYGHGWGQLEAT-----RTLGAYTRDIIKNNPRDFRIFGPDETA 439

     ----BBBBB---BB-------------BB----BBB----HHHHHHHHHHHHH---BBBBBBBHHHHHHHHHHHHHHHHH
6gua SNRFWEMFKVTNRQW-MQVIKNPNDEFISPEGRIIDSQLSEHQAEGWLEGYTLTGRTGAFASYESFLRVVDSMLTQHFKW 525
3ahc SNRLNATYEVTDKQWDNGYLSGLVDEHMAVTGQVTE-QLSEHQCEGFLEAYLLTGRHGIWSSYESFVHVIDSMLNQHAKW 538
3ai7 SNRLQASYEVTNKQWDAGYISDEVDEHMHVSGQVVE-QLSEHQMEGFLEAYLLTGRHGIWSSYESFVHVIDSMLNQHAKW 518

     HHHHH-H-------B-BBBBBB--HHH-----HHH----HHHHHHHH-------BBBBB---HHHHHHHHHHHHH-B--B
6gua IRQAA-DQKWRHDYPSLNVISTSTVFQQDHNGYTHQDPGMLTHLAEK---KSDFIRQYLPADGNTLLAVFDRAFQDRSKI 601
3ahc LEATVREIPWRKPISSVNLLVSSHVWRQDHNGFSHQDPGVTSLLINKTFNNDHVTNIYFATDANMLLAISEKCFKSTNKI 618
3ai7 LEATVREIPWRKPIASMNLLVSSHVWRQDHNGFSHQDPGVTSVLLNKCFHNDHVIGIYFATDANMLLAIAEKCYKSTNKI 598

     BBBB-----BBB---HHHHHHHHHH-BBB--------------BBBBBB-HHHHHHHHHHHHHHHH------BBBBBB-B
6gua NHIVASKQPRQQWFTKEEAEKLATDGIATIDWASTAKDGEAVDLVFASAGAEPTIETLAALHLVNEVFPQAKFRYVNVVE 681
3ahc NAIFAGKQPAPTWVTLDEARAELEAGAAEWKWASNAENNDEVQVVLASAGDVPTQELMAASDALNKM--GIKFKVVNVVD 696
3ai7 NAIIAGKQPAATWLTLDEARAELEKGAAAWDWASTAKNNDEAEVVLAAAGDVPTQEIMAASDKLKEL--GVKFKVVNVAD 676

     HHHH--------------HHHHHHHH------BBBBB---HHHHHHH------HHHBBBB--------------------
6gua LGRLQKKKGALNQERELSDEEFEKYFGPSGTPVIFGFHGYEDLIESIFYQR-GHDGLIVHGYREDGDITTTYDMRVYSEL 760
3ahc LLKLQSRE---NNDEALTDEEFTELF-TADKPVLFAYHSYAQDVRGLIYDRPNHDNFHVVGYKEQGSTTTPFDMVRVNDM 772
3ai7 LLSLQSAK---ENDEALTDEEFADIF-TADKPVLFAYHSYAHDVRGLIYDRPNHDNFNVHGYEEEGSTTTPYDMVRVNRI 752
```

```
     --------------------------------------------------------------------------------
6gua DRFHQAIDAMQVLYVNRKVNQGLAKAFIDRMKRTLVKHFEVTRNEGVDIPDFTEWVWSDLKK------------------ 822
3ahc DRYALQAAALKLIDADKYADK------IDELNAFRKKAFQFAVDNGYDIPEFTDWVYPDVKVDETQMLSATAATAGDNE- 845
3ai7 DRYELTAEALRMIDADKYADK------IDELEKFRDEAFQFAVDNGYDHPDYTDWVYSGVNTDKKGAVTATAATAGDNEH 826

     -----
6gua ----- 822
3ahc ----- 845
3ai7 HHHHH 831
```

**Supplemental Figure S1H:** Multiple sequence alignment with example secondary structure for the pyruvate ferredoxin oxidoreductase (PFOR) group of ThDP enzymes. The PDB ID used in this figure is given. For the 5c4i protein, several chains were stitched together to give a chain of the same length as the other members of this class. The three state secondary structure assignment (H=helix, B=strand) for the repeat regions of PDB 5c4i is shown on top. The identified repeats for each domain are colored red, yellow, green, cyan, blue, and magenta respectively. The conserved residues in the functional domains (repeat 3 position 10 alanine, repeat 5 positions 5, 6) are indicated with bold underlined yellow text where they could be identified from sequence (and red where they could only be identified in the structures). Residue numbers are shown at the end of each line.

```
     --------HHHHHHHHHHH----BBBB------HHHHHHHHHHHH----------BBBB---HHHHHHHHHHHHH----
5c4i MGKVRNISGCVAVAHGVRLADVDVICSYPIRPYTGIMSELARMVADGELDA-----EFVHGEGEHAQLSVVYGASAAGAR 75
6cin MPK-QTLDGNTAAAH-VAYAMSEVATIYPITPSSPMAEIADEWAAHGRKNIFGKTLQVAEMQSEAGAAGAVHGSLAAGAL 78
1b0p GKKMMTTDGNTATAH-VAYAMSEVAAIYPITPSSTMGEEADDWAAQGRKNIFGQTLTIREMQSEAGAAGAVHGALAAGAL 79

     BBBBB--HHHHH-HHHHHHHHH-----BBBBB---------------HHHH-------BBBBB--HHHHHHHHHHHHHH
5c4i VFTGSSGVGVTYAMEVYSPISGERLPVQMAIADRTLDPPG--DFGEEHTDAECCRDQGWIQGWASTPQEALDNTLIYYRV 153
6cin TTTFTASQGLLLMIPNMYKIAGELLPCVFHVAARALSTHALSIFG-DHADVMAARQTGFAMLSSASVQEVMDLALVAHLA 157
1b0p TTTFTASQGLLLMIPNMYKISGELLPGVFHVTARAIAAHALSIFG-DHQDIYAARQTGFAMLASSSVQEAHDMALVAHLA 158

     H--------BBBBB-----------------------------------------------------------------
5c4i GEDQRVLLPQYACLDGYFVSHILGPVDIPDEAQVKEF-----LPPYKNHHVLDPRKPQIIGPQIEPAM---GPPLQYQRY 225
6cin TLKARVPFVHF--FDGFRTSHEVQKIDVIEYEDMAKLVDWDAIRAFR-QRALNPEHPHQRGTAQNPDIYFQSREAANPYY 234
1b0p AIESNVPFMHF--FDGFRTSHEIQKIEVLDYADMASLVNQKALAEFR-AKSMNPEHPHVRGTAQNPDIYFQGREAANPYY 235

     -HHH-HHHHHHHHHHHHHHHH-------BBBB----------B---HHHHHHHHHHHHH-----BBBBBB---B---HH
5c4i QAVKGVHKVLEEACDEFARIFGRKYDPYLDEYL-TDDAEVIIFGQGAHMETAKAVARRLRNLGEKVGVARLRTFRPFPTE 304
6cin LATPG---IVAQVMEQVAGLTGRHYHLF--DYAGAPDAERVIVSMGSSCEVIEETVNYLVEKGEKVGLIKVRLFRPFSAE 309
1b0p LKVPG---IVAEYMQKVASLTGRSYKLF--DYVGAPDAERVIVSMGSSCETIEEVINHLAAKGEKIGLIKVRLYRPFVSE 310

     HHHHH------BBBBBB------HHH--HHHHHHHHH-HHH-----BBBBBB--HHH----HHHHHHHHHHHHHHH-HH
5c4i QIKERL-SKFKAIGVLDVSANFGISCSGGVLLSELRAALYDYGDKV-KTVGFVAGLGGEVVTHDEFYRMFQKLKEIA-KT 381
6cin HFLKVLPASVKRIAVLDRTKEPG--SLGEPLYEDVQTVLAEHGKNI-LVVGGRYGLGSKEFNPSMVKAVFDNLAATTPKN 386
1b0p AFFAALPASAKVITVLDRTKEPG--APGDPLYLDVCSAFVERGEAMPKILAGRYGLGSKEFSPAMVKSVYDNMSGAK-KN 387

     ------BBB-----------------------------------HHHHHHHHHHHHHHHH----BBBBBB-HHH-----
5c4i GKVEQTSYWIPFELMSTKDLFAEPNLK---QITVWARG----VVMNKDARDIVVALTEAAAKEGKYVQAWENYVDLPDRI 454
6cin KFTVGITDDVTHTSLEIKE-HIDTSPKGTFRCKFFGLGSDGTVGANKNSIKIIGDHTDM------YAQGYFVY-DSKKSG 458
1b0p HFTVGIEDDVTGTSLPVDNAFADTTPKGTIQCQFWGLGADGTVGANKQAIKIIGDNTDL------FAQGYFSY-DSKKSG 460

      --BBBBBBBBB---------------------HHHH-----------BBBBBB----HHHHHHHH-----BBBBBBB-
5c4i YVPVRAYARISSDPIESKYIYENETPDIVVLVEESLIKGVPILKGIRPGSTLVVNTKRSIDTILEFLGDTGNLAQIVTVD 534
6cin GVTI-SHLRFGKQPIQSAYLIDQ--ADLIACHNPSYVGRYNLLEGIKPGGIFLLNSTWSAEE-MD--------SRLPADM 526
1b0p GITI-SHLRFGEKPIQSTYLVNR--ADYVACHNPAYVGIYDILEGIKDGGTFVLNSPWSSLEDMD--------KHLPSGI 529

     ---------------------------------------HH----HHHHH-----HHHHHHHHH-BBBB----------
5c4i ANSMAEAVMTLSGAEGATDATGIGAG--IAAPIAGAVVKATGIVDVENLAAVVKNPAAMRRGYAEAQVRQLP-------- 604
6cin KRTIATKKLKFYNIDAVKIAQEIGLGSRINVIMQTAFFKIANVIPVDEAIKYIKDSIVKTYGKKGDKILNMNFAAVDRAL 606
1b0p KRTIANKKLKFYNIDAVKIATDVGLGGRINMIMQTAFFKLAGVLPFEKAVDLLKKSIHKAYGKKGEKIVKMNTDAVDQAV 609

     --------------------------------------------------------------------------------
5c4i --------PHEAVEEAAVSATELLRQMPFAGTVPSPVTENEG------------MVTGNWRIQR-------PIIDREAC 656
6cin EALEEIKYPASWADAVDEAAATVTEEPEFIQKVLRPINALKGDELPVSTFTPDGVFPVGTTKYEKRGIAVNIPQWQPENC 686
1b0p TSLQEFKYPDSWKDAPAETKAEPMTN-EFFKNVVKPILTQQGDKLPVSAFEADGRFPLGTSQFEKRGVAINVPQWVPENC 688

     --------------------------------------------------------------------------------
5c4i TECYTCWIYCPDSCIT----------------------RTEEGPVFNMKY----CKGCGLCTAVCPS--GALT------ 701
6cin IQCNQCSLVCPHAAIRPYLAKPADLAGAPETFVTKDAIGKEAAGLKFRIQVSPLDCTGCGNCADVCPAKVKALTMVPLEE 766
1b0p IQCNQCAFVCPHSAILPVLAKEEELVGAPANFTALEAKGKELKGYKFRIQINTLDCMGCGNCADICPPKEKALVMQPLDT 768
```

```
     ----------------------------------------------HHHHHHHHHHHHH---BBBBBB--HHHHHHH---
5c4i ----NVPELDFKDMLDRI------ASIKKAPDEEYYVPGHRTCAGCGPALTYRLVAKAAGPNTIFIGPTGCMYVANTSYG 771
6cin VTAVEEANYNFAEQLPEVKVNFNPATVKGSQFRQPLLEFSGACAGCGETPYVKLVTQLFGDRMIIANATGCSSIWGGSAP 846
1b0p QRDAQVPNLEYAARIPVKSEVLPRDSLKGSQFQEPLMEFSGACSGCGETPYVRVITQLFGERMFIANATGCSSIWGASAP 848

     ---B----------BBBB----HHHHHHHH------------HHHHHHH----------------------HH-------
5c4i CGPWRV-------PWIHAQITNGGAVASGI------------EAAYKAM----------------------IRKKKTDAE 810
6cin ACPYTVNRQGHGPAWASSLFEDNAEFGYGMALAVAKRQDELATAISKALEAPVSAAFKAACEGWLAGKDDADRSREYGDR 926
1b0p SMPYKTNRLGQGPAWGNSLFEDAAEYGFGMNMSMFARRTHLADLAAKALESDASGDVKEALQGWLAGKNDPIKSKEYGDK 928

     --BBB-------------------------------BBBB-HHHH---HHHHHHHHHH----BBBBBB------------
5c4i FPNII-------------------------------VMAGDGGAVDIGLQALSAMLYRGHDVLFICYDNESYANTGIQTS 859
6cin IKALLPGEISQASGEVKDLLLDIDRQKDYLTKKSIWIIGGDGWAYDIGYGGLDHVLASGANVNVLVLDTEVYSNTGGQSS 1006
1b0p LKKLLAGQKDGLLGQIAAM-------SDLYTKKSVWIFGGDGWAYDIGYGGLDHVLASGEDVNVFVMDTEVYSNTGGQSS 1001

     ----------------------------HHHHHH-------BBBBBB---H-HHHHHHHHHHH-----BBBBBB------
5c4i PTTPYGANTTFTPPGEVVPEGKKLFPKDNPKVIAHGHPELKYVATASIGWP-VDLMNKVRKGLNQEGPAYIHIHAPC-PK 937
6cin KATQTGAVARFAAGGKFT--KKKDLGL---MAMSYGYV---YVASVAMGASHSQLMKALIEAEKYDGPSLIIAYAPCINH 1078
1b0p KATPTGAVAKFAAAGKRT--GKKDLAR---MVMTYGYV---YVATVSMGYSKQQFLKVLKEAESFPGPSLVIAYATCINQ 1073

     --------------------------------------------------------------------------------
5c4i GWQFPADKTIEMAKLAVQTGMFQLYEYE---NGEYKLSVKVDKRKP---VSEYMKLQKRFAHLKPEHIAKMQAFVDARCA 1011
6cin GIN--MTYSQREAKKAVEAGYWPLYRYNPQLAQEGKNPFILDYKTPTASFRDFLMGEIRYTSLKKQFPEKAEQLFAKAEA 1156
1b0p GLRKGMGKSQDVMNTAVKSGYWPLFRYDPRLAAQGKNPFQLDSKAPDGSVEEFLMAQNRFAVLDRSFPEDAKRLRAQVAH 1153

     ------------------------------------------------------------------------------
5c4i EVGITVPVVASNA----------------------------------------------------------------- 1024
6cin DAKARLEQYKKLA----------------------EG----------------------------------------- 1171
1b0p ELDVRFKELEHMAATNIFESFAPAGGKADGSVDFGEGAEFCTRDDTPMMARPDSGEACDQNRAGTSEQQGDLSKRTKK 1231
```

**Supplemental Figure S1I:** Multiple sequence alignment with example secondary structure for the pyruvate oxidase (POX) group of ThDP enzymes. The PDB ID used in this figure is given. The three state secondary structure assignment (H=helix, B=strand) for the repeat regions of PDB 3ey9 is shown on top. The identified repeats for each domain are colored red, yellow, green, cyan, blue, and magenta respectively. The conserved residues in the functional domains (repeat 3 position 10 alanine, repeat 5 positions 5, 6) are indicated with bold underlined yellow text where they could be identified from sequence (and red where they could only be identified in the structures). Residue numbers are shown at the end of each line.

```
     --------------HHHHHHHHHH---BBBBB--HHHHHHHHHHH-HH---BBBB---HHHHHHHHHHHHHHH---BBBB
3ey9 --------MKQTVAAYIAKTLESAGVKRIWGVTGDSLNGLSDSLN-RMGTIEWMSTRHEEVAAFAAGAEAQLSGELAVCA 71
2dji -------DNKINIGLAVMKILESWGADTIYGIPSGTLSSLMDAMGEEENNVKFLQVKHEEVGAMAAVMQSKFGGNLGVTV 73
1y9d MVMKQTKQTNILAGAAVIKVLEAWGVDHLYGIPGGSINSIMDALSAERDRIHYIQVRHEEVGAMAAAADAKLTGKIGVCF 80

     B---HHHH--HHHHHHHHH----BBBBBBB--HHH----------HHHH------BBBB---HHHHHHHHHHHHHHHHHH-
3ey9 GSCGPGNLHLINGLFDCHRNHVPVLAIAAHIPSSEIGSGYFQETHPQELFRECSHYCELVSSPEQIPQVLAIAMRKAVLN 151
2dji GSGGPGASHLINGLYDAAMDNIPVVAILGSRPQRELNMDAFQELNQNPMYDHIAVYNRRVAYAEQLPKLVDEAARMAIAK 153
1y9d GSAGPGGTHLMNGLYDAREDHVPVLALIGQFGTTGMNMDTFQEMNENPIYADVADYNVTAVNAATLPHVIDEAIRRAYAH 160

     --BBBBBBB--------------------------------HHHHHHHHHHHH----BBBBB-HHH---HHHHHHHHHHH
3ey9 RGVSVVVLPGDVALKPAPEGATMHWYHAPQPV-----VTPEEEELRKLAQLLRYSSNIALMCGSGCAGAHKELVEFAGKI 226
2dji RGVAVLEVPGDFAKVEIDND---QWYSSANSLRKYAPIAPAAQDIDAAVELLNNSKRPVIYAGIGTMGHGPAVQELARKI 230
1y9d QGVAVVQIPVDLPWQQIPAE---DWYASANSYQTPLLPEPDVQAVTRLTQTLLAAERPLIYYGIGARKAGKELEQLSKTL 237

     ---BBB-HHHHHHH------BBBB------HHHHHHHH---BBBBB------HHH--------BBBBBB--HHH------
3ey9 KAPIVHALRGKEHVEYDNPYDVGMTGLIGFSSGFHTMMNADTLVLLGTQFPYRAF---YPTDAKIIQIDINPASIGAHSK 303
2dji KAPVITTGKNFETFEWDFEALTGSTYRVGWKPANETILEADTVLFAGSNFPFSEVEGTFRNVDNFIQIDIDPAMLGKRHH 310
1y9d KIPLMSTYPAKGIVADRYPAYLGSANRAAQKPANEALAQADVVLFVGNNYPFAEVSKAFKNTRYFLQIDIDPAKLGKRHK 317

     --BBBB--------------------------------------------------HHHHHHHHHHH-----BBBB---HH
3ey9 VDMALVGDIKSTLRALLPLVEEKADRKFLDKALEDYRDARKGLDDLAKPSEKAIHPQYLAQQISHFAADDAIFTCDVGTP 383
2dji ADVAILGDAALAIDEILNKVDAVEESAWWTANLKNIANWREYINMLETKEEGDLQFYQVYNAINNHADEDAIYSIDVGNS 390
1y9d TDIAVLADAQKTLAAILAQVSERESTPWWQANLANVKNWRAYLASLEDKQEGPLQAYQVLRAVNKIAEPDAIYSIDVGDI 397

     HHHHHHH-------BBBB---------HHHHHHHHHHH-----BBBBBBHHHHHH-HHHHHHHHH-----BBBBBB----
3ey9 TVWAARYLKMNGKRRLLGSFNHGSMANAMPQALGAQATEPERQVVAMCGDGGFSMLMGDFLSVVQMKLPVKIVVFNNSVL 463
2dji TQTSIRHLHMTPKNMWRTSPLFATMGIAIPGGLGAKNTYPDRQVWNIIGDGAFSMTYPDVVTNVRYNMPVINVVFSNTEY 470
1y9d NLNANRHLKLTPSNRHITSNLFATMGVGIPGAIAAKLNYPERQVFNLAGDGGASMTMQDLATQVQYHLPVINVVFTNCQY 477

     -------------------------HHHHHHH--BBBBB--HHHHHHHHHHHHH-------BBBB---------------
3ey9 GFVAMEMK-AGGYLTDGTELHDTNFARIAEACGITGIRVEKASEVDEALQRAFSID--G-PVLVDVVVAKEELAIPPQIK 539
2dji AFIKNKYEDTNKNLF-GVDFTDVDYAKIAEAQGAKGFTVSRIEDMDRVMAEAVAANKAGHTVVIDCKITQDRPIPVETLK 549
1y9d GFIKDEQEDTNQNDFIGVEFNDIDFSKIADGVHMQAFRVNKIEQLPDVFEQAKAIAQHE-PVLIDAVITGDRPLPAEKLR 556

     -----------------------------------------------
3ey9 LEQA--------------KGFSLYMLRAIISGRGDEVIELAKTNWLR 572
2dji LDSKLYSEDEIKAYKERYEAANLVPFREYLEAEGLE------SKYIK 590
1y9d LDSAMSSAADIEAFKQRYEAQDLQPLSTYLKQFGLDDLQHQIGQGGF 603
```

**Supplemental Figure S1J:** Multiple sequence alignment with example secondary structure for the transketolase (TK) group of ThDP enzymes. The PDB ID used in this figure is given. The three state secondary structure assignment (H=helix, B=strand) for the repeat regions of PDB 4kxu is shown on top. The identified repeats for each domain are colored red, yellow, green, cyan, blue, and magenta respectively. The conserved residues in the functional domains (repeat 3 position 10 alanine, repeat 5 positions 5, 6) are indicated with bold underlined yellow text where they could be identified from sequence (and red where they could only be identified in the structures). Residue numbers are shown at the end of each line.

```
     ---------------------------------------------------------------------HHHHH--HHHH
4kxu -----------------MESYHKPDQQKLQALKDT---------------ANRLRISSIQATTAAGSGHPTSCCSAAEIM 48
3rim ---------MTTLEEISALTRPRHPDYWTEIDSAA---------------VDTIRVLAADAVQKVGNGHPGTAMSLAPLA 56
3m34 -----------------------SNAMNIQILQEQ---------------ANTLRFLSADMVQKANSGHPGAPLGLADIL 42
3m49 --MHHHHHHSSGVDLGTENLYFQSNAMSHSIEQLS---------------INTIRTLSIDAIEKANSGHPGMPMGAAPMA 63
4c7v --MAHHHHHHSSGLEVLF------QGPYDQVDQLG---------------VNTLRTLSIDAIQRANSGHPGLPMGAAPMA 57
2r5n ------------------------MSSRKEL-------------------ANAIRALSMDAVQKAKSGHPGAPMGMADIA 37
4xeu --MAHHHHHHM-------------PSRRER--------------------ANAIRALSMDAVQKANSGHPGAPMGMADIA 45
3uk1 --MAHHHHHHMGTLEAQTQGPGSMPPVPRFLDSFSGLDMTTSSPASTTLMANAIRALAMDAVQQANSGHPGMPMGMAEIG 78
5vrb --MAHHHHHHMSQL------------------------------------ANVIRFLSADAVQKANSGHPGAPMGMAEMA 42
1r9j -------------------------RHMASIEKV----------------ANCIRCLAADIVQGGKSGHPGTPMGMAPMS 39
1ay0 ------------------------MTQFTDIDKLA---------------VSTIRILAVDTVSKANSGHPGAPLGMAPAA 41
5hje MGSSHHHHHHSSGLVPR-------GSHMSSVDQKA---------------ISTIRLLAVDAVAAANSGHPGAPLGLAPAA 58
1itz ----------------GAVETLQGKAATGELLEKS---------------VNTIRFLAIDAVEKANSGHPGLPMGCAPMG 49
5nd6 --MHHHHHHHMAAQAAPAAAKAAAPSISRDEVEKC---------------INAIRFLAIDAINKSKSGHPGMPMGCAPMG 63

     HHHHH---B-------------BBBB--HHHHHHHHHHHHH-------HHHHHH--------------------------
4kxu AVLFFHTMRYKSQDP-RNPHNDRFVLSKGHAAPILYAVWAEAGF--LAEAELLNLRKISSDLDGHPV-PKQAFTDVATGS 124
3rim YTLFQRTMRHDPSDT-HWLGRDRFVLSAGHSSLTLYIQLYLGGFG-LELSDIESLRTWGSKTPGHPEFRHTPGVEITTGP 134
3m34 SVL-SYHLKHNPKNP-TWLNRDRLVFSGGHASALLYSFLHLSGYD-LSLEDLKNFRQLHSKTPGHPE-ISTLGVEIATGP 118
3m49 YTLWTQFMKHNPNNP-TWFNRDRFVLSAGHGSMLLYSLLHLSGYD-VTMDDLKNFRQWGSKTPGHPEYGHTAGVDATTGP 141
4c7v YVLWTRHLKINPKTHMNWVNRDRFVLSAGHGSALLYSLAHLAGYD-VSMDDLKNFREWKSNTPGHPEYGCTDGVEATTGP 136
2r5n EVLWRDFLKHNPQNP-SWADRDRFVLSNGHGSMLIYSLLHLTGYD-LPMEELKNFRQLHSKTPGHPEVGYTAGVETTTGP 115
4xeu EVLWRDYMQHNPSNP-QWANRDRFVLSNGHGSMLIYSLLHLTGYD-LGIEDLKNFRQLNSRTPGHPEYGYTAGVETTTGP 123
3uk1 VALWSRHLKHNPTNP-HWADRDRFVLSNGHGSMLLYSLLHLTGYD-LPIEELKNFRQLHSKTPGHPEYGITPGVETTTGP 156
5vrb ETLWTKFLNHNPANP-KFYNRDRFVLSNGHASMLLYSLLHLTGYN-LSIEDLKNFRQLHSKTPGHPEYGYTDGVETTTGP 120
1r9j AVLWTEVMKYNSQDP-DWVDRDRFVMSNGHGCALQYALLHMAGYN-LTMDDLKGFRQDGSRTPGHPERFVTPGVEVTTGP 117
1ay0 HVLWSQ-MRMNPTNP-DWINRDRFVLSNGHAVALLYSMLHLTGYD-LSIEDLKQFRQLGSRTPGHPE-FELPGVEVTTGP 117
5hje HAVFKK-MRFNPKDT-KWINRDRFVLSNGHACALLYSMLVLYGYD-LTVEDLKKFRQLGSKTPGHPENTDVPGAEVTTGP 135
1itz HVLYDEVMRYNPKNP-YWFNRDRFVLSAGHGCMLQYALLHLAGYDSVKEEDLKQFRQWGSRTPGHPENFETPGVEVTTGP 128
5nd6 YVLWNEVMKYNPKNP-DFFNRDRFVLSAGHGSMFQYSMMHLTGYDSVPLDQIKQFRQWNSLTPGHPENFVTPGVEVTTGP 142

     ---HHHHHHHHHHHHHH--------------------BBBBBB-HHHH-HHHHHHHHHHHH-----BBBBBBB--B----
4kxu LGQGLGAACGMAYTGKY----FD---KASY------RVYCLLGDGELSEGSVWEAMAFASIYKLDNLVAILDINRLGQSD 191
3rim LGQGLASAVGMAMASRYERGLFDPDAEPGA-SPFDHYIYVIASDGDIEEGVTSEASSLAAVQQLGNLIVFYDRNQISIED 213
3m34 LGQGVANAVGFAMAAKKAQNLLG---SD----LIDHKIYCLCGDGDLQEGISYEACSLAGLHKLDNFILIYDSNNISIEG 191
3m49 LGQGIATAVGMAMAERHLAAKYN---RDAY-NIVDHYTYAICGDGDLMEGVSAEASSLAAHLQLGRLVVLYDSNDISLDG 217
4c7v LGQGISMAVGMAMAEAHLGKKFN---REGY-PVMDHYTYALIGDGDLMEGVASEAASLAGHLKLGKLIALYDSNGISLDG 212
2r5n LGQGIANAVGMAIAEKTLAAQFN---RPGH-DIVDHYTYAFMGDGCMMEGISHEVCSLAGTLKLGKLIAFYDDNGISIDG 191
4xeu LGQGIANAVGMALAEKVLAAQFN---RDGH-AVVDHYTYAFLGDGCMMEGISHEVASLAGTLRLNKLIAFYDDNGISIDG 199
3uk1 LGQGLANAVGMALGEALLAAEFN---RDDA-KIVDHHTYVFLGDGCLMEGISHEACSLAGTLKLNKLIALYDDNGISIDG 232
5vrb LGQGIANAVGMALAEKILAAEFN---KDGL-NIVDHYTYVFMGDGCLMEGVSHEACSLAGTLGLGKLIVLYDDNNISIDG 196
1r9j LGQGIANAVGLAIAEAHLAATFN---RPGY-NIVDHYTYVYCGDGCLMEGVCQEALSLAGHLALEKLIVIYDSNYISIDG 193
1ay0 LGQGISNAVGMAMAQANLAATYN---KPGF-TLSDNYTYVFLGDGCLQEGISSEASSLAGHLKLGNLIAIYDDNKITIDG 193
5hje LGQGICNGVGIALAQAQFAATYN---KPDF-PISDSYTYVFLGDGCLMEGVSSEASSLAGHLQLGNLIAFWDDNKISIDG 211
1itz LGQGIANAVGLALAEKHLAARFN---KPDS-EIVDHYTYVILGDGCQMEGIANEACSLAGHWGLGKLIAFYDDNHISIDG 204
5nd6 LGQGICNAVGLAVAEAHLAARFN---KPDVKPIVDHYTYCILGDGCMMEGISNEACSLAGHWGLGKLIALYDDNKISIDG 219
```

```
     B------HHHHHHHHHH---BBB-BBB----HHHHHHHH----------BBBBBB-------------------------
4kxu PAPLQHQMDIYQKRCEAFGWHAI-IVDGH-SVEELCKAFGQAKHQ---PTAIIAKTFKGRGITGVEDKESWHGKPLPKNM 266
3rim DTNIALCEDT-AARYRAYGWHVQEVEGGE-NVVGIEEAIANAQAVTDRPSFIALRTVIGYPAPNLMDTGKAHGAALGDDE 291
3m34 DVGLAFNENV-KMRFEAQGFEVLSI-NGH-DYEEINKALEQAKKST-KPCLIIAKTTIAKGAGELEGSHKSHGAPLGEEV 267
3m49 DLNRSFSESV-EDRYKAYGWQVIRVEDGN-DIEAIAKAIEEAKADEKRPTLIEVRTTIGFGSPNKSGKSASHGSPLGVEE 295
4c7v KTSASFTENV-GARFEAYGWQYILVEDGF-NLEEIDKAIVQAKAESDKPTIIEIKTTIGYGSEN-QGTHKVHGSPLGEEG 289
2r5n HVEGWFTDDT-AMRFEAYGWHVIRDIDGH-DAASIKRAVEEARAVTDKPSLLMCKTIIGFGSPNKAGTHDSHGAPLGDAE 269
4xeu EVHGWFTDDT-PKRFEAYGWQVIRNVDGH-DADEIKTAIDTAR-KSDQPTLICCKTVIGFGSPNKQGKEECHGAPLGADE 276
3uk1 DVVNWFHDDT-PKRFEAYGWNVIPNVNGH-DVDAIDAAIAKAK-RSDKPSLICCKTRIGNGAATKAGGHDVHGAPLGADE 309
5vrb KVDGWFTENI-PQRFESYGWHVVPNVNGH-DTAAIQTAIEAARAETGKPSIICCKTLIGKGSANKEGSHKTHGAPLGADE 274
1r9j STSLSFTEQC-HQKYVAMGFHVIEVKNGDTDYEGLRKALAEAKATKGKPKMIVQTTTIGFGSSK-QGTEKVHGAPLGEED 271
1ay0 ATSISFDEDV-AKRYEAYGWEVLYVENGNEDLAGIAKAIAQAKLSKDKPTLIKMTTTIGYGSLH-AGSHSVAGAPLKADD 271
5hje STEVAFTEDV-IARYKSYGWHIVEVSDADTDITAIAAAIDEAKKVTNKPTLVRLTTTIGFGSLA-QGTHGVHGAPLKADD 289
1itz DTEIAFTEDV-STRFEALGWHTIWVKNGNTGYDDIRAAIKEAKAVTDKPTLIKVTTTIGFGSPNKANSYSVHGSALGAKE 283
5nd6 HTDISFTEDV-AKRYEALGWHVIHVINGNTDVDGLRAAIAQAKAVKDKPTLIKVSTLIGYGSPNKADSHDVHGAPLGPDE 298

     --------------------------------------------------------------------------------
4kxu AEQ--------------IIQEIYSQIQSKKKI----------------LATPPQEDAPSVDIANIRMPS-----LPSY-- 309
3rim VAAVKKIVGFDPDKTFQVREDVLTHTRGLVAR-GKQAHERWQLEFDAWARREPERKALLDRLLAQKLPDGWDADLPHW-- 368
3m34 IKKAKEQAGFDPNISFHIPQASKIRFESAVEL-GDLEEAKWKDKLEKSAKK------ELLERLLN--PDFNKIAYPDF-- 336
3m49 TKLTKEAYAWTAEQDFHVAEEVYENFRKTVQDVGETAQAEWNTMLGEYAQAYPELANELQAAMNGLLPEGWEQNLPTY-- 373
4c7v VAHAKEVYNW-NYPPFTVPEEVSQRFKECLQDKGVKAENKWNEMFEAYKKEYSDLAQKFSDGFSNKVPNTLGDILPQY-- 366
2r5n IALTREQLGW-KYAPFEIPSEIYAQWDAKEA--GQAKESAWNEKFAAYAKAYPQEAAEFTRRMKGEMPSDFDAKAKEFIA 346
4xeu IAATRAALGW-EHAPFEIPAQIYAEWDAKET--GAAQEAEWNKRFAAYQAAHPELAAELLRRLKGELPADFAEKAAAYVA 353
3uk1 IAKTREALGW-TWAPFVIPQEVYAAWDAKEA--GKRSEDDWNAAFAQYRAKYPAEAAEFERRMAGTLPADWAAKAAAIVA 386
5vrb IEATRKHLGW-AYPAFEIPQEIYDAWNAKEK--GAKLEAGWNELFAQYQAKYPAEAAEFVRRMDKKLPENFDEYVQTALK 351
1r9j IANIKAKFGRDPQKKYDVDDDVRAVFRMHIDK-CSAEQKAWEELLAKYTAAFPAEGAAFVAQMRGELPSGWEAKLPT--- 347
1ay0 VKQLKSKFGFNPDKSFVVPQEVYDHYQKTILKPGVEANNKWNKLFSEYQKKFPELGAELARRLSGQLPANWESKLPTY-- 349
5hje IKQLKTKWGFNPEESFAVPAEVTASYNEHVAE-NQKIQQQWNELFAAYKQKYPELGAELQRRLDGKLPENWDKALPVY-- 366
1itz VEATRQNLGW-PYDTFFVPEDVKSHWSRHTPE-GAALEADWNAKFAEYEKKYADDAATLKSIITGELPTGWVDALPKY-- 359
5nd6 TAATRKNLNW-PYGEFEVPQDVYDVFRGAIKR-GAEEEANWHKACAEYKAKYPKEWAEFEALTSCKLPENWEAALPHF-- 374

     ---------BHHHHHHHHHHHHHHH---BBBBB---HHHH--HHH----HHH-H-------------HHBBB----HHHH
4kxu ---KVGDKIATRKAYGQALAKLGHASDRIIALDGDTKNSTFSEIF----KKEHP-------------DRFIECYIAEQNM 369
3rim --EPGSKALATRAASGAVLSALGPKLPELWGGSADLAGSNNTTIKGA--DSFGPPSISTKEYTAHWYGRTLHFGVREHAM 444
3m34 ----KGKDLATRDSNGEILNVLAKNLEGFLGGSADLGPSNKTELHSM--GDFVE-------------GKNIHFGIREHAM 397
3m49 ---ELGSKAATRNSSGAVINAIAESVPSFFGGSADLAGSNKTYMNNE--KDFTR-----DDYS----GKNIWYGVREFAM 439
4c7v ---GEDDSIATRAASQKAINALAKEVSSLWGGAADLASSNKTVIAGE--GDFQP-----ESYE----GRNIWFGVREFGM 432
2r5n KLQANPAKIASRKASQNAIEAFGPLLPEFLGGSADLAPSNLTLWSGS--KAINE------DAA----GNYIHYGVREFGM 414
4xeu DVANKGETIASRKASQNALNAFGPLLPELLGGSADLAGSNLTLWKGC--KGVSA-----DDAA----GNYVFYGVREFGM 422
3uk1 GANERGETVATRKASQQTIEGLAAVLPELLGGSADLTGSNLTNWKAS--KAVRANADGPGVQW----GNHINYGVREFGM 460
5vrb EVCAKAETVATRKASQNSIEILAKELPELVGGSADLTPSNLTDWSNS--VSVTR------DKG----GNYIHYGVREFGM 419
1r9j ----NSSAIATRKASENCLAVLFPAIPALMGGSADLTPSNLTRPASANLVDFSS-----SSKE----GRYIRFGVREHAM 414
1ay0 --TAKDSAVATRKLSETVLEDVYNQLPELIGGSADLTPSNLTRWKEA--LDFQPPSSGSGNYS----GRYIRYGIREHAM 421
5hje --TPADAAVATRKLSEIVLSKIIPEVPEIIGGSADLTPSNLTKAKGT--VDFQPAATGLGDYS----GRYIRYGVREHAM 438
1itz --TPESPGDATRNLSQQCLNALANVVPGLIGGSADLASSNMTLLKMF--GDFQK-----DTAE----ERNVRFGVREHGM 426
5nd6 --KPEDKGLATRQHSQTMINALAPALPGLIGGSADLAPSNLTLMKIS--GDFQK-----GSYA----ERNLRFGVREHAM 441

     HHHHHHHHHHH---BBBBBBBHHHHHHHHHHHHHHHH----BBBBBB-B-HHH----HHH-B--HHHHHH-----BBB--
4kxu VSIAVGCATRN-RTVPFCSTFAAFFTRAFDQIRMAAISESNINLCGSHCGVSIGEDGPSQMALEDLAMFRSVPTSTVFYP 448
3rim GAILSGIVLHG-PTRAYGGTFLQFSDYMRPAVRLAALMDIDTIYVWTHDSIGLGEDGPTHQPIEHLSALRAIPRLSVVRP 523
3m34 AAINNAFARYG-IFLPFSATFFIFSEYLKPAARIAALMKIKHFFIFTHDSIGVGEDGPTHQPIEQLSTFRAMPNFLTFRP 476
3m49 GAAMNGIALHG-GLKTYGGTFFVFSDYLRPAIRLAALMQLPVTYVFTHDSIAVGEDGPTHEPIEQLAALRAMPNVSVIRP 518
4c7v ACAMNGIMLHG-GTRIFGSTFFVFSDYLKAAIRLSAIQKLPVIYVLTHDSVAVGKDGPTHEPIEQLASLRTIPNVQVFRP 511
2r5n TAIANGISLHG-GFLPYTSTFLMFVEYARNAVRMAALMKQRQVMVYTHDSIGLGEDGPTHQPVEQVASLRVTPNMSTWRP 493
4xeu SAIMNGVALHG-GFIPYGATFLIFMEYARNAVRMSALMKQRVLYVFTHDSIGLGEDGPTHQPIEQLASLRLTPNLDTWRP 501
3uk1 SAAINGLVLHG-GYKPFGGTFLTFSDYSRNALRVAALMKVPSIFVFTHDSIGLGEDGPTHQSVEHVASLRLIPNLDVWRP 539
5vrb GAIMNGLVLHG-GVKPFGATFLMFSEYERNALRMAALMKINPVFVFTHDSIGLGEDGPTHQPIEQTATLRLIPNMDVWRP 498
1r9j CAILNGLDAHD-GIIPFGGTFLNFIGYALGAVRLAAISHHRVIYVATHDSIGVGEDGPTHQPVELVAALRAMPNLQVIRP 493
1ay0 GAIMNGISAFGANYKPYGGTFLNFVSYAAGAVRLSALSGHPVIWVATHDSIGVGEDGPTHQPIETLAHFRSLPNIQVWRP 501
5hje GAIMNGIAAFGANYKNYGGTFLNFVSYAAGAVRLSALSEFPITWVATHDSIGLGEDGPTHQPIETLAHFRATPNISVWRP 518
1itz GAICNGIALHSPGFVPYCATFFVFTDYMRGAMRISALSEAGVIYVMTHDSIGLGEDGPTHQPIEHLVSFRAMPNILMLRP 506
5nd6 GAICNGIALHKSGLIPYCATFYIFTDYMRNAMRMSALSEAGVVYVMTHDSIGLGEDGPTHQPIEHLASFRAMPDMLMIRP 521
```

```
      --HHHHHHHHHHHH----------BBBBB----BBB--------------BBBB-----------------HHHHHHH
4kxu  SDGVATEKAVELAANTKG------ICFIRTSRPENAIIYNNN-EDFQVGQAKVVLKSK-----DDQVTVIGAGVTLHEAL 516
3rim  ADANETAYAWRTILARRNGSGPVGLILTRQGVPVLDGTDA---EGVARGGYVLSDAGGLQPGEEPDVILIATGSEVQLAV 600
3m34  ADGVENVKAWQIALNAD---IPSAFVLSRQKLKALNEPVF---GDVKNGAYLLKESK------EAKFTLLASGSEVWLCL 544
3m49  ADGNESVAAWRLALESTN--KPTALVLTRQDLPTLEGAKDDTYEKVAKGAYVVSASKK---E-TADVILLATGSEVSLAV 592
4c7v  ADGNETSAAWKVALETLD--KPTILVLSRQNLDTLPISKEKVFDGVEKGGYVVQGAEN-----EADGILIATGSEVGLAL 584
2r5n  CDQVESAVAWKYGVERQD--GPTALILSRQNLAQQERTEEQL-ANIARGGYVLKDCAG-----QPELIFIATGSEVELAV 565
4xeu  ADAVESAVAWKHAIERAD--GPSALIFSRQNLPHQARDVAQV-ADIARGGYVLKDCEG-----EPELILIATGSEVGLAV 573
3uk1  ADTVETAVAWTYAVAHQH---PSCLIFSRQNLAFNARTDAQL-ANVEKGGYVLRDWDEEIV--ARKIILIATGSEVELAM 613
5vrb  CDTAESLVAWAEAAKAED--HPSCLIFSRQNLKFQARSEQQL-NDIKRGAYVISEAQG-----NAQAVIIATGSEVGLAV 570
1r9j  SDQTETSGAWAVALSSIH--TPTVLCLSRQNTEPQSGSSI---EGVRHGAYSVVDVP------DLQLVIVASGSEVSLAV 562
1ay0  ADGNEVSAAYKNSLESKH--TPSIIALSRQNLPQLEGSSI---ESASKGGYVLQDVA------NPDIILVATGSEVSLSV 570
5hje  ADGNETSAAYKSAIESTH--TPHILALTRQNLPQLEGSSI---EKASKGGYTLVQQD------KADIIIVATGSEVSLAV 587
1itz  ADGNETAGAYKVAVLNRK--RPSILALSRQKLPHLPGTSI---EGVEKGGYTISDNST---GNKPDLIVMGTGSELEIAA 578
5nd6  AGGNETAGAYKVAIANRK--RPTTIALSRQNMPNIPNCSV---EGVAKGAYTIHDTKA---GVKPDVILMGTGSELELAT 593

      HHHHHHH----BBBBBB--BBB---HHHHHHHHHH---BBBBBBBBB----HHHHHHHHH------BB----------BB
4kxu  AAAELLKKEKINIRVLDPFTIKPLDRKLILDSARATKGRILTVEDHYYEGGIGEAVSSAVVGEPGITV----------TH 586
3rim  AAQTLLADNDILARVVSMPCLEWFEAQ----------------PYEYRDAVLPPTVSARVAVEAGVAQCWHQLVGDTGEI 664
3m34  ESANELEKQGFACNVVSMPCFELFEKQ----------------DKAYQERLLKGEV---IGVEAAHSNELYKF---CHKV 602
3m49  EAQKALAVDGVDASVVSMPSMDRFEAQ----------------TAEYKESVLPKAVTKRFAIEMGATFGWHRYVGLEGDV 656
4c7v  KAKEELQKKGKDVIVVSLPSWERFEAQ----------------SEEYKNTVIPPELKKRMTIEAGTTYGWAKYAGDHGVM 648
2r5n  AAYEKLTAEGVKARVVSMPSTDAFDKQ----------------DAAYRESVLPKAVTARVAVEAGIADYWYKYVGLNGAI 629
4xeu  QAYDKLSEQGRKVRVVSMPCTSVYEQQ----------------DESYKQSVLPVEVGARIAIEAAHADYWYKYVGLDGRI 637
3uk1  KAVEPLAQQGIAARVVSMPSSDVFDRQ----------------DAEYRERVLPHGVRR-VAIEAGVTDFWRKYVGLEGGV 676
5vrb  EAQKVLAGQGIAVRVVSMPSTSVFDRQ----------------DAAYQAAVLPEGLPR-IAVEAGHTNGWYKYVGLNGAV 633
1r9j  DAAKALSGE-LRVRVVSMPCQELFDAQ----------------PDTYRQAVLPAGVPV-VSVEAYVSFGWEKY---SHAH 621
1ay0  EAAKTLAAKNIKARVVSLPDFFTFDKQ----------------PLEYRLSVLPDNVPI-MSVEVLATTCWGKY---AHQS 630
5hje  DALKVLEGQGIKAGVVSLPDQLTFDKQ----------------SEEYKLSVLPDGVPI-LSVEVMSTFGWSKY---SHQQ 647
1itz  KAADELRKEGKTVRVVSFVSWELFDEQ----------------SDEYKESVLPAAVTARISIEAGSTLGWQKYVGAQGKA 642
5nd6  AAAGILEKEGKNVRVVSFPCWELFEEQ----------------SAEYKESVLPSDVTARVSVEAATSFGWAKYIGLKGKH 657

      B-----------------------------------------------------
4kxu  LAVNRVPRSGKPAELLKMFGIDRDAIAQAVRGLITKALVPRGSLEHHHHHH--- 637
3rim  VSIEHYGESADHKTLFREYGFTAEAVAAAAERALDN------------------ 700
3m34  YGIESFGESGKDKDVFERFGFSVSKLVNFILSK--------------------- 635
3m49  LGIDTFGASAPGEKIMEEYGFTVENVVRKVKEML-------------------- 690
4c7v  IGIDEFGMSAPSDIVLRELGMSVENIVDKYLEK--------------------- 681
2r5n  VGMTTFGESAPAELLFEEFGFTVDNVVAKAKELLHHHHHH-------------- 669
4xeu  IGMTSFGESAPAPALFEHFGFTLDNVLAVAEELLED------------------ 673
3uk1  VGIDTFGESAPAGVLFKHFGFTVEHVIETAKAVLA------------------- 711
5vrb  VGINRFGESAPADLLFKAFGFTVDNVVDTVKSVL-------------------- 667
1r9j  VGMSGFGASAPAGVLYKKFGITVEEVVRTGRELAKRFP--DGTAPLKNSSFSKM 673
1ay0  FGIDRFGASGKAPEVFKFFGFTPEGVAERAQKTIAFYKGDKLISPLKKAF---- 680
5hje  FGLNRFGASGKAPEIFKLFEFTPEGVAERAAKTVAFYKGKDVVSPLRSAF---- 697
1itz  IGIDKFGASAPAGTIYKEYGITVESIIAAAKSF--------------------- 675
5nd6  VGIDTFGASAPAPTLYEKFGITVNHVVEAAKATLQH------------------ 693
```

**Supplemental Figure S2:** Amino acid conservation bias in the repeats. Sequences in the TEED database that were homologous to the known crystal structures were used to identify biases in conservation relative to overall sequence identity and conservation at a given position as determined by the Simpson metric. Data is given as relative enrichment compared to the normal amino acid distribution as calculated from the PDB. A) Simpson value = 0.3 B) Simpson value = 0.4, C) Simpson value = 0.5 D) Simpson value = 0.6, E) Simpson value = 0.7 F) Simpson value = 0.8 G) Amino acid distribution (total) within the repeats.

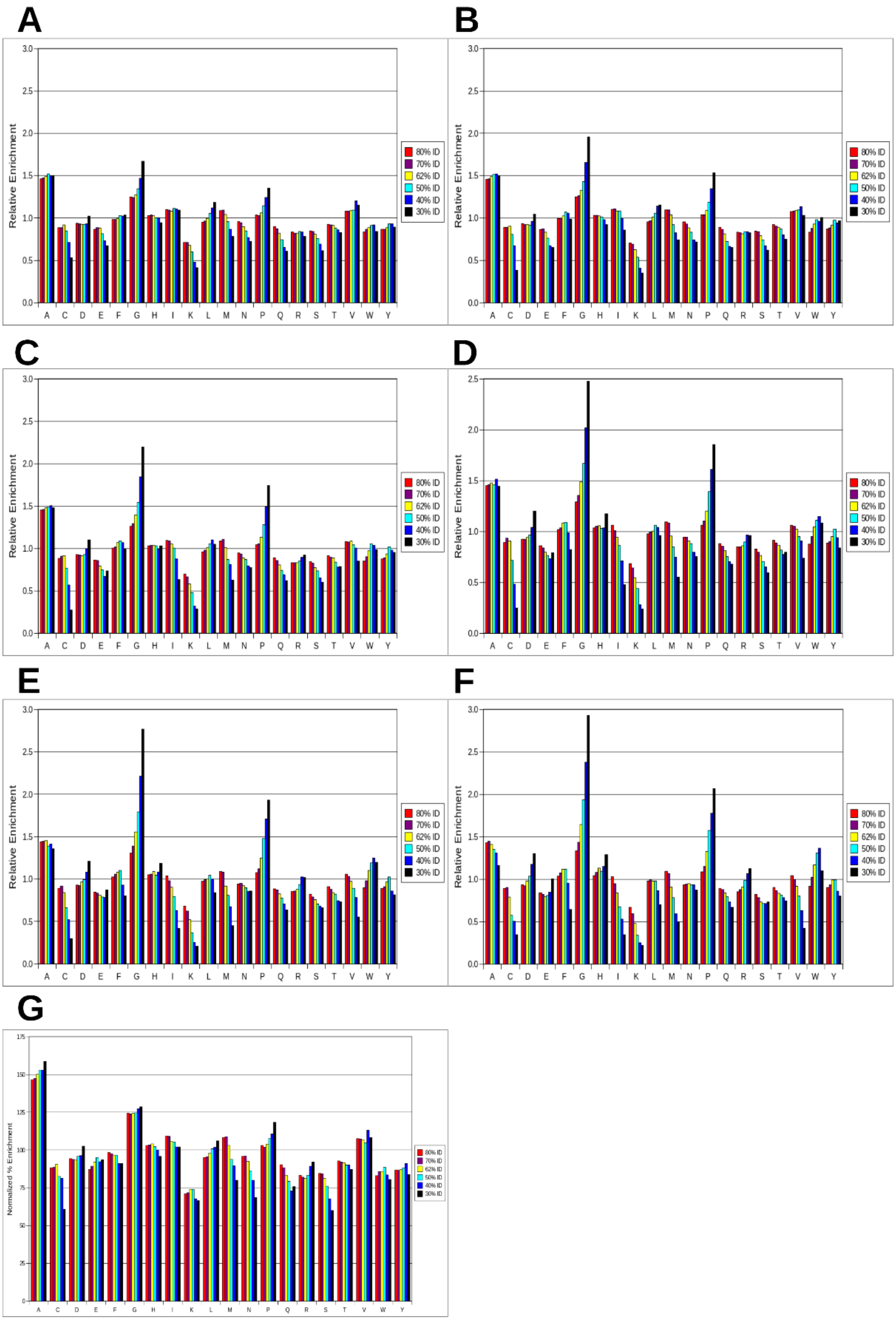

**Supplemental Figure S3:** Self-conserved sequence features within the functional repeats of the transketolases using a combination of automatic and manual alignment. The PDB ID and the functional domain in which the repeat is located are identified. Green highlight indicates overall conserved features identified either from sequence alignments or of the starting (G/A){X(1,2)}(G/A) motif. Yellow highlight indicates homologous substitutions while cyan indicates generally conserved features (such as a hydrophobic residue). Grey highlight indicates a self-conserved residue not included in the previous categories. A consensus sequence is given at the bottom of each repeat in which positions of conserved identity are indicated by a capital letter, g = Gly or Ala, ϕ = hydrophobic residue, i = Val, Ile or Leu.

**REPT 1**

```
4KXU_PP            GHPTSCCSAAEIMAVLF-FHTMRYKSQDPRNPHN----DRFVLSK
4KXU_PYR           ATRKAYGQALAKLGHAS-DRIIALD-----------------GD-
3RIM_PP            GHPGTAMSLAPLAYTLFQRTMRH----DPSDTHWLG-RDRFVLS-
3RIM_PYR           AL-ATRAASGAVLSALG-----------PKLPELWG------GS-
3M34_PP            GHPGAPLGLADILSVLSYH-LKH----NPKNPTWLN-RDRLVFS-
3M34_PYR           GKDLATRDSNGEILNVL---AKN---LEGFLG----------GS-
3M49_PP            GHPGMPMGAAPMAYTLWTQFMKH----NPNNPTWFN-RDRFVLS-
3M49_PYR           GSKAATRNSSGAVINAIAE--------SVPSFFG--------GS-
4C7V_PP            GHPGLPMGAAPMAYVLWTRHLKI----NPKTHMNWVNRDRFVLS-
4C7V_PYR           ATRAASQKAINALAKEVSSLWG--------------------GA-
2R5N_PP            GHPGAPMGMADIAEVLWRDFLKH----NPQNPSWAD-RDRFVLSN
2R5N_PYR           ANPAKIASRKASQNAIE--------AFGPLLPEFLG------GS-
4XEU_PP            GHPGAPMGMADIAEVLW----RDYMQHNPSNPQWAN-RDRFVLSN
4XEU_PYR           ANKGETIASRKASQNAL-------NAFGPLLPELL----------
3UK1_PP            GHPGMPMGMAEIGVALW----SRHLKHNPTNPHWAD-RDRFVLSN
3UK1_PYR           AG-ANERGETVATRKASQQTIEGLAAVLPELL------------
5VRB_PP            GHPGAPMGMAEMAETLW----TKFLNHNPANPKFYN-RDRFVLSN
5VRB_PYR           AK-AETVATRKASQNS-------IEILAKELPELV----------
1R9J_PP            GHPGTPMGMAPMSAVLW----TEVMKYNSQDPDWVD-RDRFVMSN
1R9J_PYR           AI-ATRKASENCLAVL------------FPAIPALMG---------
1AY0_PP            GHPGAPLGMAPAAHVLW-----SQMRMNPTNPDWIN-RDRFVLSN
1AY0_PYR           AV-ATRKLSETVLEDV------------YNQLPELIG---------
5HJE_PP            GHPGAPLGLAPAAHAVF-----KKMRFNPKDTKWIN-RDRFVLSN
5HJE_PYR           AD-AAVATRKLSEIVL--------SKIIPEVPEIIG----GSADL
1ITZ_PP            GHPGLPMGCAPMGHVLY----DEVMRYNPKNPYWFN-RDRFVLSA
1ITZ_PYR           GD-ATRNLSQQCLNAL--------ANVVPGLIG---------GS-
5ND6_PP            GHPGMPMGCAPMGYVLW----NEVMKYNPKNPDFFN-RDRFVLS-
5ND6_PYR           GL-ATRQHSQTMINAL------------APALPGLIG------GS-

CONSENSUS:         g--g-------φφ--L------------P--P-----------S-
```

**REPT 2**

```
4KXU_PP        GHAAPILYAVWAEAGFLAEAELLNL----RKISSDLDGHPVPKQAFTDVAT
4KXU_PYR       TKNSTFSEIFKKEHPDRFIECYIAE
3RIM_PP        AGHSSLTLYIQLYLGGF-GLELSDIESLRTWGSKTPGHPEFRHTPGVEITT
3RIM_PYR       ADLAGSNNTTIKGADSF-------GPPSISTKEYTAHWYGRTLHFGVREH
3M34_PP        GGHASALLYSFLHLSG--YDLSLEDLKNFRQLHSKTPGHPEISTLGVEIAT
3M34_PYR       ADLGPSNKTELHSMG--------------------DFVEGKNIHFGIREH
3M49_PP        AGHGSMLLYSLLHLSG-YDVTMDDLKNFRQWGSKTPGHPEYGHTAGVDATT
3M49_PYR       ADLAGSNKTYMNNE----------------KDFTRDDYSGKNIWYGVREF
4C7V_PP        AGHGSALLYSLAHLAG-YDVSMDDLKNFREWKSNTPGHPEYGCTDGVEATT
4C7V_PYR       ADLASSNKTVIAGE----------------GDFQPESYEGRNIWFGVREF
2R5N_PP        GHGSMLIY-SLLHLTGY-DLPMEELKNFRQLHSKTPGHPEVGYTAGVETTT
2R5N_PYR       ADLAP----SNLTLW-------------SGSKAINEDAAGNYIHYGVREF
4XEU_PP        GHGSMLIY-SLLHLTGY-DLGIEDLKNFRQLNSRTPGHPEYGYTAGVETTT
4XEU_PYR       GGSADLAG-SNLTLW------------KGCKGVSADDAAGNYVFYGVREF
3UK1_PP        GHGSMLLY-SLLHLTGY-DLPIEELKNFRQLHSKTPGHPEYGITPGVETTT
3UK1_PYR       GGSADLTG-SNLTNWKA-------SKAVRANADGPGVQWGNHINYGVREF
5VRB_PP        GHASMLLY-SLLHLTGY-NLSIEDLKNFRQLHSKTPGHPEYGYTDGVETTT
5VRB_PYR       GGSADLTP-SNLTDWSN-------------SVSVTRDKGGNYIHYGVREF
1R9J_PP        GHGCALQY-ALLHMAGY-NLTMDDLKGFRQDGSRTPGHPERFVTPGVEVTT
1R9J_PYR       GSADLTP--SNLTRPAS----------ANLVDFSSSSKEGRYIRFGVREH
1AY0_PP        GHAVALLY-SMLHLTGY--DLSIEDLKQFRQLGSRTPGHPEFELPGVEVTT
1AY0_PYR       GSADLTP--SNLTRWKE-------ALDFQPPSSGSGNYSGRYIRYGIREH
5HJE_PP        GHACALLY-SMLVLYGY-DLTVEDLKKFRQLGSKTPGHPENTDVPGAEVTT
5HJE_PYR       TP-------SNLTKAKG-------TVDFQPAATGLGDYSGRYIRYGVREH
1ITZ_PP        GHGCMLQY-ALLHLAGYDSVKEEDLKQFRQWGSRTPGHPENFETPGVEVTT
1ITZ_PYR       ADLASSNM-TLLKMFGD---------------FQKDTAEERNVRFGVREH
5ND6_PP        AGHGSMFQYSMMHLTGYDSVPLDQIKQFRQWNSLTPGHPENFVTPGVEVTT
5ND6_PYR       ADLAP----SNLTLMK------------ISGDFQKGSYAERNLRFGVREH

CONSENSUS:     g--g-L---SLL-L-------------------------------GV---
```

```
REPT_3
4KXU_PP      GSLGQGLGAACGMAYTGK---YFDKASYRVYCLL
4KXU_PYR     QNMVSIAVGCAT-RNRTVPFCSTF
3RIM_PP      GPLGQGLASAVGMAMASRYERGLFDPDAEPGASPFDHYIYVI
3RIM_PYR     AMGAILSGIVLH--GPTRAYGGTF
3M34_PP      GPLGQGVANAVGFAM-----AAKKAQNLLGSDLIDHKIYCLC
3M34_PYR     AMAAINNAFARY--GIFLPFSATF
3M49_PP      GPLGQGIATAVGMAMAERHLAAKYNRDAYNIVDHYTYAIC
3M49_PYR     AMGAAMNGIALH--GGLKTYGGTF
4C7V_PP      GPLGQGISMAVGMAMAEAHLGKKFNREGYPVMDHYTYALI
4C7V_PYR     GMACAMNGIMLH--GGTRIFGSTF
2R5N_PP      GPLGQGIANAVGMAIAEKTLAAQFNRPGHDIVDHYTYAFM
2R5N_PYR     GMTAIANGISLH--GGFLPYTSTFLMFVEY
4XEU_PP      GPLGQGIANAVGMALAEKVLAAQFNRDGHAVVDHYTYAFL
4XEU_PYR     GMSAIMNGVALH--GGFIPYGATFLIFMEY
3UK1_PP      GPLGQGLANAVGMALGEALLAAEFNRDDAKIVDHHTYVFL
3UK1_PYR     GMSAAINGLVLH--GGYKPFGGTF
5VRB_PP      GPLGQGIANAVGMALAEKILAAEFNKDGLNIVDHYTYVFM
5VRB_PYR     GMGAIMNGLVL--HGGVKPFGATF
1R9J_PP      GPLGQGIANAVGLAIAEAHLAATFNRPGYNIVDHYTYVYC
1R9J_PYR     AMCAILNGLD--AHDGIIPFGGTFLNFI
1AY0_PP      GPLGQGISNAVGMAMAQANLAATYNKPGFTLSDNYTYVFL
1AY0_PYR     AMGAIMNGIS--------AFGANYKPY
5HJE_PP      GPLGQGICNGVGIALAQAQFAATYNKPDFPISDSYTYVFL
5HJE_PYR     AMGAIMNGIAAF-GANYKNYGGTFLNFVSY
1ITZ_PP      GPLGQGIANAVGLALAEKHLAARFNKPDSEIVDHYTYVIL
1ITZ_PYR     GMGAICNGIALH-SPGFVPYCATFFVF
5ND6_PP      GPLGQGICNAVGLAVAEAHLAARFNKPDVKPIVDHYTYCIL
5ND6_PYR     AMGAICNGIALH-KSGLIPYCATF

CONSENSUS:   g--g---G-A----------AATF---
```

**REPT 4**

```
4KXU_PP            -G-DGELSEGSVWEAMAFASIYKLDNLVAILDINR
4KXU_PYR           -------AAFFTRAFDQIRMAAISESNINLCGSHC
3RIM_PP            -ASDGDIEEGVTSEASSLAAVQQLGNLIVFYDRNQ
3RIM_PYR           ----LQFSDYMRPAVRLAALMDIDTIYVWTHDSI
3M34_PP            -G-DGDLQEGISYEACSLAGLHKLDNFILIYDSNN
3M34_PYR           ----FIFSEYLKPAARIAALMKIKHFFIFTHDSI
3M49_PP            -G-DGDLMEGVSAEASSLAAHLQLGRLVVLYDSND
3M49_PYR           ----FVFSDYLRPAIRLAALMQLPVTYVFTHDSI
4C7V_PP            -G-DGDLMEGVASEAASLAGHLKLGKLIALYDSN
4C7V_PYR           ----FVFSDYLKAAIRLSAIQKLPVIYVLTHDSV
2R5N_PP            -G-DGCMMEGISHEVCSLAGTLKLGKLIAFYDDN
2R5N_PYR           ----------ARNAVRMAALMKQRQVMVYTHDSI
4XEU_PP            -G-DGCMMEGISHEVASLAGTLRLNKLIAFYDDN
4XEU_PYR           ----------ARNAVRMSALMKQRVLYVFTHDSI
3UK1_PP            -G-DGCLMEGISHEACSLAGTLKLNKLIALYDDN
3UK1_PYR           ----LTFSDYSRNALRVAALMKVPSIFVFTHDSI
5VRB_PP            -G-DGCLMEGVSHEACSLAGTLGLGKLIVLYDDN
5VRB_PYR           ----LMFSEYERNALRMAALMKINPVFVFTHDSI
1R9J_PP            -G-DGCLMEGVCQEALSLAGHLALEKLIVIYDSN
1R9J_PYR           --------GYALGAVRLAAISHHRVIYVATHDSI
1AY0_PP            -G-DGCLQEGISSEASSLAGHLKLGNLIAIYDDN
1AY0_PYR           GGTFLNFVSYAAGAVRLSALSGHPVIWVATHDSI
5HJE_PP            -G-DGCLMEGVSSEASSLAGHLQLGNLIAFWDDN
5HJE_PYR           ----------AAGAVRLSALSEFPITWVATHDSI
1ITZ_PP            -G-DGCQMEGIANEACSLAGHWGLGKLIAFYDDN
1ITZ_PYR           -------TDYMRGAMRISALSEAGVIYVMTHDSI
5ND6_PP            -G-DGCMMEGISNEACSLAGHWGLGKLIALYDDN
5ND6_PYR           ----YIFTDYMRNAMRMSALSEAGVVYVMTHDSI

CONSENSUS:         -G-DG---------ϕ---A--------i---DS-
```

**REPT 5**

```
4KXU_PP             LGQS--DPAPLQHQMDIYQK-RCEAFGWHAIIVD
4KXU_PYR            GVSIGEDGPSQMAL-EDLAMFR-------SVPTSTVFYP
3RIM_PP             ISI--EDDTNIALC-EDTAA-RYRAYGWHVQEVE
3RIM_PYR            GLG--EDGPTHQPI-EHLSALR-------AIPRLSVVRP
3M34_PP             ISI--EGDVGLAFN-ENVKM-RFEAQGFEVLSINGHD
3M34_PYR            GVG--EDGPTHQPI-EQLSTFR-------AMPNFLTFRP
3M49_PP             -ISL--DGDLNRSFSESVED-RYKAYGWQVIRVEDGND
3M49_PYR            AVG--EDGPTHEPI-EQLAALR-------AMPNVSVIRP
4C7V_PP             GISL--DGKTSASFTENVGA-RFEAYGWQYILVED
4C7V_PYR            AVGK--DGPTHEPI-EQLASLR-------TIPNVQVFRP
2R5N_PP             GISI--DGHVEGWFTDDTAM-RFEAYGWHVIRDID
2R5N_PYR            GLG--EDGPTHQPV-EQVASLR-------VTPNMSTWRP
4XEU_PP             GISI--DGEVHGWFTDDTPK-RFEAYGWQVIRNVD
4XEU_PYR            GLG--EDGPTHQPI-EQLASLR-------LTPNLDTWRP
3UK1_PP             GISI--DGDVVNWFHDDTPK-RFEAYGWNVIPNVNGHDVD
3UK1_PYR            GLG--EDGPTHQSV-EHVASLR-------LIPNLDVWRP
5VRB_PP             NISI--DGKVDGWFTENIPQ-RFESYGWHVVPNVNGHDTAAIQT
5VRB_PYR            GLG--EDGPTHQPI-EQTATLR-------LIPNMDVWRP
1R9J_PP             YISI--DGSTSLSFTEQCHQ-KYVAMGFHVIEVKN
1R9J_PYR            GVG--EDGPTHQPV-ELVAALR-------AMPNLQVIRP
1AY0_PP             KITI--DGATSISFDEDVAK-RYEAYGWEVLYVENGNEDL
1AY0_PYR            GVG--EDGPTHQPI-ETLAHFR-------SLPNIQVWRP
5HJE_PP             KISI--DGSTEVAFTEDVIA-RYKSYGWHIVEVSDADTDIT
5HJE_PYR            GLG--EDGPTHQPI-ETLAHFR-------ATPNISVWRP
1ITZ_PP             HISI--DGDTEIAFTEDVST-RFEALGWHTIWVKN
1ITZ_PYR            GLG--EDGPTHQPI-EHLVSFR-------AMPNILMLRP
5ND6_PP             KISI--DGHTDISFTEDVAK-RYEALGWHVIHVINGNTDVD
5ND6_PYR            GLG--EDGPTHQPI-EHLASFR-------AMPDMLMIRP

CONSENSUS:          giG--EDG-T-----E--A--R-------φφPN------
```

**REPT 6**

```
4KXU_PP        GHSVEELCKAFGQAKHQ----PTAIIAKTFK
4KXU_PYR       SDGVATEKAVELAANTKGICFIRTSRP
3RIM_PP        GGENVVGIEEAIANAQAVTDRPSFIALRTVI
3RIM_PYR       ADANETAYAWRTILARRNGSGPVGLILTRQ
3M34_PP        YEEINKALEQAKKS-----TKPCLIIAKTTI
3M34_PYR       ADGVENVKAWQIALNA---DIPSAFVLSRQ
3M49_PP        IEAIAKAIEEAKADE----KRPTLIEVRTTI
3M49_PYR       ADGNESVAAWRLALEST--NKPTALVLTRQDLPTLE
4C7V_PP        GFNLEEIDKAIVQAKAES-DKPTIIEIKTTI
4C7V_PYR       ADGNETSAAWKVALETL--DKPTILVLSRQNLDTLP
2R5N_PP        GHDAASIKRAVEEARAVT-DKPSLLMCKTII
2R5N_PYR       CDQVESAVAWKYGVERQ--DGPTALILSRQNLAQQERT
4XEU_PP        GHDADEIKTAIDTARKS--DQPTLICCKTVI
4XEU_PYR       ADAVESAVAWKHAIERA--DGPSALIFSRQN
3UK1_PP        AIDAAIAKAKRS-------DKPSLICCKTRI
3UK1_PYR       ADTVETAVAWTYAVAH---QHPSCLIFSRQNL
5VRB_PP        AIEAARAET----------GKPSIICCKTLIGKGSANKE
5VRB_PYR       CDTAESLVAWAEAAKAE--DHPSCLIFSRQNLKFQ
1R9J_PP        GDTDYEGLRKALAEAKATKGKPKMIVQTTTI
1R9J_PYR       SDQTETSGAWAVALSSI--HTPTVLCLS
1AY0_PP        AGIAKAIAQAKLSK-----DKPTLIKMTTTI
1AY0_PYR       ADGNEVSAAYKNSLESK--HTPSIIALSRQN
5HJE_PP        AIAAAIDEAKKVT------NKPTLVRLTTTI
5HJE_PYR       ADGNETSAAYKSAIEST--HTPHILALTRQNLP
1ITZ_PP        GNTGYDDIRAAIKEAKAVTDKPTLIKVTTTI
1ITZ_PYR       ADGNETAGAYKVAVLNR--KRPSILALS
5ND6_PP        GLRAAIAQAKAVK------DKPTLIKVSTLI
5ND6_PYR       AGGNETAGAYKVAIANR--KRPTTIALSRQNMP

CONSENSUS:     gDg-E-A-A---A------DKPTφφ--S-----
```

**Supplemental Figure S4:** Self-conserved sequence features within the functional repeats of the acetohydroxyacid synthases using a combination of automatic and manual alignment. The PDB ID and the functional domain in which the repeat is located are identified. Green highlight indicates overall conserved features identified either from sequence alignments or of the starting (G/A){X(1,2)}(G/A) motif. Yellow highlight indicates homologous substitutions while cyan indicates generally conserved features (such as a hydrophobic residue, indicated by ϕ). Grey highlight indicates a self-conserved residue not included in the previous categories. A consensus sequence is given at the bottom of each repeat in which positions of conserved identity are indicated by a capital letter, g = Gly or Ala, ϕ = hydrophobic residue, i = Val, Ile or Leu, + = cationic residue, a = aromatic residue, s = Ser or Asp.

**REPT 1**

```
2PAN_PYR        AVDAAMYVLEKEGIT---TAFGVP
2PAN_PP         PQRVYEEMNKAFGRD----VCYVT
5AHK_PYR        ASDAVAKILADNNVL---YGFELI
5AHK_PP         PFGLFTQLNKLTERV---ALDYILD
1YBH_PYR        GADILVEALERQGVE---TVFAYP
1YBH_PP         GEAIPPQYAIKVLDE---LTDGKAIIST
6DEK_PYR        GLTGGEIFHEMMLRHKVDTVFGYA
6DEK_PP         -PQTLIKEISDQAQTYNKEVIVTT
1N0H_PYR        GLTGGQIFNEMMSRQNVDTVFGYP
1N0H_PP         -PQTVIKKLSKVANDTGRHVIVTT

CONSENSUS:      g--g--------g------VFGi-
```

**REPT 2**

```
2PAN_PYR        GAAINPFYSAMRKH-GGIRHILARHVE
2PAN_PP         TIGLSQIAAAQMLHVFKDRHWINCGQA
5AHK_PYR        GGMITHLVDSINLLGKTKLVSMHHEQ
5AHK_PP         -VGNNQMWAAHTLRLNAQQAMHHSGGL
1YBH_PYR        GGASMEIHQAL-TRSSSIRNVLPRHEQ
1YBH_PP         GVGQHQMWAAQFYNYKKPRQWLSSGGL
6DEK_PYR        GGAILPVFDAIY-NSDKFKFVLPRHEQ
6DEK_PP         GVGQHQMWAAQHFTWTQPRTMITSGGL
1N0H_PYR        GGAILPVYDAIHNSDK-FNFVLPKHEQ
1N0H_PP         GVGQHQMWAAQHWTWRNPHTFITSGGL

CONSENSUS:      Gφg-----AA---+----Ra-L-----
```

**REPT 3**

```
2PAN_PYR        GA-SHMAEGYTRAT--AGNIGVCLGTS
2PAN_PP         GPLGWT-IPAALGVCAADPKRNVVAIS
5AHK_PYR        GA-AFAASAVSRVT-HHKTLGLALATS
5AHK_PP         GSMGFA-IPAAIGA-CYAGKKPIIVIT
1YBH_PYR        GG-VFAAEGYARSSGK--P-GICIATS
1YBH_PP         GAMGFG-LPAAIGASVANPDAIVVDID
6DEK_PYR        GA-GHMAEGYARASGK--P-GVVLVTS
6DEK_PP         GTMGYG-LPAAIGAQVAKPDAIVIDID
1N0H_PYR        GA-GHMAEGYARASGK--P-GVVLVTS
1N0H_PP         GTMGYG-LPAAIGAQVAKPESLVIDID

CONSENSUS:      Gg-gφg----A-----A-P---Vφ--s
```

**REPT 4**

```
2PAN_PYR     GPAGTDMIT---ALYS-ASADSI---PILCIT--GQ
2PAN_PP      GDFDFQFLI-EELAVG-AQFNI----PYIHVLV-NN
5AHK_PYR     GPGATNLIT-GIADCW-LDSH-----PCIFLTGQVN
5AHK_PP      GDGGAQLNI-QELDII-ARDKL----PILTIV-MNN
1YBH_PYR     GPGATNLVS-GLAD---ALLDSV---PLVAITGQVP
1YBH_PP      GDGSFIMNV-QEL----ATIRVENL-PVKVLL-LNN
6DEK_PYR     GPGATNVIT--PMAD--ALMDGV---PLVVFS-GQV
6DEK_PP      GDASFNMTL-TELSS--AVQAGA---PIKVCV-LNN
1N0H_PYR     GPGATNVVT--PMADAFADGIPMVVFTGQV---PTS
1N0H_PP      GDASFNMTL-TELSS--AVQAGT---PVKIL-ILNN

CONSENSUS:   G-g--N-----------A--------P--------N
```

**REPT 5**

```
2PAN_PYR     APRARLHKEDFQAVDIEAIAKPVSKMAVTVRE
2PAN_PP      GYGVDHVKVAEGLGCKAIRVFKPEDI
5AHK_PYR     THELKGKRDIRQQGFQELDSVALVTSITKYAYQIK
5AHK_PP      GYTSQFKKIGEAYRVESKTII
1YBH_PYR     RRMIGTDAFQETPIVEVTRSITKHNYLVM
1YBH_PP      GDPAQEDEIFPNMLLFAAACGIPAAR
6DEK_PYR     AIGTDAFQEADIVGISRSCTKWNVMVKN
6DEK_PP      QSNPDFMKLAESMNVKGIRITN
1N0H_PYR     AIGTDAFQEADVVGISRSCTKWNVMVKS
1N0H_PP      QLNPDFIKLAEAMGLKGLRVK

CONSENSUS:   g-g-Dg-K-AE--G------------
```

**REPT 6**

```
2PAN_PYR     AALVPRVLQQAFHLMRS-GRPGPVLVDLP
2PAN_PP      APAFEQAKALMAQYRV------PVVVEVILE
5AHK_PYR     SADELVPCLRKAIQIAKEGRPGPVLLDIP
5AHK_PP      SMQAFSSALESFLESPR-----PLLLEVSMSD
1YBH_PYR     DVEDIPRIIEEAFFLATSGRPGPVLVDVP
1YBH_PP      VTKKADLREAIQTMLDT---PGPYLLDVIC
6DEK_PYR     VAELPRRINEAFEIATT-GRPGPVLVDLP
6DEK_PP      QQELKSGVKEFLDATE------PVLLEVIVE
1N0H_PYR     VEELPLRINEAFEIATS-GRPGPVLVDLP
1N0H_PP      KQEELDAKLKEFVSTK-----GPVLLEVE

CONSENSUS:   --E--------F----T---PGPVLLDφ-
```

**Supplemental Figure S5A:** Cartoon illustration of the protein repeats present in the AHAS group of enzymes using the representative PDB 1n0h. The repeats are colored in the order red, yellow, green, cyan, blue, and magenta from N to C. Helical regions are indicated by a cylindrical tube and strand regions by an arrow when present in the PDB file. The number of the first helical residue in the helical region and the last beta residue in the strand region are also indicated. Insertions that contain secondary structure are indicated in grey while missing repeats are indicated by missing cartoon images. The three domains are shown in the order they occur in the protein. Secondary structure indication is derived from PyMol.

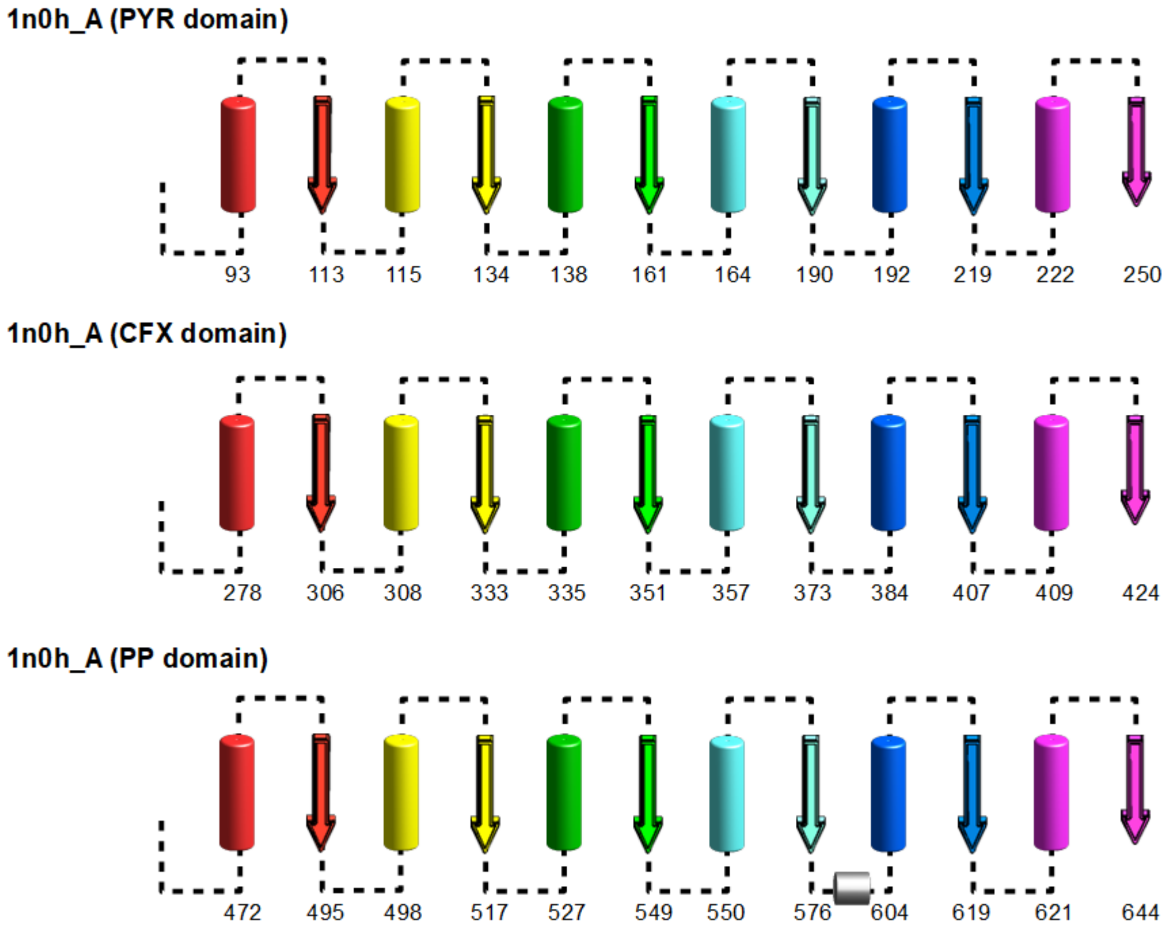

**Supplemental Figure S5B:** Cartoon illustration of the protein repeats present in the ALS group of enzymes using the representative PDB 4rji. The repeats are colored in the order red, yellow, green, cyan, blue, and magenta from N to C. Helical regions are indicated by a cylindrical tube and strand regions by an arrow when present in the PDB file. The number of the first helical residue in the helical region and the last beta residue in the strand region are also indicated. Insertions that contain secondary structure are indicated in grey while missing repeats are indicated by missing cartoon images. The three domains are shown in the order they occur in the protein. Secondary structure indication is derived from PyMol.

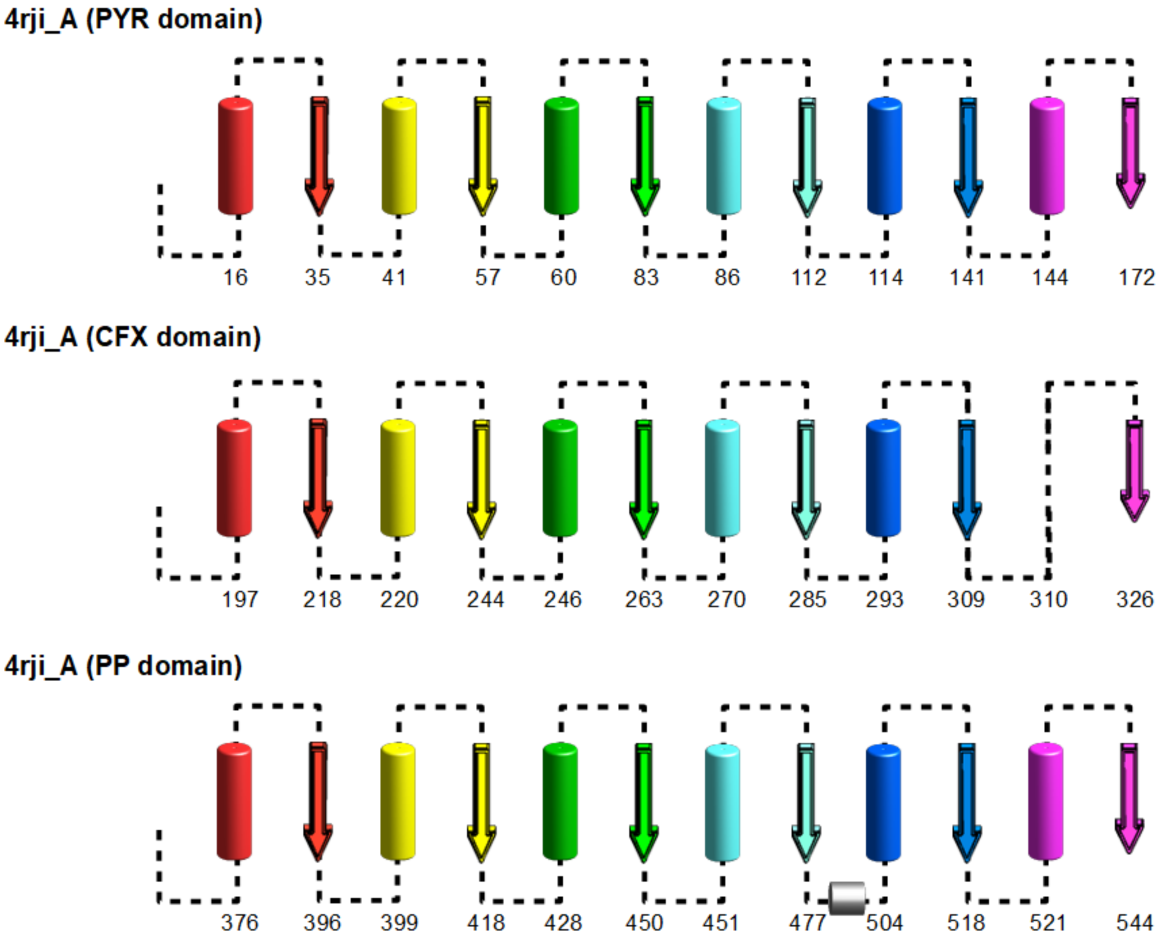

**Supplemental Figure S5C:** Cartoon illustration of the protein repeats present in the BFD group of enzymes using the representative PDB 2jlc. The repeats are colored in the order red, yellow, green, cyan, blue, and magenta from N to C. Helical regions are indicated by a cylindrical tube and strand regions by an arrow when present in the PDB file. The number of the first helical residue in the helical region and the last beta residue in the strand region are also indicated. Insertions that contain secondary structure are indicated in grey while missing repeats are indicated by missing cartoon images. The three domains are shown in the order they occur in the protein. Secondary structure indication is derived from PyMol. The CFX domain in PDB 2jlc only contains 4 identifiable repeats.

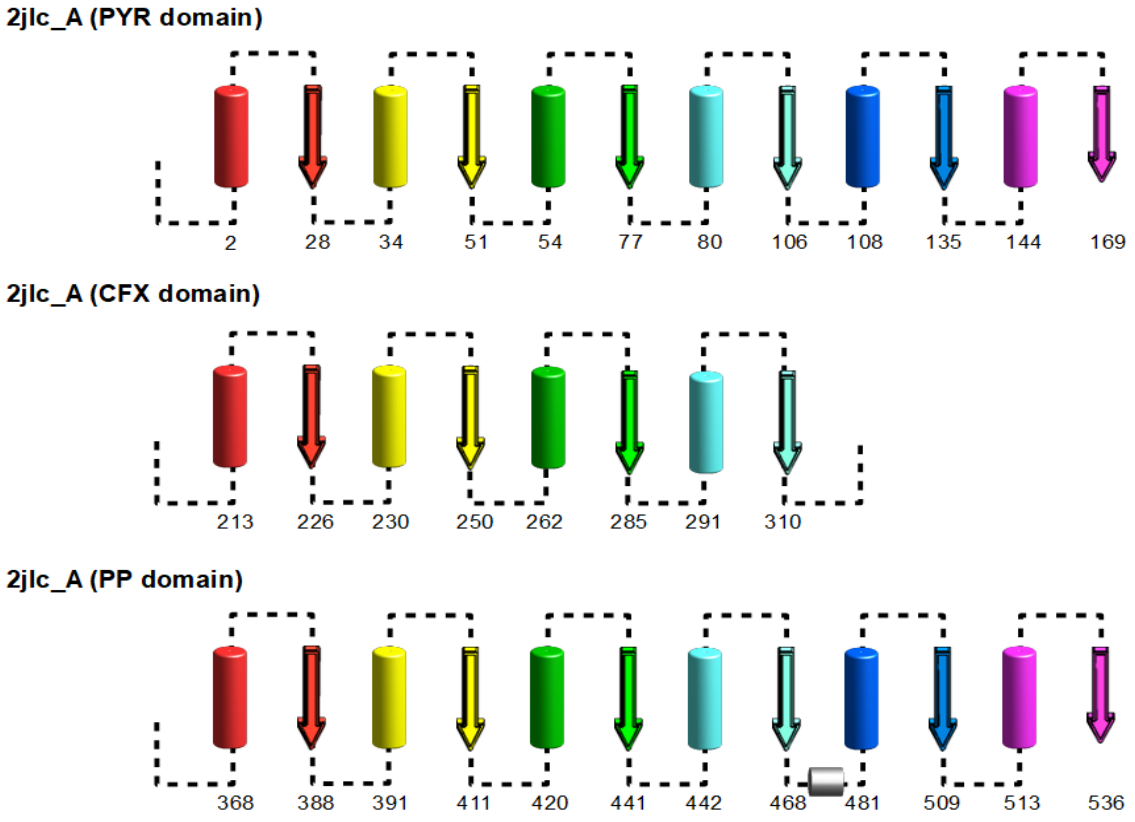

**Supplemental Figure S5D:** Cartoon illustration of the protein repeats present in the OxCDC group of enzymes using the representative PDB 4qq8. The repeats are colored in the order red, yellow, green, cyan, blue, and magenta from N to C. Helical regions are indicated by a cylindrical tube and strand regions by an arrow when present in the PDB file. The number of the first helical residue in the helical region and the last beta residue in the strand region are also indicated. Insertions that contain secondary structure are indicated in grey while missing repeats are indicated by missing cartoon images. The three domains are shown in the order they occur in the protein. Secondary structure indication is derived from PyMol.

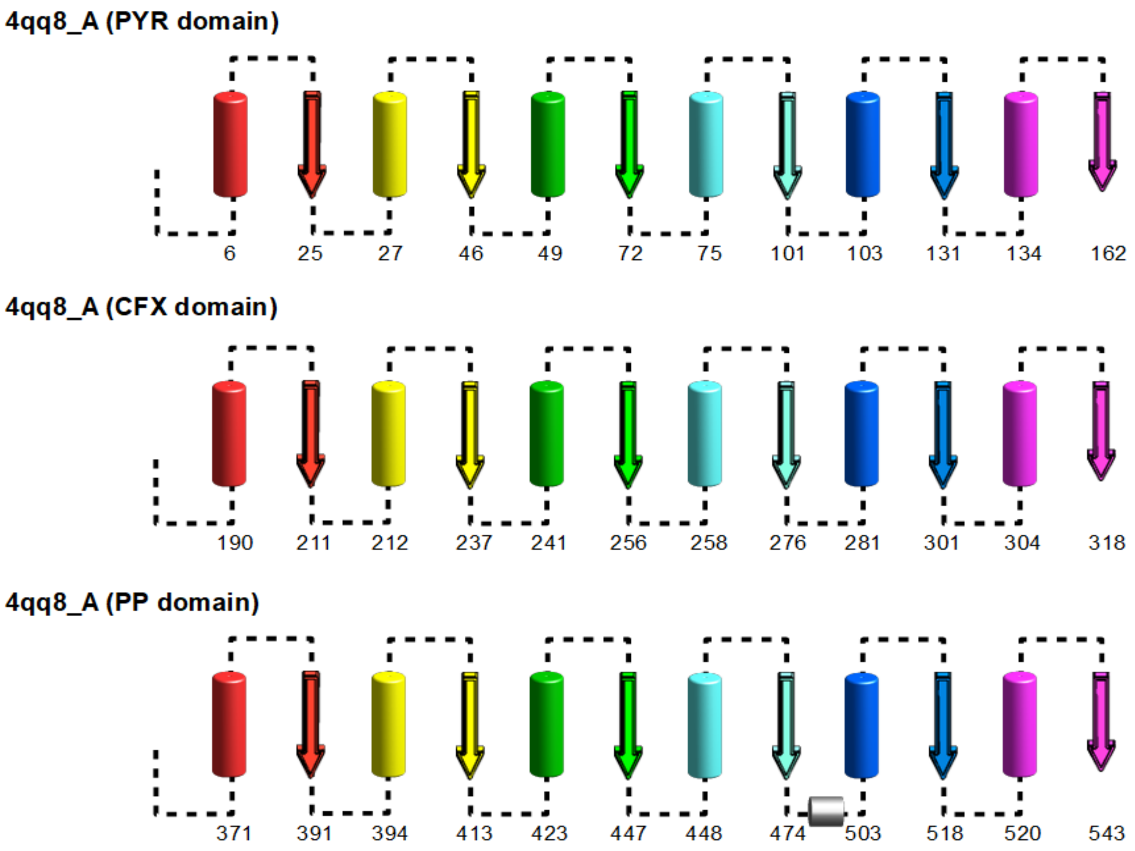

**Supplemental Figure S5E:** Cartoon illustration of the protein repeats present in the PDC group of enzymes using the representative PDB 5euj. The repeats are colored in the order red, yellow, green, cyan, blue, and magenta from N to C. Helical regions are indicated by a cylindrical tube and strand regions by an arrow when present in the PDB file. The number of the first helical residue in the helical region and the last beta residue in the strand region are also indicated. Insertions that contain secondary structure are indicated in grey while missing repeats are indicated by missing cartoon images. The three domains are shown in the order they occur in the protein. Secondary structure indication is derived from PyMol. The CFX domain in PDB 5euj only contains 5 repeats.

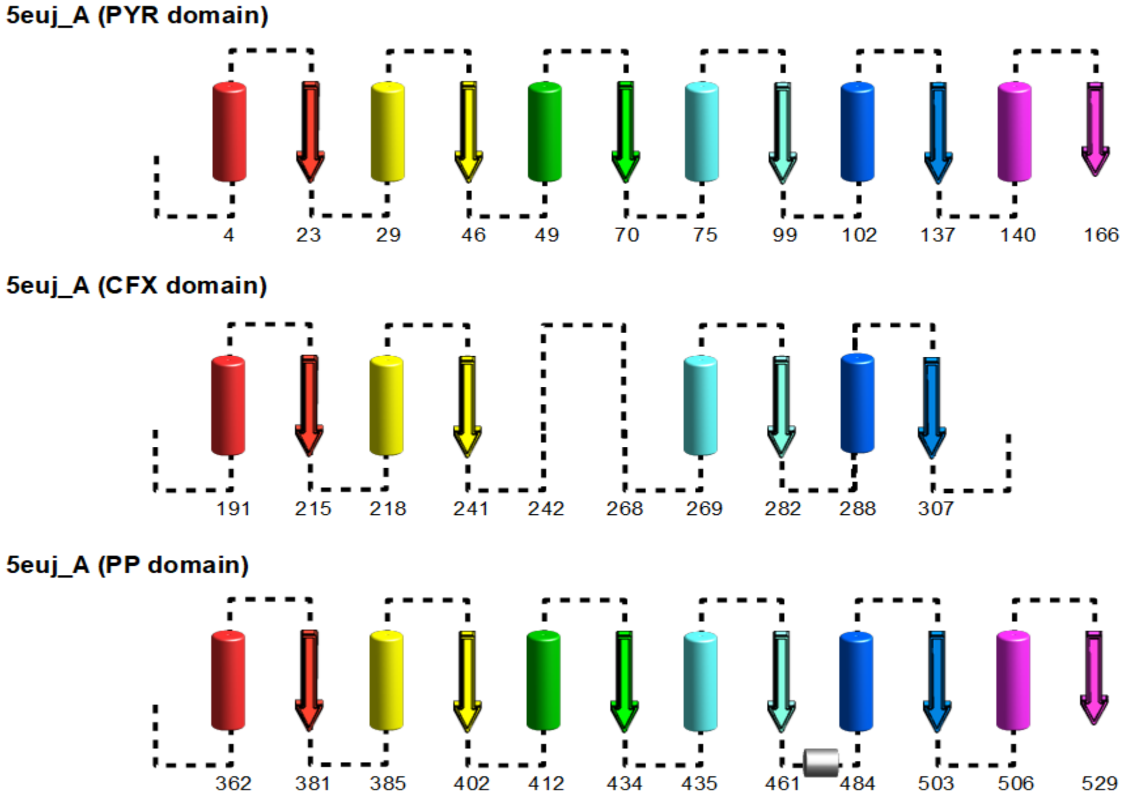

**Supplemental Figure S5F:** Cartoon illustration of the protein repeats present in the PDH group of enzymes using the representative PDB 2bp7. The repeats are colored in the order red, yellow, green, cyan, blue, and magenta from N to C. Helical regions are indicated by a cylindrical tube and strand regions by an arrow when present in the PDB file. The number of the first helical residue in the helical region and the last beta residue in the strand region are also indicated. Insertions that contain secondary structure are indicated in grey while missing repeats are indicated by missing cartoon images. The three domains are shown in the order they occur in the protein. Secondary structure indication is derived from PyMol.

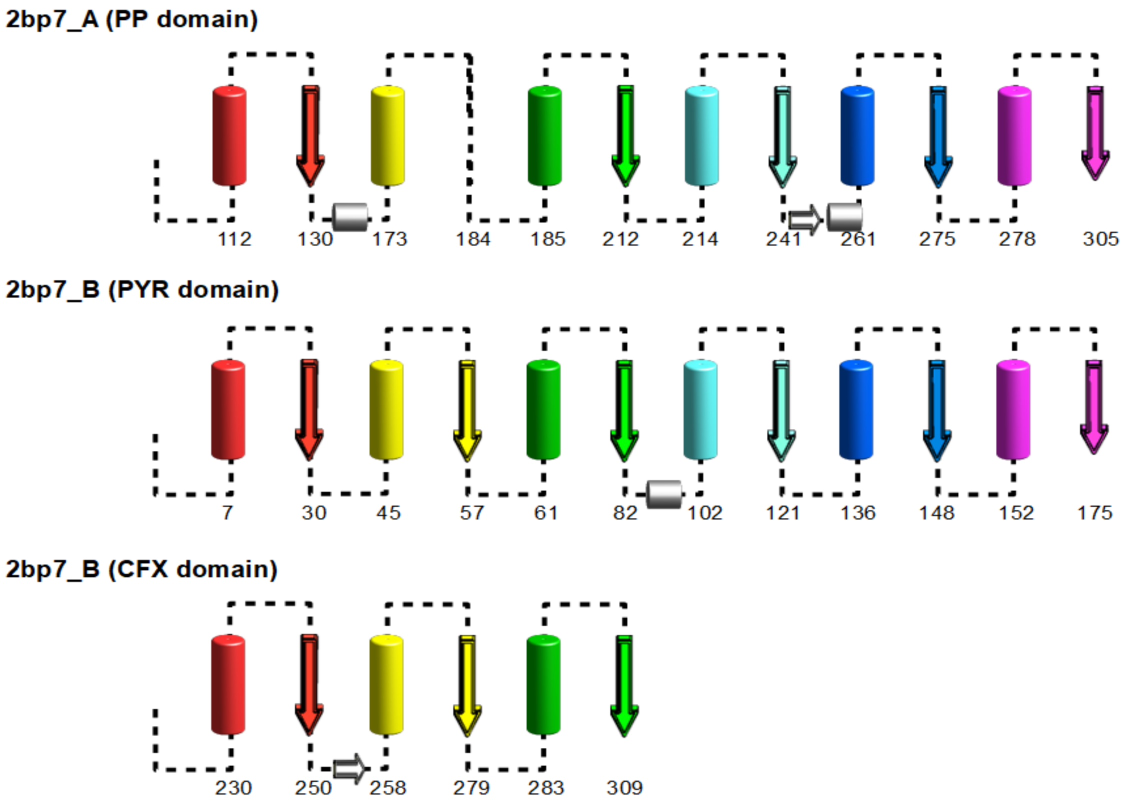

**Supplemental Figure S5G:** Cartoon illustration of the protein repeats present in the PhK group of enzymes using the representative PDB 6gua. The repeats are colored in the order red, yellow, green, cyan, blue, and magenta from N to C. Helical regions are indicated by a cylindrical tube and strand regions by an arrow when present in the PDB file. The number of the first helical residue in the helical region and the last beta residue in the strand region are also indicated. Insertions that contain secondary structure are indicated in grey while missing repeats are indicated by missing cartoon images. The three domains are shown in the order they occur in the protein. Secondary structure indication is derived from PyMol.

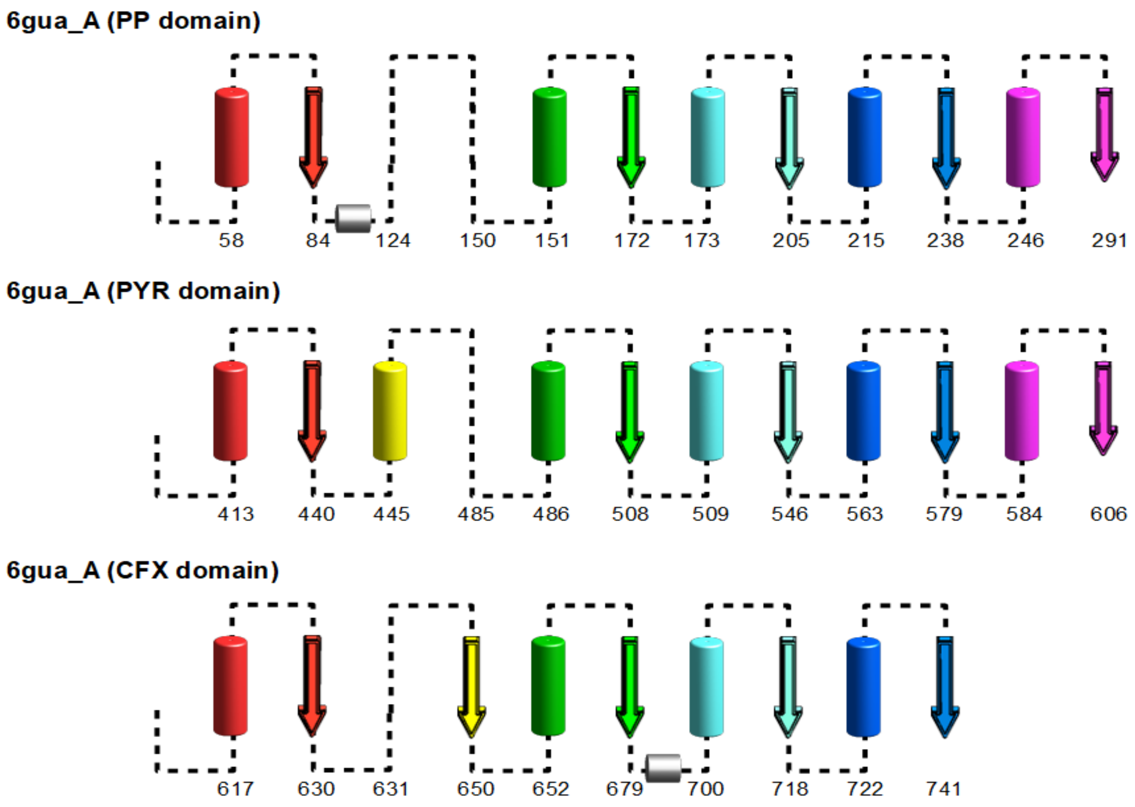

**Supplemental Figure S5H:** Cartoon illustration of the protein repeats present in the PFOR group of enzymes using the representative PDB 5c4i with each chain as indicated. The repeats are colored in the order red, yellow, green, cyan, blue, and magenta from N to C. Helical regions are indicated by a cylindrical tube and strand regions by an arrow when present in the PDB file. The number of the first helical residue in the helical region and the last beta residue in the strand region are also indicated. Insertions that contain secondary structure are indicated in grey while missing repeats are indicated by missing cartoon images. The three domains are shown in the order they occur in the protein. Secondary structure indication is derived from PyMol. The three CFX domains all lack one or two of the six repeats normally found in a domain.

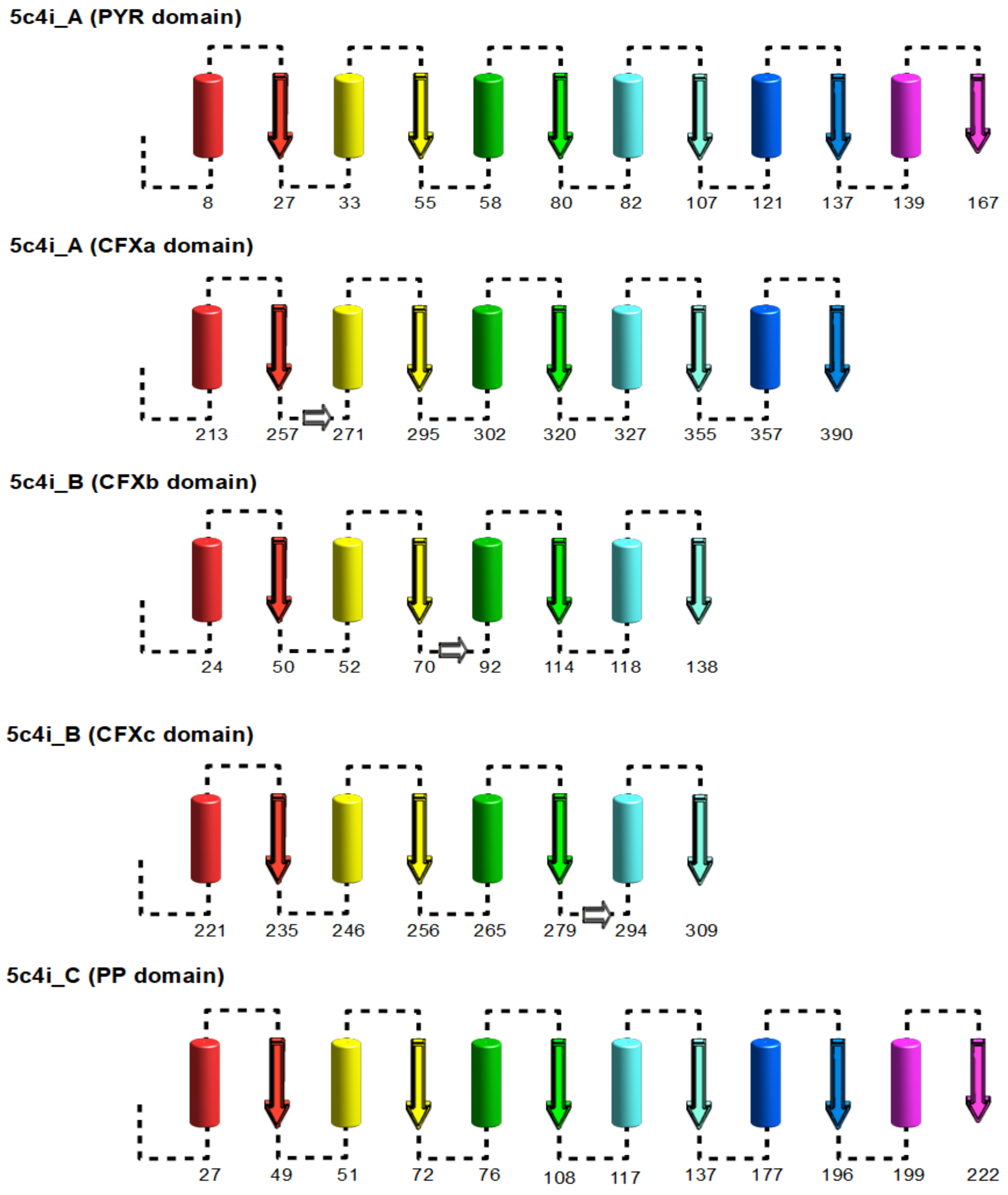

**Supplemental Figure S5I:** Cartoon illustration of the protein repeats present in the POX group of enzymes using the representative PDB 1y9d. The repeats are colored in the order red, yellow, green, cyan, blue, and magenta from N to C. Helical regions are indicated by a cylindrical tube and strand regions by an arrow when present in the PDB file. The number of the first helical residue in the helical region and the last beta residue in the strand region are also indicated.  Insertions that contain secondary structure are indicated in grey while missing repeats are indicated by missing cartoon images. The three domains are shown in the order they occur in the protein. Secondary structure indication is derived from PyMol. Note that the helical region in the fifth repeat of the CFX domain is not resolved in the structure so it is colored grey here.

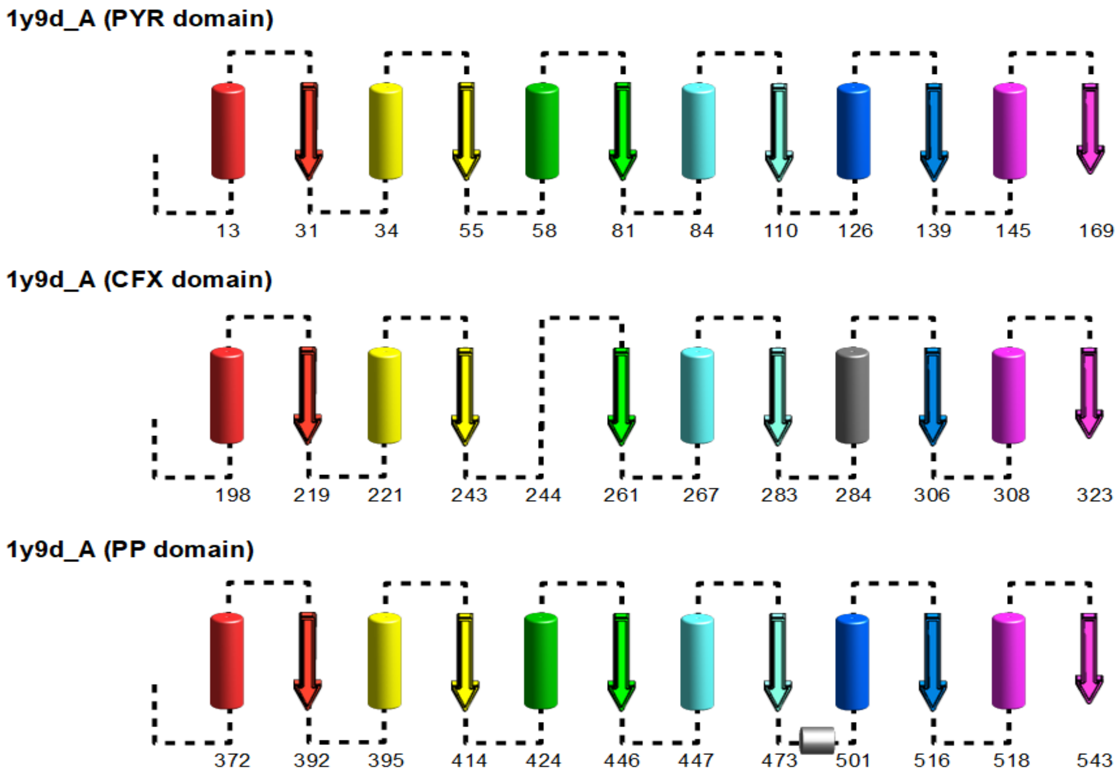

**Supplemental Figure S5J:** Cartoon illustration of the protein repeats present in the TK group of enzymes using the representative PDB 1itz. The repeats are colored in the order red, yellow, green, cyan, blue, and magenta from N to C. Helical regions are indicated by a cylindrical tube and strand regions by an arrow when present in the PDB file. The number of the first helical residue in the helical region and the last beta residue in the strand region are also indicated. Insertions that contain secondary structure are indicated in grey while missing repeats are indicated by missing cartoon images. The three domains are shown in the order they occur in the protein. Secondary structure indication is derived from PyMol.

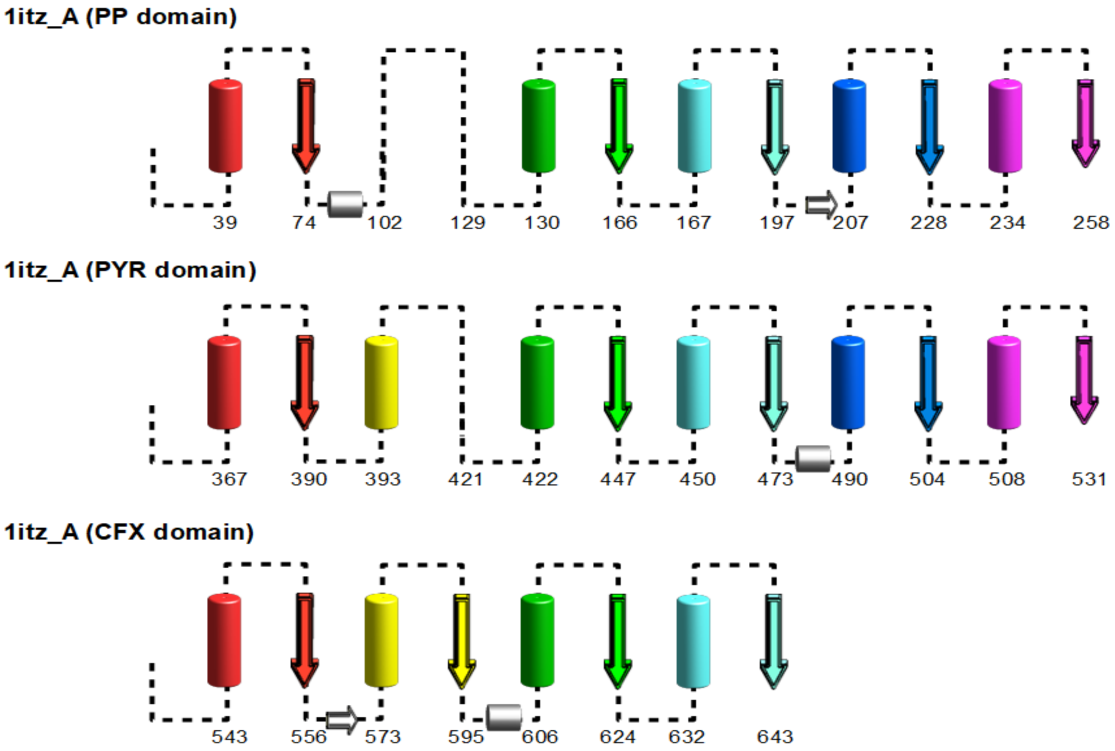

**Supplemental Figure S6A:** Cartoon illustration of the three dimensional repeat organization in the functional (PP & PYR) ThDP enzyme domains. Repeats are colored and numbered from N to C: red, yellow, green, cyan, blue, magenta with strands represented as arrows and helices as cylinders. In this domain all the β-strands in a 213465 arrangement are all parallel while the sets of helices (1-3 & 4-6) each have helices both above and below the plane of the β-sheet as indicated in the cartoon.

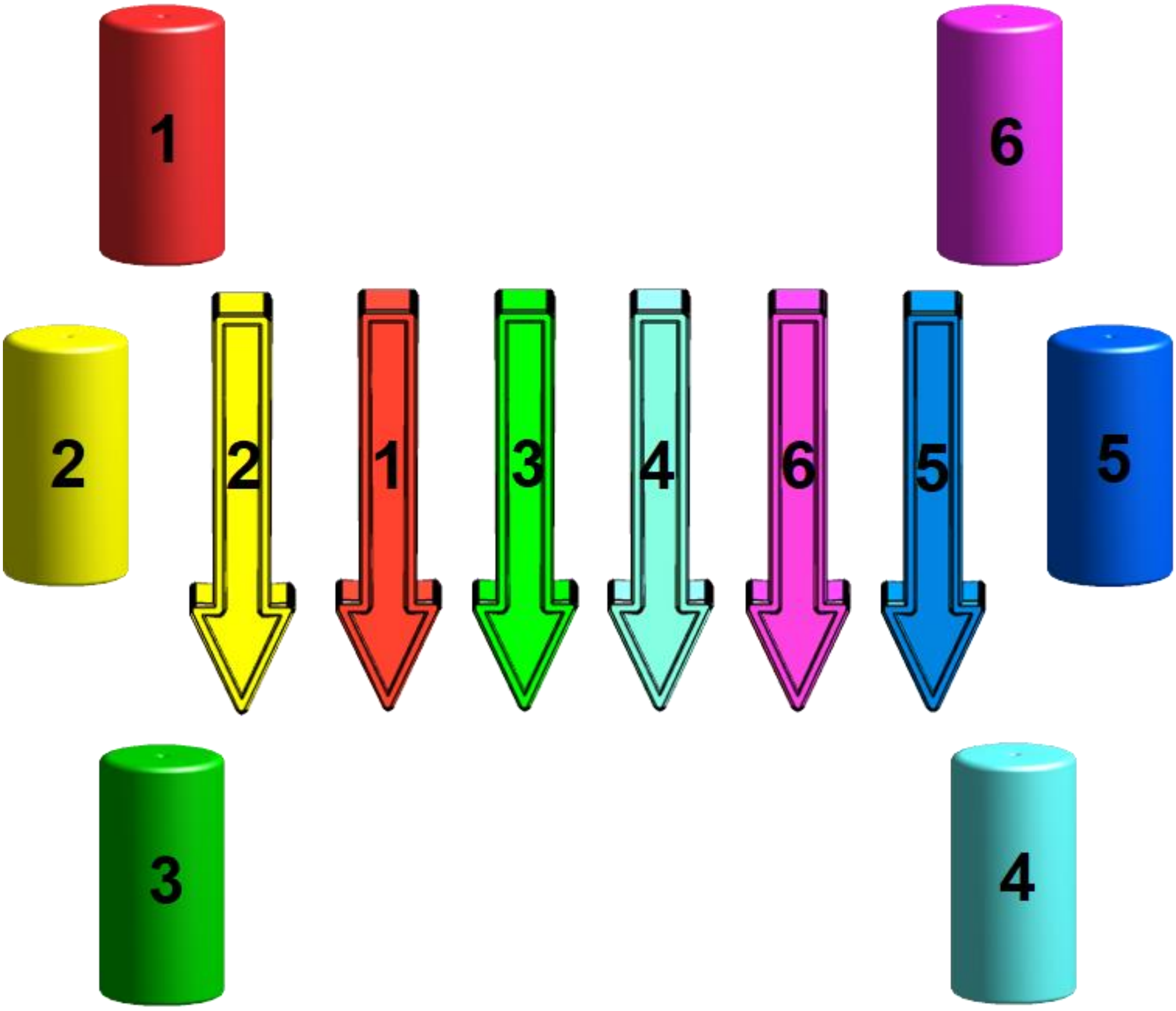

**Supplemental Figure S6B:** Cartoon illustration of the three dimensional repeat organization in the CFX domains from the DC class of ThDP enzymes. Repeats are colored and numbered from N to C: red, yellow, green, cyan, blue, magenta with strands represented as arrows and helices as cylinders. The β-strands in this domain are arranged in a parallel Rossmann-like 321456 arrangement while the sets of helices (1-3 & 4-6) are separate units either above or below the plane of the β-sheet as indicated in the cartoon. Similarity to the Rossmann domain appears to be the result of convergent evolution as the enzymes that bind a second ribose cofactor (FADH or ATP) do not maintain the conserved binding geometry that standard Rossman domain enzymes possess (see SI table S7).

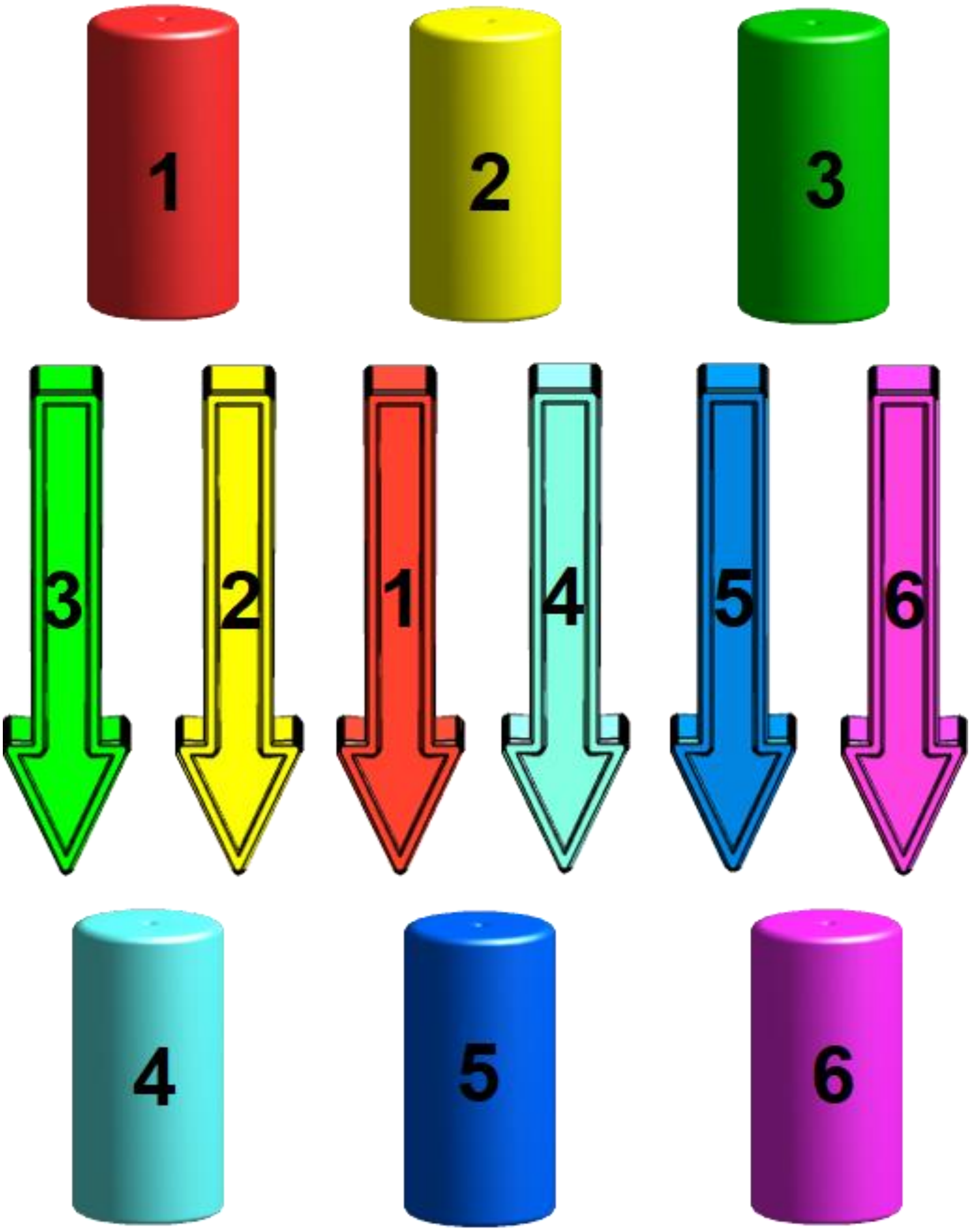

**Supplemental Figure S6C:** Cartoon illustration of the three dimensional repeat organization in the CFX domains of the DH class of ThDP enzymes. Repeats are colored and numbered from N to C: red, yellow, green, cyan, blue, magenta with strands represented as arrows and helices as cylinders. Type C domains are present in the CFX domains of the dehydrogenase (DH) class of ThDP enzymes. In this domain all the β-strands in a 1(-2)(-3)(-4) arrangement with the last three strands being anti-parallel to the first one with the sets of helices (1-2 & 3-4) are separate units either above or below the plane of the β-sheet as indicated in the cartoon.

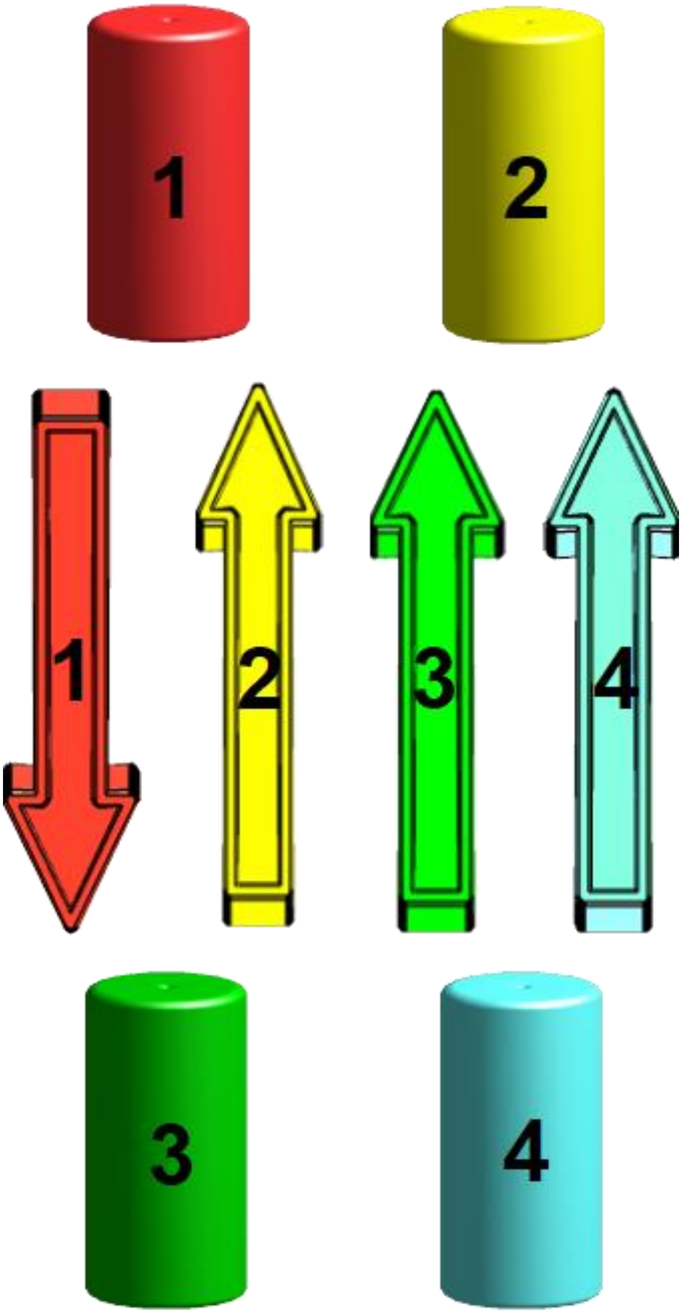

**Supplemental Figure S6D:** Cartoon illustration of the three dimensional repeat organization in the first type of CFX domains found in PFOR enzymes. Repeats are colored and numbered from N to C: red, yellow, green, cyan, blue, magenta with strands represented as arrows and helices as cylinders. In these domains all the β-strands in a 2(-3)1(-4) arrangement with the last two strands being anti-parallel to the first one. The sets of helices (1-2 & 3-4) each have helices both above and below the plane of the β-sheet as indicated in the cartoon.

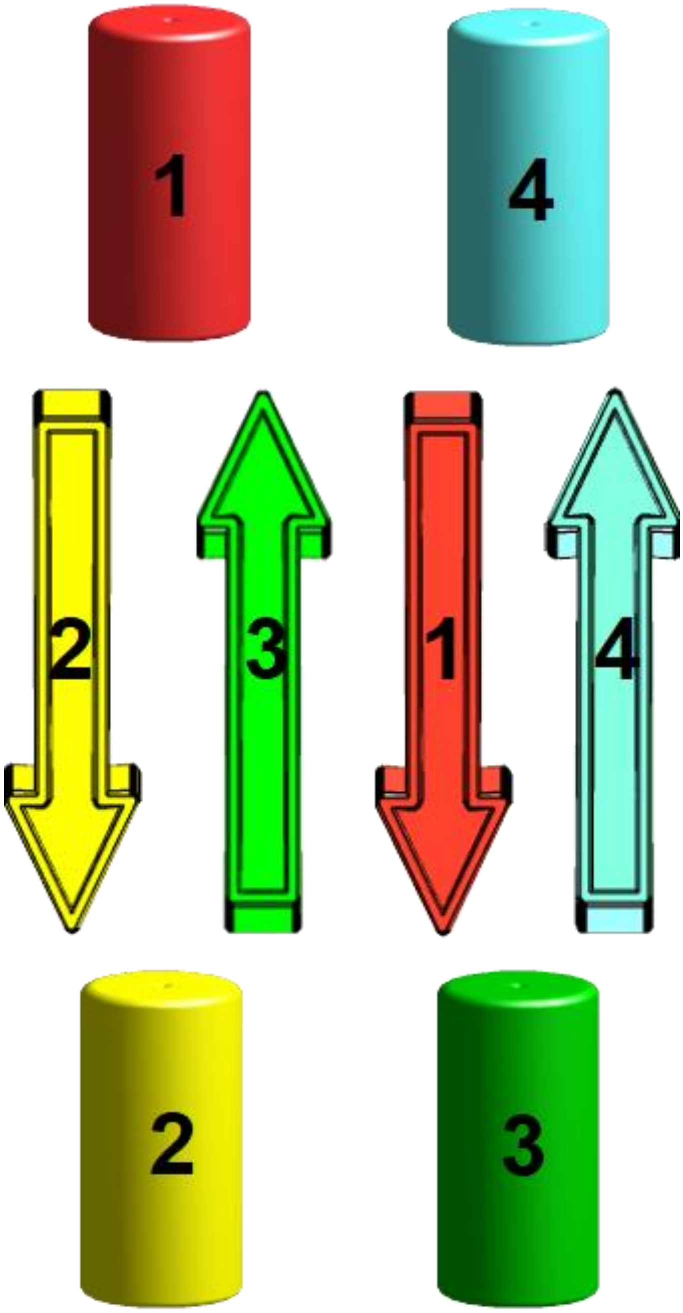

**Supplemental Figure S6E:** Cartoon illustration of the three dimensional repeat organization in the second type of CFX domains found in PFOR enzymes. Repeats are colored and numbered from N to C: red, yellow, green, cyan, blue, magenta with strands represented as arrows and helices as cylinders. In this domain all the β-strands in a 1(-2)34(-5) arrangement with the second strand being anti-parallel to the others as indicated in the cartoon. The sets of helices (1-3 & 4-5) both appear to have helices both above and below the plane of the β-sheet. This domain is likely related to a C domain with a 180˚ rotation of the second half of the domain (strands 3 & 4) and the addition of a fifth, antiparallel repeat.

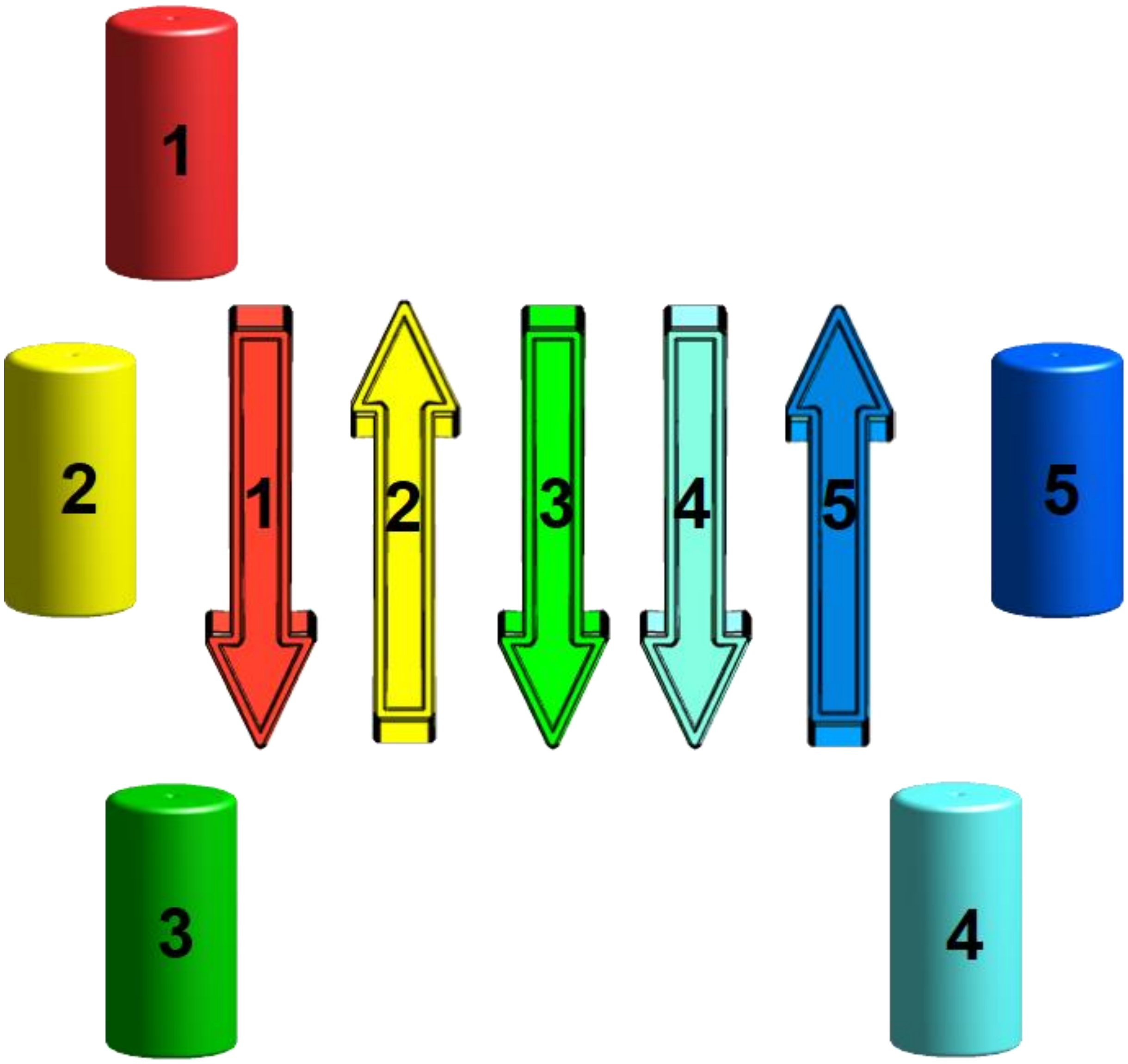

**Supplemental Figure S7A:** Cartoon illustration of the three dimensional structure of the conserved alanine in position 10 of repeat 3 in the functional domains of the transketolases (figure made from PDB ID 3m49 ) The PP domain is in light green and the PYR domain repeat is in dark green. The conserved alanine are indicated in red.

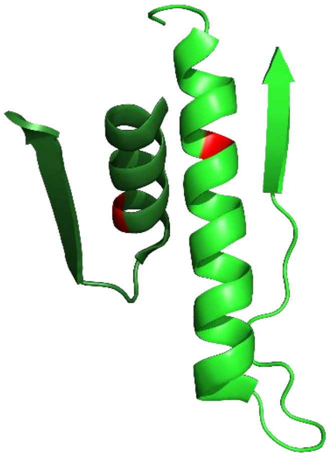

**Supplemental Figure S7B:** Cartoon illustration of the three dimensional structure of the conserved alanine in position 10 of repeat 3 in the functional domains of the acetohydroxyacid synthases (figure made from PDB ID 1n0h) The PP domain is in light green and the PYR domain repeat is in dark green. The conserved alanines are indicated in red.

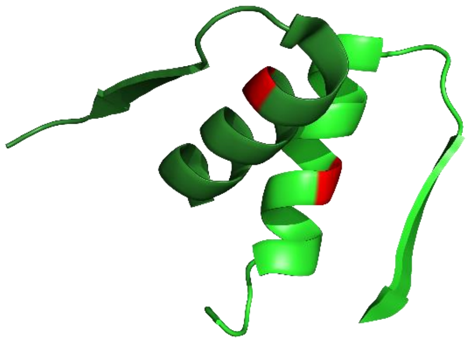

**Supplemental Figure S8A:** Cartoon illustration of the three dimensional structure of the inter chain contacts made by repeats 4 & 5 in the functional domains of the transketolases (figure made from PDB ID 3m49). Repeat 4 is shown in cyan and repeat 5 is shown in blue. Conserved positions 5 & 6 in repeat 5 are shown in red and conserved position 16 in yellow.

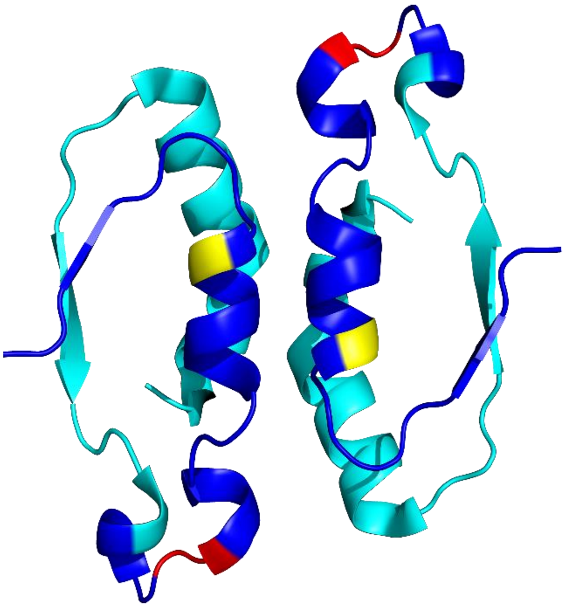

**Supplemental Figure S8B:** Cartoon illustration of the three dimensional structure of the inter chain contacts made by repeats 4 & 5 in the functional domains of the acetohydroxyacid synthases (figure made from PDB ID 1n0h). Repeat 4 is shown in cyan and repeat 5 is shown in blue.  Conserved positions 5 & 6 in repeat 5 are shown in red and conserved position 14 in yellow

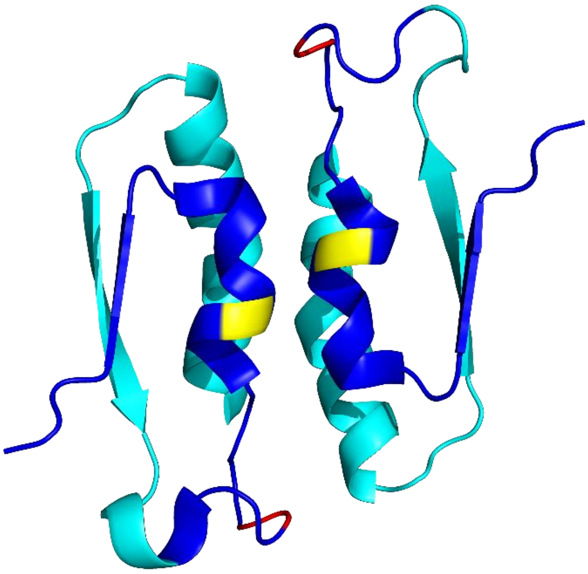

.

**Supplemental Figure S9:** Cartoon representation of the inter chain contact formed by repeat 5 (shown with grey carbons) in benzoylformate dehydrogenase (figure made from PDB ID 6a50). While not often modeled this way, this structure very clearly shows a bound magnesium mediating the inter chain contact.

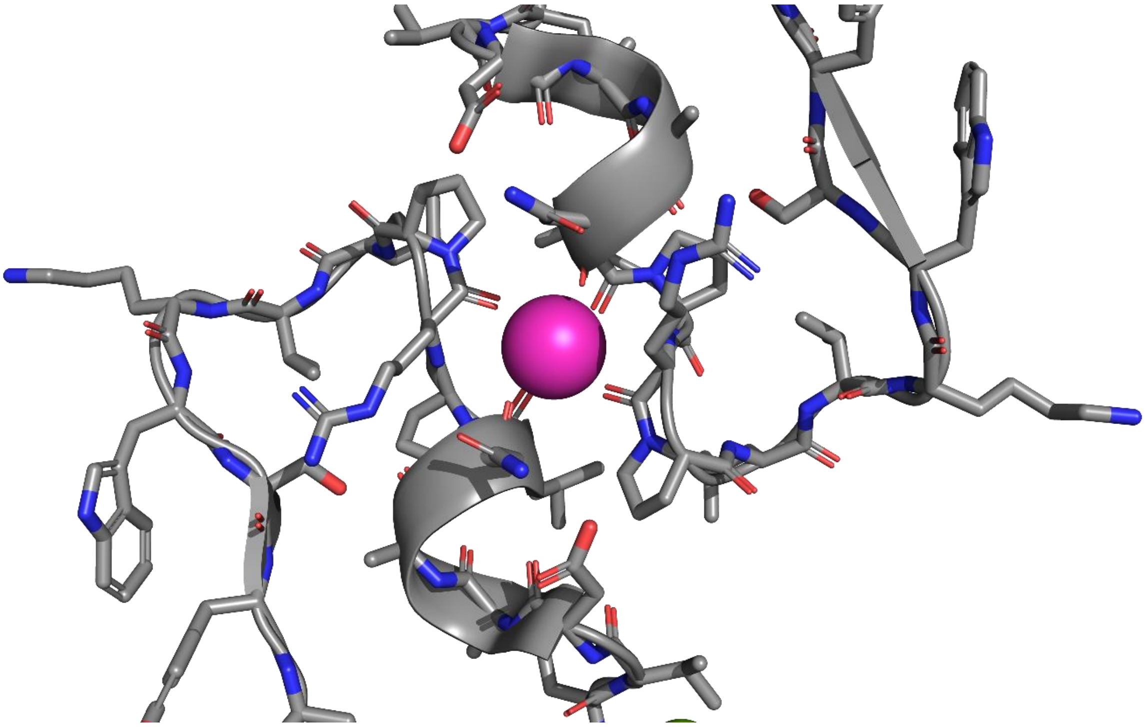

**Supplemental Data S1:** Results from searching other web-available repeat detection methods using the 10 proteins used as secondary structure examples from each of the sub-groups of ThDP enzymes in Supplemental Data 1 as query sequences.

**RADAR Nucleic Acids Research, 01 Jul 2019, 47(W1):W636-W641**
**DOI: 10.1093/nar/gkz268**

```
>2PAN:A|PDBID|CHAIN|SEQUENCE
---------------------------------------------------------------------------
No. of Repeats|Total Score|Length  |Diagonal| BW-From|   BW-To|   Level
             2|      47.66|      14|      51|      46|      59|       1
---------------------------------------------------------------------------
   46-   59 (27.81/12.71)       FGVPGAA....INPFYSA
   96-  113 (19.84/ 7.56)       LGTSGPAgtdmITALYSA
---------------------------------------------------------------------------
---------------------------------------------------------------------------
No. of Repeats|Total Score|Length  |Diagonal| BW-From|   BW-To|   Level
             2|      62.15|      17|      61|       2|      22|       2
---------------------------------------------------------------------------
    2-   22 (30.30/24.99)       GSSHHHHHHSSGlvprGSHMA
   65-   81 (31.85/16.70)       GIRHILARHVEG....ASHMA
---------------------------------------------------------------------------
---------------------------------------------------------------------------
No. of Repeats|Total Score|Length  |Diagonal| BW-From|   BW-To|   Level
             4|     227.06|      61|     341|     167|     232|       3
---------------------------------------------------------------------------
  136-  165 (11.64/ 7.57)       .................D.FQAVDIE.........AIAKP.VSKMAVtvrEAAlVPRV.........
  171-  232 (102.42/57.52)      HLMRSGRPGPVLVDLPFD.VQVAEIEfDPDMYEPLPVYKPAASRMQI...EKA.VEMLIQAERPVIV
  434-  459 (31.17/11.62)       HWINCGQAGPLGWTIPAA.LGVCAA..DP.....................................
  520-  574 (81.83/40.56)       ENINSSEVNGYGVD....hVKVAE....GLGCKAIRVFKPEDIAPAF...EQA.KALMAQYRVPVVV
---------------------------------------------------------------------------

>2PGN:A|PDBID|CHAIN|SEQUENCE
---------------------------------------------------------------------------
No. of Repeats|Total Score|Length  |Diagonal| BW-From|   BW-To|   Level
             2|      68.51|      19|      21|     338|     358|       1
---------------------------------------------------------------------------
  338-  358 (30.80/26.50)       GFkaVRYQERENFRQATEFRA
  362-  380 (37.71/25.65)       GW..VREQESGDGMPASMFRA
---------------------------------------------------------------------------
---------------------------------------------------------------------------
No. of Repeats|Total Score|Length  |Diagonal| BW-From|   BW-To|   Level
             2|      65.70|      22|      26|     389|     414|       2
---------------------------------------------------------------------------
  389-  414 (32.30/25.29)       RpedIiVTDIGNHTL....PMFGGAILQRP
  416-  441 (33.41/15.11)       R...L.VTSMAEGILgcgfPMALGAQLAEP
---------------------------------------------------------------------------
---------------------------------------------------------------------------
No. of Repeats|Total Score|Length  |Diagonal| BW-From|   BW-To|   Level
             2|      76.30|      23|      25|     252|     275|       3
---------------------------------------------------------------------------
  252-  275 (38.90/27.41)       LAMGSaGFCGWKSANDMMA.AADFV
  276-  299 (37.40/21.76)       LVLGS.RLSDWGIAQGYITkMPKFV
---------------------------------------------------------------------------
---------------------------------------------------------------------------
No. of Repeats|Total Score|Length  |Diagonal| BW-From|   BW-To|   Level
             2|      56.32|      17|      24|     542|     565|       5
---------------------------------------------------------------------------
  542-  564 (22.40/29.68)       EIPVSktQGLASDPvggvGPNLL
  569-  585 (33.92/16.72)       EIPVD..TGGSMYP....GENLL
---------------------------------------------------------------------------
---------------------------------------------------------------------------
No. of Repeats|Total Score|Length  |Diagonal| BW-From|   BW-To|   Level
             2|      58.30|      15|      19|     479|     493|       6
---------------------------------------------------------------------------
  479-  493 (29.50/17.44)       ESYGANWTLMNHQFG
  499-  513 (28.80/16.87)       EFMNPDWVGIAKAFG
---------------------------------------------------------------------------

>6A50:A|PDBID|CHAIN|SEQUENCE
---------------------------------------------------------------------------
No. of Repeats|Total Score|Length  |Diagonal| BW-From|   BW-To|   Level
```

```
             3|     558.64|     133|     209|      41|     177|       1
---------------------------------------------------------------------------
   41-  174 (222.16/140.81)
        VFGNPGS......NELPFLKDFPEDFRYILALQEACVVG.IADGY..AQASRKPAFINLHSAA..GTGNAMGALSNARTShSPLIVTA...GQQT
RAMIGVEAGETNVDAANL....P....RPLVKWSYEPASA...AEVPHAMSRAIHMASMAPQGP
  252-  391 (185.89/112.07)
        VWVAPSA......PRCPFPTRHPC.FRGLMPAGIAAISQ.LLEGHdvVLVIGAPVFRYVFYDP..GQYLKPGTRLISVTC.DPLEAARapmGDAI
VADIGAMASAL....ANLveesS....RQLPTAAPEPAKVdqdAGRLHPETVFDTLNDMAPENA
  412-  537 (150.59/91.24)
        ...NPGSyyfcaaGGLGF..ALPAAIGVQLAEPERQVIAvIGDGS..ANYS....ISALWTAAqyNIPTIFVIMNNG.........TY...G.ML
RWFAGVLEAE.NVPGLDV....PgidfRALAKGYGVQALK...ADNLEQLKGSLQEA.LSAKGP
---------------------------------------------------------------------------

>4QQ8:A|PDBID|CHAIN|SEQUENCE
---------------------------------------------------------------------------
No. of Repeats|Total Score|Length  |Diagonal| BW-From|   BW-To|   Level
             2|     593.77|     189|     344|      16|     212|       1
---------------------------------------------------------------------------
   16-  212 (298.01/167.38)
        KAGVEHlfGLHGIHIDTIFQACLDHDVPII.DTRHEAAAGHAAEGYARAGAKL..GVaLVTAGGGFTNAV.TPIANARTDRTPVLfLTGSGAL..
RDDETNTL.QAGIDQVAMAAPITKWAHrvmaTEHIPRLVMQAIRAALSAPRGPVLLDLPWDILMNQIDEDSVIIPDLVLSAHGAHPDPADLDQALALLRKAER
PVIVLG
  363-  558 (295.76/148.51)
        KSSSEH..ALHPFHASQVIAKHVDAGVTVVaDGGLTYLWLSEVMSRVKPGGFLchGY.LNSMGVGFGTALgAQVADLEAGRRTIL.VTGDGSVgy
SIGEFDTLvRKQLPLIVIIMNNQSWGW....TLHFQQLAVGPNRVTGTRLENGSYHGVAAAFGADGYHVDSVESFSAALAQALAHNRPACINVAVALDPIPPE
ELILIG
---------------------------------------------------------------------------
---------------------------------------------------------------------------
No. of Repeats|Total Score|Length  |Diagonal| BW-From|   BW-To|   Level
             2|     118.03|      39|      67|     230|     271|       2
---------------------------------------------------------------------------
  230-  271 (64.53/38.15)        ATGVPVFADYEGLSMLSGLPDAMR...GGLVQNLysfAKA...DAA.PD
  296-  341 (53.50/24.93)        AQVIQVDPDACELGRLQGIALGIVadvGGTIEAL...AQAtaqDAAwPD
---------------------------------------------------------------------------

>2NXW:A|PDBID|CHAIN|SEQUENCE
---------------------------------------------------------------------------
No. of Repeats|Total Score|Length  |Diagonal| BW-From|   BW-To|   Level
             2|     412.31|     143|     161|      72|     232|       1
---------------------------------------------------------------------------
   72-  232 (212.11/150.57)
        GFAAdAAARYSSTLGVAAVT..YGAGAfnMvnavagAYAEKSPVVVISGAPG......TTEGNAGLLLhhQGRTL.DTQFqvfkeiTVAQARLDD
PAKAPA.EIARVLGAARAQSRPVYLEIP...RNMVNAEVEPVGDDP.AWP....VDRDALAA..CADEVLAAMRSATSPvLMVCVE
  237-  399 (200.19/107.05)
        GLEA.KVAELAQRLGVPVVTtfMGRGL..L......ADAPTPPLGTYIGVAGdaeitrLVEESDGLFL..LGAILsDTNF......AVSQRKIDL
RKTIHAfDRAVTLGYHTYADIPLAGLVDallERLPPSDRTTRGKEPhAYPtglqADGEPIAPmdIARAVNDRVRAGQEP.LLIAAD
---------------------------------------------------------------------------
---------------------------------------------------------------------------
No. of Repeats|Total Score|Length  |Diagonal| BW-From|   BW-To|   Level
             3|     108.02|      26|      38|     445|     470|       2
---------------------------------------------------------------------------
  414-  434 (28.69/10.84)        GLMAPGYY.AGMGFGVPA.....GIGA.
  445-  470 (45.93/21.11)        TVVGDGAF.QMTGWELGN.CRRLGIDPI
  484-  511 (33.40/13.64)        TFQPESAFnDLDDWRFADmAAGMGGDGV
---------------------------------------------------------------------------

>1NI4:A|PDBID|CHAIN|SEQUENCE
---------------------------------------------------------------------------
No. of Repeats|Total Score|Length  |Diagonal| BW-From|   BW-To|   Level
             2|      60.49|      17|      36|     241|     257|       1
---------------------------------------------------------------------------
  241-  257 (31.44/21.07)        TRFAAAYCRSGKGPILM
  278-  294 (29.05/18.96)        TREEIQEVRSKSDPIML
---------------------------------------------------------------------------
---------------------------------------------------------------------------
No. of Repeats|Total Score|Length  |Diagonal| BW-From|   BW-To|   Level
             2|     126.43|      36|     166|      40|      77|       2
---------------------------------------------------------------------------
   40-   77 (61.66/44.56)        YRMMQTVRRMELKADQLYKQKIIRGFchLCDGQEACCV
  202-  237 (64.77/39.36)        YGMGTSVERAAASTDYYKRGDFIPGL..RVDGMDILCV
---------------------------------------------------------------------------
>1NI4:B|PDBID|CHAIN|SEQUENCE
---------------------------------------------------------------------------
No. of Repeats|Total Score|Length  |Diagonal| BW-From|   BW-To|   Level
```

```
             2|      93.89|      25|     201|      71|      97|       1
---------------------------------------------------------------------------
    71-   97 (43.41/32.54)        EMGFAGIAVGAAMAGlrPICEFMTFNF
   275-  299 (50.48/31.68)        EGGWPQFGVGAEICA..RIMEGPAFNF
---------------------------------------------------------------------------

>3AHC:A|PDBID|CHAIN|SEQUENCE
---------------------------------------------------------------------------
No. of Repeats|Total Score|Length  |Diagonal| BW-From|   BW-To|   Level
             4|     503.27|     149|     332|      51|     244|       1
---------------------------------------------------------------------------
    10-  172 (268.72/153.21)
        HSSGLV.PRGSHMTNPVIGTPWQKLDRPvseeaiegmdkywrVTNYMSIGQIYLRSNPLMKEPFTRDDVKH...........RLVGH.WGTTPGL
NFLLAHINRLI.ADHQQNTVF.........IMGPGHGGPAGTSQSYVDGTYTEYYPN...ITKDE..AG..........LQKFFRQFSY.PGGIPSHFAPETP
GSIH
   215-  280 (85.28/108.68)
        QSNKLVnPRTDGIVLPILHLNGYKIANP....................................................................
......................................................................tilarisdeeLHDFFRGMGYhPYEFVAGFDNEDH
MSIH
   380-  467 (83.57/34.49)
        .........................................VTAFMPKGELRIGANPNANGGVIREDLKLpeldqyevtgvKEYGHgWGQVEAP
RALGAY.CRDI.IKNNPDSFR.........IFGPDE.....TASNRLNATY.....................................................
....
   556-  642 (65.71/22.87)
        .......................................................NLLVSSHVWRQD..H..........NGFSH...QDPGV
TSLL..INKTFnNDHVTNIYFatdanmllaI.....SEKCFKSTNKINAIFAGKQPAptwVTLDEarAE..........LE......................
....
---------------------------------------------------------------------------
---------------------------------------------------------------------------
No. of Repeats|Total Score|Length  |Diagonal| BW-From|   BW-To|   Level
             2|      39.77|      12|     332|     341|     352|       2
---------------------------------------------------------------------------
   341-  352 (22.59/14.45)        QVPLASARD..TEE
   661-  674 (17.18/ 9.36)        QVVLASAGDvpTQE
---------------------------------------------------------------------------
---------------------------------------------------------------------------
No. of Repeats|Total Score|Length  |Diagonal| BW-From|   BW-To|   Level
             2|      79.83|      25|     209|     495|     524|       3
---------------------------------------------------------------------------
   495-  524 (35.49/36.17)        EQLSEhqcEGFLEayLLTGRHGIWSSYESF
   708-  732 (44.33/26.57)        EALTD...EEFTE..LFTADKPVLFAYHSY
---------------------------------------------------------------------------
---------------------------------------------------------------------------
No. of Repeats|Total Score|Length  |Diagonal| BW-From|   BW-To|   Level
             2|      56.80|      14|     490|     316|     329|       4
---------------------------------------------------------------------------
   316-  329 (30.34/18.19)        FRTPKGWTCPKFID
   807-  820 (26.45/14.95)        FAVDNGYDIPEFTD
---------------------------------------------------------------------------

>5C4I:A|PDBID|CHAIN|SEQUENCE
---------------------------------------------------------------------------
No. of Repeats|Total Score|Length  |Diagonal| BW-From|   BW-To|   Level
             3|     147.78|      34|      37|     178|     214|       1
---------------------------------------------------------------------------
   140-  174 (43.55/26.26)        ....PQEA.LDNTLIYYRvgeDQRVLL...PQYACLDGYFVSH
   178-  214 (56.35/45.55)        PvdiPDEAQVKEFLPPYK...NHHVLD...PRKPQIIGPQIEP
   218-  252 (47.87/30.63)        P...P..LQYQRYQAVKG...VHKVLEeacDEFARIFGRKYDP
---------------------------------------------------------------------------
---------------------------------------------------------------------------
No. of Repeats|Total Score|Length  |Diagonal| BW-From|   BW-To|   Level
             2|     196.95|      63|     205|      50|     118|       2
---------------------------------------------------------------------------
    50-  118 (100.05/91.97)
        DAE.FVHGEGEHAQlsvvyGASAAGARVFTGSSGVGVTyAMEVYSPISGERLPVQMA..IADRTLDPPGDFG
   261-  326 (96.90/71.50)
        DAEvIIFGQGAHME.....TAKAVARRLRNLGEKVGVA.RLRTFRPFPTEQIKERLSkfKAIGVLDVSANFG
---------------------------------------------------------------------------
>5C4I:B|PDBID|CHAIN|SEQUENCE
---------------------------------------------------------------------------
No. of Repeats|Total Score|Length  |Diagonal| BW-From|   BW-To|   Level
             2|      55.50|      17|      22|     153|     169|       2
---------------------------------------------------------------------------
   153-  169 (28.21/17.39)        AEGATDATGIGAGIAAP
```

```
  177-  193 (27.29/16.61)        ATGIVDVENLAAVVKNP
--------------------------------------------------------------------------------
>5C4I:C|PDBID|CHAIN|SEQUENCE
--------------------------------------------------------------------------------
No. of Repeats|Total Score|Length  |Diagonal| BW-From|   BW-To|   Level
             2|      82.61|      24|      32|      68|      98|       1
--------------------------------------------------------------------------------
   68-   92 (40.35/35.90)        PWIHAQITNGGAVASGIEAaYKAMI
  102-  125 (42.25/20.49)        PNIIVMAGDGGAVDIGLQA.LSAML
--------------------------------------------------------------------------------
--------------------------------------------------------------------------------
No. of Repeats|Total Score|Length  |Diagonal| BW-From|   BW-To|   Level
             2|      66.18|      16|      34|      12|      29|       2
--------------------------------------------------------------------------------
   12-   29 (30.78/20.04)        PDEEYYVPghRTCAGCGP
   49-   64 (35.40/16.92)        PTGCMYVA..NTSYGCGP
--------------------------------------------------------------------------------
--------------------------------------------------------------------------------
No. of Repeats|Total Score|Length  |Diagonal| BW-From|   BW-To|   Level
             2|     111.97|      32|      34|     154|     186|       3
--------------------------------------------------------------------------------
  154-  186 (54.35/33.17)        YGANTTFTPPGEVVPEGKKLFPKDNPKVIaHGH
  191-  222 (57.63/31.46)        YVATASIGWPVDLMNKVRKGLNQEGPAYI.HIH
--------------------------------------------------------------------------------

>3EY9:A|PDBID|CHAIN|SEQUENCE
--------------------------------------------------------------------------------
No. of Repeats|Total Score|Length  |Diagonal| BW-From|   BW-To|   Level
             2|     184.01|      57|     174|     214|     277|       1
--------------------------------------------------------------------------------
  214-  277 (86.30/76.10)        GAHKELVEFA.GKIKAPIVHALrGKEHVEYDNPYdVGMTGLIGFSsgfhtMMNADTLVLLGTQFP
  395-  452 (97.70/63.32)        GKRRLLGSFNhGSMANAMPQAL.GAQATEPERQV.VAMCGDGGFS.....MLMGDFLSVVQMKLP
--------------------------------------------------------------------------------
--------------------------------------------------------------------------------
No. of Repeats|Total Score|Length  |Diagonal| BW-From|   BW-To|   Level
             2|      86.26|      31|     174|     325|     369|       2
--------------------------------------------------------------------------------
  325-  357 (46.17/31.07)        EKADRkfLDKALE.DYR.DARKGLDDLAKPSEKAI
  502-  534 (40.09/18.69)        EKASE..VDEALQrAFSiDGPVLVDVVVAKEELAI
--------------------------------------------------------------------------------
--------------------------------------------------------------------------------
No. of Repeats|Total Score|Length  |Diagonal| BW-From|   BW-To|   Level
             2|      50.20|      12|     233|     116|     127|       4
--------------------------------------------------------------------------------
  116-  127 (25.78/17.02)        HPQELFRECSHY
  358-  369 (24.42/15.79)        HPQYLAQQISHF
--------------------------------------------------------------------------------

>4KXU:A|PDBID|CHAIN|SEQUENCE
--------------------------------------------------------------------------------
No. of Repeats|Total Score|Length  |Diagonal| BW-From|   BW-To|   Level
             2|      42.12|      14|      75|      85|     103|       1
--------------------------------------------------------------------------------
   88-  103 (18.92/25.16)        EAgfLAEAELLNLRKI
  514-  527 (23.20/ 9.53)        EA..LAAAELLKKEKI
--------------------------------------------------------------------------------
--------------------------------------------------------------------------------
No. of Repeats|Total Score|Length  |Diagonal| BW-From|   BW-To|   Level
             2|      51.12|      15|      75|     317|     338|       3
--------------------------------------------------------------------------------
  226-  240 (28.46/12.51)        KAFGQA....KHQPTAIIA
  319-  337 (22.66/19.15)        KAYGQAlaklGHASDRIIA
--------------------------------------------------------------------------------
--------------------------------------------------------------------------------
No. of Repeats|Total Score|Length  |Diagonal| BW-From|   BW-To|   Level
             2|     142.16|      42|     414|     160|     201|       4
--------------------------------------------------------------------------------
  160-  201 (74.46/49.60)        EGSVWEAMAFASIYKLDNLVAILDINRLGQSD.PAPLQHQMDI
  565-  607 (67.70/44.45)        EGGIGEAVSSAVVGEPGITVTHLAVNRVPRSGkPAELLKMFGI
--------------------------------------------------------------------------------
```

**T-REKS Jorda J, Kajava AV(2009). Bioinformatics 25 (20), 2632-2638**

```
repeat not found in sequence >2PAN:A|PDBID|CHAIN|SEQUENCE
repeat not found in sequence >2PGN:A|PDBID|CHAIN|SEQUENCE
repeat not found in sequence >6A50:A|PDBID|CHAIN|SEQUENCE
repeat not found in sequence >4QQ8:A|PDBID|CHAIN|SEQUENCE
repeat not found in sequence >2NXW:A|PDBID|CHAIN|SEQUENCE
repeat not found in sequence >1NI4:A|PDBID|CHAIN|SEQUENCE
repeat not found in sequence >1NI4:B|PDBID|CHAIN|SEQUENCE
repeat not found in sequence >3AHC:A|PDBID|CHAIN|SEQUENCE
repeat not found in sequence >5C4I:A|PDBID|CHAIN|SEQUENCE
repeat not found in sequence >5C4I:B|PDBID|CHAIN|SEQUENCE
repeat not found in sequence >5C4I:C|PDBID|CHAIN|SEQUENCE
>3EY9:A|PDBID|CHAIN|SEQUENCE
Length: 7 residues - nb: 2  from  27 to 40 - Psim:0.7857142857142857 region Length:14
DSLNGLS
DSLNRMG
**********************
repeat not found in sequence >4KXU:A|PDBID|CHAIN|SEQUENCE
1 sequences have been detected as tandem repeats containing.
```

**HHREPID Biegert A., Söding J. (2008) HHrepID: de novo protein repeat identification by probabilistic consistency. Bioinformatics 24(6):807-814**

```
>2PAN:A|PDBID|CHAIN|SEQUENCE
No. of repeats: 2
P-value: 1.2E-09
Length:
ID      Prob    P-val   Loc     Sequence
A1      63.01   3.7e-03 463-483 -VVAISGDFDFQFLIEELAVGAqfnip
A2      51.59   1.5e-03 489-501 YIHVLVNNAYLGL---------.....

>2PGN:A|PDBID|CHAIN|SEQUENCE
No. of repeats: 2
P-value: 2.1E-05
Length:
ID      Prob    P-val   Loc     Sequence
A1      26.72   1.8e-02 9-17    -------LIVEALEEY-------gteqvvg
A2      52.10   7.9e-03 25-47   FIGHTSHFVADAFSKSHLGKRVI.......

No. of repeats: 2
P-value: 6.6E-06
Length:
ID      Prob    P-val   Loc     Sequence
B1      54.44   5.0e-03 445-464 -VFLGTGDGALYYHFNEFRVAvehklp
B2      39.01   1.4e-02 471-483 VITMVFTNESYGA-------......

>6A50:A|PDBID|CHAIN|SEQUENCE
No. of repeats: 2
P-value: 7.6E-08
Length:
ID      Prob    P-val   Loc     Sequence
A1      58.71   2.9e-03 442-462 -QVIAVIGDGSANYSISALWTAaqyni
A2      42.83   3.3e-03 468-481 PTIFVIMNNGTYGM--------.....

>4QQ8:A|PDBID|CHAIN|SEQUENCE
No. of repeats: 3
P-value: 2.3E-05
Length:
ID      Prob    P-val   Loc     Sequence
A1      66.27   1.0e-03 8-27    ---ELVVRTLIKAGVEHLFGLHG
A2      63.01   8.5e-03 28-50   IHIDTIFQACLDHDVPIIDTRHE
A3      13.94   9.3e-01 51-62   AAAGHAAEGYAR-----------

No. of repeats: 2
P-value: 3.9E-05
Length:
ID      Prob    P-val   Loc     Sequence
B1      49.19   1.4e-02 442-461 -TILVTGDGSVGYSIGEFDTLvrkqlp
B2      35.80   6.1e-03 468-480 LIVIIMNNQSWGW--------......

>2NXW:A|PDBID|CHAIN|SEQUENCE
No. of repeats: 2
P-value: 9.5E-06
Length:
```

```
ID      Prob    P-val   Loc     Sequence
A1      60.66   8.2e-03 443-462 -ILTVVGDGAFQMTGWELGNCrrlgid
A2      42.17   3.5e-03 469-481 PIVILFNNASWEM-------......

>1NI4:A|PDBID|CHAIN|SEQUENCE
none
>1NI4:B|PDBID|CHAIN|SEQUENCE
none

>3AHC:A|PDBID|CHAIN|SEQUENCE
none

>5C4I:A|PDBID|CHAIN|SEQUENCE
none

>5C4I:B|PDBID|CHAIN|SEQUENCE
No. of repeats: 2
P-value: 1.1E-06
Length:
ID      Prob    P-val   Loc     Sequence
A1      85.97   3.3e-11 255-281 IIDREACTECYTCWIYCPDSCITRTEEgp
A2      87.01   6.2e-11 284-307 VFNMKYCKGCGLCTAVCPSGALTN---..

>5C4I:C|PDBID|CHAIN|SEQUENCE
none

>3EY9:A|PDBID|CHAIN|SEQUENCE
No. of repeats: 2
P-value: 6.8E-07
Length:
ID      Prob    P-val   Loc     Sequence
A1      55.91   1.6e-02 59-79   ---AEAQLSGELAVCAGSCGPGNLhli
A2      61.91   1.2e-03 83-101  NGLFDCHRNHVPVLAIAAH-----...

No. of repeats: 2
P-value: 0.0033
Length:
ID      Prob    P-val   Loc     Sequence
B1      16.39   6.0e-02 8-16    ------YIAKTLESA-------gvkriwgv
B2      58.50   2.0e-03 25-46   TGDSLNGLSDSLNRMGTIEWMS........

No. of repeats: 2
P-value: 0.0019
Length:
ID      Prob    P-val   Loc     Sequence
C1      54.22   3.1e-03 427-446 -VVAMCGDGGFSMLMGDFLSVvqmklp
C2      39.29   4.3e-03 453-465 VKIVVFNNSVLGF--------......

>4KXU:A|PDBID|CHAIN|SEQUENCE
none
```

**XSTREAM  BMC Bioinformatics 2007, 8:382**

```
>2PAN:A|PDBID|CHAIN|SEQUENCE
none

>2PGN:A|PDBID|CHAIN|SEQUENCE
none

>6A50:A|PDBID|CHAIN|SEQUENCE
Positions>

Period

Copy
Number

Consensus
Error
494-503

5

2.00
```

```
0.10

VPGLD
VPGID
=====
VPGID

>4QQ8:A|PDBID|CHAIN|SEQUENCE
none

>2NXW:A|PDBID|CHAIN|SEQUENCE
none

>1NI4:A|PDBID|CHAIN|SEQUENCE
None

>1NI4:B|PDBID|CHAIN|SEQUENCE
None

>3AHC:A|PDBID|CHAIN|SEQUENCE
None

>5C4I:A|PDBID|CHAIN|SEQUENCE
None

>5C4I:B|PDBID|CHAIN|SEQUENCE
None

>5C4I:C|PDBID|CHAIN|SEQUENCE
None

>3EY9:A|PDBID|CHAIN|SEQUENCE
None

>4KXU:A|PDBID|CHAIN|SEQUENCE
none
```

**TRUST ISMB/ECCB 2004 conference (Glasgow, UK), appeared in Bioinformatics. 2004 Aug 4;20 Suppl 1:i311-i317**

```
>2PAN:A|PDBID|CHAIN|SEQUENCE
none

>2PGN:A|PDBID|CHAIN|SEQUENCE
none

>6A50:A|PDBID|CHAIN|SEQUENCE
none

>4QQ8:A|PDBID|CHAIN|SEQUENCE
none

>2NXW:A|PDBID|CHAIN|SEQUENCE
none

>1NI4:A|PDBID|CHAIN|SEQUENCE
None

>1NI4:B|PDBID|CHAIN|SEQUENCE
None

>3AHC:A|PDBID|CHAIN|SEQUENCE
None

>5C4I:A|PDBID|CHAIN|SEQUENCE
None

>5C4I:B|PDBID|CHAIN|SEQUENCE
Repeat type 1
id      sequence        start   size
1       RIQR-PIIDREACTECYTCWIYCPDSCIT    250     28
2       RTEEgPVFNMKYCKGCGLCTAVCPSGALT    278     29
```

```
>5C4I:C|PDBID|CHAIN|SEQUENCE
none

>3EY9:A|PDBID|CHAIN|SEQUENCE
None

>4KXU:A|PDBID|CHAIN|SEQUENCE
none

>4KXU:A|PDBID|CHAIN|SEQUENCE
```

**Supplemental Data S2:** Table listing proteins in the PDB found by searching for at least 3 copies of a ThDP repeat pattern. Known ThDP proteins are indented and preceded by an asterisk.

| | |
|---|---|
| 1a9x | CARBAMOYL PHOSPHATE SYNTHETASE: CAUGHT IN THE ACT OF GLUTAMINE |
| 1ami | STERIC AND CONFORMATIONAL FEATURES OF THE ACONITASE MECHANISM |
| 1aor | STRUCTURE OF A HYPERTHERMOPHILIC TUNGSTOPTERIN ENZYME, |
| 1azy | STRUCTURAL AND THEORETICAL STUDIES SUGGEST DOMAIN MOVEMENT PRODUCES AN |
| * 1b0p | CRYSTAL STRUCTURE OF PYRUVATE-FERREDOXIN OXIDOREDUCTASE FROM |
| 1b1x | STRUCTURE OF DIFERRIC MARE LACTOFERRIN AT 2.62A RESOLUTION |
| 1b7u | STRUCTURE OF MARE APOLACTOFERRIN: THE N AND C LOBES ARE IN |
| 1b7z | STRUCTURE OF OXALATE SUBSTITUTED DIFERRIC MARE LACTOFERRIN FROM |
| 1ba2 | D67R MUTANT OF D-RIBOSE-BINDING PROTEIN FROM ESCHERICHIA |
| 1bxb | XYLOSE ISOMERASE FROM THERMUS THERMOPHILUS |
| 1bxc | XYLOSE ISOMERASE FROM THERMUS CALDOPHILUS |
| 1bxr | STRUCTURE OF CARBAMOYL PHOSPHATE SYNTHETASE COMPLEXED WITH THE ATP |
| 1c3f | ENDO-BETA-N-ACETYLGLUCOSAMINIDASE H, D130N MUTANT |
| 1c3o | CRYSTAL STRUCTURE OF THE CARBAMOYL PHOSPHATE SYNTHETASE: SMALL SUBUNIT |
| 1c4g | PHOSPHOGLUCOMUTASE VANADATE BASED TRANSITION STATE ANALOG COMPLEX |
| 1c8x | ENDO-BETA-N-ACETYLGLUCOSAMINIDASE H, D130E MUTANT |
| 1c8y | ENDO-BETA-N-ACETYLGLUCOSAMINIDASE H, D130A MUTANT |
| 1c90 | ENDO-BETA-N-ACETYLGLUCOSAMINIDASE H, E132Q MUTANT |
| 1c91 | ENDO-BETA-N-ACETYLGLUCOSAMINIDASE H, E132D |
| 1c92 | ENDO-BETA-N-ACETYLGLUCOSAMINIDASE H, E132A MUTANT |
| 1c93 | ENDO-BETA-N-ACETYLGLUCOSAMINIDASE H, D130N/E132Q DOUBLE MUTANT |
| 1c96 | S642A:CITRATE COMPLEX OF ACONITASE |
| 1ce8 | CARBAMOYL PHOSPHATE SYNTHETASE FROM ESCHERICHIS COLI WITH COMPLEXED |
| 1cqi | CRYSTAL STRUCTURE OF THE COMPLEX OF ADP AND MG2+ WITH DEPHOSPHORYLATED |
| 1cqj | CRYSTAL STRUCTURE OF DEPHOSPHORYLATED E. COLI SUCCINYL-COA SYNTHETASE |
| 1cs0 | CRYSTAL STRUCTURE OF CARBAMOYL PHOSPHATE SYNTHETASE COMPLEXED AT |
| 1d0n | THE CRYSTAL STRUCTURE OF CALCIUM-FREE EQUINE PLASMA GELSOLIN. |
| 1drj | PROBING PROTEIN-PROTEIN INTERACTIONS: THE RIBOSE-BINDING PROTEIN IN |
| 1drk | PROBING PROTEIN-PROTEIN INTERACTIONS: THE RIBOSE-BINDING PROTEIN IN |
| 1ea0 | ALPHA SUBUNIT OF A. BRASILENSE GLUTAMATE SYNTHASE |
| 1edt | CRYSTAL STRUCTURE OF ENDO-BETA-N-ACETYLGLUCOSAMINIDASE H AT |
| 1exp | BETA-1,4-GLYCANASE CEX-CD |
| 1f9b | MELANIN PROTEIN INTERACTION: X-RAY STRUCTURE OF THE COMPLEX OF MARE |
| 1ffu | CARBON MONOXIDE DEHYDROGENASE FROM HYDROGENOPHAGA |
| 1ffv | CARBON MONOXIDE DEHYDROGENASE FROM HYDROGENOPHAGA |
| 1fgh | COMPLEX WITH 4-HYDROXY-TRANS-ACONITATE |
| 1fh7 | CRYSTAL STRUCTURE OF THE XYLANASE CEX WITH XYLOBIOSE- |
| 1fh8 | CRYSTAL STRUCTURE OF THE XYLANASE CEX WITH XYLOBIOSE-DERIVED |
| 1fh9 | CRYSTAL STRUCTURE OF THE XYLANASE CEX WITH XYLOBIOSE-DERIVED LACTAM |
| 1fhd | CRYSTAL STRUCTURE OF THE XYLANASE CEX WITH XYLOBIOSE-DERIVED IMIDAZOLE |
| 1g8m | CRYSTAL STRUCTURE OF AVIAN ATIC, A BIFUNCTIONAL TRANSFORMYLASE AND |
| 1gc0 | CRYSTAL STRUCTURE OF THE PYRIDOXAL-5'-PHOSPHATE DEPENDENT L-METHIONINE |
| 1gca | THE 1.7 ANGSTROMS REFINED X-RAY STRUCTURE OF THE PERIPLASMIC |
| 1gcg | THE 1.9 ANGSTROMS X-RAY STRUCTURE OF A CLOSED UNLIGANDED FORM OF THE |
| 1glg | CRYSTALLOGRAPHIC ANALYSIS OF THE EPIMERIC AND ANOMERIC |
| 1gq2 | MALIC ENZYME FROM PIGEON LIVER |
| 1gub | HINGE-BENDING MOTION OF D-ALLOSE BINDING PROTEIN FROM ESCHERICHIA |
| 1hcu | ALPHA-1,2-MANNOSIDASE FROM TRICHODERMA REESEI |
| 1ir2 | CRYSTAL STRUCTURE OF ACTIVATED RIBULOSE-1,5-BISPHOSPHATE |
| 1iwg | CRYSTAL STRUCTURE OF BACTERIAL MULTIDRUG EFFLUX TRANSPORTER ACRB |
| 1j01 | CRYSTAL STRUCTURE OF THE XYLANASE CEX WITH XYLOBIOSE-DERIVED INHIBITOR |
| 1j3n | CRYSTAL STRUCTURE OF 3-OXOACYL-(ACYL-CARRIER PROTEIN) |
| 1jdb | CARBAMOYL PHOSPHATE SYNTHETASE FROM ESCHERICHIA COLI |
| 1jkj | E. COLI SCS |
| 1kee | INACTIVATION OF THE AMIDOTRANSFERASE ACTIVITY OF CARBAMOYL PHOSPHATE |
| 1kek | CRYSTAL STRUCTURE OF THE FREE RADICAL INTERMEDIATE OF |
| 1kp8 | STRUCTURAL BASIS FOR GROEL-ASSISTED PROTEIN FOLDING FROM THE CRYSTAL |
| 1l1l | CRYSTAL STRUCTURE OF B-12 DEPENDENT (CLASS II) |
| 1leh | LEUCINE DEHYDROGENASE FROM BACILLUS SPHAERICUS |
| 1m6v | CRYSTAL STRUCTURE OF THE G359F (SMALL SUBUNIT) POINT MUTANT OF |
| 1m9n | CRYSTAL STRUCTURE OF THE HOMODIMERIC BIFUNCTIONAL TRANSFORMYLASE AND |
| 1mnf | DOMAIN MOTIONS IN GROEL UPON BINDING OF AN OLIGOPEPTIDE |
| 1mum | STRUCTURE OF THE 2-METHYLISOCITRATE LYASE (PRPB) FROM ESCHERICHIA COLI |
| 1n5w | CRYSTAL STRUCTURE OF THE CU,MO-CO DEHYDROGENASE (CODH); OXIDIZED FORM |
| 1n60 | CRYSTAL STRUCTURE OF THE CU,MO-CO DEHYDROGENASE (CODH); CYANIDE- |
| 1n61 | CRYSTAL STRUCTURE OF THE CU,MO-CO DEHYDROGENASE (CODH); DITHIONITE |
| 1n63 | CRYSTAL STRUCTURE OF THE CU,MO-CO DEHYDROGENASE (CODH); CARBON |
| 1nbw | GLYCEROL DEHYDRATASE REACTIVASE |

1np2 CRYSTAL STRUCTURE OF THERMOSTABLE BETA-GLYCOSIDASE FROM
1o68 CRYSTAL STRUCTURE OF 3-METHYL-2-OXOBUTANOATE HYDROXYMETHYLTRANSFERASE
1o7t METAL NANOCLUSTERS BOUND TO THE FERRIC BINDING PROTEIN FROM NEISSERIA
1otp STRUCTURAL AND THEORETICAL STUDIES SUGGEST DOMAIN MOVEMENT PRODUCES AN
1oy8 STRUCTURAL BASIS OF MULTIPLE DRUG BINDING CAPACITY OF THE ACRB
1oz0 CRYSTAL STRUCTURE OF THE HOMODIMERIC BIFUNCTIONAL
* 1ozf THE CRYSTAL STRUCTURE OF KLEBSIELLA PNEUMONIAE ACETOLACTATE SYNTHASE
* 1ozg THE CRYSTAL STRUCTURE OF KLEBSIELLA PNEUMONIAE ACETOLACTATE SYNTHASE
* 1ozh THE CRYSTAL STRUCTURE OF KLEBSIELLA PNEUMONIAE ACETOLACTATE
1p4r CRYSTAL STRUCTURE OF HUMAN ATIC IN COMPLEX WITH FOLATE-
1pg8 CRYSTAL STRUCTURE OF L-METHIONINE ALPHA-, GAMMA-LYASE
1pj5 CRYSTAL STRUCTURE OF DIMETHYLGLYCINE OXIDASE OF
1pj6 CRYSTAL STRUCTURE OF DIMETHYLGLYCINE OXIDASE OF ARTHROBACTER
1pj7 STRUCTURE OF DIMETHYLGLYCINE OXIDASE OF ARTHROBACTER GLOBIFORMIS IN
1pkx CRYSTAL STRUCTURE OF HUMAN ATIC IN COMPLEX WITH XMP
1pl0 CRYSTAL STRUCTURE OF HUMAN ATIC IN COMPLEX WITH FOLATE-
* 1pow THE REFINED STRUCTURES OF A STABILIZED MUTANT AND OF WILD-TYPE
1q3g MURA (ASP305ALA) LIGANDED WITH TETRAHEDRAL REACTION INTERMEDIATE
1qjd FLAVOCYTOCHROME C3 FROM SHEWANELLA FRIGIDIMARINA
1qjm CRYSTAL STRUCTURE OF A COMPLEX OF LACTOFERRIN WITH A LANTHANIDE ION
1qpq STRUCTURE OF QUINOLINIC ACID PHOSPHORIBOSYLTRANSFERASE FROM
1qpr QUINOLINATE PHOSPHORIBOSYLTRANSFERASE (QAPRTASE) FROM
1qvr CRYSTAL STRUCTURE ANALYSIS OF CLPB
1r0m STRUCTURE OF DEINOCOCCUS RADIODURANS N-ACYLAMINO ACID
1r1n TRI-NUCLEAR OXO-IRON CLUSTERS IN THE FERRIC BINDING PROTEIN
1rbl STRUCTURE DETERMINATION AND REFINEMENT OF RIBULOSE 1,5 BISPHOSPHATE
1rco SPINACH RUBISCO IN COMPLEX WITH THE INHIBITOR D-XYLULOSE-2,
1rcx NON-ACTIVATED SPINACH RUBISCO IN COMPLEX WITH ITS SUBSTRATE
1rf4 STRUCTURAL STUDIES OF STREPTOCOCCUS PNEUMONIAE EPSP
1rf5 STRUCTURAL STUDIES OF STREPTOCOCCUS PNEUMONIAE EPSP
1rlu MYCOBACTERIUM TUBERCULOSIS FTSZ IN COMPLEX WITH GTP-GAMMA-S
1rpj CRYSTAL STRUCTURE OF D-ALLOSE BINDING PROTEIN FROM ESCHERICHIA COLI
1rq2 MYCOBACTERIUM TUBERCULOSIS FTSZ IN COMPLEX WITH CITRATE
1rq7 MYCOBACTERIUM TUBERCULOSIS FTSZ IN COMPLEX WITH GDP
1rsc STRUCTURE OF AN EFFECTOR INDUCED INACTIVATED STATE OF RIBULOSE
1scu THE CRYSTAL STRUCTURE OF SUCCINYL-COA SYNTHETASE FROM ESCHERICHIA COLI
1ss8 GROEL
1svd THE STRUCTURE OF HALOTHIOBACILLUS NEAPOLITANUS RUBISCO
1svt CRYSTAL STRUCTURE OF GROEL14-GROES7-(ADP-ALFX)7
1sx3 GROEL14-(ATPGAMMAS)14
1sx4 GROEL-GROES-ADP7
1t36 CRYSTAL STRUCTURE OF E. COLI CARBAMOYL PHOSPHATE SYNTHETASE SMALL
* 1t9a CRYSTAL STRUCTURE OF YEAST ACETOHYDROXYACID SYNTHASE IN COMPLEX WITH A
1thz CRYSTAL STRUCTURE OF AVIAN AICAR TRANSFORMYLASE IN COMPLEX
1u7h STRUCTURE AND A PROPOSED MECHANISM FOR ORNITHINE
1uls CRYSTAL STRUCTURE OF TT0140 FROM THERMUS THERMOPHILUS HB8
1uoz STRUCTURE OF THE ENDOGLUCANASE CEL6 FROM MYCOBACTERIUM
1up0 STRUCTURE OF THE ENDOGLUCANASE CEL6 FROM MYCOBACTERIUM
1up2 STRUCTURE OF THE ENDOGLUCANASE CEL6 FROM MYCOBACTERIUM
1up3 STRUCTURE OF THE ENDOGLUCANASE CEL6 FROM MYCOBACTERIUM
* 1upa CARBOXYETHYLARGININE SYNTHASE FROM STREPTOMYCES
* 1upb CARBOXYETHYLARGININE SYNTHASE FROM STREPTOMYCES
* 1upc CARBOXYETHYLARGININE SYNTHASE FROM STREPTOMYCES
1upm ACTIVATED SPINACH RUBISCO COMPLEXED WITH
1uqt TREHALOSE-6-PHOSPHATE FROM E. COLI BOUND WITH UDP-2-FLUORO GLUCOSE.
1uqu TREHALOSE-6-PHOSPHATE FROM E. COLI BOUND WITH UDP-GLUCOSE.
1urp D-RIBOSE-BINDING PROTEIN FROM ESCHERICHIA COLI
1us2 XYLANASE10C (MUTANT E385A) FROM CELLVIBRIO JAPONICUS IN COMPLEX WITH
1us3 NATIVE XYLANASE10C FROM CELLVIBRIO JAPONICUS
1uw9 L290F-A222T CHLAMYDOMONAS RUBISCO MUTANT
1uwa L290F MUTANT RUBISCO FROM CHLAMYDOMONAS
1uzd CHLAMYDOMONAS,SPINACH CHIMERIC RUBISCO
1uzh A CHIMERIC CHLAMYDOMONAS, SYNECHOCOCCUS RUBISCO ENZYME
1uzl MABA FROM MYCOBACTERIUM TUBERCULOSIS
1uzm MABA FROM MYCOBACTERIUM TUBERCULOSIS
1uzn MABA FROM MYCOBACTERIUM TUBERCULOSIS
1w46 P4 PROTEIN FROM BACTERIOPHAGE PHI12 IN COMPLEX WITH ADP AND MG
1w48 P4 PROTEIN FROM BACTERIOPHAGE PHI12 IN COMPLEX WITH AMPCPP
1w4a P4 PROTEIN FROM PHI12 IN COMPLEX WITH AMPCPP AND MN
1wcv STRUCTURE OF THE BACTERIAL CHROMOSOME SEGREGATION PROTEIN SOJ
1wdd CRYSTAL STRUCTURE OF ACTIVATED RICE RUBISCO COMPLEXED WITH 2-
1wdl FATTY ACID BETA-OXIDATION MULTIENZYME COMPLEX FROM
1wmb CRYSTAL STRUCTURE OF NAD DEPENDENT D-3-HYDROXYBUTYLATE DEHYDROGENASE
1wyt CRYSTAL STRUCTURE OF GLYCINE DECARBOXYLASE (P-PROTEIN) OF THE GLYCINE

2926 1wyu CRYSTAL STRUCTURE OF GLYCINE DECARBOXYLASE (P-PROTEIN) OF THE GLYCINE
2927 1wyv CRYSTAL STRUCTURE OF GLYCINE DECARBOXYLASE (P-PROTEIN) OF THE GLYCINE
2928 1x1t CRYSTAL STRUCTURE OF D-3-HYDROXYBUTYRATE DEHYDROGENASE FROM
2929 1xc1 OXO ZIRCONIUM(IV) CLUSTER IN THE FERRIC BINDING PROTEIN
2930 1xck CRYSTAL STRUCTURE OF APO GROEL
2931 1xg3 CRYSTAL STRUCTURE OF THE C123S 2-METHYLISOCITRATE LYASE
2932 1xg4 CRYSTAL STRUCTURE OF THE C123S 2-METHYLISOCITRATE LYASE
2933 1xpy STRUCTURAL BASIS FOR CATALYTIC RACEMIZATION AND SUBSTRATE
2934 1xr4 X-RAY CRYSTAL STRUCTURE OF PUTATIVE CITRATE LYASE ALPHA CHAIN/CITRATE-
2935 1xs2 STRUCTURAL BASIS FOR CATALYTIC RACEMIZATION AND SUBSTRATE
2936 * 1y9d PYRUVATE OXIDASE VARIANT V265A FROM LACTOBACILLUS PLANTARUM
2937 * 1ybh CRYSTAL STRUCTURE OF ARABIDOPSIS THALIANA ACETOHYDROXYACID SYNTHASE IN
2938 1ygy CRYSTAL STRUCTURE OF D-3-PHOSPHOGLYCERATE DEHYDROGENASE FROM
2939 * 1yi1 CRYSTAL STRUCTURE OF ARABIDOPSIS THALIANA ACETOHYDROXYACID SYNTHASE IN
2940 1yt8 CRYSTAL STRUCTURE OF THIOSULFATE SULFURTRANSFERASE FROM PSEUDOMONAS
2941 1zlp PETAL DEATH PROTEIN PSR132 WITH CYSTEINE-LINKED GLUTARALDEHYDE FORMING
2942 1zxi RECONSTITUTED CO DEHYDROGENASE FROM OLIGOTROPHA CARBOXIDOVORANS
2943 2a9f CRYSTAL STRUCTURE OF A PUTATIVE MALIC ENZYME ((S)-MALATE:NAD+
2944 * 2ag0 CRYSTAL STRUCTURE OF BENZALDEHYDE LYASE (BAL)- NATIVE
2945 * 2ag1 CRYSTAL STRUCTURE OF BENZALDEHYDE LYASE (BAL)- SEMET
2946 2ahv CRYSTAL STRUCTURE OF ACYL-COA TRANSFERASE FROM E. COLI O157:H7 (YDIF)-
2947 2ahw CRYSTAL STRUCTURE OF ACYL-COA TRANSFERASE FROM E. COLI O157:H7 (YDIF)-
2948 2b1g CRYSTAL STRUCTURES OF TRANSITION STATE ANALOGUE INHIBITORS OF INOSINE
2949 2b1i CRYSTAL STRUCTURES OF TRANSITION STATE ANALOGUE INHIBITORS OF INOSINE
2950 2bej STRUCTURE OF THE BACTERIAL CHROMOSOME SEGREGATION PROTEIN SOJ
2951 2bek STRUCTURE OF THE BACTERIAL CHROMOSOME SEGREGATION PROTEIN SOJ
2952 * 2c31 CRYSTAL STRUCTURE OF OXALYL-COA DECARBOXYLASE IN COMPLEX
2953 * 2c3m CRYSTAL STRUCTURE OF PYRUVATE-FERREDOXIN OXIDOREDUCTASE
2954 * 2c3o CRYSTAL STRUCTURE OF THE FREE RADICAL INTERMEDIATE OF
2955 * 2c3p CRYSTAL STRUCTURE OF THE FREE RADICAL INTERMEDIATE OF
2956 * 2c3u CRYSTAL STRUCTURE OF PYRUVATE-FERREDOXIN OXIDOREDUCTASE
2957 * 2c3y CRYSTAL STRUCTURE OF THE RADICAL FORM OF
2958 * 2c42 CRYSTAL STRUCTURE OF PYRUVATE-FERREDOXIN OXIDOREDUCTASE
2959 2c7c FITTED COORDINATES FOR GROEL-ATP7-GROES CRYO-EM COMPLEX (EMD-1180)
2960 2c7d FITTED COORDINATES FOR GROEL-ADP7-GROES CRYO-EM COMPLEX (EMD-1181)
2961 2c7e REVISED ATOMIC STRUCTURE FITTING INTO A GROEL(D398A)-ATP7 CRYO-EM MAP
2962 2cfc STRUCTURAL BASIS FOR STEREO SELECTIVITY IN THE (R)- AND
2963 2csu CRYSTAL STRUCTURE OF PH0766 FROM PYROCOCCUS HORIKOSHII OT3
2964 2d0o STRCUTURE OF DIOL DEHYDRATASE-REACTIVATING FACTOR COMPLEXED
2965 2d0p STRCUTURE OF DIOL DEHYDRATASE-REACTIVATING FACTOR IN
2966 2dkh CRYSTAL STRUCTURE OF 3-HYDROXYBENZOATE HYDROXYLASE FROM COMAMONAS
2967 2dki CRYSTAL STRUCTURE OF 3-HYDROXYBENZOATE HYDROXYLASE FROM COMAMONAS
2968 2dq4 CRYSTAL STRUCTURE OF THREONINE 3-DEHYDROGENASE
2969 2dr6 CRYSTAL STRUCTURE OF A MULTIDRUG TRANSPORTER REVEAL A
2970 2drd CRYSTAL STRUCTURE OF A MULTIDRUG TRANSPORTER REVEAL A
2971 2dri PROBING PROTEIN-PROTEIN INTERACTIONS: THE RIBOSE BINDING PROTEIN IN
2972 2dsj CRYSTAL STRUCTURE OF PROJECT ID TT0128 FROM THERMUS THERMOPHILUS HB8
2973 2ejv CRYSTAL STRUCTURE OF THREONINE 3-DEHYDROGENASE COMPLEXED WITH NAD+
2974 2exo CRYSTAL STRUCTURE OF THE CATALYTIC DOMAIN OF THE BETA-1,4-
2975 * 2ez4 PYRUVATE OXIDASE VARIANT F479W
2976 * 2ez8 PYRUVATE OXIDASE VARIANT F479W IN COMPLEX WITH REACTION INTERMEDIATE
2977 * 2ez9 PYRUVATE OXIDASE VARIANT F479W IN COMPLEX WITH REACTION INTERMEDIATE
2978 * 2ezt PYRUVATE OXIDASE VARIANT F479W IN COMPLEX WITH REACTION INTERMEDIATE
2979 * 2ezu PYRUVATE OXIDASE VARIANT F479W IN COMPLEX WITH REACTION INTERMEDIATE
2980 2fep STRUCTURE OF TRUNCATED CCPA IN COMPLEX WITH P-SER-HPR AND
2981 2fgh ATP BOUND GELSOLIN
2982 2fkp THE MUTANT G127C-T313C OF DEINOCOCCUS RADIODURANS N-
2983 2fvy HIGH RESOLUTION GLUCOSE BOUND CRYSTAL STRUCTURE OF GGBP
2984 2fw0 APO OPEN FORM OF GLUCOSE/GALACTOSE BINDING PROTEIN
2985 2gbp SUGAR AND SIGNAL-TRANSDUCER BINDING SITES OF THE ESCHERICHIA COLI
2986 2ggg THE MUTANT A68C-D72C OF DEINOCOCCUS RADIODURANS N-ACYLAMINO
2987 2ggh THE MUTANT A68C-D72C-NLQ OF DEINOCOCCUS RADIODURANS
2988 2ggi THE MUTANT E149C-A182C OF DEINOCOCCUS RADIODURANS N-
2989 2ggj THE MUTANT Y218C OF DEINOCOCCUS RADIODURANS N-ACYLAMINO
2990 2gif ASYMMETRIC STRUCTURE OF TRIMERIC ACRB FROM ESCHERICHIA COLI
2991 2gl5 CRYSTAL STRUCTURE OF PUTATIVE DEHYDRATASE FROM SALMONELLA THYPHIMURIUM
2992 2grv CRYSTAL STRUCTURE OF LPQW
2993 2gx6 RATIONAL STABILIZATION OF E. COLI RIBOSE BINDING PROTEIN
2994 2hg4 STRUCTURE OF THE KETOSYNTHASE-ACYLTRANSFERASE DIDOMAIN OF MODULE 5
2995 2his CELLULOMONAS FIMI XYLANASE/CELLULASE DOUBLE MUTANT
2996 2hjg THE CRYSTAL STRUCTURE OF THE B. SUBTILIS YPHC GTPASE IN COMPLEX WITH
2997 2hk0 CRYSTAL STRUCTURE OF D-PSICOSE 3-EPIMERASE (DPEASE) IN THE ABSENCE OF
2998 2hk1 CRYSTAL STRUCTURE OF D-PSICOSE 3-EPIMERASE (DPEASE) IN THE PRESENCE OF
2999 2hph HIGH RESOLUTION STRUCTURE OF E. COLI GLUCOSE/GALACTOSE

3000 2hqg CONFORMATION OF THE ACRB MULTIDRUG EFFLUX PUMP IN MUTANTS OF THE
3001 2hrt ASYMMETRIC STRUCTURE OF TRIMERIC ACRB FROM ESCHERICHIA COLI
3002 2hzg CRYSTAL STRUCTURE OF PREDICTED MANDELATE RACEMASE FROM RHODOBACTER
3003 2i56 CRYSTAL STRUCTURE OF L-RHAMNOSE ISOMERASE FROM PSEUDOMONAS
3004 2i6w CRYSTAL STRUCTURE OF THE MULTIDRUG EFFLUX TRANSPORTER ACRB
3005 * 2iht CARBOXYETHYLARGININE SYNTHASE FROM STREPTOMYCES CLAVULIGERUS: SEMET
3006 * 2ihu CARBOXYETHYLARGININE SYNTHASE FROM STREPTOMYCES CLAVULIGERUS: PUTATIVE
3007 * 2ihv CARBOXYETHYLARGININE SYNTHASE FROM STREPTOMYCES CLAVULIGERUS: 5-
3008 2ipl CRYSTAL STRUCTURE OF A DISULFIDE MUTANT GLUCOSE BINDING PROTEIN
3009 2ipm CRYSTAL STRUCTURE OF A DISULFIDE MUTANT GLUCOSE BINDING PROTEIN
3010 2ipn CRYSTAL STRUCTURE OF A DISULFIDE MUTANT GLUCOSE BINDING PROTEIN
3011 2iu0 CRYSTAL STRUCTURES OF TRANSITION STATE ANALOGUE INHIBITORS OF INOSINE
3012 2iu3 CRYSTAL STRUCTURES OF TRANSITION STATE ANALOGUE INHIBITORS OF INOSINE
3013 2ivd STRUCTURE OF PROTOPORPHYRINOGEN OXIDASE FROM MYXOCOCCUS
3014 2ive STRUCTURE OF PROTOPORPHYRINOGEN OXIDASE FROM MYXOCOCCUS
3015 2j8s DRUG EXPORT PATHWAY OF MULTIDRUG EXPORTER ACRB REVEALED BY
3016 * 2ji6 X-RAY STRUCTURE OF OXALYL-COA DECARBOXYLASE IN COMPLEX WITH 3-DEAZA-
3017 * 2ji7 X-RAY STRUCTURE OF OXALYL-COA DECARBOXYLASE WITH COVALENT
3018 * 2ji8 X-RAY STRUCTURE OF OXALYL-COA DECARBOXYLASE IN COMPLEX WITH
3019 * 2ji9 X-RAY STRUCTURE OF OXALYL-COA DECARBOXYLASE IN COMPLEX WITH
3020 * 2jib X-RAY STRUCTURE OF OXALYL-COA DECARBOXYLASE IN COMPLEX WITH
3021 * 2jla CRYSTAL STRUCTURE OF E.COLI MEND, 2-SUCCINYL-5-ENOLPYRUVYL-
3022 * 2jlc CRYSTAL STRUCTURE OF E.COLI MEND, 2-SUCCINYL-5-ENOLPYRUVYL-
3023 2l26 RV0899 FROM MYCOBACTERIUM TUBERCULOSIS CONTAINS TWO SEPARATED DOMAINS
3024 2ntn CRYSTAL STRUCTURE OF MABA-C60V/G139A/S144L
3025 2nu6 C123AA MUTANT OF E. COLI SUCCINYL-COA SYNTHETASE
3026 2nu7 C123AS MUTANT OF E. COLI SUCCINYL-COA SYNTHETASE
3027 2nu8 C123AT MUTANT OF E. COLI SUCCINYL-COA SYNTHETASE
3028 2nu9 C123AT MUTANT OF E. COLI SUCCINYL-COA SYNTHETASE
3029 2nua C123AV MUTANT OF E. COLI SUCCINYL-COA SYNTHETASE
3030 2nwc A 3.02 ANGSTROM CRYSTAL STRUCTURE OF WILD-TYPE APO GROEL IN
3031 2o15 MYCOBACTERIUM TUBERCULOSIS EPSP SYNTHASE AFTER PARTIAL PRODUCTS
3032 2o3j STRUCTURE OF CAENORHABDITIS ELEGANS UDP-GLUCOSE DEHYDROGENASE
3033 2oej CRYSTAL STRUCTURE OF A RUBISCO-LIKE PROTEIN FROM GEOBACILLUS
3034 2oek CRYSTAL STRUCTURE OF A RUBISCO-LIKE PROTEIN FROM GEOBACILLUS
3035 2oel CRYSTAL STRUCTURE OF A RUBISCO-LIKE PROTEIN FROM GEOBACILLUS
3036 2oem CRYSTAL STRUCTURE OF A RUBISCO-LIKE PROTEIN FROM GEOBACILLUS
3037 2oqh CRYSTAL STRUCTURE OF AN ISOMERASE FROM STREPTOMYCES COELICOLOR A3(2)
3038 2osw ENDO-GLYCOCERAMIDASE II FROM RHODOCOCCUS SP.
3039 2oyk ENDO-GLYCOCERAMIDASE II FROM RHODOCOCCUS SP.: CELLOBIOSE-LIKE
3040 2oyl ENDO-GLYCOCERAMIDASE II FROM RHODOCOCCUS SP.: CELLOBIOSE-LIKE
3041 * 2pan CRYSTAL STRUCTURE OF E. COLI GLYOXYLATE CARBOLIGASE
3042 2pua CRYSTAL STRUCTURE OF THE LACI FAMILY MEMBER, PURR, BOUND TO DNA: MINOR
3043 2pub CRYSTAL STRUCTURE OF THE LACI FAMILY MEMBER, PURR, BOUND TO DNA: MINOR
3044 2puc CRYSTAL STRUCTURE OF THE LACI FAMILY MEMBER, PURR, BOUND TO
3045 2pud CRYSTAL STRUCTURE OF THE LACI FAMILY MEMBER, PURR, BOUND TO DNA: MINOR
3046 2q1x CRYSTAL STRUCTURE OF CELL DIVISION PROTEIN FTSZ FROM MYCOBACTERIUM
3047 2q1y CRYSTAL STRUCTURE OF CELL DIVISION PROTEIN FTSZ FROM MYCOBACTERIUM
3048 * 2q5o X-RAY STRUCTURE OF PHENYLPYRUVATE DECARBOXYLASE IN COMPLEX WITH 3-
3049 2qo3 CRYSTAL STRUCTURE OF [KS3][AT3] DIDOMAIN FROM MODULE 3 OF 6-
3050 2qq6 CRYSTAL STRUCTURE OF MANDELATE RACEMASE/MUCONATE
3051 2qw1 GLUCOSE/GALACTOSE BINDING PROTEIN BOUND TO 3-O-METHYL D-GLUCOSE
3052 2r79 CRYSTAL STRUCTURE OF A PERIPLASMIC HEME BINDING PROTEIN FROM
3053 2rb9 CRYSTAL STRUCTURE OF E.COLI HYPE
3054 2rdd X-RAY CRYSTAL STRUCTURE OF ACRB IN COMPLEX WITH A NOVEL
3055 2scu A DETAILED DESCRIPTION OF THE STRUCTURE OF SUCCINYL-COA
3056 2tpt STRUCTURAL AND THEORETICAL STUDIES SUGGEST DOMAIN MOVEMENT PRODUCES AN
3057 * 2uz1 1.65 ANGSTROM STRUCTURE OF BENZALDEHYDE LYASE COMPLEXED
3058 * 2uza CRYSTAL STRUCTURE OF THE FREE RADICAL INTERMEDIATE OF
3059 2v63 CRYSTAL STRUCTURE OF RUBISCO FROM CHLAMYDOMONAS REINHARDTII
3060 2v67 CRYSTAL STRUCTURE OF CHLAMYDOMONAS REINHARDTII RUBISCO WITH
3061 2v68 CRYSTAL STRUCTURE OF CHLAMYDOMONAS REINHARDTII RUBISCO WITH LARGE-
3062 2v69 CRYSTAL STRUCTURE OF CHLAMYDOMONAS REINHARDTII RUBISCO WITH A LARGE-
3063 2v6a CRYSTAL STRUCTURE OF CHLAMYDOMONAS REINHARDTII RUBISCO WITH
3064 2vdc THE 9.5 A RESOLUTION STRUCTURE OF GLUTAMATE SYNTHASE FROM CRYO-
3065 2vdh CRYSTAL STRUCTURE OF CHLAMYDOMONAS REINHARDTII RUBISCO WITH
3066 2vdi CRYSTAL STRUCTURE OF CHLAMYDOMONAS REINHARDTII RUBISCO WITH A LARGE-
3067 2vqj STRUCTURE OF HDAC4 CATALYTIC DOMAIN BOUND TO A
3068 2vqm STRUCTURE OF HDAC4 CATALYTIC DOMAIN BOUND TO A HYDROXAMIC
3069 2vqo STRUCTURE OF HDAC4 CATALYTIC DOMAIN WITH A GAIN-OF-FUNCTION
3070 2vqq STRUCTURE OF HDAC4 CATALYTIC DOMAIN (A DOUBLE CYSTEINE-TO-
3071 2vsn STRUCTURE AND TOPOLOGICAL ARRANGEMENT OF AN O-GLCNAC
3072 2vyc CRYSTAL STRUCTURE OF ACID INDUCED ARGININE DECARBOXYLASE
3073 2vz8 CRYSTAL STRUCTURE OF MAMMALIAN FATTY ACID SYNTHASE

3074 2vz9 CRYSTAL STRUCTURE OF MAMMALIAN FATTY ACID SYNTHASE IN
3075 2w1b THE STRUCTURE OF THE EFFLUX PUMP ACRB IN COMPLEX WITH BILE
3076 2wpg SUCROSE HYDROLASE
3077 2wvw CRYO-EM STRUCTURE OF THE RBCL-RBCX COMPLEX
3078 2x8u SPHINGOMONAS WITTICHII SERINE PALMITOYLTRANSFERASE
3079 2xyl CELLULOMONAS FIMI XYLANASE/CELLULASE COMPLEXED WITH 2-DEOXY-
3080 2y0c BCEC MUTATION Y10S
3081 2y0d BCEC MUTATION Y10K
3082 2y0e BCEC AND THE FINAL STEP OF UGDS REACTION
3083 2yey CRYSTAL STRUCTURE OF THE ALLOSTERIC-DEFECTIVE CHAPERONIN
3084 2yfn GALACTOSIDASE DOMAIN OF ALPHA-GALACTOSIDASE-SUCROSE KINASE,
3085 2yfo GALACTOSIDASE DOMAIN OF ALPHA-GALACTOSIDASE-SUCROSE KINASE,
3086 2yri CRYSTAL STRUCTURE OF ALANINE-PYRUVATE AMINOTRANSFERASE WITH 2-
3087 2yrr HYPOTHETICAL ALANINE AMINOTRANSFERASE (TTH0173) FROM THERMUS
3088 2yz7 X-RAY ANALYSES OF 3-HYDROXYBUTYRATE DEHYDROGENASE FROM
3089 2zsh STRUCTURAL BASIS OF GIBBERELLIN(GA3)-INDUCED DELLA
3090 2zsi STRUCTURAL BASIS OF GIBBERELLIN(GA4)-INDUCED DELLA
3091 2ztl CLOSED CONFORMATION OF D-3-HYDROXYBUTYRATE DEHYDROGENASE COMPLEXED
3092 2ztm T190S MUTANT OF D-3-HYDROXYBUTYRATE DEHYDROGENASE
3093 2ztu T190A MUTANT OF D-3-HYDROXYBUTYRATE DEHYDROGENASE COMPLEXED
3094 2ztv THE BINARY COMPLEX OF D-3-HYDROXYBUTYRATE DEHYDROGENASE WITH NAD+
3095 2zul CRYSTAL STRUCTURE OF THERMUS THERMOPHILUS 16S RRNA METHYLTRANSFERASE
3096 2zwv CRYSTAL STRUCTURE OF THERMUS THERMOPHILUS 16S RRNA
3097 3aob STRUCTURES OF THE MULTIDRUG EXPORTER ACRB REVEAL A PROXIMAL MULTISITE
3098 3aoc STRUCTURES OF THE MULTIDRUG EXPORTER ACRB REVEAL A PROXIMAL MULTISITE
3099 3cmt MECHANISM OF HOMOLOGOUS RECOMBINATION FROM THE RECA-SSDNA/DSDNA
3100 3cmu MECHANISM OF HOMOLOGOUS RECOMBINATION FROM THE RECA-SSDNA/DSDNA
3101 3cmv MECHANISM OF HOMOLOGOUS RECOMBINATION FROM THE RECA-SSDNA/DSDNA
3102 3cmw MECHANISM OF HOMOLOGOUS RECOMBINATION FROM THE RECA-SSDNA/DSDNA
3103 3cmx MECHANISM OF HOMOLOGOUS RECOMBINATION FROM THE RECA-SSDNA/DSDNA
3104 3cny CRYSTAL STRUCTURE OF A PUTATIVE INOSITOL CATABOLISM PROTEIN IOLE
3105 3cr9 CRYSTAL STRUCTURE OF THE COMPLEX OF LACTOFERRIN WITH 6-
3106 3cuf CELLULOMONAS FIMI XYLANASE/CELLULASE CEX (CF XYN10A) IN COMPLEX WITH
3107 3cug CELLULOMONAS FIMI XYLANASE/CELLULASE CEX (CF XYN10A) IN COMPLEX WITH
3108 3cuh CELLULOMONAS FIMI XYLANASE/CELLULASE CEX (CF XYN10A) IN COMPLEX WITH
3109 3cui CELLULOMONAS FIMI XYLANASE/CELLULASE CEX (CF XYN10A) IN COMPLEX WITH
3110 3cuj CELLULOMONAS FIMI XYLANASE/CELLULASE CEX (CF XYN10A) IN COMPLEX WITH
3111 3cze CRYSTAL STRUCTURE ANALYSIS OF SUCROSE HYDROLASE (SUH)- TRIS COMPLEX
3112 3czl CRYSTAL STRUCTURE ANALYSIS OF SUCROSE HYDROLASE(SUH) E322Q-GLUCOSE
3113 * 3d7k CRYSTAL STRUCTURE OF BENZALDEHYDE LYASE IN COMPLEX WITH THE
3114 3d9b SYMMETRIC STRUCTURE OF E. COLI ACRB
3115 3dc2 CRYSTAL STRUCTURE OF SERINE BOUND D-3-PHOSPHOGLYCERATE DEHYDROGENASE
3116 3ddn CRYSTAL STRUCTURE OF HYDROXYPYRUVIC ACID PHOSPHATE BOUND D-3-
3117 3dme CRYSTAL STRUCTURE OF CONSERVED EXPORTED PROTEIN FROM BORDETELLA
3118 3dmf T. THERMOPHILUS 16S RRNA N2 G1207 METHYLTRANSFERASE (RSMC) IN COMPLEX
3119 3dmg T. THERMOPHILUS 16S RRNA N2 G1207 METHYLTRANSFERASE (RSMC) IN COMPLEX
3120 3dmh T. THERMOPHILUS 16S RRNA N2 G1207 METHYLTRANSFERASE (RSMC) IN COMPLEX
3121 3e6g CRYSTAL STRUCTURE OF XOMETC, A CYSTATHIONINE C-LYASE-LIKE
3122 * 3e9y ARABIDOPSIS THALIANA ACETOHYDROXYACID SYNTHASE IN COMPLEX WITH
3123 * 3ea4 ARABIDOPSIS THALIANA ACETOHYDROXYACID SYNTHASE IN COMPLEX WITH
3124 3ffn CRYSTAL STRUCTURE OF CALCIUM-FREE HUMAN GELSOLIN
3125 * 3flm CRYSTAL STRUCTURE OF MEND FROM E.COLI
3126 3ga5 X-RAY STRUCTURE OF GLUCOSE/GALACTOSE RECEPTOR FROM
3127 3gbp STRUCTURE OF THE PERIPLASMIC GLUCOSE/GALACTOSE RECEPTOR OF SALMONELLA
3128 3gcm CRYSTAL STRUCTURE OF E. COLI POLYNUCLEOTIDE PHOSPHORYLASE
3129 3gll CRYSTAL STRUCTURE OF POLYNUCLEOTIDE PHOSPHORYLASE (PNPASE)
3130 3go7 CRYSTAL STRUCTURE OF M. TUBERCULOSIS RIBOKINASE (RV2436) IN COMPLEX
3131 3gsi CRYSTAL STRUCTURE OF D552A DIMETHYLGLYCINE OXIDASE MUTANT OF
3132 3h1c CRYSTAL STRUCTURE OF POLYNUCLEOTIDE PHOSPHORYLASE (PNPASE)
3133 3h8e LOW PH NATIVE STRUCTURE OF LEUCINE AMINOPEPTIDASE FROM PSEUDOMONAS
3134 3h8f HIGH PH NATIVE STRUCTURE OF LEUCINE AMINOPEPTIDASE FROM PSEUDOMONAS
3135 3h8g BESTATIN COMPLEX STRUCTURE OF LEUCINE AMINOPEPTIDASE FROM PSEUDOMONAS
3136 3haz CRYSTAL STRUCTURE OF BIFUNCTIONAL PROLINE UTILIZATION A
3137 3hjr CRYSTAL STRUCTURE OF SERINE PROTEASE OF AEROMONAS SOBRIA
3138 * 3hww CRYSTAL STRUCTURE OF MENAQUINONE SYNTHESIS PROTEIN MEND FROM E. COLI
3139 * 3hwx CRYSTAL STRUCTURE OF MENAQUINONE SYNTHESIS PROTEIN MEND FROM E. COLI
3140 3i01 NATIVE STRUCTURE OF BIFUNCTIONAL CARBON MONOXIDE DEHYDROGENASE/ACETYL-
3141 3i8b THE CRYSTAL STRUCTURE OF XYLULOSE KINASE FROM
3142 * 3iae STRUCTURE OF BENZALDEHYDE LYASE A28S MUTANT WITH BENZOYLPHOSPHONATE
3143 * 3iaf STRUCTURE OF BENZALDEHYDE LYASE A28S MUTANT WITH MONOMETHYL
3144 3ihg CRYSTAL STRUCTURE OF A TERNARY COMPLEX OF AKLAVINONE-11 HYDROXYLASE
3145 3ij3 1.8 ANGSTROM RESOLUTION CRYSTAL STRUCTURE OF CYTOSOL AMINOPEPTIDASE
3146 3itl CRYSTAL STRUCTURE OF PSEUDOMONAS STUTZERI L-RHAMNOSE
3147 3ito CRYSTAL STRUCTURE OF PSEUDOMONAS STUTZERI L-RHAMNOSE

3148 3itx MN2+ BOUND FORM OF PSEUDOMONAS STUTZERI L-RHAMNOSE ISOMERASE
3149 3ity METAL-FREE FORM OF PSEUDOMONAS STUTZERI L-RHAMNOSE ISOMERASE
3150 3iui ZN2+-BOUND FORM OF PSEUDOMONAS STUTZERI L-RHAMNOSE ISOMERASE
3151 3iyl ATOMIC CRYOEM STRUCTURE OF A NONENVELOPED VIRUS SUGGESTS HOW MEMBRANE
3152 3jru CRYSTAL STRUCTURE OF LEUCYL AMINOPEPTIDASE (PEPA) FROM XOO0834,
3153 3k9d CRYSTAL STRUCTURE OF PROBABLE ALDEHYDE DEHYDROGENASE FROM LISTERIA
3154 3kdr THE CRYSTAL STRUCTURE OF A HK97 FAMILY PHAGE PORTAL PROTEIN FROM
3155 3kg2 AMPA SUBTYPE IONOTROPIC GLUTAMATE RECEPTOR IN COMPLEX WITH COMPETITIVE
3156 3ktt ATOMIC MODEL OF BOVINE TRIC CCT2(BETA) SUBUNIT DERIVED FROM A 4.0
3157 3l76 CRYSTAL STRUCTURE OF ASPARTATE KINASE FROM SYNECHOCYSTIS
3158 3lml CRYSTAL STRUCTURE OF THE SHEATH TAIL PROTEIN LIN1278 FROM LISTERIA
3159 3m0v CRYSTAL STRUCTURE OF PSEUDOMONAS STUTZERI L-RHAMNOSE ISOMERASE MUTANT
3160 3m0x CRYSTAL STRUCTURE OF PSEUDOMONAS STUTZERI L-RHAMNOSE ISOMERASE MUTANT
3161 3m0y CRYSTAL STRUCTURE OF PSEUDOMONAS STUTZERI L-RHAMNOSE ISOMERASE MUTANT
3162 3mga 2.4 ANGSTROM CRYSTAL STRUCTURE OF FERRIC ENTEROBACTIN ESTERASE (FES)
3163 3mog CRYSTAL STRUCTURE OF 3-HYDROXYBUTYRYL-COA DEHYDROGENASE FROM
3164 3my7 THE CRYSTAL STRUCTURE OF THE ACDH DOMAIN OF AN ALCOHOL DEHYDROGENASE
3165 3n2b 1.8 ANGSTROM RESOLUTION CRYSTAL STRUCTURE OF DIAMINOPIMELATE
3166 3ndz THE STRUCTURE OF THE CATALYTIC AND CARBOHYDRATE BINDING DOMAIN OF
3167 3noc DESIGNED ANKYRIN REPEAT PROTEIN (DARPIN) BINDERS TO ACRB: PLASTICITY
3168 3o9p THE STRUCTURE OF THE ESCHERICHIA COLI MUREIN TRIPEPTIDE BINDING
3169 3oqm STRUCTURE OF CCPA-HPR-SER46P-ACKA2 COMPLEX
3170 3oqn STRUCTURE OF CCPA-HPR-SER46-P-GNTR-DOWN CRE
3171 3oqo CCPA-HPR-SER46P-SYN CRE
3172 3pgj 2.49 ANGSTROM RESOLUTION CRYSTAL STRUCTURE OF SHIKIMATE 5-
3173 3pgy SERINE HYDROXYMETHYLTRANSFERASE FROM STAPHYLOCOCCUS AUREUS, S95P
3174 3ptz ROLE OF PACKING DEFECTS IN THE EVOLUTION OF ALLOSTERY AND INDUCED FIT
3175 3q9l THE STRUCTURE OF THE DIMERIC E.COLI MIND-ATP COMPLEX
3176 3qfw CRYSTAL STRUCTURE OF RUBISCO-LIKE PROTEIN FROM RHODOPSEUDOMONAS
3177 3qhx CRYSTAL STRUCTURE OF CYSTATHIONINE GAMMA-SYNTHASE METB (CGS) FROM
3178 3qi6 CRYSTAL STRUCTURE OF CYSTATHIONINE GAMMA-SYNTHASE METB (CGS) FROM
3179 3qm3 1.85 ANGSTROM RESOLUTION CRYSTAL STRUCTURE OF FRUCTOSE-BISPHOSPHATE
3180 3qp9 THE STRUCTURE OF A C2-TYPE KETOREDUCTASE FROM A MODULAR POLYKETIDE
3181 3r0x CRYSTAL STRUCTURE OF SELENOMETHIONINE INCORPORATED APO D-SERINE
3182 3r4t CRYSTAL STRUCTURE OF 4-AMINOBUTYRATE AMINOTRANSFERASE GABT FROM
3183 3r9i 2.6A RESOLUTION STRUCTURE OF MIND COMPLEXED WITH MINE (12-31) PEPTIDE
3184 3rcy CRYSTAL STRUCTURE OF MANDELATE RACEMASE/MUCONATE LACTONIZING ENZYME-
3185 3rg6 CRYSTAL STRUCTURE OF A CHAPERONE-BOUND ASSEMBLY INTERMEDIATE OF FORM I
3186 3rr1 CRYSTAL STRUCTURE OF ENOLASE PRK14017 (TARGET EFI-500653) FROM
3187 3sef 2.4 ANGSTROM RESOLUTION CRYSTAL STRUCTURE OF SHIKIMATE 5-DEHYDROGENASE
3188 3sx2 CRYSTAL STRUCTURE OF A PUTATIVE 3-KETOACYL-(ACYL-CARRIER-PROTEIN)
3189 3t4w THE CRYSTAL STRUCTURE OF MANDELATE RACEMASE/MUCONATE LACTONIZING
3190 3t51 CRYSTAL STRUCTURES OF THE PRE-EXTRUSION AND EXTRUSION STATES OF THE
3191 3t56 CRYSTAL STRUCTURE OF THE PRE-EXTRUSION STATE OF THE CUSBA ADAPTOR-
3192 3t5t VALL FROM STREPTOMYCES HYGROSCOPICUS IN APO FORM
3193 3t7d VALL FROM STREPTOMYCES HYGROSCOPICUS IN COMPLEX WITH TREHALOSE
3194 3t81 CRYSTAL STRUCTURE OF DIIRON ADENINE DEAMINASE
3195 3t8l CRYSTAL STRUCTURE OF ADENINE DEAMINASE WITH MN/FE
3196 3tdk CRYSTAL STRUCTURE OF HUMAN UDP-GLUCOSE DEHYDROGENASE
3197 3tl3 STRUCTURE OF A SHORT-CHAIN TYPE DEHYDROGENASE/REDUCTASE FROM
3198 3tlj CRYSTAL STRUCTURE OF TRM14 FROM PYROCOCCUS FURIOSUS IN COMPLEX WITH S-
3199 3tm4 CRYSTAL STRUCTURE OF TRM14 FROM PYROCOCCUS FURIOSUS IN COMPLEX WITH S-
3200 3tm5 CRYSTAL STRUCTURE OF TRM14 FROM PYROCOCCUS FURIOSUS IN COMPLEX WITH
3201 3tma CRYSTAL STRUCTURE OF TRMN FROM THERMUS THERMOPHILUS
3202 3tte CRYSTAL STRUCTURE OF ENOLASE BRADO_4202 (TARGET EFI-501651) FROM
3203 3tyh CRYSTAL STRUCTURE OF OXO-CUPPER CLUSTERS BINDING TO FERRIC BINDING
3204 3u0b CRYSTAL STRUCTURE OF AN OXIDOREDUCTASE FROM MYCOBACTERIUM SMEGMATIS
3205 3umm FORMYLGLYCINAMIDE RIBONUCLEOTIDE AMIDOTRANSFERASE FROM SALMONELLA
3206 3uwx CRYSTAL STRUCTURE OF UVRA-UVRB COMPLEX
3207 3v4z D-ALANINE--D-ALANINE LIGASE FROM YERSINIA PESTIS
3208 3vdm CRYSTAL STRUCTURE OF VLDE, THE PSEUDO-GLYCOSYLTRANSFERASE WHICH
3209 3vdn CRYSTAL STRUCTURE OF VLDE, THE PSEUDO-GLYCOSYLTRANSFERASE, IN COMPLEX
3210 3vk2 CRYSTAL STRUCTURE OF L-METHIONINE GAMMA-LYASE FROM PSEUDOMONAS PUTIDA
3211 3vkg X-RAY STRUCTURE OF AN MTBD TRUNCATION MUTANT OF DYNEIN MOTOR DOMAIN
3212 3vtf STRUCTURE OF A UDP-GLUCOSE DEHYDROGENASE FROM THE HYPERTHERMOPHILIC
3213 3w9h STRUCTURAL BASIS FOR THE INHIBITION OF BACTERIAL MULTIDRUG EXPORTERS
3214 3wt4 STRUCTURAL AND KINETIC BASES FOR THE METAL PREFERENCE OF THE M18
3215 3wy7 CRYSTAL STRUCTURE OF MYCOBACTERIUM SMEGMATIS 7-KETO-8-AMINOPELARGONIC
3216 3zqj MYCOBACTERIUM TUBERCULOSIS UVRA
3217 3zxw STRUCTURE OF ACTIVATED RUBISCO FROM THERMOSYNECHOCOCCUS ELONGATUS
3218 3zz1 CRYSTAL STRUCTURE OF A GLYCOSIDE HYDROLASE FAMILY 3 BETA-GLUCOSIDASE,
3219 4a0o SYMMETRY-FREE CRYO-EM MAP OF TRIC IN THE NUCLEOTIDE-FREE (APO) STATE
3220 4a0v MODEL REFINED AGAINST THE SYMMETRY-FREE CRYO-EM MAP OF TRIC-AMP-PNP
3221 4a0w MODEL BUILT AGAINST SYMMETRY-FREE CRYO-EM MAP OF TRIC-ADP-ALFX

| | | |
|---|---|---|
| 4a13 | MODEL REFINED AGAINST SYMMETRY-FREE CRYO-EM MAP OF TRIC-ADP | |
| 4a21 | STRUCTURE OF MYCOBACTERIUM TUBERCULOSIS FRUCTOSE 1,6- | |
| 4aaq | ATP-TRIGGERED MOLECULAR MECHANICS OF THE CHAPERONIN GROEL | |
| 4aar | ATP-TRIGGERED MOLECULAR MECHANICS OF THE CHAPERONIN GROEL | |
| 4aau | ATP-TRIGGERED MOLECULAR MECHANICS OF THE CHAPERONIN GROEL | |
| 4ai6 | DYNEIN MOTOR DOMAIN - ADP COMPLEX | |
| 4akg | DYNEIN MOTOR DOMAIN - ATP COMPLEX | |
| 4akh | DYNEIN MOTOR DOMAIN - AMPPNP COMPLEX | |
| 4aki | DYNEIN MOTOR DOMAIN - LUAC DERIVATIVE | |
| 4atp | STRUCTURE OF GABA-TRANSAMINASE A1R958 FROM ARTHROBACTER AURESCENS IN | |
| 4atq | GABA-TRANSAMINASE A1R958 IN COMPLEX WITH EXTERNAL ALDIMINE PLP-GABA | |
| 4avn | THERMOBIFIDA FUSCA CELLOBIOHYDROLASE CEL6B CATALYTIC MUTANT | |
| 4avo | THERMOBIFIDA FUSCA CELLOBIOHYDROLASE CEL6B CATALYTIC MUTANT D274A | |
| 4ayg | LACTOBACILLUS REUTERI N-TERMINALLY TRUNCATED GLUCANSUCRASE GTF180 IN | |
| 4ayo | STRUCTURE OF THE GH47 PROCESSING ALPHA-1,2-MANNOSIDASE FROM | |
| 4ayp | STRUCTURE OF THE GH47 PROCESSING ALPHA-1,2-MANNOSIDASE FROM | |
| 4ayq | STRUCTURE OF THE GH47 PROCESSING ALPHA-1,2-MANNOSIDASE FROM | |
| 4ayr | STRUCTURE OF THE GH47 PROCESSING ALPHA-1,2-MANNOSIDASE FROM | |
| 4b2t | THE CRYSTAL STRUCTURES OF THE EUKARYOTIC CHAPERONIN CCT REVEAL ITS | |
| 4b3f | CRYSTAL STRUCTURE OF IGHMBP2 HELICASE | |
| 4b3g | CRYSTAL STRUCTURE OF IGHMBP2 HELICASE IN COMPLEX WITH RNA | |
| 4b3h | CRYSTAL STRUCTURE OF MYCOBACTERIUM TUBERCULOSIS FATTY ACID | |
| 4b3i | CRYSTAL STRUCTURE OF MYCOBACTERIUM TUBERCULOSIS FATTY ACID | |
| 4b3j | CRYSTAL STRUCTURE OF MYCOBACTERIUM TUBERCULOSIS FATTY ACID | |
| 4b4f | THERMOBIFIDA FUSCA CEL6B(E3) CO-CRYSTALLIZED WITH CELLOBIOSE | |
| 4b4h | THERMOBIFIDA FUSCA CELLOBIOHYDROLASE CEL6B(E3) CATALYTIC DOMAIN | |
| 4bju | GENETIC AND STRUCTURAL VALIDATION OF ASPERGILLUS FUMIGATUS | |
| 4blp | P4 PROTEIN FROM BACTERIOPHAGE PHI13 | |
| 4c48 | CRYSTAL STRUCTURE OF ACRB-ACRZ COMPLEX | |
| 4cbt | DESIGN, SYNTHESIS, AND BIOLOGICAL EVALUATION OF POTENT AND SELECTIVE | |
| 4cdi | CRYSTAL STRUCTURE OF ACRB-ACRZ COMPLEX | |
| 4dcs | CRYSTAL STRUCTURE OF B. SUBTILIS ENGA IN COMPLEX WITH SULFATE ION AND | |
| 4dct | CRYSTAL STRUCTURE OF B. SUBTILIS ENGA IN COMPLEX WITH HALF-OCCUPACY | |
| 4dcu | CRYSTAL STRUCTURE OF B. SUBTILIS ENGA IN COMPLEX WITH GDP | |
| 4del | ACTIVE SITE LOOP DYNAMICS OF A CLASS IIA FRUCTOSE 1,6-BISPHOSPHATE | |
| 4dop | CRYSTAL STRUCTURE OF THE CUSBA HEAVY-METAL EFFLUX COMPLEX FROM | |
| 4dpp | THE STRUCTURE OF DIHYDRODIPICOLINATE SYNTHASE 2 FROM ARABIDOPSIS | |
| 4dpq | THE STRUCTURE OF DIHYDRODIPICOLINATE SYNTHASE 2 FROM ARABIDOPSIS | |
| 4dqx | CRYSTAL STRUCTURE OF A SHORT CHAIN DEHYDROGENASE FROM RHIZOBIUM ETLI | |
| 4dx5 | TRANSPORT OF DRUGS BY THE MULTIDRUG TRANSPORTER ACRB INVOLVES AN | |
| 4dx6 | TRANSPORT OF DRUGS BY THE MULTIDRUG TRANSPORTER ACRB INVOLVES AN | |
| 4dx7 | TRANSPORT OF DRUGS BY THE MULTIDRUG TRANSPORTER ACRB INVOLVES AN | |
| 4dye | CRYSTAL STRUCTURE OF AN ENOLASE (PUTATIVE SUGAR ISOMERASE, TARGET EFI- | |
| 4e4t | CRYSTAL STRUCTURE OF PHOSPHORIBOSYLAMINOIMIDAZOLE CARBOXYLASE, ATPASE | |
| 4e6e | CRYSTAL STRUCTURE OF A PUTATIVE CELL DIVISION PROTEIN FTSZ (TFU_1113) | |
| 4e6m | CRYSTAL STRUCTURE OF PUTATIVE DEHYDRATASE PROTEIN FROM SALMONELLA | |
| 4e6p | CRYSTAL STRUCTURE OF A PROBABLE SORBITOL DEHYDROGENASE (TARGET PSI- | |
| 4ezb | CRYSTAL STRUCTURE OF THE CONSERVED HYPOTHETICAL PROTEIN FROM | |
| 4f0k | UNACTIVATED RUBISCO WITH MAGNESIUM AND CARBON DIOXIDE BOUND | |
| 4f0m | UNACTIVATED RUBISCO WITH MAGNESIUM AND A WATER MOLECULE BOUND | |
| 4f4c | THE CRYSTAL STRUCTURE OF THE MULTI-DRUG TRANSPORTER | |
| 4f4f | X-RAY CRYSTAL STRUCTURE OF PLP BOUND THREONINE SYNTHASE FROM BRUCELLA | |
| 4f96 | CRYSTAL STRUCTURE OF VLDE, THE PSEUDO-GLYCOSYLTRANSFERASE, IN COMPLEX | |
| 4f97 | CRYSTAL STRUCTURE OF VLDE, THE PSEUDO-GLYCOSYLTRANSFERASE, IN COMPLEX | |
| 4f9f | CRYSTAL STRUCTURE OF VLDE, THE PSEUDO-GLYCOSYLTRANSFERASE, IN COMPLEX | |
| * | 4fee | HIGH-RESOLUTION STRUCTURE OF PYRUVATE OXIDASE IN COMPLEX WITH REACTION |
| * | 4feg | HIGH-RESOLUTION STRUCTURE OF PYRUVATE OXIDASE IN COMPLEX WITH REACTION |
| 4fsx | CRYSTAL STRUCTURE OF SE-SUBSTITUTED ZEA MAYS ZMET2 IN COMPLEX WITH SAH | |
| 4ft2 | CRYSTAL STRUCTURE OF ZEA MAYS ZMET2 IN COMPLEX H3(1-15)K9ME2 PEPTIDE | |
| 4ft4 | CRYSTAL STRUCTURE OF ZEA MAYS ZMET2 IN COMPLEX H3(1-32)K9ME2 PEPTIDE | |
| 4ggm | STRUCTURE OF LPXI | |
| 4gji | CRYSTAL STRUCTURE OF PSEUDOMONAS STUTZERI L-RHAMNOSE ISOMERASE MUTANT | |
| 4gjj | CRYSTAL STRUCTURE OF PSEUDOMONAS STUTZERI L-RHAMNOSE ISOMERASE MUTANT | |
| 4hel | CRYSTAL STRUCTURE ANALYSIS OF APO-GROEL STRUCTURE | |
| 4i3g | CRYSTAL STRUCTURE OF DESR, A BETA-GLUCOSIDASE FROM STREPTOMYCES | |
| 4ixs | NATIVE STRUCTURE OF XOMETC AT PH 5.2 | |
| 4ixz | NATIVE STRUCTURE OF CYSTATHIONINE GAMMA LYASE (XOMETC) FROM | |
| 4iy7 | CRYSTAL STRUCTURE OF CYSTATHIONINE GAMMA LYASE (XOMETC) FROM | |
| 4iyo | CRYSTAL STRUCTURE OF CYSTATHIONINE GAMMA LYASE FROM XANTHOMONAS ORYZAE | |
| 4izo | CRYSTAL STRUCTURE OF KINASE PHOSPHORIBOSYLAMINOIMIDAZOLE CARBOXYLASE, | |
| 4jk1 | X-RAY CRYSTAL STRUCTURE OF ESCHERICHIA COLI SIGMA70 HOLOENZYME IN | |
| 4k03 | CRYSTAL STRUCTURE OF DROSOPHILA CRYPROCHROME | |
| 4k0j | X-RAY CRYSTAL STRUCTURE OF A HEAVY METAL EFFLUX PUMP, CRYSTAL FORM I | |
| 4k28 | 2.15 ANGSTROM RESOLUTION CRYSTAL STRUCTURE OF A SHIKIMATE | |

*       4kgd   HIGH-RESOLUTION CRYSTAL STRUCTURE OF PYRUVATE OXIDASE FROM L.
4kn4   X-RAY CRYSTAL STRUCTURE OF THE ESCHERICHIA COLI RNA POLYMERASE IN
4kn7   X-RAY CRYSTAL STRUCTURE OF THE ESCHERICHIA COLI RNA POLYMERASE IN
4kq9   CRYSTAL STRUCTURE OF PERIPLASMIC RIBOSE ABC TRANSPORTER FROM
4ksi   CRYSTAL STRUCTURE ANALYSIS OF THE ACIDIC LEUCINE AMINOPEPTIDASE OF
4kwe   GDP-BOUND, DOUBLE-STRANDED, CURVED FTSZ PROTOFILAMENT STRUCTURE
4l9y   CRYSTAL STRUCTURE OF RHODOBACTER SPHAEROIDES MALYL-COA LYASE IN
4l9z   CRYSTAL STRUCTURE OF RHODOBACTER SPHAEROIDES MALYL-COA LYASE IN
4lf1   HEXAMERIC FORM II RUBISCO FROM RHODOPSEUDOMONAS PALUSTRIS, ACTIVATED
4mex   CRYSTAL STRUCTURE OF ESCHERICHIA COLI RNA POLYMERASE IN COMPLEX WITH
4mey   CRYSTAL STRUCTURE OF ESCHERICHIA COLI RNA POLYMERASE HOLOENZYME
4n0q   CRYSTAL STRUCTURE OF AN ABC TRANSPORTER, SUBSTRATE-BINDING PROTEIN
4n44   CRYSTAL STRUCTURE OF OXIDIZED FORM OF THIOLASE FROM CLOSTRIDIUM
4ndz   STRUCTURE OF MALTOSE BINDING PROTEIN FUSION TO 2-O-SULFOTRANSFERASE
4ni5   CRYSTAL STRUCTURE OF A SHORT CHAIN DEHYDROGENASE FROM BRUCELLA SUIS
4njq   STRUCTURAL AND KINETIC BASES FOR THE METAL PREFERENCE OF THE M18
4njr   STRUCTURAL AND KINETIC BASES FOR THE METAL PREFERENCE OF THE M18
4ns4   CRYSTAL STRUCTURE OF COLD-ACTIVE ESTARASE FROM PSYCHROBACTER
4o89   CRYSTAL STRUCTURE OF RTCA, THE RNA 3'-TERMINAL PHOSPHATE CYCLASE FROM
4o8j   CRYSTAL STRUCTURE OF RTCA, THE RNA 3'-TERMINAL PHOSPHATE CYCLASE FROM
4oby   CRYSTAL STRUCTURE OF E.COLI ARGINYL-TRNA SYNTHETASE AND LIGAND BINDING
4oid   STRUCTURAL AND KINETIC BASES FOR THE METAL PREFERENCE OF THE M18
4oiw   STRUCTURAL AND KINETIC BASES FOR THE METAL PREFERENCE OF THE M18
4omu   CRYSTAL STRUCTURE OF SHIKIMATE DEHYDROGENASE (AROE) FROM PSEUDOMONAS
4ope   STREPTOMCYES ALBUS JA3453 OXAZOLOMYCIN KETOSYNTHASE DOMAIN OZMH KS7
4pj1   CRYSTAL STRUCTURE OF THE HUMAN MITOCHONDRIAL CHAPERONIN SYMMETRICAL
4pvf   CRYSTAL STRUCTURE OF HOMO SAPIENS HOLO SERINE HYDROXYMETHYLTRANSFERASE
4q0c   3.1 A RESOLUTION CRYSTAL STRUCTURE OF THE B. PERTUSSIS BVGS
4q31   THE CRYSTAL STRUCTURE OF CYSTATHIONE GAMMA LYASE (CALE6) FROM
4q71   CRYSTAL STRUCTURE OF BRADYRHIZOBIUM JAPONICUM PROLINE UTILIZATION A
4q72   CRYSTAL STRUCTURE OF BRADYRHIZOBIUM JAPONICUM PROLINE UTILIZATION A
4q73   CRYSTAL STRUCTURE OF BRADYRHIZOBIUM JAPONICUM PROLINE UTILIZATION A
4qav   THE STRUCTURE OF BETA-KETOACYL -(ACYL CARRIER PROTEIN) SYNTHASE II
*       4qpz   CRYSTAL STRUCTURE OF THE FORMOLASE FLS_V2 IN SPACE GROUP P 21
*       4qq8   CRYSTAL STRUCTURE OF THE FORMOLASE FLS IN SPACE GROUP P 43 21 2
4qwv   A PBP-LIKE PROTEIN BUILT FROM FRAGMENTS OF DIFFERENT FOLDS
4qyr   STREPTOMYCES PLATENSIS ISOMIGRASTATIN KETOSYNTHASE DOMAIN MGSE KS3
4r5d   CRYSTAL STRUCTURE OF COMPUTATIONAL DESIGNED LEUCINE RICH REPEATS
4rjt   CRYSTAL STRUCTURE OF UNLIGANDED, FULL LENGTH HUGDH AT PH 7.0
4rkq   CRYSTAL STRUCTURE OF LACI FAMILY TRANSCRIPTIONAL REGULATOR FROM
4rkr   CRYSTAL STRUCTURE OF LACI FAMILY TRANSCRIPTIONAL REGULATOR FROM
4rl6   CRYSTAL STRUCTURE OF THE Q04L03_STRP2 PROTEIN FROM STREPTOCOCCUS
4rnj   PAMORA PHOSPHODIESTERASE DOMAIN, APO FORM
4roq   CRYSTAL STRUCTURE OF MALYL-COA LYASE FROM METHYLOBACTERIUM EXTORQUENS
4rs3   CRYSTAL STRUCTURE OF CARBOHYDRATE TRANSPORTER A0QYB3 FROM
4rsm   CRYSTAL STRUCTURE OF CARBOHYDRATE TRANSPORTER MSMEG_3599 FROM
4rub   A CRYSTAL FORM OF RIBULOSE-1,5-BISPHOSPHATE CARBOXYLASE(SLASH)
4rwe   THE CRYSTAL STRUCTURE OF A SUGAR-BINDING TRANSPORT PROTEIN FROM
4ry9   CRYSTAL STRUCTURE OF CARBOHYDRATE TRANSPORTER SOLUTE BINDING PROTEIN
4tkt   STREPTOMYCES PLATENSIS ISOMIGRASTATIN KETOSYNTHASE DOMAIN MGSF KS6
4toz   MPPA PERIPLASMIC MUREIN TRIPEPTIDE BINDING PROTEIN, UNLIGANDED OPEN
4u1w   FULL LENGTH GLUA2-KAINATE-(R,R)-2B COMPLEX CRYSTAL FORM A
4u2p   FULL-LENGTH AMPA SUBTYPE IONOTROPIC GLUTAMATE RECEPTOR GLUA2 IN THE
4u2q   FULL-LENGTH AMPA SUBTYPE IONOTROPIC GLUTAMATE RECEPTOR GLUA2 IN
4u5c   CRYSTAL STRUCTURE OF GLUA2, CON-IKOT-IKOT SNAIL TOXIN, PARTIAL AGONIST
4u8v   COUPLING OF REMOTE ALTERNATING-ACCESS TRANSPORT MECHANISMS FOR PROTONS
4u8y   COUPLING OF REMOTE ALTERNATING-ACCESS TRANSPORT MECHANISMS FOR PROTONS
4u95   COUPLING OF REMOTE ALTERNATING-ACCESS TRANSPORT MECHANISMS FOR PROTONS
4u96   COUPLING OF REMOTE ALTERNATING-ACCESS TRANSPORT MECHANISMS FOR PROTONS
4wbt   CRYSTAL STRUCTURE OF HISTIDINOL-PHOSPHATE AMINOTRANSFERASE FROM
4wgl   CRYSTAL STRUCTURE OF A GROEL D83A/R197A DOUBLE MUTANT
4wky   STREPTOMCYES ALBUS JA3453 OXAZOLOMYCIN KETOSYNTHASE DOMAIN OZMN KS2
4wsc   CRYSTAL STRUCTURE OF A GROELK105A MUTANT
4x8f   VIBRIO CHOLERAE O395 RIBOKINASE IN APO FORM
4xd7   STRUCTURE OF THERMOPHILIC F1-ATPASE INHIBITED BY EPSILON SUBUNIT
4xkj   A NOVEL D-LACTATE DEHYDROGENASE FROM SPOROLACTOBACILLUS SP
4xl2   CRYSTAL STRUCTURE OF OXIDIZED FORM OF THIOLASE FROM CLOSTRIDIUM
4xq2   ENSEMBLE REFINEMENT OF CYSTATHIONE GAMMA LYASE (CALE6) D7G FROM
4xqk   ATP-DEPENDENT TYPE ISP RESTRICTION-MODIFICATION ENZYME LLABIII BOUND
4xx0   COA BOUND TO PIG GTP-SPECIFIC SUCCINYL-COA SYNTHETASE
4yln   E. COLI TRANSCRIPTION INITIATION COMPLEX - 17-BP SPACER AND 4-NT RNA
4ylo   E. COLI TRANSCRIPTION INITIATION COMPLEX - 16-BP SPACER AND 4-NT RNA
4ylp   E. COLI TRANSCRIPTION INITIATION COMPLEX - 16-BP SPACER AND 5-NT RNA
4yv7   CRYSTAL STRUCTURE OF AN ABC TRANSPORTER SOLUTE BINDING PROTEIN

4z6k   ALCOHOL DEHYDROGENASE FROM THE ANTARCTIC PSYCHROPHILE MORAXELLA SP.
4zdn   STREPTOMYCES PLATENSIS ISOMIGRASTATIN KETOSYNTHASE DOMAIN MGSF KS4
4zit   CRYSTAL STRUCTURE OF ACRB IN P21 SPACE GROUP
4ziv   CRYSTAL STRUCTURE OF ACRB TRIPLE MUTANT IN P21 SPACE GROUP
4ziw   CRYSTAL STRUCTURE OF ACRB DELETION MUTANT IN P21 SPACE GROUP
4zjl   CRYSTAL STRUCTURE OF ACRB IN COMPLEX WITH ANTIBIOTIC IN P21 SPACE
4zjo   CRYSTAL STRUCTURE OF ACRB TRIPLE MUTANT IN COMPLEX WITH ANTIBIOTIC IN
4zjp   STRUCTURE OF AN ABC-TRANSPORTER SOLUTE BINDING PROTEIN (SBP_IPR025997)
4zjq   CRYSTAL STRUCTURE OF ACRB DELETION MUTANT IN COMPLEX WITH ANTIBIOTIC
4zlj   CRYSTAL STRUCTURE OF TRANSPORTER ACRB
4zll   CRYSTAL STRUCTURE OF TRANSPORTER ACRB TRIPLE MUTANT
4zln   CRYSTAL STRUCTURE OF TRANSPORTER ACRB DELETION MUTANT
4zqi   CRYSTAL STRUCTURE OF APO D-ALANINE-D-ALANINE LIGASE(DDL) FROM YERSINIA
4ztx   NEUROSPORA CRASSA COBALAMIN-INDEPENDENT METHIONINE SYNTHASE COMPLEXED
4zty   NEUROSPORA CRASSA COBALAMIN-INDEPENDENT METHIONINE SYNTHASE COMPLEXED
5ac3   CRYSTAL STRUCTURE OF PAM12A
5b63   CRYSTAL STRUCTURES OF E.COLI ARGINYL-TRNA SYNTHETASE (ARGRS) IN
5bpf   CRYSTAL STRUCTURE OF ADP COMPLEXED D-ALANINE-D-ALANINE LIGASE(DDL)
5bph   CRYSTAL STRUCTURE OF AMP COMPLEXED D-ALANINE-D-ALANINE LIGASE(DDL)
5c1p   CRYSTAL STRUCTURE OF ADP AND D-ALANYL-D-ALANINE COMPLEXED D-ALANINE-D-
5c2g   GWS1B RUBISCO: FORM II RUBISCO DERIVED FROM UNCULTIVATED
5ch8   CRYSTAL STRUCTURE OF MDLA N225Q MUTANT FORM PENICILLIUM CYCLOPIUM
*   5d6r   ACETOLACTATE SYNTHASE FROM KLEBSIELLA PNEUMONIAE IN COMPLEX WITH
5d8n   TOMATO LEUCINE AMINOPEPTIDASE MUTANT - K354E
5dkv   CRYSTAL STRUCTURE OF AN ABC TRANSPORTER SOLUTE BINDING PROTEIN FROM
5dvy   2.95 ANGSTROM CRYSTAL STRUCTURE OF THE DIMERIC FORM OF PENICILLIN
*   5dx6   ACETOLACTATE SYNTHASE FROM KLEBSIELLA PNEUMONIAE SOAKED WITH BETA-
5edu   CRYSTAL STRUCTURE OF HUMAN HISTONE DEACETYLASE 6 CATALYTIC DOMAIN 2 IN
5eef   CRYSTAL STRUCTURE OF DANIO RERIO HISTONE DEACETYLASE 6 CATALYTIC
5ehk   CRYSTAL STRUCTURE OF TRNA DEPENDENT LANTIBIOTIC DEHYDRATASE MIBB FROM
*   5ej4   ECMEND-THDP-MN2+ COMPLEX SOAKED WITH 2-KETOGLUTARATE FOR 15 MIN
*   5ej5   ECMEND-THDP-MN2+ COMPLEX SOAKED WITH 2-KETOGLUTARATE FOR 1.5 H
*   5ej6   ECMEND-THDP-MN2+ COMPLEX SOAKED WITH 2-KETOGLUTARATE FOR 2MIN THEN
*   5ej7   ECMEND-THDP-MN2+ COMPLEX SOAKED WITH 2-KETOGLUTARATE FOR 21 S
*   5ej8   ECMEND-THDP-MN2+ COMPLEX STRUCTURE SOAKED WITH 2-KETOGLUTARATE FOR 2
*   5ej9   ECMEND-THDP-MN2+ COMPLEX SOAKED WITH 2-KETOGLUTARATE FOR 2 MIN AND
*   5eja   ECMEND-THDP-MN2+ COMPLEX SOAKED WITH 2-KETOGLUTARATE FOR 2 MIN AND
*   5ejm   THDP-MN2+ COMPLEX OF R413A VARIANT OF ECMEND SOAKED WITH 2-
5en5   APO STRUCTURE OF BACTERIAL EFFLUX PUMP.
5eno   MBX2319 BOUND STRUCTURE OF BACTERIAL EFFLUX PUMP.
5enp   MBX2931 BOUND STRUCTURE OF BACTERIAL EFFLUX PUMP.
5enq   MBX3132 BOUND STRUCTURE OF BACTERIAL EFFLUX PUMP.
5enr   MBX3135 BOUND STRUCTURE OF BACTERIAL EFFLUX PUMP.
5ens   RHODAMINE BOUND STRUCTURE OF BACTERIAL EFFLUX PUMP.
5ent   MINOCYCLINE BOUND STRUCTURE OF BACTERIAL EFFLUX PUMP.
5ezk   RNA POLYMERASE MODEL PLACED BY MOLECULAR REPLACEMENT INTO X-RAY
5f7s   CYCLOALTERNAN-DEGRADING ENZYME FROM TRUEPERELLA PYOGENES
5f9c   CRYSTAL STRUCTURE OF THE G121R MUTANT OF HUMAN PHOSPHOGLUCOMUTASE 1
5fac   ALANINE RACEMASE FROM STREPTOMYCES COELICOLOR A3(2)
5fag   ALANINE RACEMASE FROM STREPTOMYCES COELICOLOR A3(2) WITH BOUND
5faj   ALANINE RACEMASE FROM STREPTOMYCES COELICOLOR A3(2) IN COMPLEX WITH D-
5g0g   CRYSTAL STRUCTURE OF DANIO RERIO HDAC6 CD1 IN COMPLEX WITH
5g0i   CRYSTAL STRUCTURE OF DANIO RERIO HDAC6 CD1 AND CD2 (LINKER
5g0j   CRYSTAL STRUCTURE OF DANIO RERIO HDAC6 CD1 AND CD2 (LINKER
5g0x   PSEUDOMONAS AERUGINOSA HDAH BOUND TO ACETATE.
5g0y   PSEUDOMONAS AERUGINOSA HDAH UNLIGANDED.
5g10   PSEUDOMONAS AERUGINOSA HDAH BOUND TO 9,9,9 TRIFLUORO-8,8-DIHYDROY-N-
5g11   PSEUDOMONAS AERUGINOSA HDAH BOUND TO PFSAHA.
5g12   PSEUDOMONAS AERUGINOSA HDAH (Y313F) UNLIGANDED.
5g13   PSEUDOMONAS AERUGINOSA HDAH (H143A) UNLIGANDED.
5gzi   CYCLODEAMINASE_PA
5gzj   CYCLODEAMINASE_PA
5gzl   CYCLODEAMINASE_PA
5gzm   CYCLODEAMINASE_PA
5hao   STRUCTURE FUNCTION STUDIES OF R. PALUSTRIS RUBISCO (M331A MUTANT;
5hko   CRYSTAL STRUCTURE OF ABC TRANSPORTER SOLUTE BINDING PROTEIN MSMEG_3598
5hsh   CRYSTAL STRUCTURE OF THE G291R MUTANT OF HUMAN PHOSPHOGLUCOMUTASE 1
5hxa   CRYSTAL STRUCTURE OF AN UDP-FORMING ALPHA, ALPHA-TERHALOSE-PHOSPHATE
5hzg   THE CRYSTAL STRUCTURE OF THE STRIGOLACTONE-INDUCED ATD14-D3-ASK1
5i1f   CRYSTAL STRUCTURE OF UTP-GLUCOSE-1-PHOSPHATE URIDYLYLTRANSFERASE FROM
5i2h   CRYSTAL STRUCTURE OF O-METHYLTRANSFERASE FAMILY 2 PROTEIN PLIM_1147
5i47   CRYSTAL STRUCTURE OF RIMK DOMAIN PROTEIN ATP-GRASP FROM SPHAEROBACTER
5iky   APO STRUCTURE OF OBC1, A BIFUNCTIONAL ENZYME FOR QUORUM SENSING-
5ikz   GLYCEROL BOUND STRUCTURE OF OBC1, A BIFUNCTIONAL ENZYME FOR QUORUM

3444 5ipl SIGMAS-TRANSCRIPTION INITIATION COMPLEX WITH 4-NT NASCENT RNA
3445 5ipm SIGMAS-TRANSCRIPTION INITIATION COMPLEX WITH 4-NT NASCENT RNA
3446 5ipn SIGMAS-TRANSCRIPTION INITIATION COMPLEX WITH 4-NT NASCENT RNA
3447 5iu0 RUBISCO FROM ARABIDOPSIS THALIANA
3448 5iz4 CRYSTAL STRUCTURE OF A PUTATIVE SHORT-CHAIN DEHYDROGENASE/REDUCTASE
3449 5j78 CRYSTAL STRUCTURE OF AN ACETYLATING ALDEHYDE DEHYDROGENASE FROM
3450 5j7i CRYSTAL STRUCTURE OF A GEOBACILLUS THERMOGLUCOSIDASIUS ACETYLATING
3451 5ja1 ENTF, A TERMINAL NONRIBOSOMAL PEPTIDE SYNTHETASE MODULE BOUND TO THE
3452 5ja2 ENTF, A TERMINAL NONRIBOSOMAL PEPTIDE SYNTHETASE MODULE BOUND TO THE
3453 5jgf CRYSTAL STRUCTURE OF MAPE1
3454 5jh9 CRYSTAL STRUCTURE OF PRAPE1
3455 5jmn FUSIDIC ACID BOUND ACRB
3456 5jx2 CRYSTAL STRUCTURE OF MGLB-2 (TP0684) FROM TREPONEMA PALLIDUM
3457 * 5k2o CRYSTAL STRUCTURE OF ARABIDOPSIS THALIANA ACETOHYDROXYACID SYNTHASE IN
3458 * 5k3s CRYSTAL STRUCTURE OF ARABIDOPSIS THALIANA ACETOHYDROXYACID SYNTHASE IN
3459 * 5k6t CRYSTAL STRUCTURE OF ARABIDOPSIS THALIANA ACETOHYDROXYACID SYNTHASE IN
3460 5kf7 STRUCTURE OF PROLINE UTILIZATION A FROM SINORHIZOBIUM MELILOTI
3461 5koz STRUCTURE FUNCTION STUDIES OF R. PALUSTRIS RUBISCO (K192C MUTANT;
3462 5kpd MOUSE PGP 34 LINKER DELETED DOUBLE EQ MUTANT
3463 5kr3 DIRECTED EVOLUTION OF TRANSAMINASES BY ANCESTRAL RECONSTRUCTION. USING
3464 5kr4 DIRECTED EVOLUTION OF TRANSAMINASES BY ANCESTRAL RECONSTRUCTION. USING
3465 5kws CRYSTAL STRUCTURE OF GALACTOSE BINDING PROTEIN FROM YERSINIA PESTIS IN
3466 5l1b AMPA SUBTYPE IONOTROPIC GLUTAMATE RECEPTOR GLUA2 IN APO STATE
3467 5l1e AMPA SUBTYPE IONOTROPIC GLUTAMATE RECEPTOR GLUA2 IN COMPLEX WITH
3468 5l1f AMPA SUBTYPE IONOTROPIC GLUTAMATE RECEPTOR GLUA2 IN COMPLEX WITH
3469 5l1g AMPA SUBTYPE IONOTROPIC GLUTAMATE RECEPTOR GLUA2 IN COMPLEX WITH GYKI-
3470 5li3 CRYSTAL STRUCTURE OF HDAC-LIKE PROTEIN FROM P. AERUGINOSA IN COMPLEX
3471 5m6g CRYSTAL STRUCTURE GLUCAN 1,4-BETA-GLUCOSIDASE FROM SACCHAROPOLYSPORA
3472 5m7n CRYSTAL STRUCTURE OF NTRX FROM BRUCELLA ABORTUS IN COMPLEX WITH ATP
3473 5m7o CRYSTAL STRUCTURE OF NTRX FROM BRUCELLA ABORTUS PROCESSED WITH THE
3474 5m7p CRYSTAL STRUCTURE OF NTRX FROM BRUCELLA ABORTUS IN COMPLEX WITH ADP
3475 5mbs CRYSTAL STRUCTURE OF BACILLUS SUBTILIS ENGA IN SPACE GROUP P21
3476 5meh CRYSTAL STRUCTURE OF ALPHA-1,2-MANNOSIDASE FROM CAULOBACTER K31 STRAIN
3477 5msc STRUCTURE OF THE A DOMAIN OF CARBOXYLIC ACID REDUCTASE (CAR) FROM
3478 5msd STRUCTURE OF THE A DOMAIN OF CARBOXYLIC ACID REDUCTASE (CAR) FROM
3479 5n8o CRYO EM STRUCTURE OF THE CONJUGATIVE RELAXASE TRAI OF THE F/R1 PLASMID
3480 5nc5 CRYSTAL STRUCTURE OF ACRBZ IN COMPLEX WITH ANTIBIOTIC PUROMYCIN
3481 5ne5 CRYSTAL STRUCTURE OF FAMILY 47 ALPHA-1,2-MANNOSIDASE FROM CAULOBACTER
3482 5ng5 MULTI-DRUG EFFLUX; MEMBRANE TRANSPORT; RND SUPERFAMILY; DRUG
3483 5nug MOTOR DOMAINS FROM HUMAN CYTOPLASMIC DYNEIN-1 IN THE PHI-PARTICLE
3484 5o66 ASYMMETRIC ACRABZ-TOLC
3485 5oaw CRYSTAL STRUCTURE OF ASPERGILLUS FUMIGATUS N-ACETYLPHOSPHOGLUCOSAMINE
3486 5ocp THE PERIPLASMIC BINDING PROTEIN COMPONENT OF THE ARABINOSE ABC
3487 5odh HETERODISULFIDE REDUCTASE / [NIFE]-HYDROGENASE COMPLEX FROM
3488 5odq HETERODISULFIDE REDUCTASE / [NIFE]-HYDROGENASE COMPLEX FROM
3489 5odr HETERODISULFIDE REDUCTASE / [NIFE]-HYDROGENASE COMPLEX FROM
3490 5opw CRYSTAL STRUCTURE OF THE GROEL MUTANT A109C
3491 5t1e CRYSTAL STRUCTURE OF PHAEOSPAERIA NODRUM FRUCTOSYL PEPTIDE OXIDASE
3492 5t1f CRYSTAL STRUCTURE OF PHAEOSPAERIA NODRUM FRUCTOSYL PEPTIDE OXIDASE
3493 5td7 CRYSTAL STRUCTURE OF HISTONE DEACETYLASE 10
3494 5tr2 CRYSTAL STRUCTURE OF THE D263G MISSENSE VARIANT OF HUMAN PGM1
3495 5tt0 CRYSTAL STRUCTURE OF AN OXIDOREDUCTASE (SHORT CHAIN
3496 5tvg CRYSTAL STRUCTURE OF AN ALPHA,ALPHA-TREHALOSE-PHOSPHATE SYNTHASE (UDP-
3497 5u2w CRYSTAL STRUCTURE OF A SHORT CHAIN DEHYDROGENASE FROM BURKHOLDERIA
3498 5u9p CRYSTAL STRUCTURE OF A GLUCONATE 5-DEHYDROGENASE FROM BURKHOLDERIA
3499 5ugr MALYL-COA LYASE FROM METHYLOBACTERIUM EXTORQUENS
3500 5uof CRYSTAL STRUCTURE OF ALPHA,ALPHA-TREHALOSE 6-PHOSPHATE SYTHASE FROM
3501 5urm CRYSTAL STRUCTURE OF HUMAN BRR2 IN COMPLEX WITH T-1206548
3502 5ux5 STRUCTURE OF PROLINE UTILIZATION A (PUTA) FROM CORYNEBACTERIUM
3503 5uy8 CRYSTAL STRUCTURE OF AICARFT BOUND TO AN ANTIFOLATE
3504 5uyt CRYSTAL STRUCTURE OF ICE BINDING PROTEIN FROM AN ANTARCTIC BACTERIUM
3505 5uz0 CRYSTAL STRUCTURE OF AICARFT BOUND TO AN ANTIFOLATE
3506 5v5s MULTI-DRUG EFFLUX; MEMBRANE TRANSPORT; RND SUPERFAMILY; DRUG
3507 5v68 CRYSTAL STRUCTURE OF CELL DIVISION PROTEIN FTSZ FROM MYCOBACTERIUM
3508 5v7i CRYSTAL STRUCTURE OF HOMO SAPIENS SERINE HYDROXYMETHYLTRANSFERASE 2
3509 5vhw GLUA2-0XGSG1L BOUND TO ZK
3510 5vhx GLUA2-1XGSG1L BOUND TO ZK
3511 5vhy GLUA2-2XGSG1L BOUND TO ZK
3512 5vhz GLUA2-2XGSG1L BOUND TO L-QUISQUALATE
3513 5vjh CLOSED STATE CRYOEM RECONSTRUCTION OF HSP104:ATPYS AND FITC CASEIN
3514 5vp5 CRYSTAL STRUCTURE OF A 3-OXOACYL-ACYL-CARRIER PROTEIN REDUCTASE FABG4
3515 5vy9 S. CEREVISIAE HSP104:CASEIN COMPLEX, MIDDLE DOMAIN CONFORMATION
3516 5vya S. CEREVISIAE HSP104:CASEIN COMPLEX, EXTENDED CONFORMATION
3517 5vyw CRYSTAL STRUCTURE OF LACTOCOCCUS LACTIS PYRUVATE CARBOXYLASE

5vyz CRYSTAL STRUCTURE OF LACTOCOCCUS LACTIS PYRUVATE CARBOXYLASE IN
5vz0 CRYSTAL STRUCTURE OF LACTOCOCCUS LACTIS PYRUVATE CARBOXYLASE G746A
5w4x TRUNCATED HUGDH
* 5wdg ACETOLACTATE SYNTHASE FROM KLEBSIELLA PNEUMONIAE IN COMPLEX WITH A
5wek GLUA2 BOUND TO ANTAGONIST ZK AND GSG1L IN DIGITONIN, STATE 1
5wel GLUA2 BOUND TO ANTAGONIST ZK AND GSG1L IN DIGITONIN, STATE 2
5wem GLUA2 BOUND TO GSG1L IN DIGITONIN, STATE 1
5wen GLUA2 BOUND TO GSG1L IN DIGITONIN, STATE 2
* 5wj1 CRYSTAL STRUCTURE OF ARABIDOPSIS THALIANA ACETOHYDROXYACID SYNTHASE IN
5wqj CRYSTAL STRUCTURE OF 3-MERCAPTOPYRUVATE SULFURTRANSFERASE(3MST) IN
5wqk CRYSTAL STRUCTURE OF 3-MERCAPTOPYRUVATE SULFURTRANSFERASE(3MST) IN
5x2w CRYSTAL STRUCTURE OF PSEUDOMONAS PUTIDA METHIONINE GAMMA-LYASE WILD
5x2x CRYSTAL STRUCTURE OF PSEUDOMONAS PUTIDA METHIONINE GAMMA-LYASE WILD
5x2z CRYSTAL STRUCTURE OF PSEUDOMONAS PUTIDA METHIONINE GAMMA-LYASE C116H
5x30 CRYSTAL STRUCTURE OF PSEUDOMONAS PUTIDA METHIONINE GAMMA-LYASE C116H
5x3i KFLA1895 D451A MUTANT
5x3j KFLA1895 D451A MUTANT IN COMPLEX WITH CYCLOBIS-(1->6)-ALPHA-NIGEROSYL
5x3k KFLA1895 D451A MUTANT IN COMPLEX WITH ISOMALTOSE
5x3v STRUCTURE OF HUMAN SHMT2 PROTEIN MUTANT
5x7j CRYSTAL STRUCTURE OF THYMIDYLATE KINASE FROM THERMUS THERMOPHILUS HB8
5x86 CRYSTAL STRUCTURE OF TMP BOUND THYMIDYLATE KINASE FROM THERMUS
5x8a CRYSTAL STRUCTURE OF ATP BOUND THYMIDYLATE KINASE FROM THERMUS
5x8b CRYSTAL STRUCTURE OF ATP-TMP AND ADP BOUND THYMIDYLATE KINASE FROM
5x8c AMPPCP AND TMP BOUND CRYSTAL STRUCTURE OF THYMIDYLATE KINASE FROM
5x8j K16M MUTANT OF THERMUS THERMOPHILUS HB8 THYMIDYLATE KINASE
5x8k V158T MUTANT OF THERMUS THERMOPHILUS HB8 THYMIDYLATE KINASE
5x8v Y92H MUTANT OF THERMUS THERMOPHILUS HB8 THYMIDYLATE KINASE
5x98 Y162F MUTANT OF THERMUS THERMOPHILUS HB8 THYMIDYLATE KINASE
5x99 T18V MUTANT OF THERMUS THERMOPHILUS HB8 THYMIDYLATE KINASE
5xak CRYSTAL STRUCTURE (FORM II) OF THYMIDYLATE KINASE FROM THERMUS
5xal Y99F MUTANT OF THERMUS THERMOPHILUS HB8 THYMIDYLATE KINASE
5xog RNA POLYMERASE II ELONGATION COMPLEX BOUND WITH SPT5 KOW5 AND ELF1
5xt8 MAGNESIUM BOUND APO STRUCTURE OF THYMIDYLATE KINASE (FORM I) FROM
5yil HOISTING-LOOP IN BACTERIAL MULTIDRUG EXPORTER ACRB IS A HIGHLY
5yu0 STRUCTURAL BASIS FOR RECOGNITION OF L-LYSINE, L-ORNITHINE, AND L-2,4-
5yu1 STRUCTURAL BASIS FOR RECOGNITION OF L-LYSINE, L-ORNITHINE, AND L-2,4-
5yu3 STRUCTURAL BASIS FOR RECOGNITION OF L-LYSINE, L-ORNITHINE, AND L-2,4-
5yu4 STRUCTURAL BASIS FOR RECOGNITION OF L-LYSINE, L-ORNITHINE, AND L-2,4-
5yym CRYSTAL STRUCTURES OF E.COLI ARGINYL-TRNA SYNTHETASE (ARGRS) IN
5yyn CRYSTAL STRUCTURES OF E.COLI ARGINYL-TRNA SYNTHETASE (ARGRS) IN
* 5z2p THDP-MN2+ COMPLEX OF R413K VARIANT OF ECMEND SOAKED WITH 2-
* 5z2r THDP-MN2+ COMPLEX OF R395K VARIANT OF ECMEND SOAKED WITH 2-
* 5z2u THDP-MN2+ COMPLEX OF R395A VARIANT OF ECMEND SOAKED WITH 2-
5zax CRYSTAL STRUCTURE OF THYMIDYLATE KINASE IN COMPLEX WITH ADP, TDP AND
5zb0 CRYSTAL STRUCTURE OF THYMIDYLATE KINASE IN COMPLEX WITH ADP AND TDP
5zb4 CRYSTAL STRUCTURE OF THYMIDYLATE KINASE IN COMPLEX WITH ADP AND TMP
5zfm KETOREDUCTASE LBCR MUTANT - M6
5zi0 KETOREDUCTASE LBCR MUTANT - M8
5zue GTP-BOUND, DOUBLE-STRANDED, CURVED FTSZ PROTOFILAMENT STRUCTURE
5zvt STRUCTURE OF RNA POLYMERASE COMPLEX AND GENOME WITHIN A DSRNA VIRUS
6a3f LEVOGLUCOSAN DEHYDROGENASE, APO FORM
6a3g LEVOGLUCOSAN DEHYDROGENASE, COMPLEX WITH NADH
6a3i LEVOGLUCOSAN DEHYDROGENASE, COMPLEX WITH NADH AND LEVOGLUCOSAN
6a3j LEVOGLUCOSAN DEHYDROGENASE, COMPLEX WITH NADH AND L-SORBOSE
6acn STRUCTURE OF ACTIVATED ACONITASE. FORMATION OF THE (4FE-4S) CLUSTER IN
6ahc CRYO-EM STRUCTURE OF ALDEHYDE-ALCOHOL DEHYDROGENASE REVEALS A HIGH-
6alg CRYOEM STRUCTURE OF HK022 NUN - E.COLI RNA POLYMERASE ELONGATION
6azy CRYSTAL STRUCTURE OF HSP104 R328M/R757M MUTANT FROM CALCARISPORIELLA
6b5b CRYO-EM STRUCTURE OF THE NAIP5-NLRC4-FLAGELLIN INFLAMMASOME
6b6h THE CRYO-EM STRUCTURE OF A BACTERIAL CLASS I TRANSCRIPTION ACTIVATION
6b9u CRYSTAL STRUCTURE OF 3-KETOACYL-(ACYL-CARRIER-PROTEIN) REDUCTASE FROM
6bgc THE CRYSTAL STRUCTURE OF THE W145A VARIANT OF TPMGLB-2 (TP0684) WITH
6bgd THE CRYSTAL STRUCTURE OF THE W145A VARIANT OF TPMGLB-2 (TP0684) WITH
6bjs CRYOEM STRUCTURE OF E.COLI HIS PAUSE ELONGATION COMPLEX WITHOUT PAUSE
6bsn STRUCTURE OF PROLINE UTILIZATION A (PUTA) WITH PROLINE BOUND IN REMOTE
6c4j LIGAND BOUND FULL LENGTH HUGDH WITH A104L SUBSTITUTION
6c5a HUMAN UDP-GLUCOSE DEHYDROGENASE WITH UDP- GLC AND NADH BOUND
6c5z HUMAN UDP-GLUCOSE DEHYDROGENASE A225L SUBSTITUTUION WITH UDP-GLUCOSE
6c7n MONOCLINIC FORM OF MALIC ENZYME FROM SORGHUM AT 2 ANGSTROMS RESOLUTION
6c84 CRYSTAL STRUCTURE OF PBP5 FROM ENTEROCOCCUS FAECIUM
6c9u CRYSTAL STRUCTURE OF [KS3][AT3] DIDOMAIN FROM MODULE 3 OF 6-
6ca0 CRYO-EM STRUCTURE OF E. COLI RNAP SIGMA70 OPEN COMPLEX
* 6ciq PYRUVATE:FERREDOXIN OXIDOREDUCTASE FROM MOORELLA THERMOACETICA WITH
6clw CRYSTAL STRUCTURE OF TNMH

| | |
|---|---|
| 6clx | CRYSTAL STRUCTURE OF TNMH IN COMPLEX WITH SAM |
| 6cn1 | 2.75 ANGSTROM RESOLUTION CRYSTAL STRUCTURE OF UDP-N-ACETYLGLUCOSAMINE |
| 6csx | SINGLE PARTICLES CRYO-EM STRUCTURE OF ACRB D407A ASSOCIATED WITH LIPID |
| 6ct6 | CRYSTAL STRUCTURE OF LACTATE DEHYDROGENASE FROM EIMERIA MAXIMA WITH |
| 6d00 | CALCARISPORIELLA THERMOPHILA HSP104 |
| 6d6k | STRUCTURE OF POLYRIBONUCLEOTIDE NUCLEOTIDYLTRANSFERASE FROM |
| 6dem | CRYSTAL STRUCTURE OF CANDIDA ALBICANS ACETOHYDROXYACID SYNTHASE IN |
| 6den | CRYSTAL STRUCTURE OF CANDIDA ALBICANS ACETOHYDROXYACID SYNTHASE IN |
| 6deq | CRYSTAL STRUCTURE OF CANDIDA ALBICANS ACETOHYDROXYACID SYNTHASE IN |
| 6dju | MTB CLPB IN COMPLEX WITH ATPGAMMAS AND CASEIN, CONFORMER 1 |
| 6djv | MTB CLPB IN COMPLEX WITH ATPGAMMAS AND CASEIN, CONFORMER 2 |
| 6dk3 | HUMAN MITOCHONDRIAL SERINE HYDROXYMETHYLTRANSFERASE 2 |
| 6dlz | OPEN STATE GLUA2 IN COMPLEX WITH STZ AFTER MICELLE SIGNAL SUBTRACTION |
| 6dm0 | OPEN STATE GLUA2 IN COMPLEX WITH STZ AND BLOCKED BY IEM-1460, AFTER |
| 6dm1 | OPEN STATE GLUA2 IN COMPLEX WITH STZ AND BLOCKED BY NASPM, AFTER |
| 6e10 | PTEX CORE COMPLEX IN THE ENGAGED (EXTENDED) STATE |
| 6em8 | S.AUREUS CLPC RESTING STATE, C2 SYMMETRISED |
| 6eq0 | STRUCTURE OF THE PERIPLASMIC BINDING PROTEIN (PBP) MELB (ATU4661) IN |
| 6eqo | TRI-FUNCTIONAL PROPIONYL-COA SYNTHASE OF ERYTHROBACTER SP. NAP1 WITH |
| 6et9 | STRUCTURE OF THE ACETOACETYL-COA-THIOLASE/HMG-COA-SYNTHASE COMPLEX |
| 6f5d | TRYPANOSOMA BRUCEI F1-ATPASE |
| 6fij | STRUCTURE OF THE LOADING/CONDENSING REGION (SAT-KS-MAT) OF THE |
| 6fik | ACP2 CROSSLINKED TO THE KS OF THE LOADING/CONDENSING REGION OF THE |
| 6flq | CRYOEM STRUCTURE OF E.COLI RNA POLYMERASE PAUSED ELONGATION COMPLEX |
| 6g0k | CRYSTAL STRUCTURE OF ENTEROCOCCUS FAECIUM D63R PENICILLIN-BINDING |
| 6gav | EXTREMELY 'OPEN' CLAMP STRUCTURE OF DNA GYRASE: ROLE OF THE |
| 6gg2 | THE STRUCTURE OF FSQB FROM ASPERGILLUS FUMIGATUS, A FLAVOENZYME OF THE |
| 6gym | STRUCTURE OF A YEAST CLOSED COMPLEX WITH DISTORTED DNA (CCDIST) |
| 6haf | PYRUVATE OXIDASE VARIANT E59Q FROM L. PLANTARUM IN COMPLEX WITH |
| 6hb0 | CRYSTAL STRUCTURE OF MSMEG_1712 FROM MYCOBACTERIUM SMEGMATIS |
| 6hbc | STRUCTURE OF THE REPEAT UNIT IN THE NETWORK FORMED BY CCMM AND RUBISCO |
| 6hbd | CRYSTAL STRUCTURE OF MSMEG_1712 FROM MYCOBACTERIUM SMEGMATIS IN |
| 6hbm | CRYSTAL STRUCTURE OF MSMEG_1712 FROM MYCOBACTERIUM SMEGMATIS IN |
| 6hyh | CRYSTAL STRUCTURE OF MSMEG_1712 FROM MYCOBACTERIUM SMEGMATIS IN |
| 6ier | APO STRUCTURE OF A BETA-GLUCOSIDASE 1317 |
| 6ii2 | CRYSTAL STRUCTURE OF ALPHA-BETA HYDROLASE (ABH) AND MAKES CATERPILLARS |
| 6imp | CRYSTAL STRUCTURE OF ALPHA-BETA HYDROLASE (ABH) FROM VIBRIO VULNIFICUS |
| 6io4 | SILVER-BOUND GLYCERALDEHYDE-3-PHOSPHATE DEHYDROGENASE A |
| 6io6 | SILVER-BOUND GLYCERALDEHYDE-3-PHOSPHATE DEHYDROGENASE A AT NON- |
| 6ioj | GLYCERALDEHYDE-3-PHOSPHATE DEHYDROGENASE A (APO-FORM) |
| 6ir9 | RNA POLYMERASE II ELONGATION COMPLEX BOUND WITH ELF1 AND SPT4/5, |
| 6j28 | CRYSTAL STRUCTURE OF THE BRANCHED-CHAIN POLYAMINE SYNTHASE C9 MUTEIN |
| 6j4w | RNA POLYMERASE II ELONGATION COMPLEX BOUND WITH ELF1 AND SPT4/5, |
| 6j4z | RNA POLYMERASE II ELONGATION COMPLEX BOUND WITH SPT4/5 AND FOREIGN |
| 6j50 | RNA POLYMERASE II ELONGATION COMPLEX BOUND WITH SPT4/5 AND FOREIGN |
| 6jco | CRYSTAL STRUCTURE OF CALCIUM FREE HUMAN GELSOLIN AMYLOID MUTANT D187N |
| 6jeg | CRYSTAL STRUCTURE OF CALCIUM FREE HUMAN GELSOLIN AMYLOID MUTANT G167R |
| 6jeh | CRYSTAL STRUCTURE OF CALCIUM FREE HUMAN GELSOLIN AMYLOID MUTANT D187Y |
| 6mka | CRYSTAL STRUCTURE OF PENICILLIN BINDING PROTEIN 5 (PBP5) FROM |
| 6mkf | CRYSTAL STRUCTURE OF PENICILLIN BINDING PROTEIN 5 (PBP5) FROM |
| 6mkg | CRYSTAL STRUCTURE OF PENICILLIN BINDING PROTEIN 5 (PBP5) FROM |
| 6n39 | CRYSTAL STRUCTURE OF AN DEPHOSPHO-COA KINASE COAE FROM MYCOBACTERIUM |
| 6n61 | ESCHERICHIA COLI RNA POLYMERASE SIGMA70-HOLOENZYME BOUND TO UPSTREAM |
| 6n62 | ESCHERICHIA COLI RNA POLYMERASE SIGMA70-HOLOENZYME BOUND TO UPSTREAM |
| 6n8e | CRYSTAL STRUCTURE OF HOLO-OBIF1, A FIVE DOMAIN NONRIBOSOMAL PEPTIDE |
| 6n8t | HSP104DWB CLOSED CONFORMATION |
| 6n8z | HSP104DWB EXTENDED CONFORMATION |
| 6njl | STRUCTURE OF A COMPLEX |
| 6njm | STRUCTURE OF A COMPLEX |
| 6njn | STRUCTURE OF A COMPLEX |
| 6non | STRUCTURE OF CYANTHECE APO MCDA |
| 6noo | STRUCTURE OF CYANOTHECE MCDA-AMPPNP COMPLEX |
| 6nor | CRYSTAL STRUCTURE OF GEND2 FROM GENTAMICIN A BIOSYNTHESIS IN COMPLEX |
| 6nr8 | HTRIC-HPFD CLASS6 |
| 6nr9 | HTRIC-HPFD CLASS5 |
| 6nra | HTRIC-HPFD CLASS1 (NO PFD) |
| 6nrb | HTRIC-HPFD CLASS2 |
| 6nrc | HTRIC-HPFD CLASS3 |
| 6nrd | HTRIC-HPFD CLASS4 |
| 6nzi | LOW RESOLUTION CRYSTAL STRUCTURE OF THE BACTERIAL MULTIDRUG EFFLUX |
| 6o4n | CRYSTAL STRUCTURE OF ENOLASE FROM CHLAMYDIA TRACHOMATIS |
| 6o9g | OPEN STATE GLUA2 IN COMPLEX WITH STZ AND BLOCKED BY AGTX-636, AFTER |
| 6oax | STRUCTURE OF THE HYPERACTIVE CLPB MUTANT K476C, BOUND TO CASEIN, PRE- |
| 6oay | STRUCTURE OF THE HYPERACTIVE CLPB MUTANT K476C, BOUND TO CASEIN, POST- |

| | |
|---|---|
| 6om8 | CAENORHABDITIS ELEGANS UDP-GLUCOSE DEHYDROGENASE IN COMPLEX WITH UDP- |
| 6or5 | FULL-LENGTH S. POMBE MDN1 IN THE PRESENCE OF AMPPNP (RING REGION) |
| 6orb | FULL-LENGTH S. POMBE MDN1 IN THE PRESENCE OF ATP AND RBIN-1 |
| 6pz9 | CRYO-EM STRUCTURE OF THE PANCREATIC BETA-CELL SUR1 BOUND TO ATP AND |
| 6q7i | GH3 EXO-BETA-XYLOSIDASE (XLND) |
| 6q7j | GH3 EXO-BETA-XYLOSIDASE (XLND) IN COMPLEX WITH XYLOBIOSE AZIRIDINE |
| 6qb8 | HUMAN CCT:MLST8 COMPLEX |
| 6qep | ENGBF DARPIN FUSION 4B H14 |
| 6qev | ENGBF DARPIN FUSION 4B B6 |
| 6qfk | ENGBF DARPIN FUSION 4B G10 |
| 6qfo | ENGBF DARPIN FUSION 9B 3G124 |
| 6qg9 | CRYSTAL STRUCTURE OF IDEONELLA SAKAIENSIS MHETASE |
| 6qga | CRYSTAL STRUCTURE OF IDEONELLA SAKAIENSIS MHETASE BOUND TO THE NON- |
| 6qgb | CRYSTAL STRUCTURE OF IDEONELLA SAKAIENSIS MHETASE BOUND TO BENZOIC |
| 6qs4 | TWO-STEP ACTIVATION MECHANISM OF THE CLPB DISAGGREGASE FOR SEQUENTIAL |
| 6qs6 | CLPB (DWB AND K476C MUTANT) BOUND TO CASEIN IN PRESENCE OF ATPGAMMAS - |
| 6qs7 | CLPB (DWB AND K476C MUTANT) BOUND TO CASEIN IN PRESENCE OF ATPGAMMAS - |
| 6qs8 | CLPB (DWB AND K476C MUTANT) BOUND TO CASEIN IN PRESENCE OF ATPGAMMAS - |
| 6qss | CRYSTAL STRUCTURE OF IGNICOCCUS ISLANDICUS MALATE DEHYDROGENASE CO- |
| 6qvg | HUMAN SHMT2 IN COMPLEX WITH LOMETREXOL |
| 6qvl | HUMAN SHMT2 IN COMPLEX WITH PEMETREXED |
| 6r8f | CRYO-EM STRUCTURE OF THE HUMAN BRISC-SHMT2 COMPLEX |
| 6rdc | CRYOEM STRUCTURE OF POLYTOMELLA F-ATP SYNTHASE, PRIMARY ROTARY STATE |
| 6rdg | CRYOEM STRUCTURE OF POLYTOMELLA F-ATP SYNTHASE, PRIMARY ROTARY STATE |
| 6rdq | CRYO-EM STRUCTURE OF POLYTOMELLA F-ATP SYNTHASE, ROTARY SUBSTATE 1D, |
| 6rdr | CRYO-EM STRUCTURE OF POLYTOMELLA F-ATP SYNTHASE, ROTARY SUBSTATE 1D, |
| 6rds | CRYO-EM STRUCTURE OF POLYTOMELLA F-ATP SYNTHASE, ROTARY SUBSTATE 1D, |
| 6rdz | CRYO-EM STRUCTURE OF POLYTOMELLA F-ATP SYNTHASE, ROTARY SUBSTATE 2A, |
| 6re0 | CRYO-EM STRUCTURE OF POLYTOMELLA F-ATP SYNTHASE, ROTARY SUBSTATE 2A, |
| 6re1 | CRYO-EM STRUCTURE OF POLYTOMELLA F-ATP SYNTHASE, ROTARY SUBSTATE 2A, |
| 6ree | CRYO-EM STRUCTURE OF POLYTOMELLA F-ATP SYNTHASE, ROTARY SUBSTATE 3B, |
| 6ref | CRYO-EM STRUCTURE OF POLYTOMELLA F-ATP SYNTHASE, ROTARY SUBSTATE 3B, |
| 6rep | CRYO-EM STRUCTURE OF POLYTOMELLA F-ATP SYNTHASE, PRIMARY ROTARY STATE |
| 6rer | CRYO-EM STRUCTURE OF POLYTOMELLA F-ATP SYNTHASE, ROTARY SUBSTATE 3B, |
| 6rh3 | CRYO-EM STRUCTURE OF E. COLI RNA POLYMERASE ELONGATION COMPLEX BOUND |
| 6rla | STRUCTURE OF THE DYNEIN-2 COMPLEX; MOTOR DOMAINS |
| 6rn2 | CLPB (DWB MUTANT) BOUND TO CASEIN IN PRESENCE OF ATPGAMMAS - STATE WT- |
| 6rn3 | CLPB (DWB MUTANT) BOUND TO CASEIN IN PRESENCE OF ATPGAMMAS - STATE WT- |
| 6rn4 | CLPB (DWB MUTANT) BOUND TO CASEIN IN PRESENCE OF ATPGAMMAS - STATE WT- |
| 6s6t | STRUCTURE OF AZOSPIRILLUM BRASILENSE GLUTAMATE SYNTHASE IN A4B3 |
| 6sc2 | STRUCTURE OF THE DYNEIN-2 COMPLEX; IFT-TRAIN BOUND MODEL |
| 6sh9 | ENGBF DARPIN FUSION 4B D12 |
| 1N5W | Crystal Structure of the Cu,Mo-CO Dehydrogenase (CODH); Oxidized form |
| 2FKP | The mutant G127C-T313C of Deinococcus Radiodurans N-acylamino acid racemase |
| 2GGG | The mutant A68C-D72C of Deinococcus Radiodurans N-acylamino acid racemase |
| 2GGH | The mutant A68C-D72C-NLQ of Deinococcus Radiodurans Nacylamino acid racemase |
| 2GGI | The mutant E149C-A182C of Deinococcus Radiodurans N-acylamino acid racemase |
| 2GGJ | The mutant Y218C of Deinococcus Radiodurans N-acylamino acid racemase |
| 2IU0 | crystal structures of transition state analogue inhibitors of inosine monophosphate cyclohydrolase |
| 2YEY | Crystal structure of the allosteric-defective chaperonin GroEL E434K mutant |
| 3K1Q | Backbone model of an aquareovirus virion by cryo-electron microscopy and bioinformatics |
| 3NDY | The structure of the catalytic and carbohydrate binding domain of endoglucanase D from Clostridium cellulovorans |
| 3ZQJ | Mycobacterium tuberculosis UvrA |
| 4E4T | Crystal structure of Phosphoribosylaminoimidazole carboxylase, ATPase subunit from Burkholderia ambifaria |
| 4IXS | Native structure of xometc at ph 5.2 |
| 4IXZ | Native structure of cystathionine gamma lyase (XometC) from xanthomonas oryzae pv. oryzae at pH 9.0 |
| 4K28 | 2.15 Angstrom resolution crystal structure of a shikimate dehydrogenase family protein from Pseudomonas putida KT2440 in complex with NAD+ |
| 4V43 | Structural and mechanistic basis for allostery in the bacterial chaperonin GroEL |
| 4V58 | Crystal structure of fatty acid synthase from thermomyces lanuginosus at 3.1 angstrom resolution |
| 4V8L | Cryo-EM Structure of the Mycobacterial Fatty Acid Synthase |
| 4V8V | Structure and conformational variability of the Mycobacterium tuberculosis fatty acid synthase multienzyme complex |
| 4V8W | Structure and conformational variability of the Mycobacterium tuberculosis fatty acid synthase multienzyme complex |
| 4XL4 | Crystal structure of thiolase from Clostridium acetobutylicum in complex with CoA |
| 4ZDN | Streptomyces platensis isomigrastatin ketosynthase domain MgsF KS4 |
| 5O66 | Asymmetric AcrABZ-TolC |
| 5W4X | Truncated hUGDH |
| 6ALH | CryoEM structure of E.coli RNA polymerase elongation complex |

3740 6DK3 HUMAN MITOCHONDRIAL SERINE HYDROXYMETHYLTRANSFERASe 2
3741 6O9G Open state GluA2 in complex with STZ and blocked by AgTx-636, after micelle signal subtraction
3742 6OR5 Full-length S. pombe Mdn1 in the presence of AMPPNP (ring region)
3743 6ORB Full-length S. pombe Mdn1 in the presence of ATP and Rbin-1
